# Snail-trematode dynamics in central Alberta wetlands: A longitudinal survey of infections and interactions

**DOI:** 10.1101/2025.05.26.656203

**Authors:** Brooke A. McPhail, Sara Tomusiak, Hannah Veinot, Neill Dodds, Patrick C. Hanington

## Abstract

Previous research has shown that host diversity and heterogeneity promote parasite abundance and heterogeneity by creating additional niches for parasites to occupy. Central Alberta, Canada, sees a diverse array of native and migratory species each year. A previous snail–trematode survey conducted at six lakes in central Alberta from 2013-2015 uncovered 79 trematode species. However, analyses suggested that additional species remained to be uncovered. To build on this baseline, we conducted further snail–trematode collections from 2019 to 2022 at eight reclaimed wetland sites in various stages of reclamation, along with one established lake in Alberta. Across the nine sites, we collected 22,397 snails, of which 1,981 were infected with digenean trematodes. We also documented broader biodiversity at these sites using traditional survey techniques. Through DNA barcoding, we identified 74 trematode species infecting five snail species. Among these were 23 trematode species not previously reported in central Alberta and nine putative novel species. In addition, we observed several previously unreported snail–trematode interactions. While trematode richness did not vary significantly with the wetland reclamation stage, host identity did influence richness: *Physa gyrina* hosted significantly more trematode species than *Planorbella trivolvis*. When combined with data from the earlier survey, sample completeness analyses indicate that we captured 100% of the dominant species and 99% of the typical species, but only 63% of the overall species diversity in central Alberta. These findings underscore that trematode diversity in central Alberta remains underestimated and highlight the continued value of long-term and host-inclusive sampling efforts.

## 1. Introduction

Parasites are important organisms that are often overlooked when considering biodiversity (Tompkins et al., 2001; Gómez & Nichols, 2013; Frainer et al., 2018), even though they constitute 50% of the organisms on Earth (Thomas et al., 2005; Poulin, 2014). Every animal in an aquatic ecosystem can be parasitized by flatworms known as digenean trematodes (Marcogliese, 2005). Trematodes are connected to other organisms in their environment through complex life cycles incorporating up to four hosts. The first intermediate host is almost always a species of snail. After this, trematodes can parasitize a second intermediate host (species of benthic invertebrates or fish) and/or a definitive host (birds, ungulates, rodents, etc.). Even if their host is no longer present in the area (e.g., migratory birds), trematodes act as a record of their presence (Morely & Lewis, 2007). The relationship between trematodes and their snail hosts is considered much more specific than the other hosts in their life cycles, likely due to their long history of co-evolution (Adamson & Caira, 1994; Esch & Fernandez, 1994). Snails host exponentially amplify infections and have been shown to maintain trematode species in waterbodies where resident avian hosts have been removed, leaving only migratory birds responsible for miracidial deposits.

The life cycle stages of trematodes are morphologically distinct and parasitize different host groups. To assess trematode diversity, researchers can collect animal feces from potential host species to search for trematode eggs, necropsy potential host organisms to search for metacercariae or adult worms, or collect snails to shed or dissect them in order to characterize larval stages. However, many trematode life cycles are not fully elucidated, and the elucidation of life cycles is not progressing at the same rate that new species are described (Blasco-Costa & Poulin, 2017). To fully capture trematode diversity, samples that include vouchered specimens, morphological information, DNA sequences, and relevant sample collection information are necessary (Brant et al., 2006). Utilizing molecular sequencing tools in conjunction with morphological assessments and making the sequences and corresponding metadata publicly available can facilitate easier connections between life cycle stages (Brant et al., 2006).

Employing DNA barcoding in conjunction with morphology can help us avoid misidentifications associated with phenotypic plasticity or differences due to geography (Whelan, 2021; Oyston et al., 2022). Researchers suggest that genetic identifications of trematodes are most useful when considering host species, geographic location, and the ecological context of host-parasite relationships (Blasco-Costa, Cutmore, et al., 2016). Repetitive sampling is recommended, and DNA should be sequenced from many samples to avoid misidentifying cryptic species (Poulin, 2011; Blasco-Costa, Cutmore, et al., 2016).

Diverse trematode communities reflect a host community that is also diverse (Hechinger & Lafferty, 2005; Hechinger et al., 2007). Throughout the last decade, the trematode community of central Alberta has been extensively studied through snail-trematode collections (Gordy et al., 2016, 2017, 2018; Gordy & Hanington, 2019; Gordy et al., 2020). Central Alberta is home to a species-rich trematode community of at least 79 unique species, which has been thoroughly characterized (Gordy et al., 2016; Gordy & Hanington, 2019). Based on morphological identifications alone, Gordy and Hanington (2019) predicted they had found 29 trematode species. Using DNA in conjunction with morphology, they discovered 79 trematode species (Gordy et al., 2016, 2017, 2018; Gordy & Hanington, 2019; Gordy et al., 2020). However, rarefaction analyses revealed that there remained trematode species to uncover in central Alberta (Gordy & Hanington, 2019).

Across four summers from 2019 to 2022, we conducted snail collections in central Alberta wetlands at varying stages of reclamation to continue capturing the composition of the trematode community and catalogue snail-trematode interactions, while also collecting data on potential vertebrate and invertebrate host species present at these sites. This data presents a unique opportunity to assess trematode familial relationships using phylogenetics, host-trematode relationships, the effect of wetland reclamation on trematode richness, trematode seasonality, and the completeness of trematode sampling undertaken in central Alberta.

## 2. Materials and methods

### 2.1 Study sites and sample collections

Eight reclaimed wetland sites located approximately 25km northeast of Edmonton, Alberta, Canada, were chosen based on information shared by an industrial partner that had mined these sites in previous years (Figure 1; Table 1). Collections were undertaken biweekly at the wetland sites from June to September 2019-2022, with 5-7 collection days at each site throughout the summer due to occasional industrial activity that prevented site access. Heritage Lake in Morinville, Alberta, Canada (Figure 1) was chosen as an additional comparison site.

**Figure 1.**
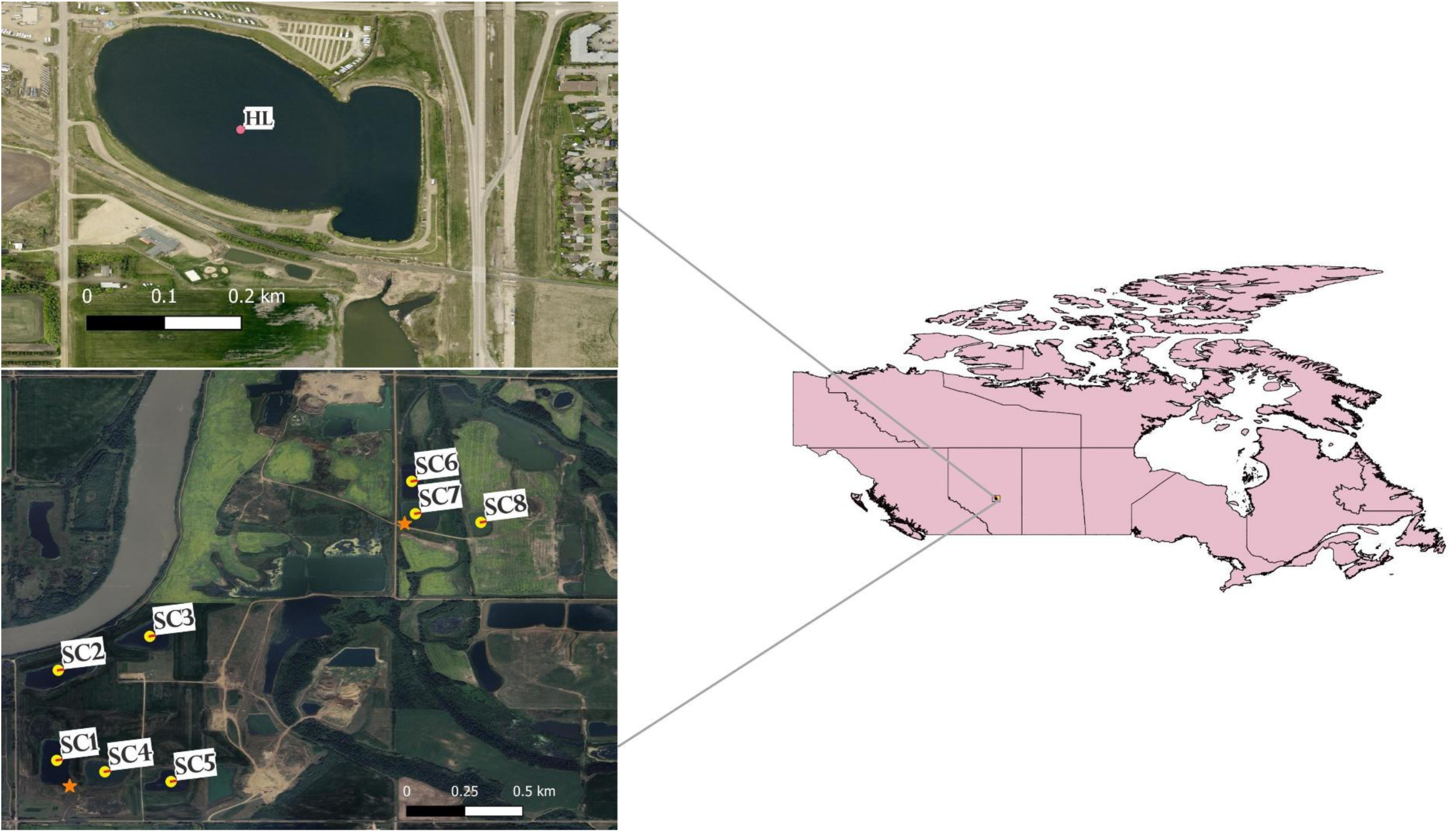
Collection sites visited during this study. HL = Heritage Lake, Morinville, Sturgeon County; SC = Strathcona County, Alberta. Orange stars indicate where the field cameras and recorders were placed. Made using QGIS (QGIS Development Team, 2024) with shapefiles from Statistics Canada (Statistics Canada, 2011).

**Table 1.** Information on collection sites, area is reported in hectares. Wetlands 0-5 years are considered early stage, 6-10 mid-stage, and 11+ years late-stage.

| Site | Latitude | Longitude | Area<br>(ha) | Age in 2019<br>(stage) |
| --- | --- | --- | --- | --- |
| SC1 | 53.63254 | -113.29271 | 3.66 | 5 (early) |
| SC2 | 53.63841 | -113.29261 | 3.44 | 8 (mid) |
| SC3 | 53.64062 | -113.28664 | 3.84 | 13 (late) |
| SC4 | 53.63179 | -113.28957 | 2.44 | 14 (late) |
| SC5 | 53.63116 | -113.28526 | 1.81 | 14 (late) |
| SC6 | 53.65072 | -113.26956 | 1.79 | 13 (late) |
| SC7 | 53.64861 | -113.26933 | 2.20 | 6 (mid) |
| SC8 | 53.64805 | -113.26506 | 1.40 | 2 (early) |
| Heritage Lake | 53.79940 | -113.66621 | 14.13 | Unknown |

Heritage Lake is located approximately 40km from Edmonton, AB. Snails were collected at Heritage Lake once in 2020 to determine if infected snails were present, followed by three collection days in 2021 and 2022 (one each in June, July, and August).

Snails were collected using a handheld sieve to separate the snails from the sediment and vegetation, or by gently handpicking the snails off rocks and vegetation where visible. The goal was to collect 200 snails from each site on every collection day. Collections were timed, so no more than half an hour was spent at each site. The snails were then brought to the laboratory and placed individually into well plates with artificial spring water (Ulmer, 1970). The well plates were placed under fluorescent lights that followed a 12-hour light-dark cycle. At 24 hours after collection, the wells were studied under a dissecting scope to determine if any cercariae had emerged. If cercariae were present, they were pipetted into a 1.5mL tube and preserved in ethanol (approximately 95% final concentration) for future molecular characterization. Upon cercarial emergence, cercariae were identified to the family level based on morphological characteristics from photos published by Gordy and colleagues (2016). Snails were identified based on morphology, using Aquatic Invertebrates of Alberta as a guide (Clifford, 1991).

### 2.2 Trematode species identification

DNA was extracted from the preserved cercariae using DNEasy Blood and Tissue kits from Qiagen (Catalogue No: 69504) using slightly modified protocols. The first four solutions (Buffer ATL, Proteinase K, Buffer AL, and ethanol) were added in 1/4^th^ of the quantity suggested by the manufacturer (Webster, 2009). The preserved samples then underwent Polymerase Chain Reaction (PCR), to amplify a region of either the cytochrome *c* oxidase subunit I (*COI*) barcoding gene (for most observed trematode families) or the nicotinamide adenine dinucleotide dehydrogenase subunit 1 (*nad1*) gene for trematodes belonging to the family Echinostomatidae.

Polymerase Chain Reactions (PCR) primers were chosen for each sample based on morphological identifications. Reactions were performed in 20μL samples using COIF15 and COIR15 for schistosome cercariae (Brant & Loker, 2009), NDJ11 and NDJ2a for echinostome cercariae (Kostadinova et al., 2003), and Dice 1F and Dice 11R primers for most non-schistosome cercariae (Van Steenkiste et al., 2015). After which, the samples underwent gene cleaning using a PCR Cleanup kit (Model: KTS1115, Truin Science Ltd., Edmonton, Canada) before they were sent for sequencing at either Macrogen (https://dna.macrogen.com/, Seoul, South Korea) or the Molecular Biology Service Unit at the University of Alberta (https://www.ualberta.ca/en/biological-sciences/services/mbsu/index.html). The sequences were trimmed using SnapGene® software (from Dotmatics; snapgene.com) and then edited and aligned using Geneious Prime 2024.0.7 (https://www.geneious.com).

The consensus sequences for each trematode sample were aligned and identified using either the Basic Local Alignment Search Tool (BLAST) in GenBank (Benson et al., 2013) or the Barcode of Life Database (Ratnasingham et al., 2024). A phylogenetic tree was run for each family in Geneious Prime using the MrBayes plugin (Huelsenbeck & Ronquist, 2001). The final alignment for each tree was exported into MEGA X (Tamura et al., 2021) to determine the best substitution model for each tree. Substitution models were chosen based on the lowest Bayesian information criterion (BIC) score. The parameters of the phylogenetic trees included four chains run at a chain length of 500 000 and the first 5 000 trees discarded as burn-in. Most trees used the *COI* gene. For the Echinostomatidae, both the *COI* and *nad1* genes were analyzed when available. Publicly available sequences from GenBank (Benson et al., 2013) or BOLD (Ratnasingham et al., 2024) were added to analyses to supplement sequences from the present study. GenBank accession numbers or BOLD Sequence IDs indicate such sequences. All trees were visualized using FigTree v1.4.5.

#### 2.2.1 Diplostomidae

Members of the Diplostomidae were divided into two groups based on previous work (Hernández-Mena et al., 2017; Achatz et al., 2019; Gordy & Hanington, 2019). Diplostomidae I consisted of species belonging to the following genera: *Diplostomum* and *Alaria,* with the outgroup of *Ornithodiplostomum scardinii* (Gordy & Hanington, 2019). Diplostomidae II was comprised of the genera *Posthodiplostomum*, *Ornithodiplostomum*, and *Bolbophorus,* with the outgroup being *Crocodilicola pseudostoma* (Hernández-Mena et al., 2017; Achatz et al., 2019; Gordy & Hanington, 2019).

#### 2.2.2 Echinostomatidae

Five phylogenetic trees were run for the Echinostomatidae. The first set of trees included species belonging to the genera *Echinoparyphium*, *Hypoderaeum*, and *Drepanocephalus*, with one tree run with the *COI* gene and the other with the *nad1* gene. The outgroups used were *Euparyphium capitaneum* for the *COI* tree and *Isthmiophora melis* for the *nad1* tree (Tkach et al., 2016; Soldánová et al., 2017; Gordy & Hanington, 2019). The second group included species in the genus *Echinostoma*, with one tree each for *COI* and *nad1*. The outgroups used were the same as above. The final tree included species belonging to the genera *Petasiger* and *Neopetasiger* using *nad1* sequences. The outgroup used was *Fasciola hepatica* (Gordy & Hanington, 2019).

#### 2.2.3 Leucochloridiidae

Three specimens of *Leucochloridium* sp. were found during snail collections for this study in 2019. All three specimens were collected at the same pond (SC3; Figure 1) from *Oxyloma* sp. snails. These trematodes differ from the others found during this study, the parasite brood sac inhabits the eyestalk of its snail host as this is thought to attract its avian definitive host (Casey et al., 2003; Kagan, 1951; Nakao et al., 2019; Núñez et al., 2020; Wesołowska & Wesołowski, 2014; Yamada & Fukumoto, 2011). The snail was crushed between two glass slides to extract the brood sac. The brood sacs were preserved in 95% ethanol and stored at −20°C until DNA extraction was performed as described above.

#### 2.2.4 Notocotylidae

For the Notocotylidae phylogenetic analysis, species in the genus *Notocotylus* and *Ogmocotyle* were included, and the outgroup used was *Echinostoma hortense* (Gordy & Hanington, 2019). Other *COI* sequences of *Notocotylus* were available on GenBank; however, they were not from the Folmer region (Folmer et al., 1994) of the *COI* gene and did not align with the sequences from this study.

#### 2.2.5 Plagiorchiidae

One phylogenetic tree contained only species belonging to the genus *Plagiorchis,* and another was run with *Manodistomum* species. This was due to the short fragment length of the *Manodistomum* sp. sample from this study. The outgroup for both trees was *Haematoloechus* sp. (Zikmundová et al., 2014; Gordy & Hanington, 2019; Li et al., 2022).

#### 2.2.6 Psilostomidae

The tree for the Psilostomidae used the outgroup *Echinocasmus japonicus* (Tkach et al., 2016; Gordy & Hanington, 2019) and included Psilostomidae gen sp. A and *Riberoia ondatrae* sequences.

#### 2.2.7 Schistosomatidae

Two trees were run for the Schistosomatidae, and included species known to cause swimmer’s itch. The first tree included members of the avian schistosomes, with species belonging to the genera *Allobilharzia*, *Anserobilharzia*, *Dendritobilharzia*, *Gigantobilharzia, Nasusbilharzia, Trichobilharzia*, along with undescribed Avian schistosomatid species, and used two Schistosomatidae sp. sequences for the outgroup (McPhail et al., 2021). The second tree included mammalian schistosomes known to cause swimmer’s itch, belonging to the genera *Heterobilharzia, Schistosoma,* and *Schistosomatium* (Loker et al., 2022), and the outgroup used was *Schistosoma bovis* (Gordy & Hanington, 2019).

#### 2.2.8 Strigeidae

Members of the Strigeidae were divided into two groups based on previous research (Hernández-Mena et al., 2017; Gordy & Hanington, 2019). Strigeidae I consisted of the genera *Cotylurus, Cardiocephaloides*, and *Ichthyocotylurus,* while Strigeidae II comprised species belonging to *Australapatemon* and *Apatemon*. The outgroup used for Strigeidae I was *Tylodelphys scheuringi* (Hernández-Mena et al., 2017; Gordy & Hanington, 2019), while the outgroup *Apharyngostrigea pipientis* was used for Strigeidae II (Hernández-Mena et al., 2017; Gordy & Hanington, 2019).

### 2.3 Traditional biodiversity assessments

Bushnell Core DS Low Glow Trail Cameras (Model 119975C) (Bushnell Corporation, Kansas, USA) were employed. Two cameras were in the vicinity of the wetlands (Figure 1) from mid-May to the end of August in 2020, 2021 and 2022, each being secured to six-foot metal stakes driven into the ground. The cameras were motion-activated, and three photographs were taken during each activation. Wildlife Acoustics Song Meter Minis (Wildlife Acoustics, Inc., Massachusetts, USA) were also employed and set to record 24 hours a day, which resulted in hour-long files for the analyses. The recorders were attached to the metal stakes below the field cameras with their microphones oriented upward. No field cameras or recorders were placed at Heritage Lake.

Visual bird identifications were aided by the Sibley Birds application v1 for iPhone (Cool Ideas LLC, Florida, USA) and the web page All About Birds from Cornell University (https://www.allaboutbirds.org/news/). The bird song recordings were analyzed using the program BirdNet (Kahl et al., 2021), which analyzes files in 3-second intervals to determine species identifications. As the recordings were used to supplement field camera identifications, only results with a confidence ≥ 90% were retained. All species identifications were reviewed to ensure that they could plausibly be found in central Alberta.

Benthic kick netting was performed during the 2021 and 2022 collection seasons by stirring up the sediment in several spots at each site and pulling the 500μm kick net through the water above the disturbed area to capture any invertebrates. The net was then emptied into a plastic, sealable 1.2L container with water from the site and brought to the lab. All kick net samples from a single site were combined into one container. Invertebrates from each site were grouped based on morphology, preserved in 95% ethanol, and frozen at −20° C while awaiting DNA sequencing. A subset of the samples collected in 2022 was chosen for DNA sequencing based on distinct morphological features. DNA was amplified using the pan-invertebrate primers HCO2198 and LCO1490 in 20µL reactions following published thermocycling protocols (Folmer et al., 1994).

### 2.4 Diversity calculations, statistical analyses, and visualizing host-parasite interactions

#### 2.4.1 Community diversity

The R packages vegan (Oksanen et al., 2025) and BiodiversityR (Kindt & Coe, 2005) were used to calculate diversity measures: species richness (vegan::specnumber), Shannon diversity (H) (BiodiversityR::diversityresult), and pooled gamma (γ) diversity (Hγ) (BiodiversityR::diversityresult). Effective species were calculated for each site and year combination as exp(H) (Jost, 2006). Evenness was calculated for each site and year combination as exp(H)/species richness (Kindt & Coe, 2005). Beta diversity (β) was calculated using exp(Hγ)/exp(H) for each site and year combination (Jost, 2006).

We explored whether trematode richness differed significantly between snail hosts using Analysis of Variance (ANOVA). *Aplexa* sp. and *Oxyloma* sp. snails were excluded from the analyses because *Aplexa* sp. did not produce any patent infections, and *Oxyloma* sp. was only infected with one trematode species, at SC3 in 2019. Following the ANOVA, a Tukey’s Honest Significant Difference (HSD) test was run to compare differences among snail species.

#### 2.4.2 Sampling completeness

The R package iNEXT.4steps (Chao et al., 2020) was used to analyze sampling completeness, and the iNEXT package (Hsieh et al., 2016) was used for rarefaction analyses, the plots for each of the analyses were made using the package ggplot2 (Wickham, 2016). Sampling completeness analyses were run for vertebrates, invertebrates, trematodes, snails, and snail-trematode interaction counts. Rarefaction analyses were run for trematode species and snail-trematode interactions.

Additionally, we combined the data collected in the present study with previous snail and trematode collections in central Alberta (Gordy et al., 2016; Gordy & Hanington, 2019) to assess sampling completeness and determine whether trematode species discoveries in the region have plateaued using rarefaction analysis. Gordy and Hanington (2019) reported that snails were collected at six lakes across three years [see (Gordy et al., 2016) for site locations]. Rarefaction curves were extrapolated up to an end-point of 7 000 individuals, approximately double the observed trematode infections with DNA sequence data (Chao et al., 2020) from snail-trematode collections in central Alberta.

#### 2.4.3 Effect of wetland age on trematode species richness

We tested the effect of wetland remediation age (stage) and year on trematode species richness, with site included as a random intercept. Wetlands 0-5 years into their remediation were considered early, 6-10 years were considered mid-stage, and 11+ years were considered late-stage (Alberta Environment, 2008; CPP Environmental, 2017; Browne et al., 2018). The trematode data from Heritage Lake was excluded from this analysis. We fit a generalized linear mixed model (GLMM) in R (R Core Team, 2023) with a negative binomial using the package glmmTMB (Brooks et al., 2017) to account for overdispersion. Overdispersion was assessed using the performance package (Lüdecke et al., 2021). Likelihood ratio tests (LRTs) were run using the package lmtest (Zeilis & Hothorn, 2002) to determine the significance of model components.

#### 2.4.4 Visualizing host-parasite interactions

A tripartite diagram was assembled using the bipartite package (Dormann et al., 2024). The first level represented observed snail-trematode relationships, while the second level included trematode-definitive host interactions, based on host associations reported in the literature. Each possible interaction was entered as a value of 1 if it was observed (for level one) or reported (for level 2), and 0 if it had not. For the snail-trematode level, interactions were assigned based on trematode sequence identifications. The trematode-definitive host level was inferred from existing literature rather than direct observation, meaning some interactions may be missing.

A host-parasite web was assembled using MindMup (Sauf Pompiers Limited, 2025) to infer host-parasite interactions based on the traditional biodiversity assessment results. The information included in the tripartite diagram was also used for the trematode web. Inferred host relationships are based on information available in the Natural History Museum Host-Parasite Database (Gibson et al., 2005) using only species identified at the study sites through field camera and recorder data.

#### 2.4.5 Trematode seasonality

We visualized trematode seasonality across study years using ggplot2 (Wickham, 2016). The trematode species were separated by life cycle type and designated as either sporocyst (species that produce cercariae within the sporocysts) or rediae (species that develop rediae within the sporocysts, which then produce cercariae). We opted to plot them based on sporocyst or rediae life cycle category to visualize the seasonal patterns of both life cycle types.

## 3. Results

### 3.1 Study sites and sample collections

From the eight wetland sites and Heritage Lake, 22 397 snails were collected between 2019-2022 (Table 2), and 1 981 were infected with a digenean trematode (8.85%) (Tables 2, 3, and 4). Of the trematode infections, species identifications were obtained for 1 482 (74.81%) (Tables 2, 3, and 4). Snails from six genera were observed in the wetlands and Heritage Lake, but the composition differed slightly between sites. These include: *Lymnaea stagnalis* (10 367, 46.29%), *Physa gyrina* (6 531, 29.16%), *Stagnicola elodes* (3 210, 14.33%), *Oxyloma* sp. (1 096, 4.89%), *Planorbella trivolvis* (995, 4.44%), and *Aplexa* sp. (198, 0.88%) (Figure 2, Table 5). We have also added new snail host records for 20 trematode species (see Supplementary Table A1). Of the snail species infected with trematodes, *Lymnaea stagnalis* snails had the most infections, and *Oxyloma* sp. had the least (Table 5). *Aplexa* sp. had no observed infections and was an interesting observation at site SC6. *Aplexa* sp. snails appeared on June 27, 2022, had dwindled by July 11^th^, 2022, and disappeared from the site by July 25, 2022. No evidence of *Aplexa* sp. was observed at other collection sites.

**Figure 2.**
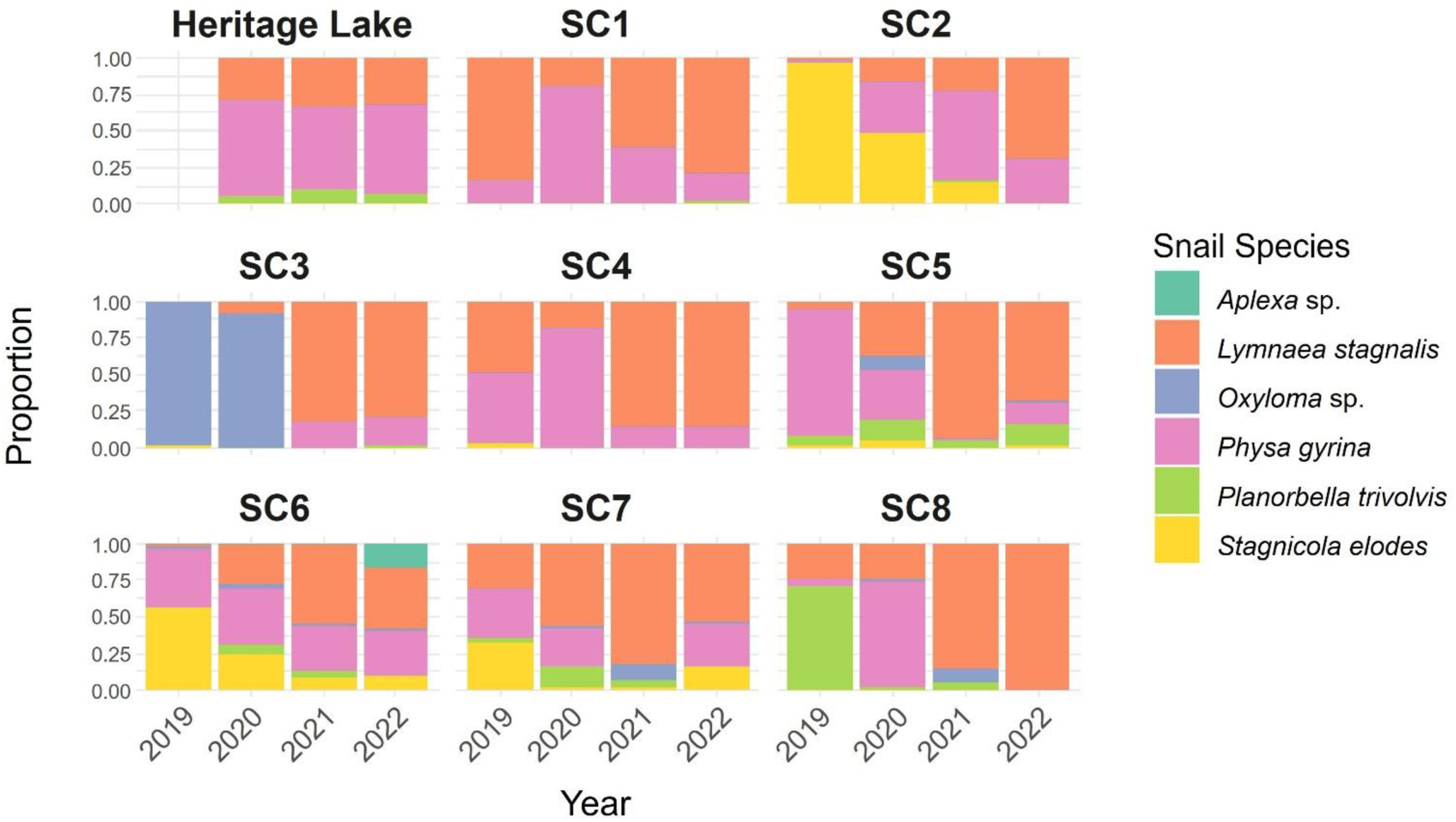
Plot of snail proportions at each site during each collection year. Collections did not start at Heritage Lake until 2020.

**Table 2.** Summary of snail collections from the eight wetland sites near Edmonton, Alberta, Canada. SC = Strathcona County sites. Infections with IDs are those identified using DNA barcoding.

| Site | Snail<br>Richness | Snails<br>Collected | Snails<br>Infected | Infections<br>with IDs |
| --- | --- | --- | --- | --- |
| **SC1** |  |  |  |  |
| 2019 | 4 | 768 | 125 | 90 |
| 2020 | 3 | 744 | 23 | 19 |
| 2021 | 2 | 229 | 25 | 21 |
| 2022 | 4 | 434 | 18 | 15 |
| **Total** | - | **2 175** | **191** | **145** |
| **SC2** |  |  |  |  |
| 2019 | 3 | 1 210 | 88 | 79 |
| 2020 | 3 | 890 | 91 | 68 |
| 2021 | 4 | 152 | 30 | 24 |
| 2022 | 5 | 1 126 | 65 | 49 |
| **Total** | - | **3 378** | **274** | **220** |
| **SC3** |  |  |  |  |
| 2019 | 3 | 627 | 4 | 4 |
| 2020 | 2 | 349 | 1 | 0 |
| 2021 | 4 | 1 535 | 109 | 85 |
| 2022 | 4 | 1 003 | 67 | 51 |
| **Total** | - | **3 514** | **181** | **140** |
| **SC4** |  |  |  |  |
| 2019 | 3 | 624 | 160 | 103 |
| 2020 | 2 | 195 | 10 | 9 |
| 2021 | 2 | 520 | 81 | 59 |
| 2022 | 2 | 481 | 73 | 55 |
| **Total** | - | **1 820** | **324** | **226** |
| **SC5** |  |  |  |  |
| 2019 | 4 | 545 | 10 | 5 |
| 2020 | 5 | 300 | 11 | 4 |
| 2021 | 3 | 145 | 19 | 10 |
| 2022 | 5 | 125 | 6 | 5 |
| **Total** | - | **1 115** | **46** | **24** |
| **SC6** |  |  |  |  |
| 2019 | 4 | 1 449 | 85 | 60 |
| 2020 | 5 | 600 | 45 | 35 |
| 2021 | 5 | 654 | 69 | 62 |
| 2022 | 6 | 1 240 | 33 | 24 |
| **Total** | - | **3 943** | **232** | **181** |

Table 2 – *Continued from previous page*
| Site | Snail<br>Richness | Snails<br>Collected | Snails<br>Infected | Infections<br>with IDs |
| --- | --- | --- | --- | --- |
| **SC7** |  |  |  |  |
| 2019 | 5 | 863 | 101 | 83 |
| 2020 | 5 | 590 | 148 | 95 |
| 2021 | 5 | 292 | 35 | 33 |
| 2022 | 4 | 615 | 18 | 14 |
| **Total** | - | **2 360** | **302** | **225** |
| **SC8** |  |  |  |  |
| 2019 | 3 | 733 | 84 | 82 |
| 2020 | 3 | 60 | 15 | 10 |
| 2021 | 3 | 20 | 2 | 2 |
| 2022 | 2 | 1 439 | 135 | 88 |
| **Total** | - | **2 252** | **236** | **182** |
| **Heritage Lake** |  |  |  |  |
| 2020 | 3 | 207 | 25 | 20 |
| 2021 | 3 | 821 | 114 | 96 |
| 2022 | 3 | 812 | 56 | 23 |
| **Total** | - | **1 840** | **195** | **139** |

**Table 3.**
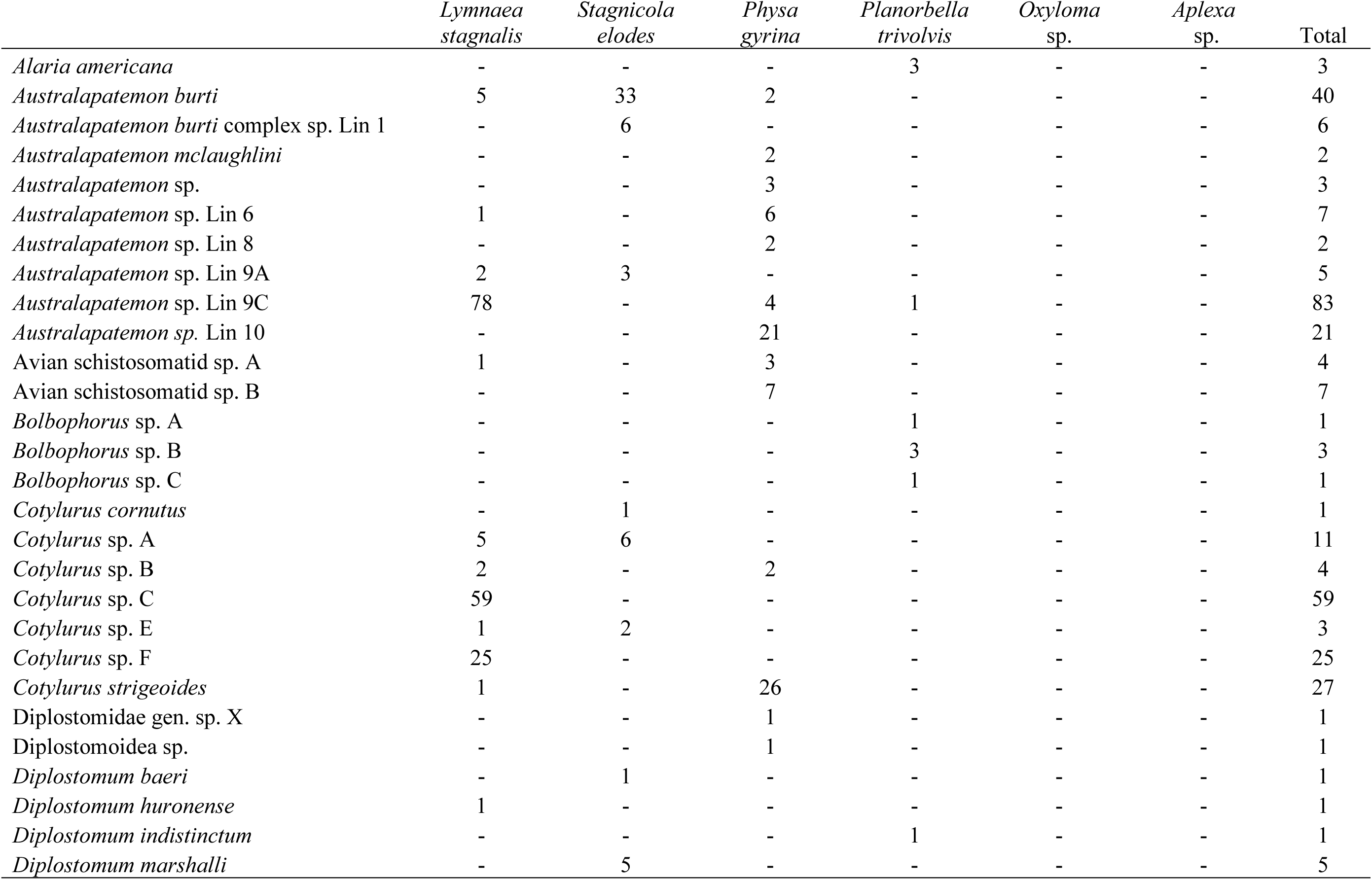
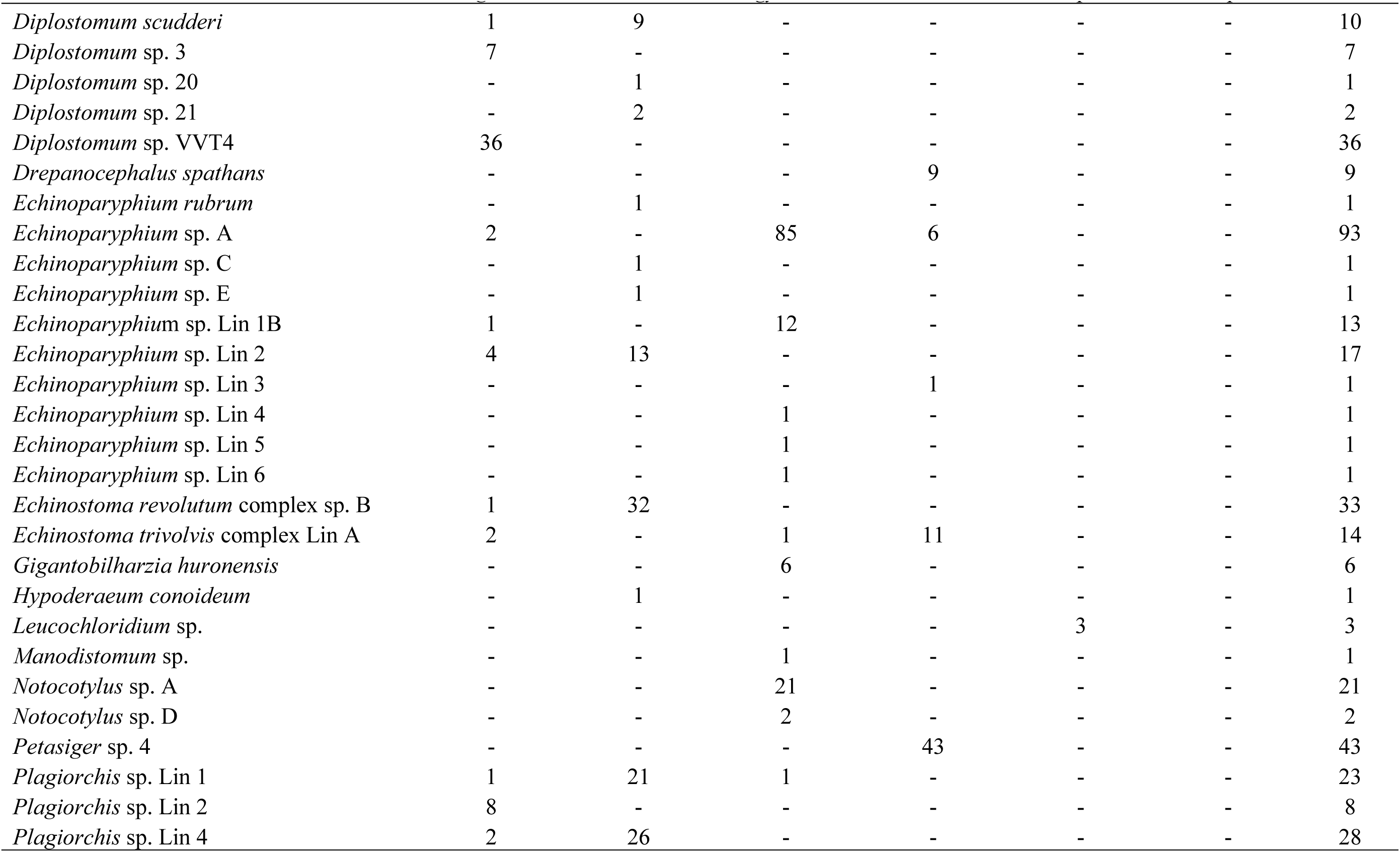
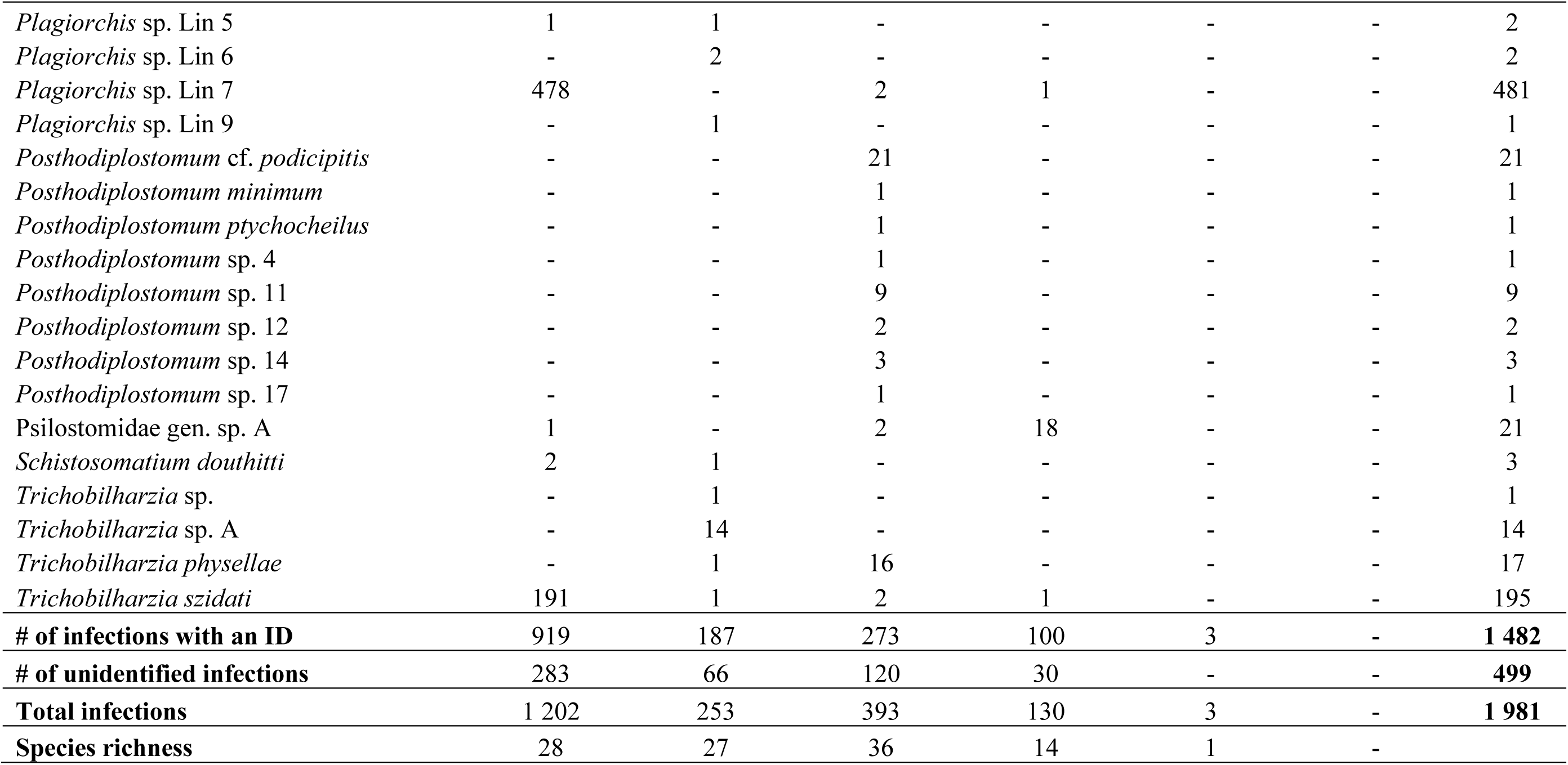
Count of each trematode species in snail hosts across all sampling years combined.

**Table 4.** Count of identifications of each trematode species at collection sites across all sampling years combined. SC = Strathcona County. HL = Heritage Lake, Morinville.

| Species | SC1 | SC2 | SC3 | SC4 | SC5 | SC6 | SC7 | SC8 | HL | Total |
| --- | --- | --- | --- | --- | --- | --- | --- | --- | --- | --- |
| *Alaria americana* | - | - | - | - | - | 1 | - | 2 | - | 3 |
| *Australapatemon burti* | - | 30 | - | - | - | 7 | 2 | 1 | - | 40 |
| *Australapatemon burti* complex sp. Lin 1 | - | 6 | - | - | - | - | - | - | - | 6 |
| *Australapatemon mclaughlini* | - | 1 | - | 1 | - | - | - | - | - | 2 |
| *Australapatemon* sp. | - | - | - | 2 | - | 1 | - | - | - | 3 |
| *Australapatemon* sp. Lin 6 | 1 | 3 | - | - | - | - | - | 1 | 2 | 7 |
| *Australapatemon* sp. Lin 8 | - | 1 | - | - | - | - | - | - | 1 | 2 |
| *Australapatemon* sp. Lin 9A | 1 | 2 | - | - | - | - | 2 | - | - | 5 |
| *Australapatemon* sp. Lin 9C | 13 | 1 | - | 7 | 2 | 11 | 40 | 7 | 2 | 83 |
| *Australapatemon* sp. Lin 10 | 2 | 3 | 3 | 4 | 3 | 5 | - | 1 | - | 21 |
| Avian schistosomatid sp. A | - | - | - | - | - | - | - | - | 4 | 4 |
| Avian schistosomatid sp. B | - | 1 | 1 | 1 | 2 | 1 | - | - | 1 | 7 |
| *Bolbophorus* sp. A | - | - | - | - | - | - | 1 | - | - | 1 |
| *Bolbophorus* sp. B | - | - | - | - | - | 2 | 1 | - | - | 3 |
| *Bolbophorus* sp. C | - | - | - | - | - | - | 1 | - | - | 1 |
| *Cotylurus cornutus* | - | 1 | - | - | - | - | - | - | - | 1 |
| *Cotylurus* sp. A | 1 | 2 | - | - | - | 7 | 1 | - | - | 11 |
| *Cotylurus* sp. B | - | 1 | - | 2 | 1 | - | - | - | - | 4 |
| *Cotylurus* sp. C | 12 | 10 | 1 | 2 | 1 | 20 | 8 | 3 | 2 | 59 |
| *Cotylurus* sp. E | - | 1 | - | - | - | 1 | 1 | - | - | 3 |
| *Cotylurus* sp. F | 2 | 2 | - | 1 | 2 | 3 | 6 | 8 | 1 | 25 |
| *Cotylurus strigeoides* | 8 | 7 | 3 | 2 | - | 2 | 3 | - | 2 | 27 |
| Diplostomidae gen. sp. X | - | - | - | 1 | - | - | - | - | - | 1 |
| Diplostomoidea sp. | - | - | - | - | - | 1 | - | - | - | 1 |
| *Diplostomum baeri* | - | - | - | - | - | 1 | - | - | - | 1 |
| *Diplostomum huronense* | - | 1 | - | - | - | - | - | - | - | 1 |
| *Diplostomum indistinctum* | - | - | - | - | - | - | - | 1 | - | 1 |

Table 4 – Continued from previous page
| Species | SC1 | SC2 | SC3 | SC4 | SC5 | SC6 | SC7 | SC8 | HL | Total |
| --- | --- | --- | --- | --- | --- | --- | --- | --- | --- | --- |
| *Diplostomum marshalli* | - | - | - | - | - | 5 | - | - | - | 5 |
| *Diplostomum scudleri* | - | - | - | - | - | 9 | 1 | - | - | 10 |
| *Diplostomum* sp. 3 | - | - | - | - | - | - | - | - | 7 | 7 |
| *Diplostomum* sp. 20 | - | - | - | - | - | 1 | - | - | - | 1 |
| *Diplostomum* sp. 21 | - | - | - | - | - | 2 | - | - | - | 2 |
| *Diplostomum* sp. VVT4 | 4 | - | - | 4 | - | 9 | 9 | 5 | 5 | 36 |
| *Drepanocephalus spathans* | - | - | - | - | - | - | 1 | 5 | 3 | 9 |
| *Echinoparyphium rubrum* | - | 1 | - | - | - | - | - | - | - | 1 |
| *Echinoparyphium* sp. A | 10 | 18 | 12 | 12 | 2 | 7 | 12 | 2 | 18 | 93 |
| *Echinoparyphium* sp. C | - | - | - | - | - | - | 1 | - | - | 1 |
| *Echinoparyphium* sp. E | - | 1 | - | - | - | - | - | - | - | 1 |
| *Echinoparyphium* sp. Lin 1B | 1 | 7 | - | 1 | - | - | 1 | 1 | 2 | 13 |
| *Echinoparyphium* sp. Lin 2 | - | 14 | - | - | - | 2 | 1 | - | - | 17 |
| *Echinoparyphium* sp. Lin 3 | - | - | - | - | - | - | 1 | - | - | 1 |
| *Echinoparyphium* sp. Lin 4 | - | - | - | 1 | - | - | - | - | - | 1 |
| *Echinoparyphium* sp. Lin 5 | - | - | - | 1 | - | - | - | - | - | 1 |
| *Echinoparyphium* sp. Lin 6 | - | - | - | - | - | 1 | - | - | - | 1 |
| *Echinostoma revolutum* complex sp. B | 1 | 29 | - | - | - | 3 | - | - | - | 33 |
| *Echinostoma trivolvis* complex Lin A | - | 1 | - | 2 | - | 1 | 1 | 3 | 6 | 14 |
| *Gigantobilharzia huronensis* | - | - | - | 3 | - | 2 | - | - | 1 | 6 |
| *Hypoderaeum conoideum* | - | - | - | - | - | 1 | - | - | - | 1 |
| *Leucochloridium* sp. | - | - | 3 | - | - | - | - | - | - | 3 |
| *Manodistomum* sp. | - | - | - | 1 | - | - | - | - | - | 1 |
| *Notocotylus* sp. A | - | - | - | 13 | - | - | - | - | 8 | 21 |
| *Notocotylus* sp. D | - | - | - | 2 | - | - | - | - | - | 2 |
| *Petasiger* sp. 4 | - | - | 1 | - | 2 | 12 | 14 | 7 | 7 | 43 |
| *Plagiorchis* sp. Lin 1 | - | 19 | - | - | - | 3 | 1 | - | - | 23 |
| *Plagiorchis* sp. Lin 2 | 3 | - | 2 | 2 | - | - | 1 | - | - | 8 |

Table 4 – Continued from previous page
| Species | SC1 | SC2 | SC3 | SC4 | SC5 | SC6 | SC7 | SC8 | HL | Total |
| --- | --- | --- | --- | --- | --- | --- | --- | --- | --- | --- |
| *Plagiorchis* sp. Lin 4 | - | 21 | - | 1 | - | 5 | 1 | - | - | 28 |
| *Plagiorchis* sp. Lin 5 | - | 1 | - | - | 1 | - | - | - | - | 2 |
| *Plagiorchis* sp. Lin 6 | - | 1 | - | - | - | - | 1 | - | - | 2 |
| *Plagiorchis* sp. Lin 7 | 44 | 2 | 112 | 132 | 6 | 20 | 42 | 61 | 62 | 481 |
| *Plagiorchis* sp. Lin 9 | - | - | - | - | - | 1 | - | - | - | 1 |
| *Posthodiplostomum* cf. *podicipitis* | 3 | - | - | 5 | - | 2 | 7 | 2 | 2 | 21 |
| *Posthodiplostomum* *minimum* | - | - | - | - | - | 1 | - | - | - | 1 |
| *Posthodiplostomum* *ptychocheilus* | - | - | - | - | - | - | 1 | - | - | 1 |
| *Posthodiplostomum* sp. 4 | - | - | - | - | - | 1 | - | - | - | 1 |
| *Posthodiplostomum* sp. 11 | 2 | 1 | - | 5 | - | 1 | - | - | - | 9 |
| *Posthodiplostomum* sp. 12 | - | - | - | - | - | 1 | - | 1 | - | 2 |
| *Posthodiplostomum* sp. 14 | - | - | - | - | - | 1 | - | 2 | - | 3 |
| *Posthodiplostomum* sp. 17 | - | - | - | - | - | - | 1 | - | - | 1 |
| Psilostomidae gen. sp. A | - | - | - | 1 | - | - | 3 | 16 | 1 | 21 |
| *Schistosomatium* *douthitti* | - | 1 | - | 1 | 1 | - | - | - | - | 3 |
| *Trichobilharzia* sp. | - | 1 | - | - | - | - | - | - | - | 1 |
| *Trichobilharzia* sp. A | - | 14 | - | - | - | - | - | - | - | 14 |
| *Trichobilharzia* *physellae* | 2 | 6 | - | - | - | 6 | 1 | 1 | 1 | 17 |
| *Trichobilharzia* *szidati* | 35 | 9 | 2 | 16 | 1 | 21 | 58 | 52 | 1 | 195 |
| **Total** | 145 | 220 | 140 | 226 | 24 | 181 | 225 | 182 | 139 | **1482** |
| **Species richness** | 18 | 35 | 10 | 28 | 12 | 39 | 32 | 21 | 22 | - |

**Table 5.** Counts of snail species combined across all collection sites. SC = Strathcona County, Alberta. HL = Heritage Lake, Morinville.

| Snail species | SC1 | SC2 | SC3 | SC4 | SC5 | SC6 | SC7 | SC8 | HL | Total |
| --- | --- | --- | --- | --- | --- | --- | --- | --- | --- | --- |
| *Lymnaea stagnalis* | 1 256 | 970 | 2 093 | 1 204 | 365 | 1 073 | 1 163 | 1 646 | 597 | 10 367 |
| *Stagnicola elodes* | 4 | 1 640 | 6 | 16 | 19 | 1 135 | 390 | - | - | 3 210 |
| *Physa gyrina* | 905 | 762 | 456 | 600 | 600 | 1 406 | 623 | 83 | 1 096 | 6 531 |
| *Planorbella trivolvis* | 8 | 5 | 12 | - | 103 | 70 | 130 | 520 | 147 | 995 |
| *Oxyloma* sp. | 2 | 1 | 947 | - | 28 | 61 | 54 | 3 | - | 1 096 |
| *Aplexa* sp. | - | - | - | - | - | 198 | - | - | - | 198 |
| Total | 2 175 | 3 378 | 3 514 | 1 820 | 1 115 | 3 943 | 2 360 | 2 252 | 1 840 | 22 397 |

Seven co-infections were observed during this study: one each in 2019 and 2020, two in 2021, and three in 2022. Each co-infection occurred in a *Lymnaea stagnalis* snail. The two types of cercariae from each snail were preserved separately based on morphological differences.

Based on morphological characteristics, all co-infections presented one species of *Plagiorchis,* with the second cercarial type being either an avian schistosome (6/7) or a strigeid (1/7). For four of these co-infections, no high-quality sequence data was obtained for either cercarial morphotype. For the remaining three co-infections, only one cercarial morphotype from each snail yielded high-quality sequence data, one *Trichobilharzia szidati*, one *Plagiorchis* sp. Lin 7, and one *Australapatemon* sp. Lin 9C.

### 3.2 Trematode species identification

Throughout this study, we identified 74 trematode species, 23 of which had not been previously identified in central Alberta. Combined with the trematode species identified by previous research in the province (Gordy & Hanington, 2019), our total for central Alberta is 102 species. The family Plagiorchiidae made up the largest proportion of trematodes identified (36.8%), while Leucochloridiidae made up the smallest (0.2%) (Supplementary Figure A1). Two species, Avian schistosomatid sp. A, and *Diplostomum* sp. 3, were identified at Heritage Lake, but were not present at the wetlands. Within the trematode species observed between the sites, we identified novel operational taxonomic units (OTUs) that presented no matches to publicly available DNA sequences on GenBank (Benson et al., 2013) or the Barcode of Life Data Systems (BOLD) (Ratnasingham et al., 2024). We acknowledge that these have been identified from cercarial larval stages and indicated as distinct OTUs in single-gene phylogenies, with the exception of two putative novel species for which we have both *COI* and *nad1* sequences. We will refer to these OTUs as species for consistency throughout. The phylogenies presented herein use representative sequences for each species. While we agree with the call to action from Hechinger (2023), the sample collections for this project ended prior to its publication, and we lack detailed morphological identification beyond gross morphology, sufficient only for assigning specimens to a trematode family. Therefore, we will proceed using collective group names.

We observed 74 species from five snail hosts based on DNA sequences and molecular phylogenies. The site with the greatest species richness was SC6, and the lowest was SC3 (Table 6). Notably, the snail richness at SC6 ranged between four and six snail species across collection seasons, while the range at SC3 was between two and four snail species (Table 2). Other sites had lower snail richness (i.e., SC4 and SC8, Table 2), though SC3 had many *Oxyloma* sp. snails (Table 4). The most common trematode species identified during the study was *Plagiorchis* sp. Lineage 7 (Table 4).

**Table 6.** Community diversity measures by site and year combinations. Effective species were calculated as exp(H). Evenness was calculated as Effective species/Species richness. Pooled Shannon diversity was calculated from pooled data across all sites and years. Pooled γ diversity was calculated as exp(pooled γ)/exp(H). SC = Strathcona County, Alberta. HL = Heritage Lake, Morinville.

| Site | Year | Species richness | Shannon diversity (H) | Effective species | Evenness | $\beta$ diversity |
| --- | --- | --- | --- | --- | --- | --- |
| SC1 | 2019 | 8 | 1.43 | 4.17 | 0.52 | 4.16 |
|  | 2020 | 9 | 1.95 | 7.04 | 0.78 | 2.47 |
|  | 2021 | 9 | 2.07 | 7.93 | 0.88 | 2.19 |
|  | 2022 | 7 | 1.77 | 5.85 | 0.84 | 2.97 |
| SC2 | 2019 | 9 | 1.84 | 6.27 | 0.70 | 2.77 |
|  | 2020 | 20 | 2.33 | 10.33 | 0.52 | 1.68 |
|  | 2021 | 11 | 2.05 | 7.74 | 0.70 | 2.24 |
|  | 2022 | 13 | 2.08 | 7.97 | 0.61 | 2.18 |
| SC3 | 2019 | 2 | 0.56 | 1.75 | 0.88 | 9.89 |
|  | 2020 | 0 | 0.00 | 1.00 | - | 17.36 |
|  | 2021 | 7 | 0.52 | 1.67 | 0.24 | 10.37 |
|  | 2022 | 4 | 0.79 | 2.20 | 0.55 | 7.88 |
| SC4 | 2019 | 20 | 1.90 | 6.70 | 0.34 | 2.59 |
|  | 2020 | 4 | 1.15 | 3.16 | 0.79 | 5.50 |
|  | 2021 | 12 | 1.50 | 4.46 | 0.37 | 3.89 |
|  | 2022 | 11 | 1.24 | 3.45 | 0.31 | 5.03 |
| SC5 | 2019 | 3 | 1.05 | 2.87 | 0.96 | 6.04 |
|  | 2020 | 3 | 1.04 | 2.83 | 0.94 | 6.14 |
|  | 2021 | 7 | 1.83 | 6.26 | 0.89 | 2.77 |
|  | 2022 | 4 | 1.33 | 3.79 | 0.95 | 4.58 |
| SC6 | 2019 | 20 | 2.68 | 14.63 | 0.73 | 1.19 |
|  | 2020 | 15 | 2.45 | 11.54 | 0.77 | 1.50 |
|  | 2021 | 13 | 2.24 | 9.42 | 0.72 | 1.84 |
|  | 2022 | 10 | 2.06 | 7.86 | 0.79 | 2.21 |
| SC7 | 2019 | 25 | 2.38 | 10.82 | 0.43 | 1.60 |
|  | 2020 | 11 | 1.87 | 6.46 | 0.59 | 2.69 |
|  | 2021 | 10 | 2.03 | 7.63 | 0.76 | 2.28 |
|  | 2022 | 4 | 1.12 | 3.05 | 0.76 | 5.68 |
| SC8 | 2019 | 12 | 1.69 | 5.44 | 0.45 | 3.19 |
|  | 2020 | 8 | 2.03 | 7.58 | 0.95 | 2.29 |
|  | 2021 | 2 | 0.69 | 2.00 | 1.00 | 8.68 |
|  | 2022 | 6 | 1.29 | 3.62 | 0.60 | 4.79 |
| HL | 2020 | 8 | 1.73 | 5.64 | 0.70 | 3.08 |
|  | 2021 | 18 | 1.87 | 6.46 | 0.36 | 2.69 |
|  | 2022 | 10 | 2.15 | 8.57 | 0.86 | 2.03 |
| Pooled Shannon diversity |  |  | 2.85 |  |  |  |
| Pooled $\gamma$ diversity | | | 17.36 | | | |

#### 3.2.1 Diplostomidae

The final alignment for Diplostomidae I was 385 bp and included 60 sequences. The species of *Alaria americana* identified during this study grouped with a specimen collected in Quebec, Canada (Locke et al., 2018) (Figure 3). During this study, two novel species of *Diplostomum* were observed that presented no strong matches to publicly available sequences. We have labelled them as *Diplostomum* sp. 20 and *Diplostomum* sp. 21 (Figure 3).

**Figure 3.**
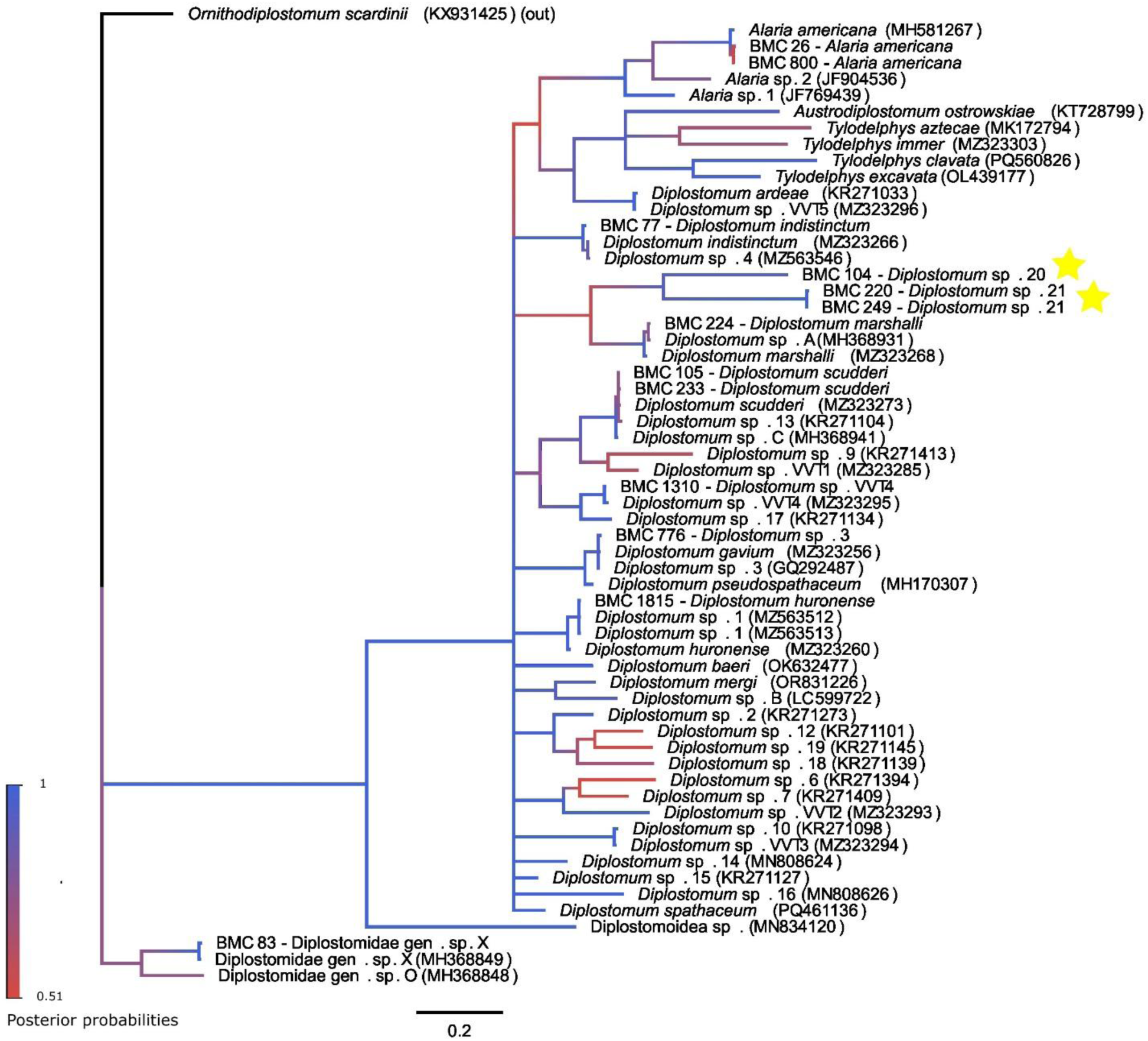
Bayesian inference molecular phylogeny of the Diplostomidae-I based on *COI* using the GTR + G + I nucleotide substitution model. Accession numbers follow species names. Samples with a “BMC” label indicate sequences obtained during this study. Star symbols indicate species newly identified by this study.

The final alignment for Diplostomidae II was 314 bp and included 53 sequences. During this study, we found three unique species of *Bolbophorus,* which we have called *Bolbophorus* sp.

A, B, and C, all collected from *Planorbella trivolvis* snails. They share between 80-92% similarity (Supplementary Table B2). *Bolbophorus* sp. A groups within the *Bolbophorus* clade, but shares only 81-93% similarity with members of the genus included from this study and the literature (Supplementary Table B2), and we consider this a putative novel species (Figure 4). *Bolbophorus* sp. B groups with the species of *Bolbophorus* previously identified in Alberta (Gordy et al., 2016; Gordy & Hanington, 2019), as well as with a *Bolbophorus* sp. collected in Mississippi, USA (Rosser, Baumgartner, et al., 2016) (Figure 4, Supplementary Table A1). *Bolbophorus* sp. C groups with a *Bolbophorus* sp. collected in southern Alberta (Van Steenkiste et al., 2015).

**Figure 4.**
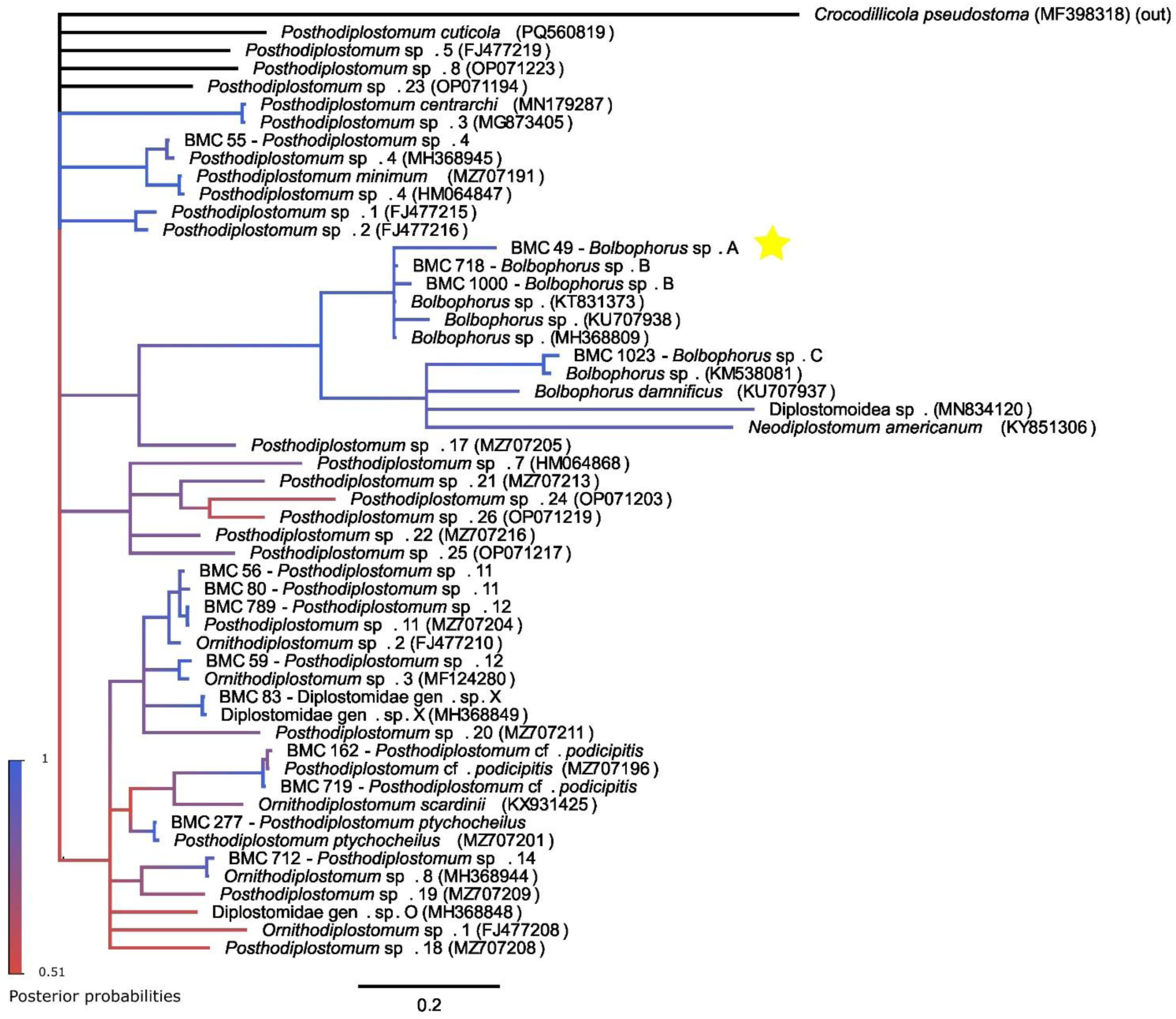
Bayesian inference molecular phylogeny of the Diplostomidae-II based on *COI* using the HKY + G + I substitution model. Accession numbers follow species names. Samples with a “BMC” label indicate sequences obtained during this study. Star symbols indicate species newly identified by this study.

Additionally, sequences belonging to the genera *Posthodiplostomum* and *Ornithodiplostomum* form a clade, with some species of *Posthodiplostomum* separated near the top of the tree (Figure 4). Recently, researchers have demonstrated a close relationship between the genera *Posthodiplostomum* and *Ornithodiplostomum,* along with a third genus, *Mesoophorodiplostomum,* which was not observed during this study (Achatz et al., 2021). This has led to *Ornithodiplostomum* and *Mesoophorodiplostomum* being synonymized with *Posthodiplostomum* (Achatz et al., 2021). The phylogenetic tree of the Diplostomidae II reflects this close relationship (Figure 4), and as such, we have identified our specimens of *Ornithodiplostomum* species 2, 3 and 8 as *Posthodiplostomum* species 11, 12, and 14, respectively, following the updated species names from Achatz and colleagues (2021).

#### 3.2.2 Echinostomatidae

Five phylogenetic trees were run for species within the family Echinostomatidae (Figure 5A, B and Supplementary Figures A2 A, B, A3). There were fewer sequences available for the *COI* tree, as *nad1* is the preferred barcoding gene for the Echinostomatidae (Morgan & Blair, 1998a; Detwiler et al., 2010; Gordy & Hanington, 2019; Pantoja et al., 2021). Additionally, only some of our samples have *COI* and *nad1* sequences to compare.

**Figure 5.**
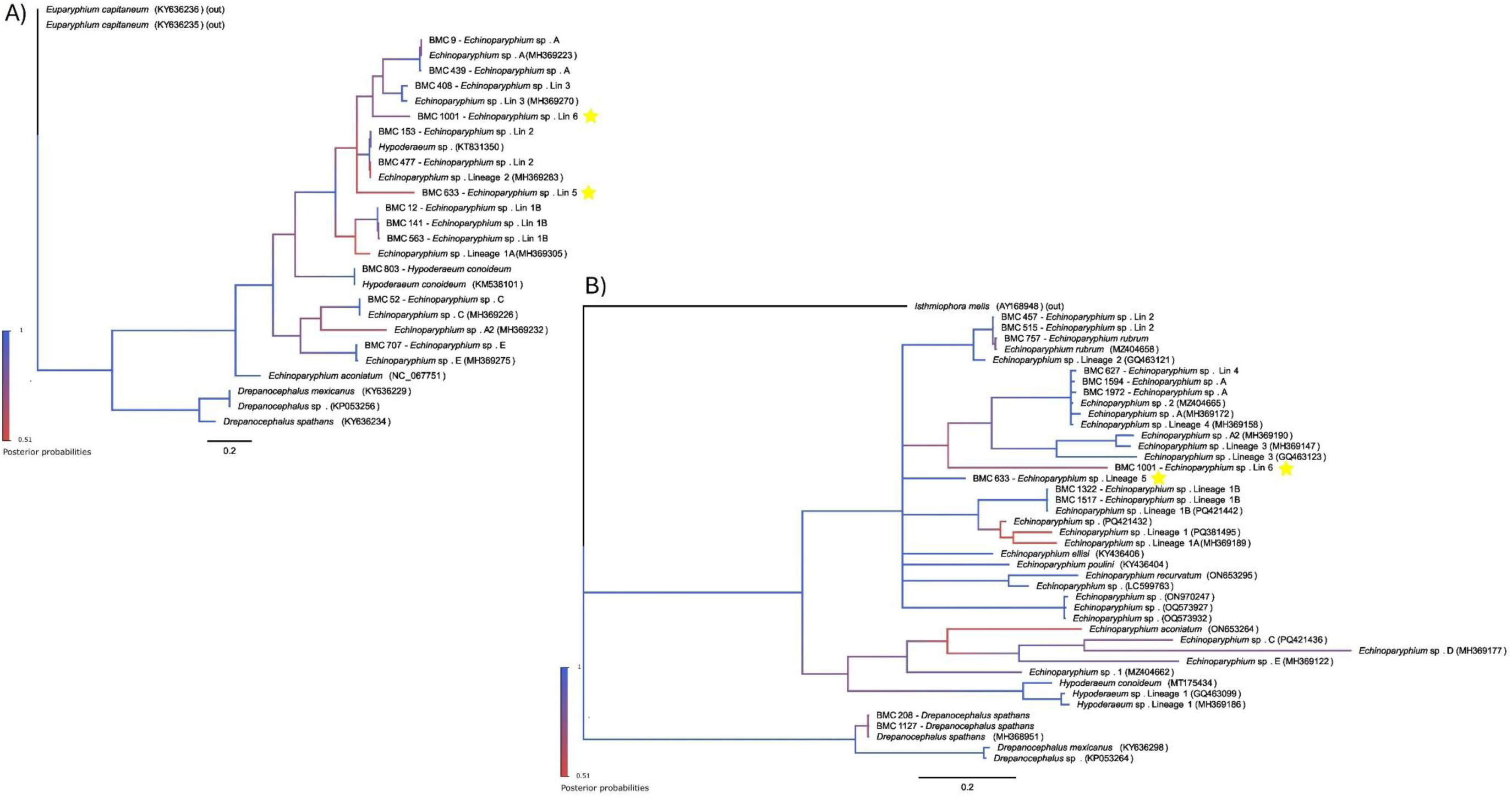
Bayesian inference molecular phylogenies of the Echinostomatidae (genera *Echinoparyphium*, *Hypoderaeum*, and *Drepanocephalus*) based on A) *COI*, with substitution model HKY + G + I, and B) *nad1* using substitution model GTR + G. Accession numbers follow species names. Samples with a “BMC” label indicate sequences obtained during this study. Star symbols indicate species newly identified by this study.

The first set of trees included species belonging to the genera *Echinoparyphium*, *Hypoderaeum*, and *Drepanocephalus* (Figure 5A, B). The *COI* tree for this group had a final alignment of 492 base pairs and included 28 sequences, while the *nad1* tree had a final alignment of 351 base pairs and 43 sequences. We observed two new lineages of *Echinoparyphium* sp. during this study and have called them *Echinoparyphium* sp. Lineage 5 and 6 (Figure 5A).

*Echinoparyphium* sp. Lin 5 and Lin 6 were found to be infecting one *P. gyrina* snail each (Table 3). These lineages share 86.8% identity (Supplementary Table B3) and are both putative novel species which was confirmed with both *COI* and *nad1* genes (Figure 5A, B). Within the *nad1* tree (Figure 5B), the representatives of *Drepanocephalus spathans* group with *D. spathans* previously collected in Alberta (Gordy & Hanington, 2019). While representatives of *Echinoparyphium* sp. A formed a distinct clade in the *COI* tree (Figure 5A), the corresponding clade in the *nad1* tree (Figure 5B) also included representatives of *Echinoparyphium* sp. Lineage 4 and *Echinoparyphium* sp. 2. These taxa require further taxonomic clarification.

The next set of trees for the Echinostomatidae included species belonging to the genus *Echinostoma* (Supplementary Figure A2 A, B) and for both the *COI* and *nad1* genes. The *COI* tree had a final alignment of 458 base pairs and included 16 sequences, while the *nad1* tree had a final alignment of 423 bp and included 36 sequences. As above, there were fewer *COI* sequences available for this group. Only two species within the genus *Echinostoma* identified during this study were *Echinostoma revolutum* and *Echinostoma trivolvis.* In both trees, the representative sequences of *Echinostoma revolutum* group with *Echinostoma revolutum* complex sp. B from Gordy and Hanington (2019) (Supplementary Figure A2 A, B). Again, the representative sequences for *Echinostoma trivolvis* groups with *Echinostoma trivolvis* complex Lineage A from Gordy and Hanington (2019) and the same lineage collected in Indiana, USA (Supplementary Table A1).

The final tree was comprised of species belonging to the genera *Petasiger* and *Neopetasiger* (Supplementary Figure A3). The final alignment was 394 base pairs and included 16 sequences. The representatives of *Petasiger* sp. 4 identified in this study grouped with a sequence of *Petasiger* sp. 4 from the previous survey in Alberta (Gordy & Hanington, 2019), but this clade was separate from another species identified as *Petasiger* sp. 4 collected from a *Biomphalaria pfeifferi* snail in Kenya (Laidemitt et al., 2019) (Supplementary Figure A3), which only shows 70% similarity to *Petasiger* sp. 4 in Alberta (Supplementary Table B7).

#### 3.2.3 Leucochloridiidae

The final alignment for the Leucochloridiidae was 403 bp and contained 21 sequences. Although we found three specimens of *Leucochloridium* sp. during this study, a DNA sequence was only obtained for one representative specimen. Based on the phylogenetic analysis, this is a putative novel species within the genus *Leucochloridium* (Figure 6). The other available *COI* sequences were from Japan, Russia, and Belarus (Supplementary Table A1), for which our specimen shared between 79.4 −82.1% similarity (Supplementary Table B8). Representatives of *L. subtilis* and *L. pulchrum* were omitted because their available sequences did not align with the other *COI* representative sequences used in the analyses.

**Figure 6.**
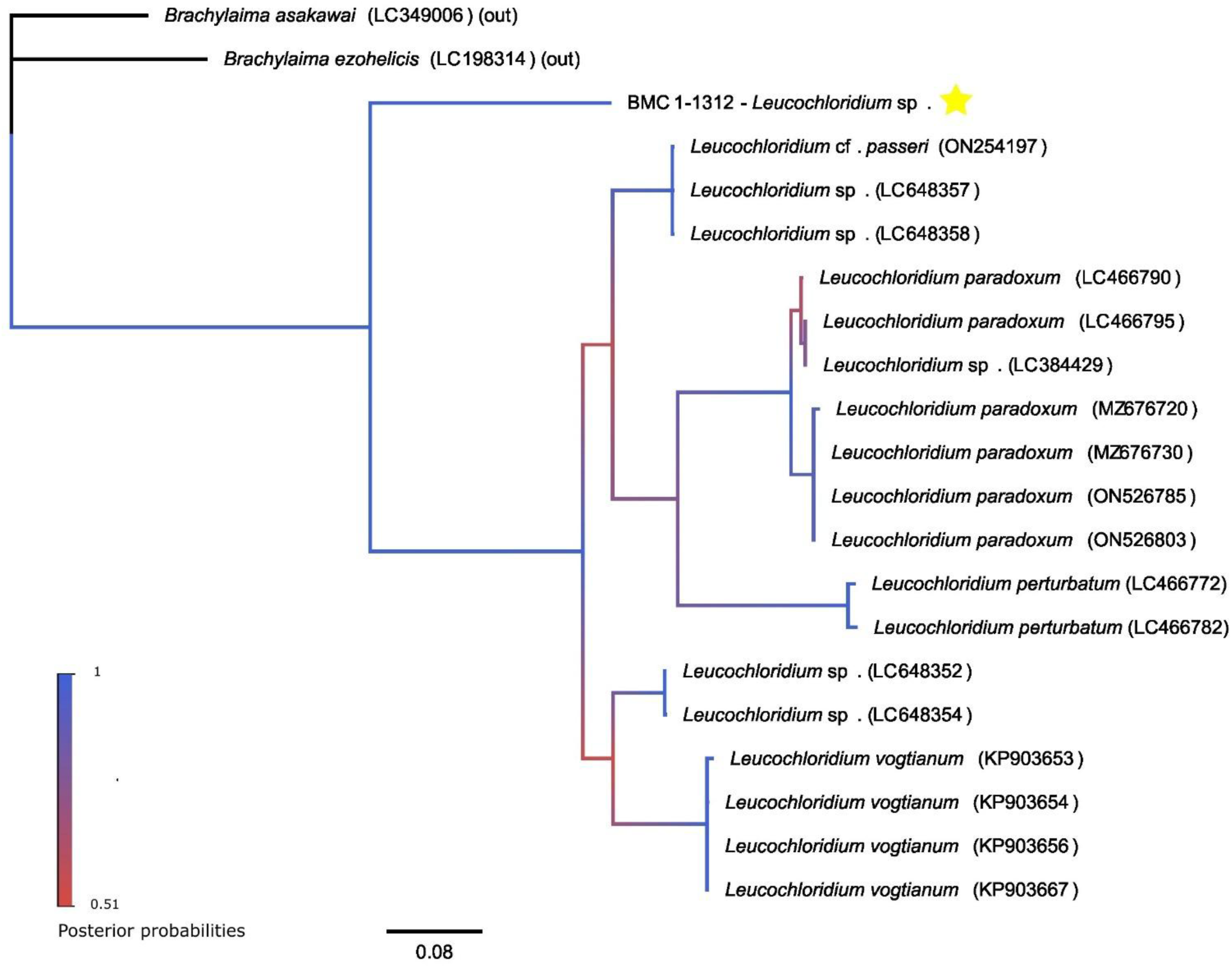
Bayesian inference molecular phylogeny of the Leucochloridiidae based on *COI* using the HKY + G + I substitution model. Accession numbers follow species names. Samples with a “BMC” label indicate sequences obtained during this study. Star symbols indicate species newly identified by this study.

#### 3.2.4 Notocotylidae

The final alignment for the Notocotylidae tree was 495 bp and contained 14 sequences. During this study, we identified *Notocotylus* sp. A, and *Notocotylus* sp. D. Our sequences of *Notocotylus* sp. A group with a representative from the previous work in Alberta and *Notocotylus* sp. B from Alberta (Gordy & Hanington, 2019) (Supplementary Figure A4). Our representative of *Notocotylus* sp. D did not group with the same species identified by Gordy and Hanington (2019).

#### 3.2.5 Plagiorchiidae

The final alignment for the tree containing species in the genus *Plagiorchis* was 329 base pairs and included 37 sequences. The lineages of *Plagiorchis* sp. collected throughout this study matched with the lineages previously collected in Alberta (Gordy & Hanington, 2019), as well as lineages collected in Alaska, USA (Kudlai et al., 2021) (Figure 7A).

**Figure 7.**
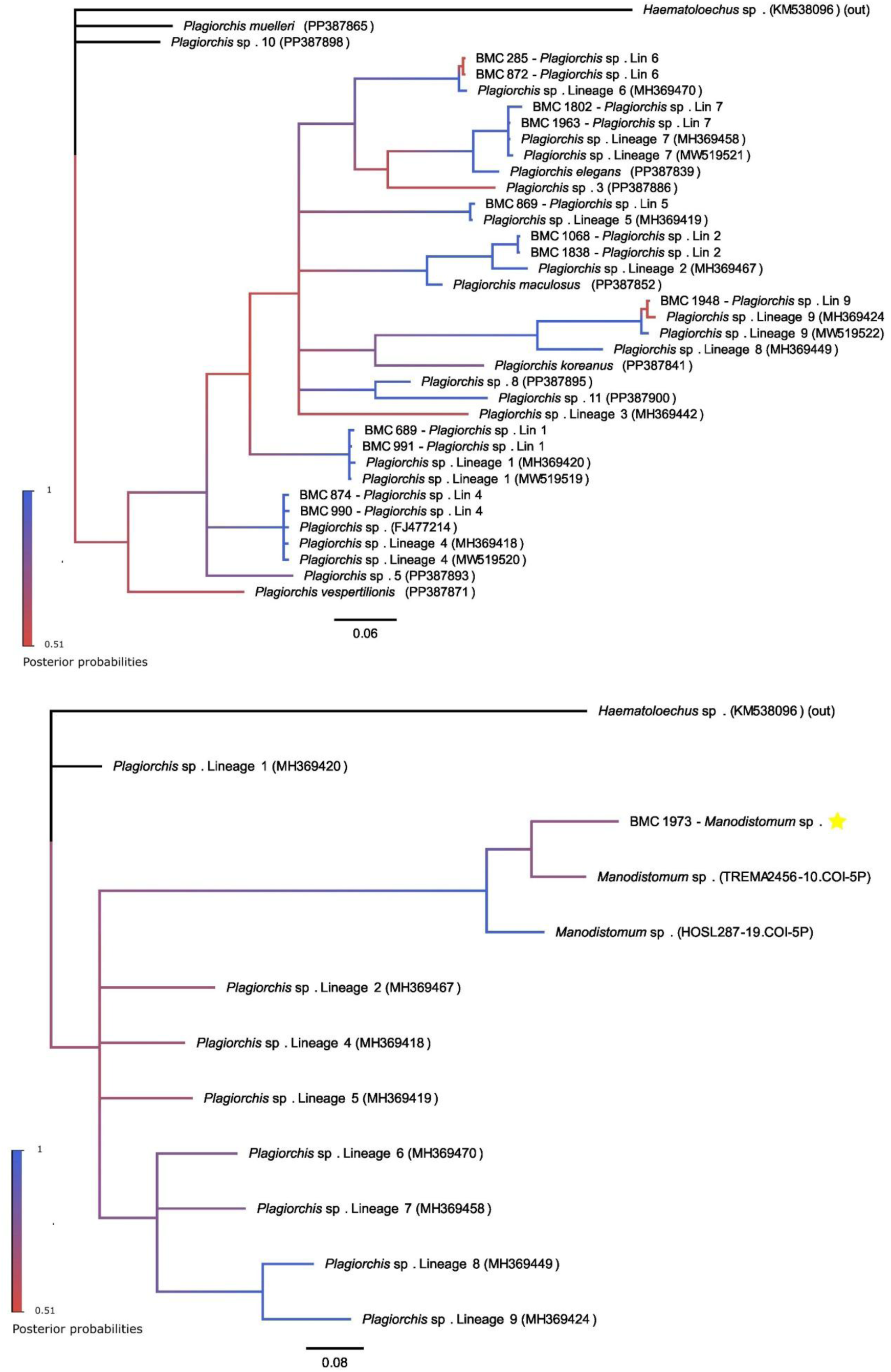
Bayesian inference molecular phylogenies of the Plagiorchiidae based on *COI*. A) *Plagiorchis* species using the HKY + G + I substitution model. B) *Manodistomum* species and *Plagiorchis* lineages previously identified in Alberta using the HKY + G substitution model. Accession numbers follow species names. Samples with a “BMC” label indicate sequences obtained during this study. Star symbols indicate species newly identified by this study.

The second tree for the family Plagiorchiidae had a final alignment of 284 base pairs and included 13 sequences. There was a paucity of sequences belonging to the genus *Manodistomum*. The only two available specimens were found on BOLD (Ratnasingham et al., 2024), with one each collected in California and Missouri, USA (Supplementary Table A1). While the suspected *Manodistomum* sp. collected during this study grouped with the two other *Manodistomum* sp. specimens (Figure 7B), they are not all the same species. The two *Manodistomum* sp. sequences collected in the USA are 87% similar, while the species identified during this study share 84-89% similarity (Supplementary Table B11).

#### 3.2.6 Psilostomidae

The Psilostomidae phylogenetic tree alignment was 473 bp and included 10 sequences. Two different isolates of *Ribeiroia ondatrae* were included in the tree. The isolate included in the analysis from Johnson and colleagues (2021) had two representatives that were collected from *Planorbella trivolvis* snails in California and Oregon (Johnson et al., 2021). The other isolate of *R. ondatrae* included was also collected in California from the California tiger salamander (*Ambystoma californiense*) by Keller and colleagues (2021). However, this sequence did not group with the *R. ondatrae* from Johnson et al. (2021). Instead, it grouped with sequences identified as Psilostomidae gen. sp. A. (Figure 8). These sequences shared approximately 64% similarity (Supplementary Table B12).

**Figure 8.**
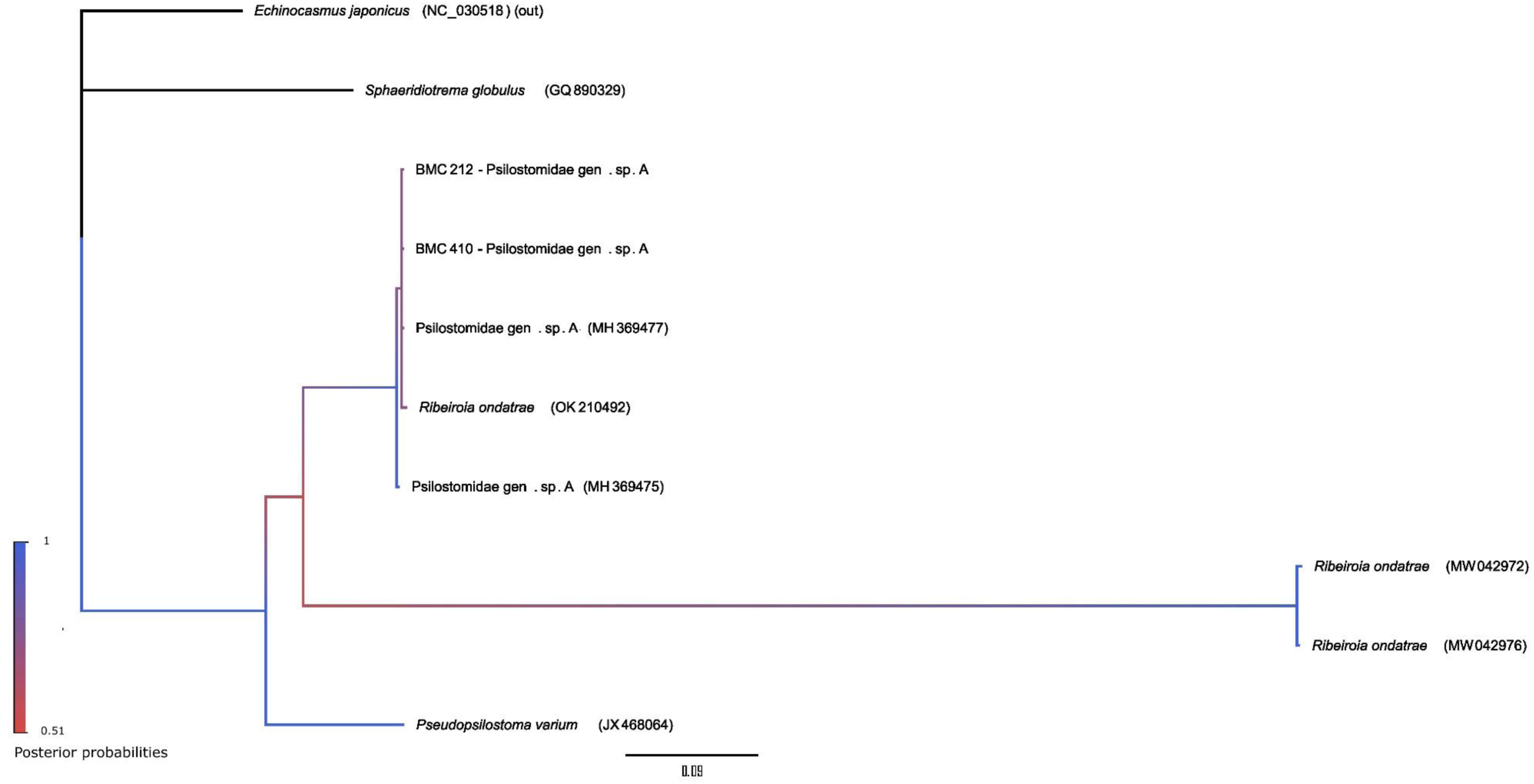
Bayesian inference molecular phylogeny of the Psilostomidae based on *COI*, using substitution model HKY + G. Accession numbers follow species names. Samples with a “BMC” label indicate sequences obtained during this study.

#### 3.2.7 Schistosomatidae

The final alignment for the avian schistosome phylogenetic tree was 413 bp and included 34 sequences. The representative sequences from the present study grouped as expected with members of the avian schistosomes (Supplementary Figure A5 A). The mammalian schistosome phylogenetic tree was run using a final alignment of 507 bp that included seven sequences. The representative sequence of *Schistosomatium douthitti* from this study grouped with other representatives as expected (Supplementary Figure A5 B).

#### 3.2.8 Strigeidae

For the Strigeidae I tree, the final alignment was 457 base pairs and included 28 sequences. The representative *Cotylurus* spp. sequences from the present study grouped as expected (Supplementary Figure A6). For the Strigeidae II analyses, the final alignment was 416 base pairs with 34 sequences. *Australapatemon* sp. Lin 9C is a putative novel species observed during this study. Notably, these cercariae were observed performing “Rat-King” behaviour numerous times. This is a rare phenomenon where the cercariae are joined by their tails to form Rat-King-like groups, after which they were described (Gordy et al., 2017). We found this species emerging from *Lymnaea stagnalis* (n = 78), *Physa gyrina* (n = 4), and *Planorbella trivolvis* (n = 1) (Table 3, Supplementary Table A1). This putative novel species groups with *Australapatemon* sp. Lin 9A and 9B (Figure 9). As such, we designated it as *Australapatemon* sp. Lin 9C, rather than assigning this species as a consecutive lineage. Another interesting clade in this tree included representatives of *Australapatemon* sp. Lin 6 and *Australapatemon* sp. (Figure 9). We found *Australapatemon* sp. Lin 6 emerging from *L. stagnalis* and *P. gyrina* snails (Table 3, Supplementary Table A1). Gordy and Hanington (2019) also collected it from *P. gyrina* snails. As for *Australapatemon* sp., we found it emerging from *P. gyrina* snails, as did Gordy and colleagues (2017) (Table 3, Supplementary Table A1). These representative sequences share ≥ 98% identity (Supplementary Table B16); however, further taxonomic clarification is required.

**Figure 9.**
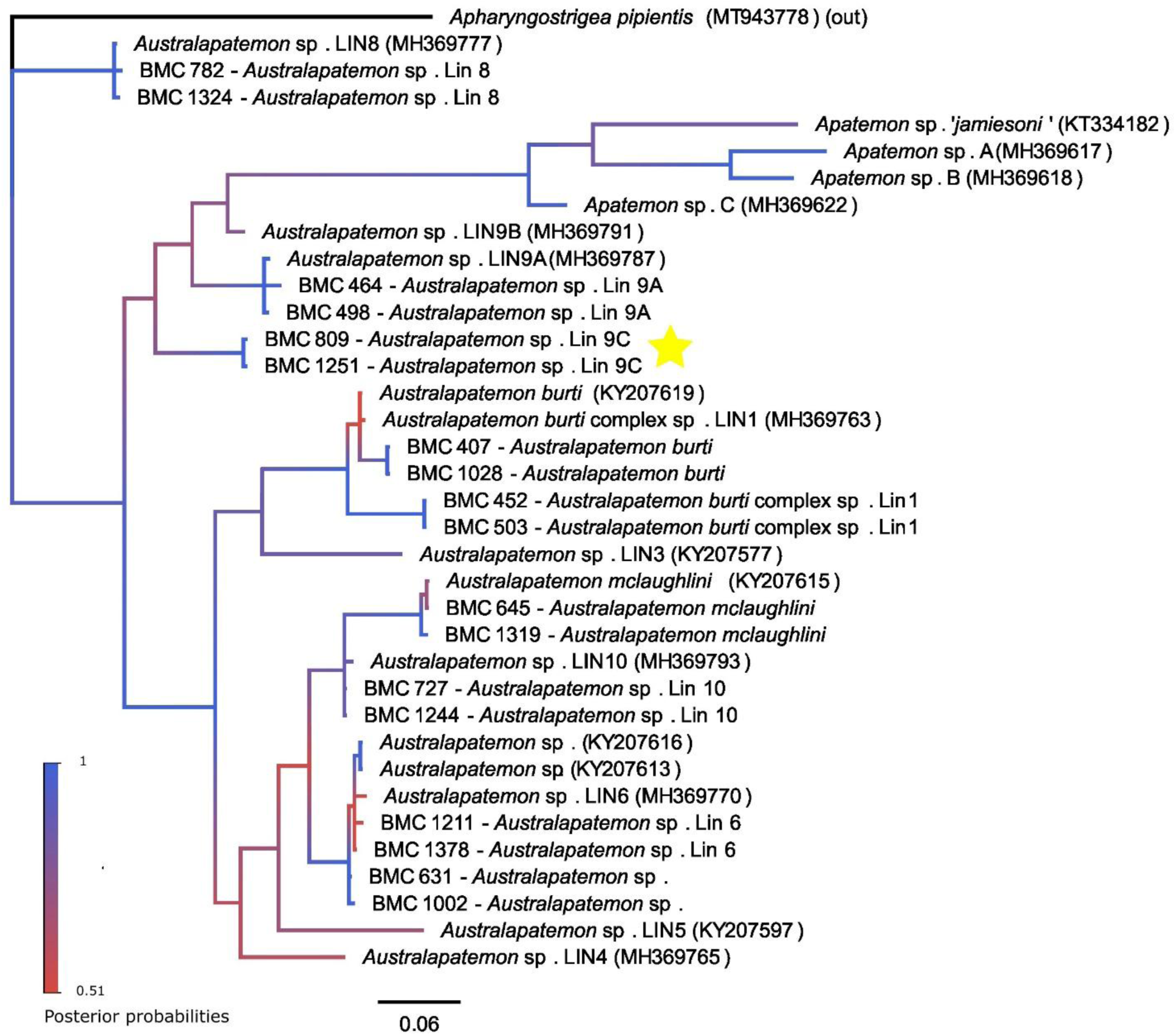
Bayesian inference molecular phylogeny of the Strigeidae II based on *COI* using the HKY + G + I substitution model. Accession numbers follow species names. Samples with a “BMC” label indicate sequences obtained during this study. Star symbols indicate species newly identified by this study.

### 3.3 Traditional biodiversity assessments

We gathered approximately 1 700 hours of recordings from the recorders, and the field cameras yielded approximately 12 000 photos with animal wildlife present within. From the field cameras and bird song recorders, 137 vertebrate species were identified in the study area (Supplementary Table A2). Using the field cameras and recorder data, 41 and 126 species were identified, respectively, and 30 species (21.9%) overlapped between the two methods. Between the two birdsong recorders, there were 63 363 identifications. Of these, 1 901 were implausible identifications (3.0%), meaning that these species were not native to central Alberta or western Canada nor were reported to visit during their migration or breeding periods (Cornell University, 2025; The Cornell Lab of Ornithology, 2025). Sequencing of invertebrates captured by the benthic kick netting was undertaken only for samples taken in 2022. Between the eight wetland sites and Heritage Lake in 2022, 93 invertebrate samples were selected for DNA sequencing based on distinct morphological features (Supplementary Table A3). Of these, 85 returned high-quality sequences that resulted in 46 unique identifications.

### 3.4 Community diversity

The lowest species richness at a site and year combination was at site SC3 in 2020 where no infected snails were found, though species richness at this site remained consistently low. The highest species richness was 25 species at site SC7 in 2019 (Table 6). Evenness among sites was variable. Site SC5 generally had low richness (between 3-7 species), but high evenness (0.94 – 0.96), indicating a relatively equal representation of a few species. While sites with higher richness, e.g. SC4 (species richness ranged from 4 – 20 species), had reduced evenness (0.31-0.79), indicating that some species dominated over others (Table 6). β-diversity between sites and years had a wide range (1.19 – 17.36), indicating that the trematode communities were highly variable (Table 6).

### 3.5 Sampling completeness

Both sampling completeness and rarefaction analyses were run for the trematodes identified during this study (from the eight wetlands and Heritage Lake. The trematode sampling completeness analysis revealed that we captured 100% of the dominant species (q = 2), 98% of the typical species (q = 1), and 64% of the overall species diversity (q = 0) (Figure 10A). The rarefaction analysis did not plateau, suggesting that trematode species remain to be discovered at these sites (Figure 10B).

**Figure 10.**
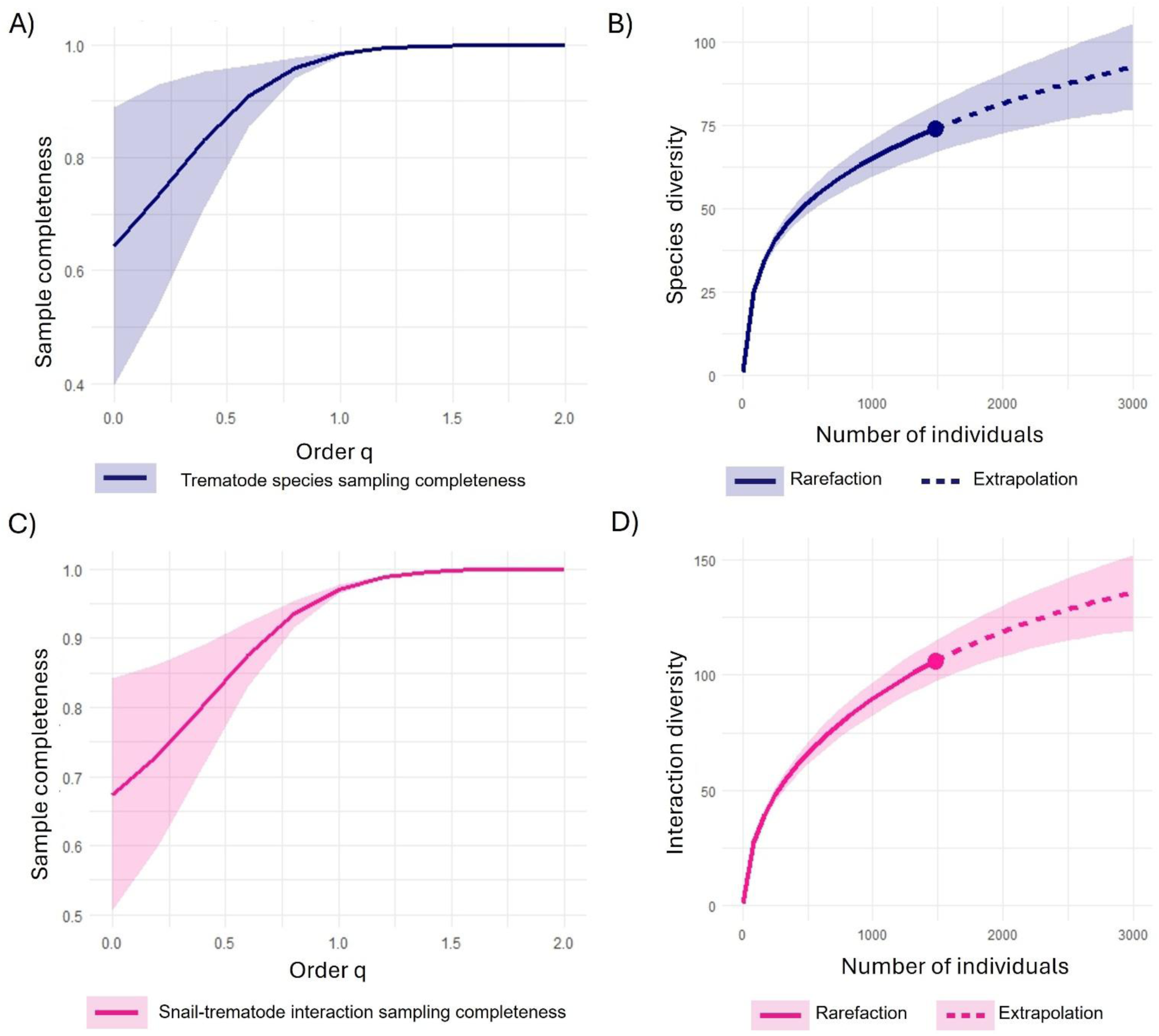
Sample completeness and rarefaction curves for trematode sampling (navy) and snail-trematode interactions (pink). The shaded areas indicated the 95% confidence interval. For the sample completeness profiles (A and C), the levels of q demonstrate: sampling completeness of dominant species (q = 2), typical species (q = 1), and species richness (q = 0). For the rarefaction curves (B and D), the solid line indicates the interpolated values and the dashed lines the rarefaction extrapolation. Data used are either trematodes collected or snail-trematode interactions from all eight wetlands and Heritage Lake across all 4 sampling years. A) Sample completeness profile for all trematodes. B) Rarefaction curve illustrating trematode species diversity. C) Sample completeness profile for all trematode-snail interactions. D) Rarefaction curve illustrating snail-trematode interaction diversity.

Likewise, sampling completeness and rarefaction analyses were conducted for trematode-snail interactions observed during this study, encompassing the eight wetlands and Heritage Lake. The trematode-snail interaction sampling completeness showed that we captured 100% of the dominant interactions (q = 2), 97% of the typical interactions (q = 1), and 69% of the overall interaction diversity (q = 0) (Figure 10C). Again, the rarefaction analysis did not plateau (Figure 10D).

Sampling completeness and rarefaction analyses were also run using combined data from central Alberta, including the eight wetland sites, Heritage Lake from the present study, and six lakes from Gordy and Hanington (2019). The central Alberta trematode sampling completeness revealed that we captured 100% of the dominant species (q = 2), 99% of the typical species (q = 1), and 63% of overall species diversity (q = 0) (Figure 11A). The trematode sample rarefaction analyses corroborated these findings, as it did not reach a plateau (Figure 11B). The sampling completeness for snail-trematode interactions in central Alberta indicated that we captured 100% of the dominant interactions (q = 2), 98% of the typical interactions (q = 1), and 63% of overall interaction diversity (q = 0) (Figure 11C). Again, the rarefaction curve did not plateau, corroborating the findings of the sample completeness analysis (Figure 11D).

**Figure 11.**
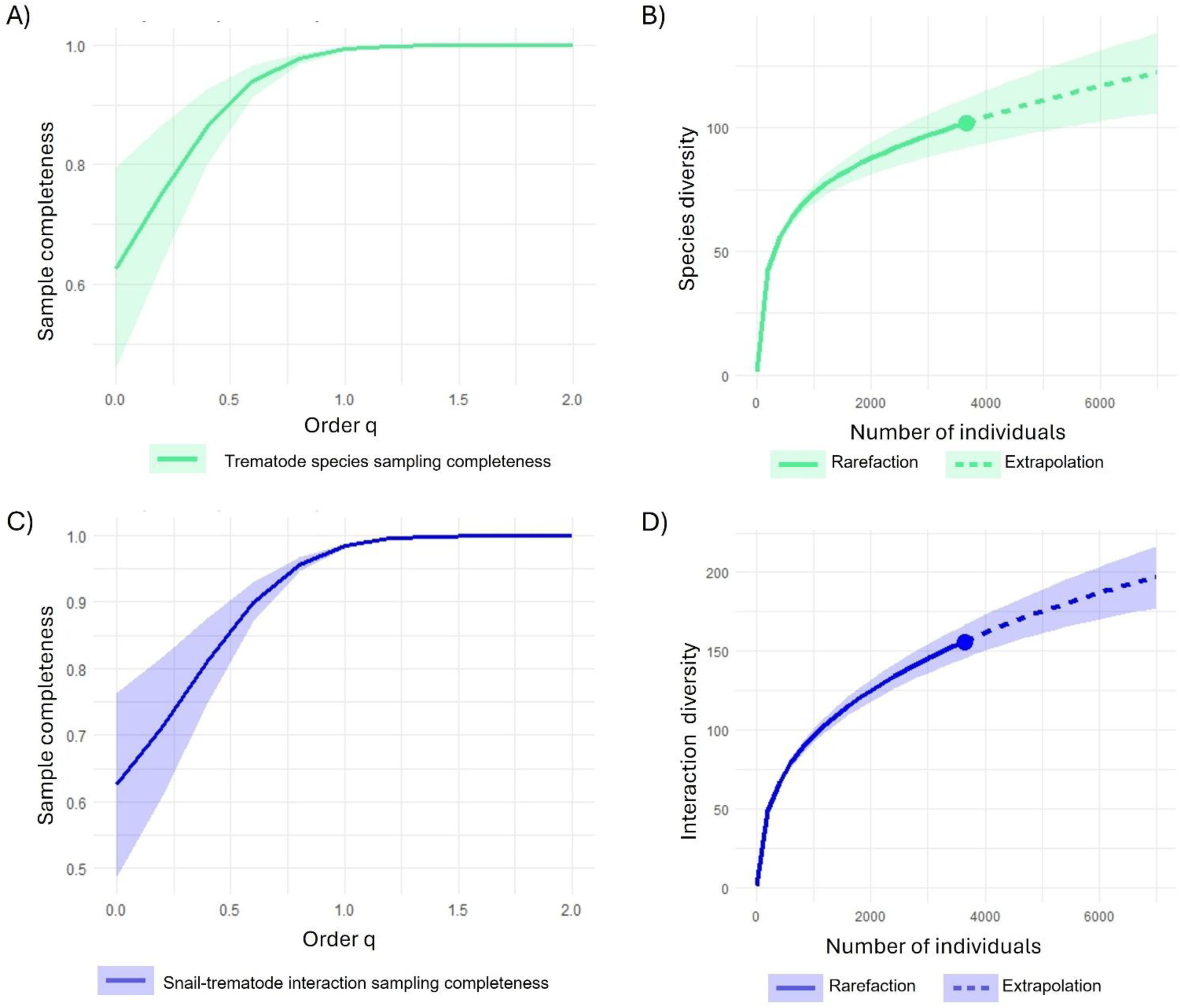
Sample completeness and rarefaction curves for trematode sampling (green) and snail-trematode interactions (blue) in central Alberta. The shaded areas indicated the 95% confidence interval. For the sample completeness profiles (A and C), the levels of q demonstrate: sampling completeness of dominant species (q = 2), typical species (q = 1), and species richness (q = 0). For the rarefaction curves (B and D), the solid line indicates the interpolated values and the dashed lines the rarefaction extrapolation. Data used are either trematodes collected or snail-trematode interactions from collections in central Alberta during this study and previous work in the province (Gordy et al., 2016, 2017, 2018; Gordy and Hanington 2019). A) Sample completeness profile for all trematodes. B) Rarefaction curve illustrating trematode species diversity. C) Sample completeness profile for all trematode-snail interactions. D) Rarefaction curve illustrating snail-trematode interaction diversity.

The sampling completeness analysis for vertebrates observed at the wetland sites indicated that at levels q = 2 (dominant species) and q = 1 (typical species), we captured 100% of the species and 83% of overall species diversity (q = 0) (Supplementary Figure A7). These results combine the bird song recorder and field camera identifications at the wetland sites across 2020-2022.

For the invertebrates, the sampling completeness analysis indicated lower sample completeness overall. We captured 90% at q = 2, 70% at q = 1, and 55% at q = 0 (Supplementary Figure A8). The invertebrate analysis comprises invertebrate species captured using benthic kick netting in 2022 with species identifications from DNA sequencing. This invertebrate data did not include snail or trematode samples. The sampling completeness analyses for the snails revealed that we captured 100% of species diversity for all three levels of q (Supplementary Figure A9). This data comprised all snails collected from the wetlands and Heritage Lake across all four collection years.

### 3.6 Effects of snail species on trematode richness

The assumptions of normality and homogeneity of variances were met for the ANOVA. We found a significant difference in trematode richness among snail species (F3,12 = 3.95, p = 0.036). Tukey’s HSD revealed that the difference between *Pl. trivolvis* and *P. gyrina* was significant (mean difference = −12.50, p = 0.030, CI [−23.88, −1.12]), while the other comparisons were not significant.

### 3.7 Effect of wetland age on trematode species richness

We tested the effect of wetland remediation stage (age) and year on trematode species richness, with site included as a random intercept. We fit a GLMM using a negative binomial distribution. A likelihood ratio test (LRT) determined that both wetland age (p = 0.52) and year (p = 0.22) were not significant to the model, so we did not continue modelling this relationship.

### 3.8 Visualizing host-parasite interactions

When mapping trematode-definitive host information from the literature, the tripartite diagram showed gaps in our life cycle knowledge for many trematode species with many species missing connections to definitive hosts entirely (Figure 12). Notably, this diagram revealed the importance of *L. stagnalis* in these wetlands, as every trematode that parasitized more than one snail species parasitized *L. stagnalis*. Using the identifications of vertebrate species at the study sites, we inferred relationships between 16 trematode species and their numerous potential definitive hosts, demonstrating the magnitude of these relationships in a relatively small geographic area (Supplementary Figure A10).

**Figure 12.**
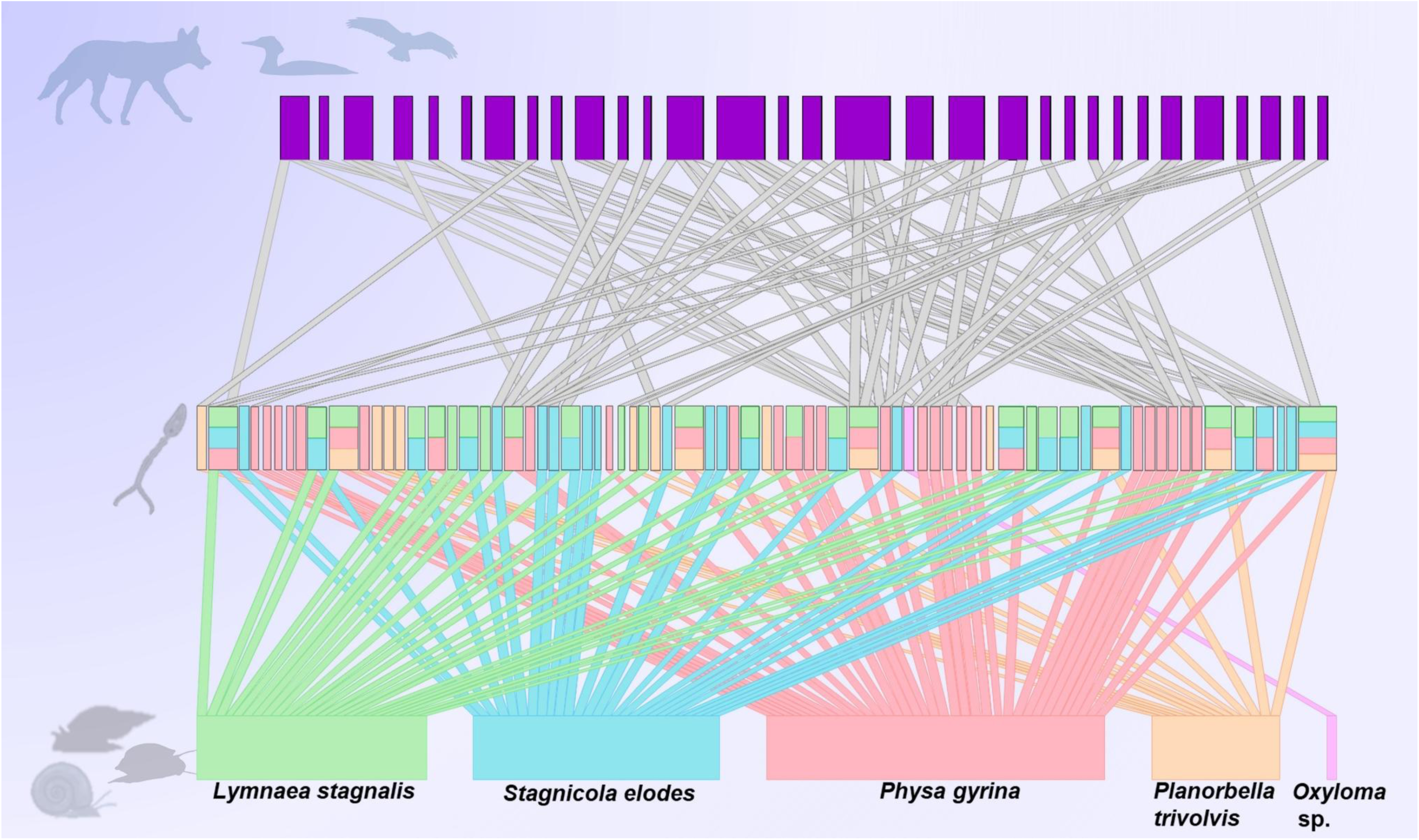
Tripartite diagram of snails, trematodes, and known host species made using the package bipartite (Dormann et al., 2024). The bottom level includes snail hosts, the middle includes trematode species, and the top level includes known definitive hosts, with the lines between each level representing connections between species. The size of each definitive host block is related to the number of connections to trematode species. The size of the block and snail host colours indicates the number of connections between a trematode species and the snail hosts. Host and parasite graphics from BioRender (https://www.biorender.com/) and PhyloPic (https://www.phylopic.org/).

### 3.9 Trematode seasonality

The trematode seasonality plot revealed that species with rediae stages, when present, typically appeared in June and remained throughout the remaining months of the collection season (Figure 13). Species that use sporocysts, however, often did not appear until July, with some appearing for the first time in August (Figure 13). Species with sporocyst stages were more common (56 species) than species with rediae stages (18 species). Trematode infections take approximately one month to reach patency in their snail hosts (Sorenson & Minchella, 1998; Bustinduy & King, 2014), suggesting that trematodes appearing later in the season are deposited by hosts arriving in June and July.

**Figure 13.**
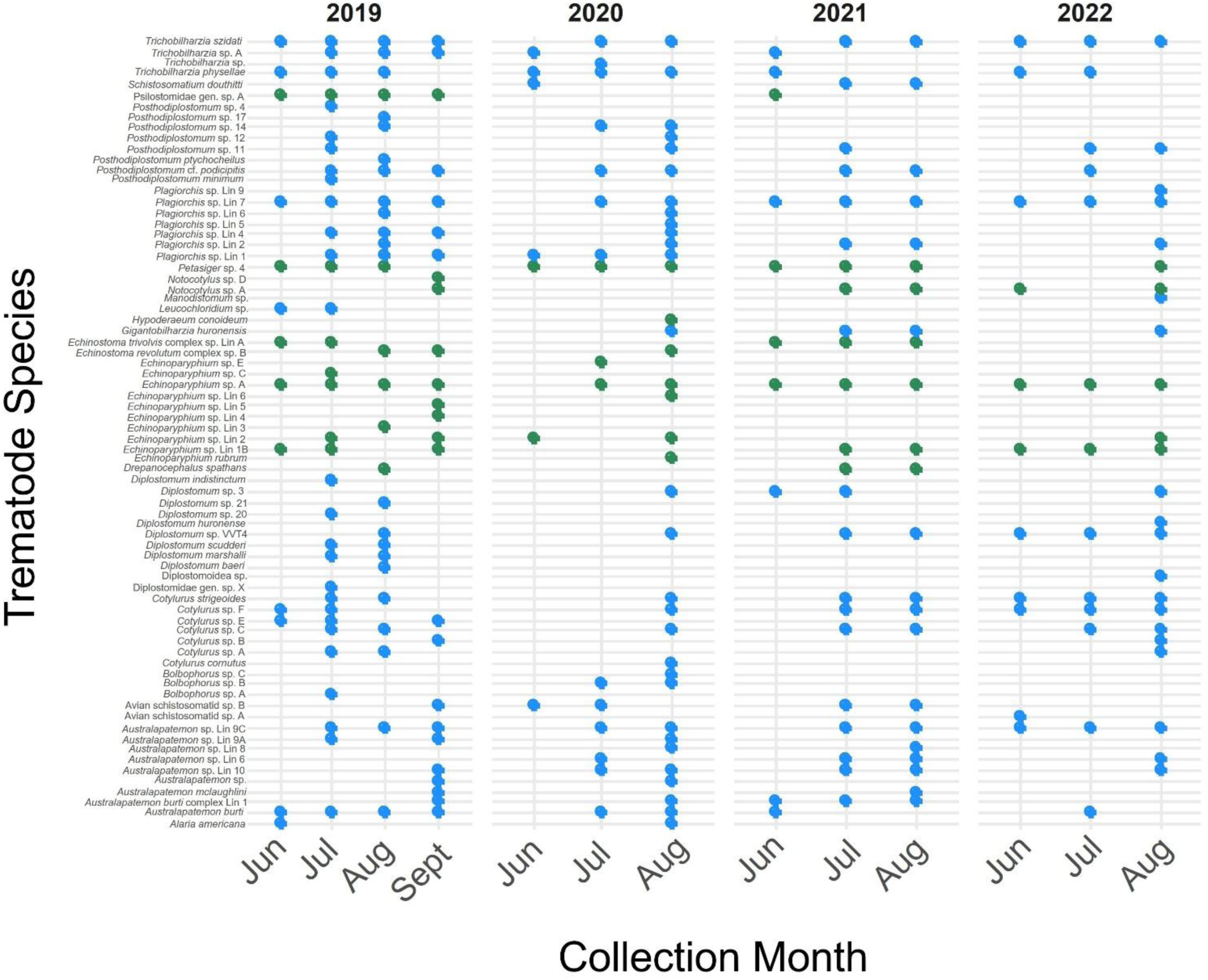
Trematode seasonality plot. Green are trematodes with rediae stages in their life cycles. All trematode species across the eight wetland sites and Heritage Lake. Green represents rediae species, blue represents sporocyst species.

## 4. Discussion

This study identified 73 trematode species, including 23 not previously reported in central Alberta, of which nine are putative novel species, confirmed through phylogenetic analyses. Through sample completeness and rarefaction analyses, we found that there are still trematode species to be discovered in central Alberta. Using traditional biodiversity assessment methods, we identified 137 vertebrate and 46 invertebrate species (excluding the six snail species) in the area, which have the potential to be parasitized by the trematodes present. We have also added new snail first-intermediate host records for 20 trematode species. These findings are important as we continue to add to snail-trematode relationship data in the province and have uncovered several species in a small geographic area.

### 4.1 Trematode and snail-trematode interaction sampling completeness

Gordy and Hanington (2019) identified 79 trematode species in central Alberta, and this study has added 23 additional species, including nine putative novel species. The sample completeness for trematodes identified from all collection sites throughout this study indicated we identified 100% of the dominant species (q = 2), 98% of the typical species (q = 1), and 64% of the overall species richness (q = 0) (Figure 10A), which aligns with the rarefaction results for these collection sites (Figure 10B).

The central Alberta rarefaction analysis revealed that combining the snail collections conducted in central Alberta over the last decade did not fully capture all digenean trematode species and snail-trematode interactions in the area (Figure 11A-D). Gordy and colleagues (Gordy et al., 2016; Gordy & Hanington, 2019) collected 17 447 snails across three years and six lakes and observed 2 452 patent infections. Throughout this study, we collected 22 397 snails across four years, at eight reclaimed wetlands, and Heritage Lake. Our total number of snails collected in central Alberta is 39 844, of which 4 433 (11.13%) were infected with a digenean trematode. From the trematodes collected in central Alberta, 3 657 have been identified, and we have identified 102 unique species combined. Based on the rarefaction results for this combined dataset, it is unlikely that snail collections alone will capture the full trematode diversity of central Alberta. If we nearly doubled our snail collections to 70 000 snails and assumed an infection rate of ∼10%, we would still not reach a plateau based on the rarefaction analyses of trematode species and snail-trematode interactions (Figure 11B, D). It is neither feasible nor a productive use of resources to double snail collections in central Alberta over the next decade.

However, molecular tools such as environmental DNA (eDNA) or cercariometry combined with metabarcoding could help us capture those trematode species that remain undetected in a less laborious, costly, and time-consuming manner (Huver et al., 2015; Sengupta et al., 2019; Alzaylaee et al., 2020).

In central Alberta we have identified 102 trematode species emerging from a handful of snail species. This finding suggests that the estimated 18 000 nominal species within the Digenea (Cribb et al., 2001; Olson et al., 2003) may be conservative. Poulin and Morand (2004) estimate that trematode diversity is 24 401 species. However, Dobson and colleagues (2008) suggest that this number is still too low as tropical vertebrate hosts were under-sampled, and posit that the species diversity of parasitic helminths (trematodes, cestodes, acanthocephalans, and nematodes) is between 75 000 - 300 000. Twelve years later, an updated estimate indicated that trematode diversity was approximately 181 000 species with helminth diversity totalling approximately 348 000 species, accounting for the presence of cryptic species (Carlson et al., 2020). Detecting over one hundred trematode species in a relatively small geographic area with limited first-intermediate host diversity suggests that global trematode diversity was likely underestimated in the early 2000s, and the species identifications and host-parasite relationship discoveries in the decades since have further improved our estimates. A key challenge in assessing trematode diversity is the presence of cryptic species, trematode species that appear morphologically similar but require molecular sequencing for accurate identification (Agapow et al., 2004; Dobson et al., 2008; Poulin, 2011; Blasco-Costa, Cutmore, et al., 2016). Expanding sampling efforts to incorporate eDNA methods across diverse ecosystems will be essential for refining trematode diversity estimates and capturing the full scope of trematode diversity.

### 4.2 Novel first-intermediate host records

During this study, we have added new snail host records for 20 trematode species (Supplementary Table A1). These include the first snail record for species described from their adult stages. Of those with known snail hosts, some parasitized a novel snail within the same family of their known hosts, while others infiltrated a new genus of snails. Those species that remained within the same family include: *Cotylurus* sp. A, *Diplostomum scudderi*, *Hypoderaeum conoideum*, and *Plagiorchis* sp. Lineages 2, 4, and 5 (Supplementary Table A1).

There were also species that were first identified from other hosts in the life cycle, with no snail records, such as *Posthodiplostomum* sp. 12, *Posthodiplostomum* sp. 17 and *Posthodiplostomum* cf. *podicipitis*. The species *Posthodiplostomum* sp. 12, formerly *Ornithodiplostomum* sp. 3 (Achatz et al., 2021), has been previously reported to infect the fish *Pimephales notatus* (Locke, McLaughlin, & Marcogliese, 2010), and was first reported by Moszczynska and colleagues (2009). To our knowledge, no snail host has been previously reported. We found it emerging from *P. gyrina* snails (n = 2) (Table 3, Supplementary Table A1). *Posthodiplostomum* sp. 17 was described by Achatz and colleagues (2021) and infected the hooded merganser (*Lophodytes cucullatus*). This species emerged from a single *P. gyrina* snail during this study (Table 3, Supplementary Table A1). Similarly, *Posthodiplostomum* cf. *podicipitis* was described from a hooded merganser (*Lophodytes cucullatus*) (Achatz et al., 2021) and shared morphological characteristics with the original *Posthodiplostomum podicipitis* described by Yamaguti (1939) in *Tachybaptus ruficollis* (= *Podiceps ruficollis*). We also found *Posthodiplostomum* cf. *podicipitis* emerging from *Physa gyrina* snails (Table 3, Supplementary Table A1).

Finally, there were trematode species that infected snails from different families than their known snail hosts, including *Australapatemon* sp. Lin 6, Avian schistosomatid sp. A, *Cotylurus* sp. B, *Diplostomum indistinctum*, *Echinoparyphium* sp. A, *Echinoparyphium* sp. Lin 1B, *Echinostoma trivolvis* Lin A, *Plagiorchis* sp. Lineages 1 and 7, Psilostomidae gen. sp. A, and *Trichobilharzia szidati*. Of these, some species infected 2 or more novel snail hosts, including *Echinoparyphium* sp. A, *Echinostoma trivolvis* complex Lin A, *Plagiorchis* sp. Lineages 1 and 7, Psilostomidae gen. sp. A, and *T. szidati*, while the others infected only one.

*Echinostoma trivolvis* complex Lin A emerged from *L. stagnalis* (n = 2), *P. gyrina* (n = 1), and *Pl. trivolvis* (n = 11), and had previously been reported from *Pl. trivolvis* snails (Gordy & Hanington, 2019). *Echinoparyphium* sp. Lin 1B was infecting *L. stagnalis* (n = 1) and *P. gyrina* (n = 12), and had been previously reported to infect *P. gyrina* snails (Gordy & Hanington, 2019). Psilostomidae gen. sp. A emerged from *Pl. trivolvis* (n = 18), *L. stagnalis* (n = 1) and *P. gyrina* (n = 2). This species was first discovered by Gordy and colleagues (2016) infecting *Pl. trivolvis* snails (Gordy et al., 2016; Gordy & Hanington, 2019). *Lymnaea stagnalis* and *P. gyrina* are both novel snail records for this species. Lineages 1 and 7 of *Plagiorchis* sp. both infected three snail species. *Plagiorchis* sp. Lin 1 infected *L. stagnalis* (n = 1), *S. elodes* (n = 21), and *P. gyrina* (n = 1), and had previously only been reported from *S. elodes* (Gordy & Hanington, 2019; Kudlai et al., 2021). *Plagiorchis* sp. Lin 7 infected *L. stagnalis* (n = 478), *P. gyrina* (n = 2), and *Pl. trivolvis* (n = 1). This species had only been previously reported from *L. stagnalis* (Gordy & Hanington, 2019; Kudlai et al., 2021), and continued to exhibit a clear preference for *L. stagnalis* throughout this study.

The avian schistosome *T. szidati* was observed to infect four snail species: *L. stagnalis, S. elodes, P. gyrina* and *Pl. trivolvis* (Table 3, Supplementary Table A1). Reports in the literature indicate that *T. szidati* is known to infect species of lymnaeids belonging to the genera *Lymnaea, Stagnicola,* and *Radix* (Špakulová et al., 1996; Kolářová et al., 1997; Brant & Loker, 2009; Loker et al., 2022). We did not find evidence that *T. szidati* has been reported from *P. gyrina* or *Pl. trivolvis* snails. Throughout this study, most *T. szidati* identifications came from *L. stagnalis* snails (n = 191), with one each identified from *S. elodes* and *Pl. trivolvis*, and two from *P. gyrina*. *Trichobilharzia szidati* infections occurring in snails outside of the Lymnaeidae family appear to be unusual and could indicate previously undocumented host generalism, or potential host-switching events.

### 4.3 Phylogenetic analyses

Representative sequences from this study used in phylogenetic analyses were submitted to GenBank (see Supplementary Table A1 for accession numbers). Where possible, two representatives for each gene sequenced were made available. Some species were only observed once, and only a partial sequence was obtained. These partial fragments shortened the alignments used for phylogenetic analyses and were omitted. They have been uploaded to GenBank and included in Supplementary Table A1.

#### 4.3.1 Diplostomidae

Gordy and Hanington (2019) identified *Diplostomum* sp. A and sp. C which Achatz et al. (2022) recently identified as *D. marshalli* and *D. scudderi*, respectively. *Diplostomum marshalli* was shown to infect Greater yellowlegs (*Tringa melanoleuca*) as the definitive host, and *D. scudderi* infected the hooded merganser (*Lophodytes cucullatus*) (Achatz, Martens, et al., 2022). Two putative novel species of *Diplostomum* were identified during this study, both parasitizing *Stagnicola elodes* snails. Several members of the Diplostomidae returned only partial sequence fragments including, *Diplostomum baeri*, Diplostomoidea sp., *Posthodiplostomum minimum,* and *Posthodiplostomum* sp. 17.

There were several publicly available, undescribed *Bolbophorus* sp. sequences deposited in GenBank. We discovered three unique species during this study: *Bolbophorus* sp. A, B, and C. *Bolbophorus* sp. A was the only one that did not form a clade with a previously published specimen of *Bolbophorus* sp. (Figure 4). *Bolbophorus* species A and C have been previously reported in Alberta (Van Steenkiste et al., 2015; Gordy et al., 2016; Gordy & Hanington, 2019). Species belonging to this genus have been reported to parasitize piscivorous birds as the definitive hosts and many fish species as the second intermediate hosts (Overstreet et al., 2002; Terhune et al., 2002; Forrester & Spalding, 2003; Kinsella et al., 2004; Rosser, Baumgartner, et al., 2016).

#### 4.3.2 Echinostomatidae

Two putative novel species of *Echinoparyphium* (Lineages 5 and 6) were discovered parasitizing *P. gyrina* during this study. Sequences were obtained for the *COI* and *nad1* genes to confirm this finding. In both the *COI* and *nad1* trees for the *Echinostoma* species, our representatives of *Echinostoma revolutum* grouped with a representative of *E. revolutum* complex sp. B previously collected in Alberta (Supplementary Table A1) (Gordy and Hanington, 2019). Similarly, our representatives of *E. trivolvis* grouped with *E. trivolvis* complex sp. Lineage A from Alberta (Supplementary Table A1) (Gordy and Hanington, 2019) using both the *COI* and *nad1* sequences.

#### 4.3.3 Leucochloridiidae

Species of *Leucochloridium* are interesting and conspicuous parasites; the pulsating brood sacs make infected snails stand out amongst their uninfected counterparts. In instances where the snail infection prevalence is known, it is often very low (Pojmańska, 1969), which holds true based on the prevalence observed in central Alberta (3 infected snails of 1 096 succineids; 0.27%, all collected from site SC3). The prevalence of *Leucochloridium* spp. infections in snails varies widely in the literature, from 0.7 – 33.3% (Magath, 1920; Woodhead, 1935; Ingram & Hewitt, 1943; McIntosh, 1948; Pojmańska, 1962, 1969a; Lewis Jr., 1974; Bakke, 1978; Yamada & Fukumoto, 2011; Wesołowska & Wesołowski, 2014; Nakao et al., 2019; Chiu et al., 2022), but the prevalence values are difficult to compare between studies because researchers were often searching for infected snails with the pulsating broodsacs, rather than collecting all succineid snails observed in the collection area (Lo & Chen, 1973). Due to the low prevalence, it is not surprising that there were few publicly available sequences.

To our knowledge, only a few published records of *Leucochloridium* species were reported from Canada. In Manitoba, Canada and Arkansas, USA, red-winged black birds (*Agelaius phoeniceus*) were surveyed for parasites, and *L. macrostomum* was found infecting one bird (Hood & Welch, 1980). The authors suggested it was likely infected during the breeding period in Manitoba, Canada (Hood & Welch, 1980). Another study in Manitoba revealed *L. melospizae* infecting the song sparrow (*Melospiza melodia juddi*) (Hodasi, 1963). Eighty-five bird specimens were examined during this study, but no infection prevalence values were reported (Hodasi, 1963). In Quebec, an unidentified species of *Leucochloridium* was found to be infecting *Succinea obliqua* (= *S. putris*; Pilsbry, 1948) snails (Hanam, 1897). No further information was given on this parasite, as the report focused on the snails present in the region. Another study in Quebec collected 34 ring-billed gulls (*Larus delawarensis*) and tagged them with GPS trackers to study their habitat use patterns (Aponte et al., 2014). An unidentified species of *Leucochloridium* was found to be infecting a single gull (Aponte et al., 2014). In New Brunswick, the adult stage of *L. cyanocittae* (= *L. actitis*) was found to be infecting two spotted sandpipers (*Actitis macularius*) (Didyk et al., 2007). This finding was reported along with many other parasites exploiting *A. macularius*, and no further information on the *L. cyanocittae* specimen was provided. Unfortunately, no DNA sequences were made available during these studies mentioned above. To our knowledge, this study provides the first *Leucohloridium* sp. sequence from Canada.

#### 4.3.4 Notocotylidae

There were few sequences available for the phylogenetic analyses of the Notocotylidae that aligned with our *COI* sequences. Other publicly available sequences (GenBank) of *Notocotylus* species included *N. atlanticus* (France; accession # OL629687) (Gonchar et al., 2019; Gonchar & Galaktionov, 2022), *N. magniovatus* (Japan; accession # LC597085) (Sasaki et al., 2021), *Notocotylus* sp. (Japan; accession # LC597061) (Sasaki et al., 2021), *Notocotylus* sp. A (Japan; accession # LC599776) (Nakao & Sasaki, 2021), and *Notocotylus* sp. B (Japan; accession # LC599785) (Nakao & Sasaki, 2021). However, these sequences were not from the same region of the *COI* gene and were not included in the phylogenetic analyses. Our representative of *Notocotylus* sp. A shared approximately 99% similarity to *Notocotylus* sp. A previously identified in Alberta (Supplementary Table B9) (Gordy and Hanington, 2019).

However, our representative of *Notocotylus* sp. D shared approximately 94% similarity with the *Notocotylus* sp. D specimens previously identified in Alberta (Supplementary Table B9) (Gordy and Hanington, 2019). This group requires further taxonomic clarification, which Gordy and Hanington (2019) also noted.

#### 4.3.5 Plagiorchiidae

Plagiorchiidae was the most prevalent family identified during this study (Supplementary Figure A1). Gordy and Hanington (2019) discovered nine lineages of *Plagiorchis* sp. During this study, we observed seven of the nine lineages, and we did not identify any snails infected with *Plagiorchis* sp. Lin 3 or 8. The Plagiorchiidae was the most common family identified using DNA sequences. *Plagiorchis* sp. Lin 7 was the most common parasite identified during this study and parasitized three snail hosts (Table 3). Members of *Plagiorchis* are vectors for Potomac Horse Fever (Vaughan et al., 2012; Greiman et al., 2013), known to have infected horses near Edmonton, Alberta, previously (Gordy & Hanington, 2019). We also identified a putative novel trematode within the genus *Manodistomum* from the family Plagiorchiidae. To our knowledge, this is the first record of this genus in the province. Members of this genus parasitize amphibians as their second intermediate hosts and reptiles as their definitive hosts (Dalzell & Sutherland, 1998; Mihaljevic et al., 2018). Few genetic sequences were available for comparison, and this genus requires further investigation.

#### 4.3.6 Psilostomidae

Within the Psilostomidae phylogenetic tree, one representative sequence identified as *Riberoia ondatrae* from Keller and colleagues (2021) grouped with the Psilostomidae gen. sp. A sequences from this study and previous collections in Alberta (Gordy & Hanington, 2019).

However, this clade was separate from another that contained *R. ondatrae* sequences from Oregon and California (Johnson et al., 2021). Keller and colleagues (2021) collected cercariae from lymnaeid and planorbid snails and metacercariae from two species of salamanders (*Ambystoma californiense*, *Ambystoma macrodactylum croceum*) in California. After sequencing the 28S, *COI* and *ITS2* genes of the parasites collected, 28S sequences originating from a single metacercaria (host: *A. californiense*) and 25 cercariae (host: *Planorbella* sp.) matched at 99.9% to an *R. ondatrae* sequence on GenBank (accession #KT956956), published by Tkach and colleagues (2016) (Keller et al., 2021). Notably, though sequences were obtained for the 28S, *COI*, and *ITS2* genes, phylogenetic analyses were only performed using the 28S gene (Keller et al., 2021). As such, we suggest that these sequences identified as *R. ondatrae* should be identified as Psilostomidae gen. sp. A.

#### 4.3.7 Schistosomatidae

Gordy and Hanington (2019) identified only 20 avian schistosome infections, and another 10 *Schistosomatium douthitti* infections. The avian schistosomes were much more prevalent at the collection sites described herein, with 244 infections identified. However, we observed only three infections of *S. douthitti. Schistosomatium douthitti* was the only mammalian schistosome observed during this study and the previous work in Alberta. This trematode parasitizes muskrats, field mice, and other rodents as the definitive host (Price, 1931; Kagan et al., 1954; Loker et al., 2022).

#### 4.3.8 Strigeidae

Within the Strigeidae II, we identified 83 patent infections of the putative novel species *Australapatemon* sp. Lin 9C from three different snail species throughout this study (Tables 3, 4). It is surprising that this species was not previously identified due to its abundance and ability to infect multiple snail species. Based on other species in this genus, this parasite likely uses anatid birds as definitive hosts and potentially leeches as second intermediate hosts (Stunkard et al., 1941; Drago et al., 2007; Hernández-Mena et al., 2014; Blasco-Costa, Poulin, et al., 2016; Gordy et al., 2017; Calhoun et al., 2020). We also identified a clade including representatives of *Australapatemon* sp. Lineage 6 and *Australapatemon* sp. that requires further clarification.

### 4.4 Community diversity

The community diversity data revealed variation in the trematode community across sites and years. Site SC3 showed particularly low species richness across years, likely because more than 25% of the snails collected there were *Oxyloma* sp. (Table 5). This site produced three infections of *Leucochloridium* sp. and had the highest prevalence of *Oxyloma* sp. snails. Several sites saw a reduction in richness over time, potentially indicating an effect of subtractive sampling at these sites across four years. This trend will be discussed further in a forthcoming publication. Generally, evenness was higher at sites with lower diversity. This suggests more even abundance among species, compared to sites with higher diversity, where evenness was generally lower, suggesting the presence of more dominant species (Table 6).

The effect of snail species on trematode richness was significant. Pairwise comparisons revealed that the only significant comparison was between *Pl. trivolvis* and *P. gyrina* snails, with *. gyrina* supporting a greater trematode richness. Though *L. stagnalis* and *S. elodes* hosted fewer trematode species than *P. gyrina*, these comparisons did not suggest that the lymnaeid snails hosted significantly fewer species.

### 4.5 Effect of wetland age on trematode species richness

We did not find an effect of wetland age on trematode species richness. This is likely due to the proximity of wetland ponds, which could facilitate trematode dispersal and reduce differences in species richness between wetlands of different ages. For example, any avian definitive host landing at one pond could be just as likely to land at any of the other seven nearby ponds. We hypothesize that the trematode composition at each pond is more likely driven by the snail community. During this study, *L. stagnalis* and *P. gyrina* were found at every site at least once. *Lymnaea stagnalis* exhibited the most infections (1,202; Table 3); however, *P. gyrina* hosted the largest diversity of trematodes (36 species; Table 3) with approximately a third as many infections.

### 4.6 Trematode seasonality

The trematode seasonality plot showed more trematode species with sporocyst life cycle types compared to those with rediae life cycle stages. Trematode species that infect marine snails have been shown to produce two castes of rediae: soldiers and reproductive (Hechinger et al., 2011). This phenomenon is known as division of labor. The reproductive rediae are responsible for producing cercariae, while the soldier rediae are responsible for defending the snail host from co-occurring infections (Hechinger et al., 2011). Until recently, the examples of division of labor in trematodes were limited to marine trematodes (Garcia-Vedrenne et al., 2016). Metz and Hechinger (2024) discovered rediae castes in the trematode *Haplorchis pumilio*. *H*. *pumilio* infects the snail *Melanoides tuberculata* as the first intermediate host, fish as the second intermediate host, and finally mammals (including humans) and birds as its definitive host (Metz & Hechinger, 2024). In southern California, *H. pumilio* was the most common parasite of *M. tuberculata* and was found to be a dominant competitor and more likely to infect larger snails (Metz & Hechinger, 2024).

### 4..7 Traditional biodiversity assessments

Identifications from field cameras and recorders were used to compile a list of species present rather than capture abundance data. We obtained approximately 1 700 hours of recordings across three summers from the recorders. Due to the large volume of data, we opted to use a program to analyze the recordings. Over 63 000 identifications resulted from the BirdNet analyses, and all the identifications provided by BirdNet were manually inspected for implausible identifications (any species that were not inhabitants or migratory visitors to western Canada).

Our false positive rate for bird identifications using BirdNet was 3.0%, using a confidence score ≥ 0.9. Although, a higher confidence score could exclude species that were present, and a lower confidence score could include misidentifications (Pérez-Granados, 2023). Additionally, decreasing the confidence score will increase the number of identifications presented, sacrificing precision (Pérez-Granados, 2023). Wood and colleagues (2021) achieved a false-positive rate of approximately 2% using a confidence score of 0.5. The Norwegian Institute for Nature Research (Sethi et al., 2021) suggests that a confidence score of 0.7 - 0.8 would be a suitable range for most studies, as higher confidence thresholds result in fewer identifications, and lower confidence thresholds result in higher false-positive rates. Cole and colleagues (2022) compared a low confidence score (> 0.1) to a high confidence score (> 0.9) on a manually annotated acoustic dataset containing 49 species. This resulted in 90% of species being detected at the low confidence score and 65% at the high confidence score. Ultimately, BirdNet, and automated classifiers should not be used without confirming the output results (Cole et al., 2022; Pérez-Granados, 2023).

Sample completeness analyses indicated that overall sample completeness was lower for invertebrates than vertebrates. This was expected as invertebrates from one sampling season were barcoded. A larger effort was placed into understanding snail-trematode relationships at the study sites. Though cameras and recorders were limited to one spot at each site cluster (Figure 1), vertebrate sample completeness was 100% at q = 2 and q =1, and 83% q= 0.

### 4.8 Study limitations and conclusions

To perform this study, it was necessary to impact these ecosystems. It is likely that while removing snails and trematodes from the environment, we altered the community composition. We acknowledge that this study is based on patent infections, or those infections that have reached maturity in the snail and began shedding. We are certain that a percentage of snails had pre-patent infections and, due to the absence of cercarial shedding, were marked as “uninfected”. However, our approach to identifying snails as infected or uninfected remained consistent across the study, allowing for valid comparisons of infection prevalence among sites and years despite the likelihood of underestimating true infection rates. Previous studies have shown that when comparing infection rates from shedding alone to infection rates from shedding combined with snail dissection, the latter will result in higher infection prevalences (Curtis & Hubbard, 1990; Born-Torrijos et al., 2014).

In conclusion, we identified 74 trematode species from nine sites in central Alberta, increasing our total to 102 trematode species in the area. There is a paucity of molecular sequence data for some trematode families, and many gaps in trematode life cycles remain. We identified nine putative novel trematode species during this study, and made educated inferences as to their definitive hosts. As we continue to identify novel species from larval stages, the goal is to connect these larval stages to metacercaria and adult stages from other researchers to fill in the gaps. Additionally, we have shown that these reclaimed wetland ponds are productive and home to many vertebrate and invertebrate species, without including any fish or plant life. There continues to be trematode species to uncover in central Alberta, and environmental molecular methods will likely be the best tool to identify these species.

## Acknowledgements

We gratefully acknowledge the ACA Grants in Biodiversity (supported by the Alberta Conservation Association). Thank you to the Edmonton Radio Control Society for letting us access the sites via their property and for their interest in the research. We are also grateful to Dr. Monica Ayala-Diaz, Danielle Barry, Christina Bowhay, Veronika Franzova, Dr. Abdullah Gharamah, Dr. Jacob Hambrook, Brittyne Hlavay, Dani Jakovljevic, Charlie Kerr, and Alyssa Turnbull for their help with water, eDNA, and snail collections throughout this research.

## Author Contributions

Conceptualization, B.A.M. and P.C.H.; Methodology, B.A.M., P.C.H.; Software, B.A.M.; Investigation, B.A.M., H.V., S.T., N.D., P.C.H.; Analysis, B.A.M.; Writing—original draft preparation, B.A.M.; Writing—review and editing, B.A.M., H.V., S.T., N.D., P.C.H.; Visualization, B.A.M; Supervision, P.C.H.; Project administration, B.A.M.; Funding acquisition, B.A.M and P.C.H. All authors have read and agreed to the published version of the manuscript.

