## Appendix A for "Snail-trematode dynamics in central Alberta wetlands: A longitudinal survey of infections and interactions"

Supplementary Table A1. Information on sequences used in phylogenies and those with novel snail host records. *L. stagnalis* = *Lymnaea stagnalis*, *S. elodes* = *Stagnicola elodes*, *P. gyrina* = *Physa gyrina*, *Pl. trivolvis* = *Planorbella trivolvis*. Accession numbers are from GenBank unless indicated otherwise (see footnotes).

| Family | Species | Host | Location | Accession #s | | Reference(s) |
|---|---|---|---|---|---|---|
| | | | | *COI* | *nad1* | |
| Brachylaimidae | *Brachylaima asakawai* (out) | *Succinea lauta* | Japan | LC349006 | | (Nakao et al., 2018) |
| Brachylaimidae | *Brachylaima ezohelicis* (out) | *Ezohelix gainesi* | Japan | LC198314 | | (Nakao et al., 2017) |
| Diplostomidae | *Alaria americana* | *Lithobates clamitans* | Quebec, Canada | MH581267 | | (Locke et al., 2018) |
| Diplostomidae | *Alaria americana* | *Pl. trivolvis* | Alberta, Canada | PV600499 PV600521 | | Present study. |
| Diplostomidae | *Alaria* sp. 1 | *Lithobates pipiens* | Quebec, Canada | JF769439 | | (Locke et al., 2011) |
| Diplostomidae | *Alaria* sp. 2 | *Anaxyrus boreas, Lithobates catesbeianus, Pseudacris regilla* | California, USA | JF904536 | | (Locke et al., 2011) |
| Diplostomidae | *Austrodiplostomum ostrowskiae* | *Dorosoma cepedianum* | Alabama, USA | KT728799 | | (Rosser, Alberson, et al., 2016) |
| Diplostomidae | *Bolbophorus damnificus* | *Pl. trivolvis* | Mississippi, USA | KU707937 | | (Rosser, Baumgartner, et al., 2016) |
| Diplostomidae | *Bolbophorus* sp. A[a] | *Pl. trivolvis* | Alberta, Canada | BMPC002-25[d] | | Present study. |
| Diplostomidae | *Bolbophorus* sp. B | *Pl. trivolvis* | Alberta, Canada | PV600517 PV600522 | | Present study. |
| Diplostomidae | *Bolbophorus* sp. C | *Pl. trivolvis* | Alberta, Canada | PV600523 | | Present study. |
| Diplostomidae | *Bolbophorus* sp. | *Pimephales promelas* | Alberta, Canada | KM538081 | | (Van Steenkiste et al., 2015) |
| Diplostomidae | *Bolbophorus* sp. | *Pl. trivolvis* | Alberta, Canada | KT831373 | | (Gordy et al., 2016) |
| Diplostomidae | *Bolbophorus* sp. | *Menidia beryllina* | Mississippi, USA | KU707938 | | (Rosser, Baumgartner, et al., 2016) |

*Continued on next page*

Supplementary Table A1 – *Continued from previous page*

| Family | Species | Host | Location | Accession #s | | Reference(s) |
|---|---|---|---|---|---|---|
| | | | | *COI* | *nad1* | |
| Diplostomidae | *Bolbophorus* sp. | *Pl. trivolvis* | Alberta, Canada | MH368809 | | (Gordy & Hanington, 2019) |
| Diplostomidae | Diplostomidae gen. sp. O | *P. gyrina* | Alberta, Canada | MH368848 | | (Gordy & Hanington, 2019) |
| Diplostomidae | Diplostomidae gen. sp. X | *P. gyrina* | Alberta, Canada | MH368849 | | (Gordy & Hanington, 2019) |
| Diplostomidae | Diplostomidae gen. sp. X | *P. gyrina* | Alberta, Canada | PV600504 | | Present study. |
| Diplostomidae | Diplostomoidea sp. | Physidae sp. | New Mexico, USA | MN834120 | | Unpublished |
| Diplostomidae | Diplostomoidea sp.[b] | *P. gyrina* | Alberta, Canada | BMPC017-25[d] | | Present study. |
| Diplostomidae | *Diplostomum ardeae* | *Ardea herodias* | Quebec, Canada | KR271033 | | (Locke et al., 2015) |
| Diplostomidae | *Diplostomum baeri* | *Radix balthica* | Ireland | OK632477 | | (Faltýnková et al., 2022) |
| Diplostomidae | *Diplostomum baeri*[b] | *S. elodes* | Alberta, Canada | PV600515 | | Present study. |
| Diplostomidae | *Diplostomum gavium* | *S. elodes* | North Dakota, USA | MZ323256 | | (Achatz, Martens, et al., 2022) |
| Diplostomidae | *Diplostomum huronense* | *Larus dominicanus* | Concepción, Chile | MZ323260 | | (Achatz, Martens, et al., 2022) |
| Diplostomidae | *Diplostomum huronense* | *L. stagnalis* | Alberta, Canada | PV600525 | | Present study. |
| Diplostomidae | *Diplostomum indistinctum* | *Recurvirostra americana* | North Dakota, USA | MZ323266 | | (Achatz, Martens, et al., 2022) |
| Diplostomidae | *Diplostomum indistinctum* | *Pl. trivolvis*[c] | Alberta, Canada | PV600502 | | Present study. |
| Diplostomidae | *Diplostomum marshalli* | *Tringa melanoleuca* | North Dakota, USA | MZ323268 | | (Achatz, Martens, et al., 2022) |
| Diplostomidae | *Diplostomum marshalli* | *S. elodes* | Alberta, Canada | PV600510 | | Present study. |
| Diplostomidae | *Diplostomum mergi* | *Gobius niger* | Finland | OR831226 | | (Martinek & Hernández-Orts, 2024) |

*Continued on next page*

Supplementary Table A1 – *Continued from previous page*

| Family | Species | Host | Location | Accession #s COI | Accession #s nad1 | Reference(s) |
|---|---|---|---|---|---|---|
| Diplostomidae | *Diplostomum pseudospathaceum* | *L. stagnalis* | United Kingdom | MH170307 | | (Enabulele et al., 2018) |
| Diplostomidae | *Diplostomum scudderi* | *Lophodytes cucullatus* | North Dakota, USA | MZ323273 | | (Achatz, Martens, et al., 2022) |
| Diplostomidae | *Diplostomum scudderi* | *L. stagnalis*[c], *S. elodes* | Alberta, Canada | PV600507 PV600511 | | Present study. |
| Diplostomidae | *Diplostomum spathaceum* | *Abramis brama* | Russia | PQ461136 | | (Lebedeva et al., 2024) |
| Diplostomidae | *Diplostomum* sp. 1 | *Neogobius melanostomus* | Quebec, Canada | MZ563512, MZ563513 | | (Marcogliese & Locke, 2021) |
| Diplostomidae | *Diplostomum* sp. 2 | *Notropis atherinoides, N. hudsonius, Pimephales notatus, P. promelas* | Canada (Ontario and Quebec) | KR271273 | | (Locke et al., 2015) |
| Diplostomidae | *Diplostomum* sp. 3 | *Micropterus salmoides* | Quebec, Canada | GQ292487 | | (Locke, McLaughlin, Dayanandan, et al., 2010) |
| Diplostomidae | *Diplostomum* sp. 3 | *L. stagnalis* | Alberta, Canada | PV600519 | | Present study. |
| Diplostomidae | *Diplostomum* sp. 4 | *Neogobius melanostomus* | Quebec, Canada | MZ563546 | | (Marcogliese & Locke, 2021) |
| Diplostomidae | *Diplostomum* sp. 6 | *Fundulus diaphanus*, *Pimephales notatus* | Canada (New Brunswick and Quebec) | KR271394 | | (Locke et al., 2015) |
| Diplostomidae | *Diplostomum* sp. 7 | *Notropis hudsonius, Onchorhynchus kisutch, Percopsis omiscomaycus, Pimephales notatus, Salvelinus fontinalis* | Canada (British Columbia, Alberta, Ontario, Quebec, and New Brunswick) | KR271409 | | (Locke et al., 2015) |

*Continued on next page*

| Family | Species | Host | Location | Accession #s | | Reference(s) |
|---|---|---|---|---|---|---|
| | | | | *COI* | *nad1* | |
| Diplostomidae | *Diplostomum* sp. 9 | *Apeltes quadracus, Cottus asper, Gasterosteus aculeatus, G. wheatlandi, Percina caprodes, Pungitius pungitius, Salvelinus fontinalis* | Canada (British Columbia, Quebec, and New Brunswick) | KR271413 | | (Locke et al., 2015) |
| Diplostomidae | *Diplostomum* sp. 10 | *Ambloplites rupestris, Pimephales promelas* | Canada (Quebec and Ontario) | KR271098 | | (Locke et al., 2015) |
| Diplostomidae | *Diplostomum* sp. 12 | *Cottus cognatus* | Canada (Alberta and Ontario); USA (Alaska) | KR271101 | | (Locke et al., 2015) |
| Diplostomidae | *Diplostomum* sp. 13 | *Gasterosteus aculeatus* | Oregon, USA | KR271104 | | (Locke et al., 2015) |
| Diplostomidae | *Diplostomum* sp. 14 | *Synodontis zambezensis* | South Africa | MN808624 | | (Hoogendoorn et al., 2020) |
| Diplostomidae | *Diplostomum* sp. 15 | *Chanodichthys dabryi, Hypophthalmichthys nobilis* | China | KR271127 | | (Locke et al., 2015) |
| Diplostomidae | *Diplostomum* sp. 16 | *Pseudocrenilabrus philander* | South Africa | MN808626 | | (Hoogendoorn et al., 2020) |
| Diplostomidae | *Diplostomum* sp. 17 | *Cottus cognatus* | Yukon, Canada | KR271134 | | (Locke et al., 2015) |
| Diplostomidae | *Diplostomum* sp. 18 | *Cottus asper, C. cognatus* | Canada (Yukon and British Columbia) | KR271139 | | (Locke et al., 2015) |
| Diplostomidae | *Diplostomum* sp. 19 | *Cottus cognatus, C. ricei* | Canada (Ontario); USA (Alaska) | KR271145 | | (Locke et al., 2015) |

*Continued on next page*

Supplementary Table A1 – *Continued from previous page*

| Family | Species | Host | Location | Accession #s COI | Accession #s nad1 | Reference(s) |
|---|---|---|---|---|---|---|
| Diplostomidae | *Diplostomum* sp. 20[a] | *S. elodes* | Alberta, Canada | PV600506 | | Present study. |
| Diplostomidae | *Diplostomum* sp. 21[a] | *S. elodes* | Alberta, Canada | PV600509 PV600512 | | Present study. |
| Diplostomidae | *Diplostomum* sp. A | *S. elodes* | Alberta, Canada | MH368931 | | (Gordy & Hanington, 2019) |
| Diplostomidae | *Diplostomum* sp. B | *Radix auricularia* | Japan | LC599722 | | (Nakao & Sasaki, 2021) |
| Diplostomidae | *Diplostomum* sp. C | *S. elodes* | Alberta, Canada | MH368941 | | (Gordy & Hanington, 2019) |
| Diplostomidae | *Diplostomum* sp. VVT1 | *Umbra limi* | Minnesota, USA | MZ323285 | | (Achatz, Martens, et al., 2022) |
| Diplostomidae | *Diplostomum* sp. VVT2 | *Perca flavescens* | Minnesota, USA | MZ323293 | | (Achatz, Martens, et al., 2022) |
| Diplostomidae | *Diplostomum* sp. VVT3 | *Lantra canadensis* | Wisconsin, USA | MZ323294 | | (Achatz, Martens, et al., 2022) |
| Diplostomidae | *Diplostomum* sp. VVT4 | *L. stagnalis* | Minnesota, USA | MZ323295 | | (Achatz, Martens, et al., 2022) |
| Diplostomidae | *Diplostomum* sp. VVT4 | *L. stagnalis* | Alberta, Canada | PV600524 | | Present study. |
| Diplostomidae | *Diplostomum* sp. VVT5 | *Egretta caerulea* | Mississippi, USA | MZ323296 | | (Achatz, Martens, et al., 2022) |
| Diplostomidae | *Neodiplostomum americanum* | *Megascops asio* | Mississippi, USA | KY851306 | | (Woodyard et al., 2017) |
| Diplostomidae | *Ornithodiplostomum scardinii* (out) | *Scardinius erythrophthalmus* | Czech Republic | KX931425 | | (Stoyanov et al., 2017) |
| Diplostomidae | *Ornithodiplostomum* sp. 1 | Not stated | Ontario, Canada | FJ477208 | | (Moszczynska et al., 2009) |
| Diplostomidae | *Ornithodiplostomum* sp. 2 | Not stated | Quebec, Canada | FJ477210 | | (Moszczynska et al., 2009) |

*Continued on next page*

Supplementary Table A1 – *Continued from previous page*

| Family | Species | Host | Location | Accession #s COI | Accession #s nad1 | Reference(s) |
|---|---|---|---|---|---|---|
| Diplostomidae | *Ornithodiplostomum* sp. 3 | *Pimephales notatus* | Quebec, Canada | MF124280 | | (Locke, McLaughlin, & Marcogliese, 2010; Blasco-Costa & Locke, 2017) |
| Diplostomidae | *Ornithodiplostomum* sp. 8 | *P. gyrina* | Alberta, Canada | MH368944 | | (Gordy & Hanington, 2019) |
| Diplostomidae | *Posthodiplostomum centrarchi* | *Lepomis gibbosus* | Hungary | MN179287 | | (Cech et al., 2020) |
| Diplostomidae | *Posthodiplostomum* cf. *podicipitis* | *Lophodytes cucullatus* | North Dakota, USA | MZ707196 | | (Achatz et al., 2021) |
| Diplostomidae | *Posthodiplostomum* cf. *podicipitis* | *P. gyrina*[c] | Alberta, Canada | PV600518 PV600508 | | Present study. |
| Diplostomidae | *Posthodiplostomum cuticola* | *Neogobius fluviatilis* | Lithuania | PQ560819 | | (Kudlai et al., 2024) |
| Diplostomidae | *Posthodiplostomum minimum* | *Nycticorax nycticorax* | Mississippi, USA | MZ707191 | | (Achatz et al., 2021) |
| Diplostomidae | *Posthodiplostomum minimum*[b] | *P. gyrina* | Alberta, Canada | PV600505 | | Present study. |
| Diplostomidae | *Posthodiplostomum ptychocheilus* | *Mergus merganser* | Minnesota, USA | MZ707201 | | (Achatz et al., 2021) |
| Diplostomidae | *Posthodiplostomum ptychocheilus* | *P. gyrina* | Alberta, Canada | PV600514 | | Present study. |
| Diplostomidae | *Posthodiplostomum* sp. 1 | Not stated | Ontario, Canada | FJ477215 | | (Moszczynska et al., 2009) |
| Diplostomidae | *Posthodiplostomum* sp. 2 | Not stated | Quebec, Canada | FJ477216 | | (Moszczynska et al., 2009) |

*Continued on next page*

Supplementary Table A1 – *Continued from previous page*

| Family | Species | Host | Location | Accession #s | | Reference(s) |
|---|---|---|---|---|---|---|
| | | | | *COI* | *nad1* | |
| Diplostomidae | *Posthodiplostomum* sp. 3 | *Lepomis macrochirus, L. cyanellus, L. megalotis, L. humidis, L. microlophus, L. gulosus, Micropterus punctulatus* | Illinois, USA | MG873405 | | (Boone et al., 2018) |
| Diplostomidae | *Posthodiplostomum* sp. 4 | *Pimephales notatus* | Quebec, Canada | HM064847 | | (Locke, McLaughlin, & Marcogliese, 2010) |
| Diplostomidae | *Posthodiplostomum* sp. 4 | *P. gyrina* | Alberta, Canada | MH368945 | | (Gordy & Hanington, 2019) |
| Diplostomidae | *Posthodiplostomum* sp. 4 | *P. gyrina* | Alberta, Canada | BMPC014-25[d] | | Present study. |
| Diplostomidae | *Posthodiplostomum* sp. 5 | Not stated | Quebec, Canada | FJ477219 | | (Moszczynska et al., 2009) |
| Diplostomidae | *Posthodiplostomum* sp. 7 | *Perca flavescens* | Quebec, Canada | HM064868 | | (Locke, McLaughlin, & Marcogliese, 2010) |
| Diplostomidae | *Posthodiplostomum* sp. 8 | *Micropterus salmoides* | New York, USA | OP071223 | | (Díaz Pernett et al., 2022) |
| Diplostomidae | *Posthodiplostomum* sp. 11 | Unidentified fish | North Dakota, USA | MZ707204 | | (Achatz et al., 2021) |
| Diplostomidae | *Posthodiplostomum* sp. 11 | *P. gyrina* | Alberta, Canada | PV600500 PV600503 | | Present study. |
| Diplostomidae | *Posthodiplostomum* sp. 12 | *P. gyrina*[c] | Alberta, Canada | PV600501 PV600520 | | Present study. |
| Diplostomidae | *Posthodiplostomum* sp. 14 | *P. gyrina* | Alberta, Canada | PV600516 | | Present study. |
| Diplostomidae | *Posthodiplostomum* sp. 17 | *Lophodytes cucullatus* | North Dakota, USA | MZ707205 | | (Achatz et al., 2021) |

*Continued on next page*

| Family | Species | Host | Location | Accession #s COI | Accession #s nad1 | Reference(s) |
|---|---|---|---|---|---|---|
| Diplostomidae | *Posthodiplostomum* sp. 17[b] | *P. gyrina*[c] | Alberta, Canada | PV600513 | | Present study. |
| Diplostomidae | *Posthodiplostomum* sp. 18 | *Pelecanus erythrorhynchos* | Oregon, USA | MZ707208 | | (Achatz et al., 2021) |
| Diplostomidae | *Posthodiplostomum* sp. 19 | *Physa* sp. | Minnesota, USA | MZ707209 | | (Achatz et al., 2021) |
| Diplostomidae | *Posthodiplostomum* sp. 20 | *P. gyrina* | Oregon, USA | MZ707211 | | (Achatz et al., 2021) |
| Diplostomidae | *Posthodiplostomum* sp. 21 | *Jabiru myceteria* | Brazil | MZ707213 | | (Achatz et al., 2021) |
| Diplostomidae | *Posthodiplostomum* sp. 22 | *Tigrisoma lineatum* | Brazil | MZ707216 | | (Achatz et al., 2021) |
| Diplostomidae | *Posthodiplostomum* sp. 23 | *Poecilia reticula* | Puerto Rico | OP071194 | | (Díaz Pernett et al., 2022) |
| Diplostomidae | *Posthodiplostomum* sp. 24 | *Poecilia reticula* | Puerto Rico | OP071203 | | (Díaz Pernett et al., 2022) |
| Diplostomidae | *Posthodiplostomum* sp. 25 | *Dajaus monticola* | Puerto Rico | OP071217 | | (Díaz Pernett et al., 2022) |
| Diplostomidae | *Posthodiplostomum* sp. 26 | *Poecilia reticula* | Puerto Rico | OP071219 | | (Díaz Pernett et al., 2022) |
| Diplostomidae | *Tylodelphys aztecae* | *Tachybaptus dominicus* | Mexico | MK172794 | | (Sereno-Uribe et al., 2019) |
| Diplostomidae | *Tylodelphys clavata* | *Neogobius melanostomus* | Lithuania | PQ560826 | | (Kudlai et al., 2024) |
| Diplostomidae | *Tylodelphys excavata* | *Pelophylax ridibundus* | Ukraine | OL439177 | | (Achatz, Chermak, et al., 2022) |
| Diplostomidae | *Tylodelphys immer* | *Gavia immer* | North Dakota, USA | MZ323303 | | (Achatz, Martens, et al., 2022) |
| Diplostomidae | *Tylodelphys scheuringi* (out) | Not stated | Quebec, Canada | FJ477223 | | (Moszczynska et al., 2009) |
| Echinochasmidae | *Echinochasmus japonicus* (out) | *Homo sapiens* | Vietnam | NC_030518 | | (Le et al., 2016) |

*Continued on next page*

Supplementary Table A1 – *Continued from previous page*

| Family | Species | Host | Location | Accession #s *COI* | Accession #s *nad1* | Reference(s) |
|---|---|---|---|---|---|---|
| Echinostomatidae | *Drepanocephalus mexicanus* | *Nannopterum brasilianus* | Mexico | KY636229 | KY636298 | (Hernández-Cruz et al., 2018) |
| Echinostomatidae | *Drepanocephalus* sp. | *Biomphalaria straminea* | Brazil | KP053256 | KP053264 | (Pinto et al., 2016) |
| Echinostomatidae | *Drepanocephalus spathans* | *Nannopterum brasilianus* | Mexico | KY636234 | | (Hernández-Cruz et al., 2018) |
| Echinostomatidae | *Drepanocephalus spathans* | *Pl. trivolvis* | Alberta, Canada | | MH368951 | (Gordy & Hanington, 2019) |
| Echinostomatidae | *Drepanocephalus spathans* | *Pl. trivolvis* | Alberta, Canada | | PV595372 PV595384 | Present study. |
| Echinostomatidae | *Echinoparyphium aconiatum* | *L. stagnalis* | Russia | NC_067751 | | (Gacad et al., 2023) |
| Echinostomatidae | *Echinoparyphium aconiatum* | *Ampullaceana balthica* | United Kingdom | | ON653264 | (Enabulele et al., 2023) |
| Echinostomatidae | *Echinoparyphium ellisi* | *Anas platyrhynchos* | New Zealand | | KY436406 | (Georgieva et al., 2017) |
| Echinostomatidae | *Echinoparyphium poulini* | *Cygnus atratus* | New Zealand | | KY436404 | (Georgieva et al., 2017) |
| Echinostomatidae | *Echinoparyphium recurvatum* | *Ampullaceana balthica* | United Kingdom | | ON653295 | (Enabulele et al., 2023) |
| Echinostomatidae | *Echinoparyphium rubrum* | *S. elodes* | Alaska, USA | | MZ404658 | (Pantoja et al., 2021) |
| Echinostomatidae | *Echinoparyphium rubrum* | *S. elodes* | Alberta, Canada | | PV595379 | Present study. |
| Echinostomatidae | *Echinoparyphium* sp. | *Radix auricularia* | Japan | | LC599763 | (Nakao & Sasaki, 2021) |
| Echinostomatidae | *Echinoparyphium* sp. | *Bulinus tropicus* | Uganda | | ON970247 | (Hammoud et al., 2022) |
| Echinostomatidae | *Echinoparyphium* sp. | *Bulinus tropicus* | Uganda | | OQ573927, OQ573932 | (Hammoud, 2023) |
| Echinostomatidae | *Echinoparyphium* sp. | *P. acuta* | Indiana, USA | | PQ421432 | Unpublished |
| Echinostomatidae | *Echinoparyphium* sp. A2 | *S. elodes* | Alberta, Canada | MH369232 | | (Gordy & Hanington, 2019) |

*Continued on next page*

| Family | Species | Host | Location | Accession #s COI | Accession #s nad1 | Reference(s) |
|---|---|---|---|---|---|---|
| Echinostomatidae | *Echinoparyphium* sp. A2 | *Pl. trivolvis* | Alberta, Canada | | MH369190 | (Gordy & Hanington, 2019) |
| Echinostomatidae | *Echinoparyphium* sp. A | *P. gyrina* | Alberta, Canada | MH369223 | MH369172 | (Gordy & Hanington, 2019) |
| Echinostomatidae | *Echinoparyphium* sp. A | *L. stagnalis*[c], *P. gyrina, Pl. trivolvis*[c] | Alberta, Canada | PV600975 PV600984 PV600982 | PV595390 PV595392 PV595391 PV595373 | Present study. |
| Echinostomatidae | *Echinoparyphium* sp. C | *S. elodes* | Alberta, Canada | MH369226 | | (Gordy & Hanington, 2019) |
| Echinostomatidae | *Echinoparyphium* sp. C | Unknown | Midwestern USA | | PQ421436 | Unpublished |
| Echinostomatidae | *Echinoparyphium* sp. C | *S. elodes* | Alberta, Canada | PV600978 | | Present study. |
| Echinostomatidae | *Echinoparyphium* sp. D | *S. elodes* | Alberta, Canada | | MH369177 | (Gordy & Hanington, 2019) |
| Echinostomatidae | *Echinoparyphium* sp. E | *S. elodes* | Alberta, Canada | MH369275 | MH369122 | (Gordy & Hanington, 2019) |
| Echinostomatidae | *Echinoparyphium* sp. E | *S. elodes* | Alberta, Canada | PV600990 | | Present study. |
| Echinostomatidae | *Echinoparyphium* sp. 1 | *Valvata macrostoma* | Finland | | MZ404662 | (Pantoja et al., 2021) |
| Echinostomatidae | *Echinoparyphium* sp. 2 | *P. acuta* | Iceland | | MZ404665 | (Pantoja et al., 2021) |
| Echinostomatidae | *Echinoparyphium* sp. Lineage 1 | *Physella acuta* | Indiana, USA | | PQ381495 | Unpublished |
| Echinostomatidae | *Echinoparyphium* sp. Lineage 1A | *S. elodes* | Alberta, USA | MH369305 | | (Gordy & Hanington, 2019) |
| Echinostomatidae | *Echinoparyphium* sp. Lineage 1A | *Physella gyrina* | Alberta, USA | | MH369189 | (Gordy & Hanington, 2019) |
| Echinostomatidae | *Echinoparyphium* sp. Lineage 1B | Unknown | Midwestern USA | | PQ421442 | Unpublished |

*Continued on next page*

Supplementary Table A1 – *Continued from previous page*

| Family | Species | Host | Location | Accession #s COI | Accession #s nad1 | Reference(s) |
|---|---|---|---|---|---|---|
| Echinostomatidae | *Echinoparyphium* sp. Lineage 1B | *P. gyrina, L. stagnalis*[c] | Alberta, Canada | PV600976 PV600980 PV600988 | PV595386 PV595389 | Present study. |
| Echinostomatidae | *Echinoparyphium* sp. Lineage 2 | *S. elodes* | Alberta, Canada | MH369283 | | (Gordy & Hanington, 2019) |
| Echinostomatidae | *Echinoparyphium* sp. Lineage 2 | *L. elodes* | Indiana, USA | | GQ463121 | (Detwiler et al., 2010) |
| Echinostomatidae | *Echinoparyphium* sp. Lineage 2 | *L. stagnalis*, *S. elodes* | Alberta, Canada | PV600981 PV600986 | PV595374 PV595375 | Present study. |
| Echinostomatidae | *Echinoparyphium* sp. Lineage 3 | *Pl. trivolvis* | Alberta, Canada | MH369270 | MH369147 | (Gordy & Hanington, 2019) |
| Echinostomatidae | *Echinoparyphium* sp. Lineage 3 | *Helisoma trivolvis* | Indiana, USA | | GQ463123 | (Detwiler et al., 2010) |
| Echinostomatidae | *Echinoparyphium* sp. Lineage 3 | *Pl. trivolvis* | Alberta, Canada | PV600983 | | Present study. |
| Echinostomatidae | *Echinoparyphium* sp. Lineage 4 | *P. gyrina* | Alberta, Canada | | MH369158 | (Gordy & Hanington, 2019) |
| Echinostomatidae | *Echinoparyphium* sp. Lineage 4 | *P. gyrina* | Alberta, Canada | | PV595376 | Present study. |
| Echinostomatidae | *Echinoparyphium* sp. Lineage 5[a] | *P. gyrina* | Alberta, Canada | PV600989 | PV595377 | Present study. |
| Echinostomatidae | *Echinoparyphium* sp. Lineage 6[a] | *P. gyrina* | Alberta, Canada | PV600991 | PV595382 | Present study. |
| Echinostomatidae | *Echinostoma bolschewense* | *Viviparus acerosus* | Slovakia | | KP065621 | (Georgieva et al., 2014) |
| Echinostomatidae | *Echinostoma bolschewense* | *Dreissena polymorpha* | Russia | MZ572871 | | (Bespalaya et al., 2022) |
| Echinostomatidae | *Echinostoma caproni* | Not stated | Madagascar and Egypt | | AF025837 | (Morgan & Blair, 1998b) |
| Echinostomatidae | *Echinostoma* cf. *friedi* | *Planorbis* sp. | United Kingdom | | AY168937 | (Kostadinova et al., 2003) |

*Continued on next page*

Supplementary Table A1 – *Continued from previous page*

| Family | Species | Host | Location | Accession #s | | Reference(s) |
|---|---|---|---|---|---|---|
| | | | | *COI* | *nad1* | |
| Echinostomatidae | *Echinostoma chankense* | *Rattus norvegicus* | Russia | | MT592856 | (Izrailskaia et al., 2021) |
| Echinostomatidae | *Echinostoma cinetorchis* | *Gallus gallus* | Russia | | MT592855 | (Izrailskaia et al., 2021) |
| Echinostomatidae | *Echinostoma hortense* | Not stated | East Asia | | AF025835 | (Morgan & Blair, 1998b) |
| Echinostomatidae | *Echinostoma hortense* (out) | "Dog" | China | KR062182 | | (Liu et al., 2016) |
| Echinostomatidae | *Echinostoma macrorchis* | *Mesocricetus auratus* | Thailand | | MT982427 | (Butboonchoo et al., 2020) |
| Echinostomatidae | *Echinostoma maldonadoi* | *Stenophysa marmorata* | Brazil | | OQ126877 | (Valadão et al., 2023) |
| Echinostomatidae | *Echinostoma mekongi* | *Homo sapiens* | Cambodia | | MT431433 | (Cho et al., 2020) |
| Echinostomatidae | *Echinostoma miyagawai* | *Anas platyrhynchos domesticus* | Russia | | MT592853 | (Izrailskaia et al., 2021) |
| Echinostomatidae | *Echinostoma nasincovae* | *Planorbarius corneus* | Ireland | | MZ404666 | (Pantoja et al., 2021) |
| Echinostomatidae | *Echinostoma novazealandense* | *Cygnus atratus* | New Zealand | | KY436397 | (Georgieva et al., 2017) |
| Echinostomatidae | *Echinostoma novazealandense* | *Anas platyrhynchos* | New Zealand | | KY436398 | (Georgieva et al., 2017) |
| Echinostomatidae | *Echinostoma paraensei* | *Glyptophysa* sp. | Australia | | AF026282 | (Morgan & Blair, 1998a) |
| Echinostomatidae | *Echinostoma paraensei* | Not stated. | Strain: Biology University of New Mexico | KT008005 | | Unpublished |
| Echinostomatidae | *Echinostoma paraulum* | *L. stagnalis* | Germany | | KP065681 | (Georgieva et al., 2014) |
| Echinostomatidae | *Echinostoma pesudorobustum* | *Gallus gallus* | Brazil | | OK564515 | (Valadão et al., 2022) |
| Echinostomatidae | *Echinostoma revolutum* | *L. stagnalis* | Czech Republic | | KP065646 | (Georgieva et al., 2014) |

*Continued on next page*

Supplementary Table A1 – *Continued from previous page*

| Family | Species | Host | Location | Accession #s COI | Accession #s nad1 | Reference(s) |
|---|---|---|---|---|---|---|
| Echinostomatidae | *Echinostoma revolutum* | *Anas platyrhynchos domesticus* | Bangladesh | | LC224104 | (Mohanta et al., 2019) |
| Echinostomatidae | *Echinostoma revolutum* | *Bithynia siamensis siamensis* | Thailand | | MT815862 | (Butboonchoo et al., 2020) |
| Echinostomatidae | *Echinostoma revolutum* | *Radix rubiginosa* | Thailand | OR030109 | | (Suwancharoen et al., 2023) |
| Echinostomatidae | *Echinostoma revolutum* complex sp. B | *S. elodes* | Alberta, Canada | MH369292 | MH369192 | (Gordy & Hanington, 2019) |
| Echinostomatidae | *Echinostoma revolutum* complex sp. B | *L. stagnalis, S. elodes* | Alberta, Canada | PV600987 PV600985 | PV595378 PV595380 | Present study. |
| Echinostomatidae | *Echinostoma robustum* | *Anas platyrhynchos domesticus* | Bangladesh | | LC224095 | (Mohanta et al., 2019) |
| Echinostomatidae | *Echinostoma* sp. | *Glyptophysa.* sp. | Australia | | AF025833 | (Morgan & Blair, 1998b) |
| Echinostomatidae | *Echinostoma* sp. | *Hydromys chrysogaster* | Australia | | AF026290 | (Morgan & Blair, 1998b) |
| Echinostomatidae | *Echinostoma* sp. | *Anas gracilis* | Australia | | OR257455 | (Ray et al., 2024) |
| Echinostomatidae | *Echinostoma* sp. | Not stated | Ontario, Canada | FJ477201 | | (Moszczynska et al., 2009) |
| Echinostomatidae | *Echinostoma* sp. | "Duck" | China | MZ407836 | | Unpublished |
| Echinostomatidae | *Echinostoma* sp. I | Not stated | Niger | | AF025836 | (Morgan & Blair, 1998b) |
| Echinostomatidae | *Echinostoma* sp. IG | *Myxas glutinosa* | Ireland | | MZ404678 | (Pantoja et al., 2021) |
| Echinostomatidae | *Echinostoma* sp. n. | *Planorbarius corneus* | Czech Republic | | KP065676 | (Georgieva et al., 2014) |
| Echinostomatidae | *Echinostoma* sp. 1 | *P. gyrina* | Alberta, Canada | KT831361 | | (Gordy et al., 2016) |
| Echinostomatidae | *Echinostoma* sp. 2 | *S. elodes* | Alberta, Canada | KT831355 | | (Gordy et al., 2016) |
| Echinostomatidae | *Echinostoma* aff. *trivolvis* | *S. elodes* | Alberta, Canada | KT831367 | | (Gordy et al., 2016) |

*Continued on next page*

Supplementary Table A1 – *Continued from previous page*

| Family | Species | Host | Location | Accession #s COI | Accession #s nad1 | Reference(s) |
|---|---|---|---|---|---|---|
| Echinostomatidae | *Echinostoma trivolvis* | *Ondatra zibethicus* | Wisconsin, USA | | GQ463051 | (Detwiler et al., 2010) |
| Echinostomatidae | *Echinostoma trivolvis* | *Helisoma anceps* | Indiana, USA | | PQ421444 | Unpublished |
| Echinostomatidae | *Echinostoma trivolvis* | *Ondatra zibethicus* | Ontario, Canada | KM538091 | | (Van Steenkiste et al., 2015) |
| Echinostomatidae | *Echinostoma trivolvis* complex sp. Lineage A | *Pl. trivolvis* | Alberta, Canada | MH369271 | MH369219 | (Gordy & Hanington, 2019) |
| Echinostomatidae | *Echinostoma trivolvis* complex sp. Lineage A | *Planorbella pilsbryi* | Wisconsin, USA | | PQ421361 | Unpublished |
| Echinostomatidae | *Echinostoma trivolvis* complex sp. Lineage A | *L. stagnalis*[c], *P. gyrina*[c], *Pl. trivolvis* | Alberta, Canada | PV600977 PV600979 | PV595383 PV595388 PV595387 PV595385 | Present study. |
| Echinostomatidae | *Euparyphium capitaneum* (out) | *Anhinga anhinga* | Mexico | KY636235, KY636236 | | (Hernández-Cruz et al., 2018) |
| Echinostomatidae | *Hypoderaeum conoideum* | *Anas discors* | Manitoba, Canada | KM538101 | | (Van Steenkiste et al., 2015) |
| Echinostomatidae | *Hypoderaeum conoideum* | *Anas platyrhynchos* | Thailand | | MT175434 | (Tantrawatpan & Saijuntha, 2020) |
| Echinostomatidae | *Hypoderaeum conoideum* | *S. elodes*[c] | Alberta, Canada | BMPC009-25[d] | | Present study. |
| Echinostomatidae | *Hypoderaeum* sp. | *S. elodes* | Alberta, Canada | KT831350 | | (Gordy et al., 2016) |
| Echinostomatidae | *Hypoderaeum* sp. Lineage 1 | *L. elodes*, *Gallus gallus* | Indiana, USA | | GQ463099 | (Detwiler et al., 2010) |
| Echinostomatidae | *Hypoderaeum* sp. Lineage 1 | *S. elodes* | Alberta, Canada | | MH369186 | (Gordy & Hanington, 2019) |
| Echinostomatidae | *Isthmiophora melis* (out) | *Planorbis* sp. | United Kingdom | | AY168948 | (Kostadinova et al., 2003) |
| Echinostomatidae | *Neopetasiger islandicus* | *Gyraulus* cf. *parvus* | Iceland | | MZ404686 | (Pantoja et al., 2021) |

*Continued on next page*

Supplementary Table A1 – *Continued from previous page*

| Family | Species | Host | Location | Accession #s COI | Accession #s nad1 | Reference(s) |
|---|---|---|---|---|---|---|
| Echinostomatidae | *Neopetasiger neocommense* | *Podiceps cristatus* | Czech Republic | | JQ425591 | (Georgieva et al., 2012) |
| Echinostomatidae | *Neopetasiger* sp. 5 | *Planorbis planorbis* | Ireland | | MZ404687 | (Pantoja et al., 2021) |
| Echinostomatidae | *Petasiger* sp. 1 | *Radix natalensis* | Kenya | | MK534361 | (Laidemitt et al., 2019) |
| Echinostomatidae | *Petasiger* sp. 2 | *Gyraulus albus* | Germany | | KM191811 | (Selbach et al., 2014) |
| Echinostomatidae | *Petasiger* sp. 2 | *Bulinus globosus* | Kenya | | MK534371 | (Laidemitt et al., 2019) |
| Echinostomatidae | *Petasiger* sp. 3 | *Radix natalensis* | Kenya | | MK534364 | (Laidemitt et al., 2019) |
| Echinostomatidae | *Petasiger* sp. 3 | *Bulinus* sp. | Kenya | | MK534375 | (Laidemitt et al., 2019) |
| Echinostomatidae | *Petasiger* sp. 3 | *Bulinus tropicus* | Uganda | | OQ574002 | (Hammoud, 2023) |
| Echinostomatidae | *Petasiger* sp. 4 | *Pl. trivolvis* | Alberta, Canada | | MH369316 | (Gordy & Hanington, 2019) |
| Echinostomatidae | *Petasiger* sp. 4 | *Biomphalaria pfeifferi* | Kenya | | MK534362 | (Laidemitt et al., 2019) |
| Echinostomatidae | *Petasiger* sp. 4 | *Pl. trivolvis* | Alberta, Canada | | PV595381 PV595393 | Present study. |
| Echinostomatidae | *Petasiger* sp. 5 | *Bulinus ugandae* | Kenya | | MK534411 | (Laidemitt et al., 2019) |
| Echinostomatidae | *Petasiger* sp. 5 | *Bulinus tropicus* | Uganda | | OQ574300 | (Hammoud, 2023) |
| Fasciolidae | *Fasciola hepatica* (out) | *Bos taurus* | Turkey | | LC864563 | Unpublished |
| Haematoloechidae | *Haematoloechus* sp. (out) | *Rana pipiens* | Ontario, Canada | KM538096 | | (Van Steenkiste et al., 2015) |
| Leucochloridiidae | *Leucochloridium paradoxum* | *Succinea lauta* | Japan | LC466790 | | (Nakao et al., 2019) |
| Leucochloridiidae | *Leucochloridium paradoxum* | *Succinea lauta* | Japan | LC466795 | | (Nakao et al., 2019) |

*Continued on next page*

Supplementary Table A1 – *Continued from previous page*

| Family | Species | Host | Location | Accession #s *COI* | *nad1* | Reference(s) |
|---|---|---|---|---|---|---|
| Leucochloridiidae | *Leucochloridium paradoxum* | *Succinea putris* | Russia | MZ676720 | | (Ataev et al., 2023) |
| Leucochloridiidae | *Leucochloridium paradoxum* | *Succinea putris* | Russia | MZ676730 | | (Ataev et al., 2023) |
| Leucochloridiidae | *Leucochloridium paradoxum* | *Succinea putris* | Russia | ON526785 | | (Ataev et al., 2023) |
| Leucochloridiidae | *Leucochloridium paradoxum* | *Succinea putris* | Belarus | ON526803 | | (Ataev et al., 2023) |
| Leucochloridiidae | *Leucochloridium perturbated* | *Succinea lauta* | Japan | LC466772 | | (Nakao et al., 2019) |
| Leucochloridiidae | *Leucochloridium perturbatum* | *Succinea lauta* | Japan | LC466782 | | (Nakao et al., 2019) |
| Leucochloridiidae | *Leucochloridium* sp. | *Succinea lauta* | Japan | LC384429 | | (Ohari et al., 2019) |
| Leucochloridiidae | *Leucochloridium* sp. | *Succinea* sp. | Japan | LC648352 | | (Sasaki et al., 2022) |
| Leucochloridiidae | *Leucochloridium* sp. | *Succinea* sp. | Japan | LC648354 | | (Sasaki et al., 2022) |
| Leucochloridiidae | *Leucochloridium* sp. | *Succinea* sp. | Japan | LC648357 | | (Sasaki et al., 2022) |
| Leucochloridiidae | *Leucochloridium* sp. | *Succinea* sp. | Japan | LC648358 | | (Sasaki et al., 2022) |
| Leucochloridiidae | *Leucochloridium* sp.[a] | *Oxyloma* sp. | Alberta, Canada | PV600055 | | Present study. |
| Leucochloridiidae | *Leucochloridium vogtianum* | *Acrocephalus scirpaceus* | Czech Republic | KP903653 | | (Heneberg et al., 2016) |
| Leucochloridiidae | *Leucochloridium vogtianum* | *Acrocephalus arundinaceus* | Czech Republic | KP903654 | | (Heneberg et al., 2016) |
| Leucochloridiidae | *Leucochloridium vogtianum* | *Locustella fluviatilis* | Czech Republic | KP903656 | | (Heneberg et al., 2016) |
| Leucochloridiidae | *Leucochloridium vogtianum* | *Acrocephalus arundinaceus* | Czech Republic | KP903667 | | (Heneberg et al., 2016) |
| Notocotylidae | Notocotylidae sp. MSB | *Biomphalaria sudanica* | Kenya | KX670216 | | (Laidemett et al., 2017) |
| Notocotylidae | *Notocotylus intestinalis* | *Cygnus atratus* | China | NC_059797 | | (Xu et al., 2021) |
| Notocotylidae | *Notocotylus* sp. A | *P. gyrina* | Alberta, Canada | MH369417 | | (Gordy & Hanington, 2019) |

*Continued on next page*

Supplementary Table A1 – *Continued from previous page*

| Family | Species | Host | Location | Accession #s | | Reference(s) |
|---|---|---|---|---|---|---|
| | | | | *COI* | *nad1* | |
| Notocotylidae | *Notocotylus* sp. A | *P. gyrina* | Alberta, Canada | PV600531 PV600532 | | Present study. |
| Notocotylidae | *Notocotylus* sp. B | *P. gyrina* | Alberta, Canada | MH369416 | | (Gordy & Hanington, 2019) |
| Notocotylidae | *Notocotylus* sp. BOLD | *Mergus merganser* | Quebec, Canada | KM538104 | | (Van Steenkiste et al., 2015) |
| Notocotylidae | *Notocotylus* sp. D | *S. elodes* | Alberta, Canada | MH369414 | | (Gordy & Hanington, 2019) |
| Notocotylidae | *Notocotylus* sp. D | *P. gyrina* | Alberta, Canada | PV600530 | | Present study. |
| Notocotylidae | *Ogmocotyle ailuri* | *Ailurus fulgens* | China | NC_071931 | | (Gao et al., 2023) |
| Notocotylidae | *Ogmocotyle sikae* | Not stated | China | NC_027112 | | Unpublished |
| Notocotylidae | *Ogmocotyle* sp. | Not stated | China | KR006935 | | Unpublished |
| Plagiorchiidae | *Manodistomum* sp. | Not stated | California, USA | HOSL287-19[d] | | (BOLDSYSTEMS, 2018) |
| Plagiorchiidae | *Manodistomum* sp. | Not stated | Missouri, USA | TREMA2456-10[d] | | (BOLDSYSTEMS, 2009) |
| Plagiorchiidae | *Manodistomum* sp.[a] | *P. gyrina* | Alberta, Canada | BMPC016-25[d] | | Present study. |
| Plagiorchiidae | *Plagiorchis elegans* | *L. stagnalis* | Czech Republic | PP387839 | | (Kundid et al., 2024) |
| Plagiorchiidae | *Plagiorchis koreanus* | *Ampullaceana balthica* | Czech Republic | PP387841 | | (Kundid et al., 2024) |
| Plagiorchiidae | *Plagiorchis maculosus* | *Ampullaceana balthica* | Czech Republic | PP387852 | | (Kundid et al., 2024) |
| Plagiorchiidae | *Plagiorchis muelleri* | *Ampullaceana balthica* | Czech Republic | PP387865 | | (Kundid et al., 2024) |
| Plagiorchiidae | *Plagiorchis* sp. | Not stated | Quebec, Canada | FJ477214 | | (Moszczynska et al., 2009) |
| Plagiorchiidae | *Plagiorchis* sp. 3 | *Ampullaceana balthica* | Czech Republic | PP387886 | | (Kundid et al., 2024) |
| Plagiorchiidae | *Plagiorchis* sp. 5 | *Ampullaceana balthica* | Czech Republic | PP387893 | | (Kundid et al., 2024) |

*Continued on next page*

Supplementary Table A1 – *Continued from previous page*

| Family | Species | Host | Location | Accession #s COI | Accession #s nad1 | Reference(s) |
|---|---|---|---|---|---|---|
| Plagiorchiidae | *Plagiorchis* sp. 8 | *Ampullaceana balthica* | Czech Republic | PP387895 | | (Kundid et al., 2024) |
| Plagiorchiidae | *Plagiorchis* sp. 10 | *Ampullaceana balthica* | Poland | PP387898 | | (Kundid et al., 2024) |
| Plagiorchiidae | *Plagiorchis* sp. 11 | *Ampullaceana lagotis* | Czech Republic | PP387900 | | (Kundid et al., 2024) |
| Plagiorchiidae | *Plagiorchis* sp. Lineage 1 | *S. elodes* | Alberta, Canada | MH369420 | | (Gordy & Hanington, 2019) |
| Plagiorchiidae | *Plagiorchis* sp. Lineage 1 | *S. elodes* | Alaska, USA | MW519519 | | (Kudlai et al., 2021) |
| Plagiorchiidae | *Plagiorchis* sp. Lineage 1 | *L. stagnalis*[c], *S. elodes, P. gyrina*[c] | Alberta, Canada | PV600944 PV600933 PV600938 BMPC008-25[d] | | Present study. |
| Plagiorchiidae | *Plagiorchis* sp. Lineage 2 | *S. elodes* | Alberta, Canada | MH369467 | | (Gordy & Hanington, 2019) |
| Plagiorchiidae | *Plagiorchis* sp. Lineage 2 | *L. stagnalis*[c] | Alberta, Canada | PV600945 PV600946 | | Present study. |
| Plagiorchiidae | *Plagiorchis* sp. Lineage 3 | *S. elodes* | Alberta Canada | MH369442 | | (Gordy & Hanington, 2019) |
| Plagiorchiidae | *Plagiorchis* sp. Lineage 4 | *S. elodes* | Alberta, Canada | MH369418 | | (Gordy & Hanington, 2019) |
| Plagiorchiidae | *Plagiorchis* sp. Lineage 4 | *S. elodes* | Alaska, USA | MW519520 | | (Kudlai et al., 2021) |
| Plagiorchiidae | *Plagiorchis* sp. Lineage 4 | *L. stagnalis*[c], *S. elodes* | Alberta, Canada | PV600942 PV600943 PV600934 | | Present study. |
| Plagiorchiidae | *Plagiorchis* sp. Lineage 5 | *S. elodes* | Alberta, Canada | MH369419 | | (Gordy & Hanington, 2019) |
| Plagiorchiidae | *Plagiorchis* sp. Lineage 5 | *L. stagnalis*[c], *S. elodes* | Alberta, Canada | PV600940 PV600939 | | Present study. |

*Continued on next page*

Supplementary Table A1 – *Continued from previous page*

| Family | Species | Host | Location | Accession #s COI | Accession #s nad1 | Reference(s) |
|---|---|---|---|---|---|---|
| Plagiorchiidae | *Plagiorchis* sp. Lineage 6 | *Pl. trivolvis* | Alberta, Canada | MH369470 | | (Gordy & Hanington, 2019) |
| Plagiorchiidae | *Plagiorchis* sp. Lineage 6 | *S. elodes* | Alberta, Canada | PV600936 PV600941 | | Present study. |
| Plagiorchiidae | *Plagiorchis* sp. Lineage 7 | *L. stagnalis* | Alberta, Canada | MH369458 | | (Gordy & Hanington, 2019) |
| Plagiorchiidae | *Plagiorchis* sp. Lineage 7 | *L. stagnalis* | Alaska, USA | MW519521 | | (Kudlai et al., 2021) |
| Plagiorchiidae | *Plagiorchis* sp. Lineage 7 | *L. stagnalis, P. gyrina*[c], *Pl. trivolvis*[c], | Alberta, Canada | PV600935 PV600937 BMPC013-25[d] BMPC015-25[d] | | Present study. |
| Plagiorchiidae | *Plagiorchis* sp. Lineage 8 | *S. elodes* | Alberta, Canada | MH369449 | | (Gordy & Hanington, 2019) |
| Plagiorchiidae | *Plagiorchis* sp. Lineage 9 | *S. elodes* | Alberta, Canada | MH369424 | | (Gordy & Hanington, 2019) |
| Plagiorchiidae | *Plagiorchis* sp. Lineage 9 | *S. elodes* | Alaska, USA | MW519522 | | (Kudlai et al., 2021) |
| Plagiorchiidae | *Plagiorchis* sp. Lineage 9 | *S. elodes* | Alberta, Canada | BMPC014-25[d] | | Present study. |
| Plagiorchiidae | *Plagiorchis vespertilionis* | *Ampullaceana balthica* | Czech Republic | PP387871 | | (Kundid et al., 2024) |
| Proterodiplostomidae | *Crocodilicola pseudostoma* (out) | *Rhamdia guatemalensis* | Mexico | MF398318 | | (Hernández-Mena et al., 2017) |
| Psilostomidae | *Pseudopsilostoma varium* | *Phalacrocorax auritus* | Mississippi, USA | JX468064 | | (O'Hear et al., 2014) |
| Psilostomidae | Psilostomidae gen. sp. A | *Pl. trivolvis* | Alberta, Canada | MH369475, MH369477 | | (Gordy & Hanington, 2019) |

*Continued on next page*

Supplementary Table A1 – *Continued from previous page*

| Family | Species | Host | Location | Accession #s | | Reference(s) |
|---|---|---|---|---|---|---|
| | | | | *COI* | *nad1* | |
| Psilostomidae | Psilostomidae gen. sp. A | *L. stagnalis*[c], *P. gyrina*[c], *Pl. trivolvis* | Alberta, Canada | PV600527 PV600528 PV600526 PV600529 | | Present study. |
| Psilostomidae | *Ribeiroia ondatrae* | *Pl. trivolvis* | California, USA | MW042972 | | (Johnson et al., 2021) |
| Psilostomidae | *Ribeiroia ondatrae* | *Pl. trivolvis* | Oregon, USA | MW042976 | | (Johnson et al., 2021) |
| Psilostomidae | *Ribeiroia ondatrae* | *Ambystoma californiense* | California, USA | OK210492 | | (Keller et al., 2021) |
| Psilostomidae | *Sphaeridiotrema globulus* | *Elimia virginica, Anas platyrhynchos* | New Jersey, US | GQ890329 | | (Bergmame et al., 2011) |
| Schistosomatidae | *Allobilharzia visceralis* | *Cygnus columbianus* | Nevada, USA | EF114219 | | (Brant, 2007) |
| Schistosomatidae | *Anserobilharzia brantae* | *Anser anser* | France | KC570956 | | (Brant et al., 2013) |
| Schistosomatidae | Avian schistosomatid sp. A | *P. gyrina* | Alberta, Canada | MH168789 | | (Gordy et al., 2018) |
| Schistosomatidae | Avian schistosomatid sp. A | *L. stagnalis*[c], *P. gyrina* | Alberta, Canada | PV606416 | | Present study. |
| Schistosomatidae | Avian schistosomatid sp. B | *P. gyrina* | Alberta, Canada | MH168785 | | (Gordy et al., 2018) |
| Schistosomatidae | Avian schistosomatid sp. B | *P. gyrina* | Alberta, Canada | PV606410 BMPC012-25[d] | | Present study. |
| Schistosomatidae | Avian schistosomatid sp. C | *Branta canadensis* | Michigan, USA | MW815104 | | (McPhail et al., 2021) |
| Schistosomatidae | *Dendritobilharzia pulverulenta* | *Gallus gallus* | New Mexico, USA | AY157187 | | (Lockyer et al., 2003) |
| Schistosomatidae | *Gigantobilharzia huronensis* | *Zenaida macroura* | Arizona, USA | KF738949 | | (Sweazea et al., 2015) |
| Schistosomatidae | *Gigantobilharzia huronensis* | *P. gyrina* | Alberta, Canada | PV606411 PV606413 | | Present study. |

*Continued on next page*

Supplementary Table A1 – *Continued from previous page*

| Family | Species | Host | Location | Accession #s COI | Accession #s nad1 | Reference(s) |
|---|---|---|---|---|---|---|
| Schistosomatidae | *Heterobilharzia americana* | *Mesocricetus auratus* | Louisiana, USA | AY157192 | | (Lockyer et al., 2003) |
| Schistosomatidae | *Nasusbilharzia melancorphya* | *Cygnus melancoryphus* | Argentina | MW012493 | | (Flores et al., 2021) |
| Schistosomatidae | *Schistosoma bovis* (out) | *Mus musculus* | Tanzania | AY157212 | | (Lockyer et al., 2003) |
| Schistosomatidae | *Schistosoma indicum* | *Bos taurus* | Bangladesh | AY157204 | | (Lockyer et al., 2003) |
| Schistosomatidae | *Schistosoma nasale* | *Capra hircus* | Sri Lanka | AY157205 | | (Lockyer et al., 2003) |
| Schistosomatidae | Schistosomatidae sp. (out) | *Chilina dombeiana* | Argentina | KC113073 KC113076 | | (Flores et al., 2015) |
| Schistosomatidae | *Schistomatium douthitti* | *Mesocricetus auratus* | Indiana, USA | AY157193 | | (Lockyer et al., 2003) |
| Schistosomatidae | *Schistomatium douthitti* | *L. stagnalis* | Alberta, Canada | MH168791 | | (Gordy et al., 2018) |
| Schistosomatidae | *Schistomatium douthitti* | *L. stagnalis, S. elodes* | Alberta, Canada | PV606414 | | Present study. |
| Schistosomatidae | *Trichobilharzia franki* | *Anas platyrhynchos* | Hungary | MZ562966 | | (Juhász et al., 2022) |
| Schistosomatidae | *Trichobilharzia mergi* | *Mergus serrator* | Iceland | JX456172 | | (Kolářová et al., 2013) |
| Schistosomatidae | *Trichobilharzia novaseelandiae* | *Austropeplea tomentosa* | New Zealand | OK357974 | | (Davis et al., 2021) |
| Schistosomatidae | *Trichobilharzia physellae* | *Anas platyrhynchos* | Michigan, USA | MK433251 | | (Rudko et al., 2019) |
| Schistosomatidae | *Trichobilharzia physellae* | *S. elodes, P. gyrina* | Alberta, Canada | PV606406 PV606408 PV606412 | | Present study. |
| Schistosomatidae | *Trichobilharzia querquedulae* | *Anas versicolor* | Argentina | KU057184 | | (Ebbs et al., 2016) |
| Schistosomatidae | *Trichobilharzia regenti* | *Anas platyrhynchos* | Iceland | HM439503 | | (Jouet et al., 2010) |

*Continued on next page*

Supplementary Table A1 – *Continued from previous page*

| Family | Species | Host | Location | Accession #s | | Reference(s) |
|---|---|---|---|---|---|---|
| | | | | *COI* | *nad1* | |
| Schistosomatidae | *Trichobilharzia* sp.[a,b] | *S. elodes* | Alberta, Canada | PV606409 | | Present study. |
| Schistosomatidae | *Trichobilharzia* sp. A | *Anas americana* | Alaska, USA | FJ174527 | | (Brant & Loker, 2009) |
| Schistosomatidae | *Trichobilharzia* sp. A | *S. elodes* | Alberta, Canada | PV606407 BMPC004-25[d] | | Present study. |
| Schistosomatidae | *Trichobilharzia* sp. B | *Anas americana* | Alaska, USA | FJ174528 | | (Brant & Loker, 2009) |
| Schistosomatidae | *Trichobilharzia* sp. C | *Lophodytes cucullatus* | Pennsylvania, USA | FJ174529 | | (Brant & Loker, 2009) |
| Schistosomatidae | *Trichobilharzia* sp. D | *Stagnicola* sp. | Canada | FJ174485 | | (Brant & Loker, 2009) |
| Schistosomatidae | *Trichobilharzia* sp. E | *Anas acuta* | Canada | FJ174487 | | (Brant & Loker, 2009) |
| Schistosomatidae | *Trichobilharzia stagnicolae* | *Mergus merganser* | Michigan, USA | MK433252 | | (Rudko et al., 2019) |
| Schistosomatidae | *Trichobilharzia szidati* | *L. stagnalis* | Czech Republic | AY157191 | | (Lockyer et al., 2003) |
| Schistosomatidae | *Trichobilharzia szidati* | *L. stagnalis, S. elodes*[c]*, P. gyrina*[c]*, Pl. trivolvis*[c] | Alberta, Canada | PV606415 PV606417 PV606418 BMPC001-25[d] BMPC011-25[d] | | Present study. |
| Strigeidae | *Apatemon* sp. '*jamiesoni*' | *Gobiomorphus cotidianus* | New Zealand | KT334182 | | (Blasco-Costa, Poulin, et al., 2016) |
| Strigeidae | *Apatemon* sp. A | *S. elodes* | Alberta, Canada | MH369617 | | (Gordy & Hanington, 2019) |
| Strigeidae | *Apatemon* sp. B | *S. elodes* | Alberta, Canada | MH369618 | | (Gordy & Hanington, 2019) |
| Strigeidae | *Apatemon* sp. C | *S. elodes* | Alberta, Canada | MH369622 | | (Gordy & Hanington, 2019) |

*Continued on next page*

Supplementary Table A1 – *Continued from previous page*

| Family | Species | Host | Location | Accession #s | | Reference(s) |
|---|---|---|---|---|---|---|
| | | | | *COI* | *nad1* | |
| Strigeidae | *Apharyngostrigea pipientis* (out) | *Ardea alba* | Tanzania | MT943778 | | (Locke et al., 2021) |
| Strigeidae | *Australapatemon burti* | *S. elodes* | Alberta, Canada | KY207619 | | (Gordy et al., 2017) |
| Strigeidae | *Australapatemon burti* | *L. stagnalis, S. elodes, P. gyrina* | Alberta, Canada | PV601149 PV601161 | | Present study. |
| Strigeidae | *Australapatemon burti* complex sp. LIN 1 | *S. elodes* | Alberta, Canada | MH369763 | | (Gordy & Hanington, 2019) |
| Strigeidae | *Australapatemon burti* complex sp. Lineage 1 | *S. elodes* | Alberta, Canada | PV601151 BMPC005-25[d] | | Present study. |
| Strigeidae | *Australapatemon mclaughlini* | *P. gyrina* | Alberta, Canada | KY207615 | | (Gordy et al., 2017) |
| Strigeidae | *Australapatemon mclaughlini* | *P. gyrina* | Alberta, Canada | PV601168 BMPC007-25[d] | | Present study. |
| Strigeidae | *Australapatemon* sp. | *P. gyrina* | Alberta, Canada | KY207613 | | (Gordy et al., 2017) |
| Strigeidae | *Australapatemon* sp. | *P. gyrina* | Alberta, Canada | KY207616 | | (Gordy et al., 2017) |
| Strigeidae | *Australapatemon* sp. | *P. gyrina* | Alberta, Canada | PV601155 PV601160 | | Present study. |
| Strigeidae | *Australapatemon* sp. Lin 3 | *S. elodes* | Alberta, Canada | KY207577 | | (Gordy et al., 2017) |
| Strigeidae | *Australapatemon* sp. Lin 4 | *P. gyrina* | Alberta, Canada | MH369765 | | (Gordy & Hanington, 2019) |
| Strigeidae | *Australapatemon* sp. Lin 5 | *S. elodes* | Alberta, Canada | KY207597 | | (Gordy et al., 2017) |
| Strigeidae | *Australapatemon* sp. Lin 6 | *P. gyrina* | Alberta, Canada | MH369770 | | (Gordy & Hanington, 2019) |
| Strigeidae | *Australapatemon* sp. Lin 6 | *L. stagnalis*[c], *P. gyrina* | Alberta, Canada | PV601162 PV601170 PV601173 | | Present study. |

*Continued on next page*

Supplementary Table A1 – *Continued from previous page*

| Family | Species | Host | Location | Accession #s | | Reference(s) |
|---|---|---|---|---|---|---|
| | | | | *COI* | *nad1* | |
| Strigeidae | *Australapatemon* sp. Lin 8 | *P. gyrina* | Alberta, Canada | MH369777 | | (Gordy & Hanington, 2019) |
| Strigeidae | *Australapatemon* sp. Lin 8 | *P. gyrina* | Alberta, Canada | PV601158 PV601169 | | Present study. |
| Strigeidae | *Australapatemon* sp. Lin 9A | *S. elodes* | Alberta, Canada | MH369787 | | (Gordy & Hanington, 2019) |
| Strigeidae | *Australapatemon* sp. Lin 9A | *L. stagnalis, S. elodes* | Alberta, Canada | PV601150 BMPC006-25[d] | | Present study. |
| Strigeidae | *Australapatemon* sp. Lin 9B | *S. elodes* | Alberta, Canada | MH369791 | | (Gordy & Hanington, 2019) |
| Strigeidae | *Australapatemon* sp. Lin 9C[a] | *L. stagnalis, P. gyrina, Pl. trivolvis* | Alberta, Canada | PV601159 PV601166 | | Present study. |
| Strigeidae | *Australapatemon* sp. Lin 10 | *P. gyrina* | Alberta, Canada | MH369793 | | (Gordy & Hanington, 2019) |
| Strigeidae | *Australapatemon* sp. Lin 10 | *P. gyrina* | Alberta, Canada | PV601156 PV601164 | | Present study. |
| Strigeidae | *Cardiocephaloides medioconiger* | *Thalasseus maximus* | Mississippi, USA | MN817946 | | (Achatz et al., 2020) |
| Strigeidae | *Cardiocephaloides physalis* | *Spheniscus magellanicus* | Chile | MN817947 | | (Achatz et al., 2020) |
| Strigeidae | *Cotylurus cornutus* | *S. elodes* | Alberta, Canada | MH369544 | | (Gordy & Hanington, 2019) |
| Strigeidae | *Cotylurus cornutus*[b] | *S. elodes* | Alberta, Canada | BMPC010-25[d] | | Present study. |
| Strigeidae | *Cotylurus flabelliformis* | *Aythya valisneria* | Manitoba, Canada | MH581275 | | (Locke et al., 2018) |
| Strigeidae | *Cotylurus* sp. A | *S. elodes* | Alberta, Canada | MH369602 | | (Gordy & Hanington, 2019) |
| Strigeidae | *Cotylurus* sp. A | *L. stagnalis, S. elodes* | Alberta, Canada | PV601146 PV601148 | | Present study. |
| Strigeidae | *Cotylurus* sp. B | *P. gyrina* | Alberta, Canada | MH369586 | | (Gordy & Hanington, 2019) |

*Continued on next page*

Supplementary Table A1 – *Continued from previous page*

| Family | Species | Host | Location | Accession #s COI | Accession #s nad1 | Reference(s) |
|---|---|---|---|---|---|---|
| Strigeidae | *Cotylurus* sp. B | *L. stagnalis*[c], *P. gyrina* | Alberta, Canada | PV601153 PV601154 PV601174 | | Present study. |
| Strigeidae | *Cotylurus* sp. C | *L. stagnalis* | Alberta, Canada | MH369566 | | (Gordy & Hanington, 2019) |
| Strigeidae | *Cotylurus* sp. C | *L. stagnalis* | Alberta, Canada | PV601157 PV601165 | | Present study. |
| Strigeidae | *Cotylurus* sp. D | *P. gyrina* | Alberta, Canada | MH369592 | | (Gordy & Hanington, 2019) |
| Strigeidae | *Cotylurus* sp. E | *S. elodes* | Alberta, Canada | MH369565 | | (Gordy & Hanington, 2019) |
| Strigeidae | *Cotylurus* sp. E | *L. stagnalis, S. elodes* | Alberta, Canada | PV601147 PV601152 | | Present study. |
| Strigeidae | *Cotylurus* sp. F | *L. stagnalis* | Alberta, Canada | MH369570 | | (Gordy & Hanington, 2019) |
| Strigeidae | *Cotylurus* sp. F | *L. stagnalis* | Alberta, Canada | PV601163 PV601171 | | Present study. |
| Strigeidae | *Cotylurus strigeoides* | *P. gyrina* | Alberta, Canada | MH369599 | | (Gordy & Hanington, 2019) |
| Strigeidae | *Cotylurus strigeoides* | *Aythya collaris* | Manitoba, Canada | MH581282 | | (Locke et al., 2018) |
| Strigeidae | *Cotylurus strigeoides* | *L. stagnalis, P. gyrina* | Alberta, Canada | PV601167 PV601172 | | Present study. |
| Strigeidae | *Ichthyocotylurus pileatus* | Not stated | Ontario, Canada | FJ477204 | | (Moszczynska et al., 2009) |
| Strigeidae | *Ichthyocotylurus* sp. 2 | Not stated | Quebec, Canada | FJ477205 | | (Moszczynska et al., 2009) |
| Strigeidae | *Ichthyocotylurus* sp. 3 | *Ambloplites rupestris* | Quebec, Canada | HM064731 | | (Locke, McLaughlin, & Marcogliese, 2010) |

[a] indicates a putative novel species discovered during this study
[b] indicates a partial sequence fragment too short to include in phylogenetic analyses
[c] indicates a novel snail host record
[d] indicates an accession number from BOLD

Supplementary Table A2. Vertebrates detected at the wetland sites from either field cameras or recorders.

| Group | Animal (common name) | Suspected species name | Camera | Recorder |
|---|---|---|---|---|
| | | **Amphibians** | | |
| Anurans | American bullfrog | *Lithobates catesbeianus* | | ✓ |
| | | **Birds** | | |
| Anatids | American wigeon | *Mareca americana* | ✓ | ✓ |
| | Blue-winged teal | *Anas discors* | ✓ | ✓ |
| | Bufflehead | *Bucephala albeola* | ✓ | ✓ |
| | Cackling goose | *Branta hutchinsii* | | ✓ |
| | Canada goose | *Branta canadensis* | ✓ | ✓ |
| | Cinnamon teal | *Anas cyanoptera* | ✓ | |
| | Common goldeneye | *Bucephala clangula* | ✓ | ✓ |
| | Common merganser | *Mergus merganser* | ✓ | ✓ |
| | Gadwall | *Mareca strepera* | ✓ | ✓ |
| | Greater scaup | *Aythya marila* | ✓ | |
| | Greater white-fronted goose | *Anser albifrons* | | ✓ |
| | Green-winged teal | *Anas carolinensis* | ✓ | ✓ |
| | Hooded merganser | *Lophodytes cucullatus* | | ✓ |
| | Lesser scaup | *Aythya affinis* | ✓ | |
| | Mallard | *Anas platyrhynchos* | ✓ | ✓ |
| | Northern shoveler | *Spatula clypeata* | ✓ | ✓ |
| | Redhead | *Aythya americana* | ✓ | |
| | Ring-necked duck | *Aythya collaris* | ✓ | ✓ |
| | Ruddy duck | *Oxyura jamaicensis* | ✓ | ✓ |
| | Snow goose | *Anser caerulescens* | | ✓ |
| | Trumpeter swan | *Cygnus buccinator* | | ✓ |
| | Tufted duck | *Aythya fuligula* | | ✓ |
| | Tundra swan | *Cygnus columbiansis* | | ✓ |
| Cormorants | Double-crested cormorant | *Phalacrocorax auritus* | ✓ | |
| Cranes | Sandhill crane | *Grus canadensis* | | ✓ |
| Grebes | Clark's grebe | *Aechmophorus clarkii* | | ✓ |
| | Pied-billed grebe | *Podilymbus podiceps* | ✓ | ✓ |
| | Red-necked grebe | *Podiceps grisegena* | ✓ | ✓ |
| Gulls | Franklin's gull | *Leucophaeus pipixcan* | ✓ | ✓ |
| | Ring-billed gull | *Larus delawarensis* | | ✓ |
| Herons | American bittern | *Botaurus lentiginous* | ✓ | |
| | Black-crowned night heron | *Nycticorax nycticorax* | ✓ | ✓ |
| | Great Blue Heron | *Ardea herodias* | ✓ | ✓ |
| Kingfishers | Belted kingfisher | *Megaceryle alcyon* | | ✓ |
| Loons | Common loon | *Gavia immer* | ✓ | ✓ |
| Partridges | Gray partridge | *Perdix perdix* | | ✓ |

*Continued on next page*

Supplementary Table A2 – *Continued from previous page*

| Group | Animal (common name) | Suspected species name | Camera | Recorder |
|---|---|---|---|---|
| Passerines | American crow | *Corvus brachyrhynchos* | ✓ | ✓ |
| | American goldfinch | *Spinus tristis* | | ✓ |
| | American pipit | *Anthus rubescens* | | ✓ |
| | American robin | *Turdus migratorius* | | ✓ |
| | American tree sparrow | *Spizelloides arborea* | | ✓ |
| | Bank swallow | *Riparia riparia* | | ✓ |
| | Barn swallow | *Hirundo rustica* | | ✓ |
| | Black-billed Magpie | *Pica hudsonia* | ✓ | ✓ |
| | Blackpoll warbler | *Setophaga striata* | | ✓ |
| | Blue jay | *Cyanocitta cristata* | | ✓ |
| | Bohemian waxwing | *Bombycilla garrulus* | | ✓ |
| | Brambling | *Fringilla montifringilla* | | ✓ |
| | Brewer's blackbird | *Euphagus cyanocephalus* | | ✓ |
| | Brewer's sparrow | *Spizella breweri* | | ✓ |
| | Brown-headed cowbird | *Molothrus ater* | | ✓ |
| | Cedar waxwing | *Bombycilla cedrorum* | | ✓ |
| | Clay-colored sparrow | *Spizella pallida* | | ✓ |
| | Cliff swallow | *Petrochelidon pyrrhonota* | | ✓ |
| | Common raven | *Corvus corvax* | | ✓ |
| | Eastern kingbird | *Tyrannus tyrannus* | | ✓ |
| | European starling | *Sturnus vulgaris* | | ✓ |
| | Gray-crowned Rosy-finch | *Leucosticte tephrocotis* | | ✓ |
| | Great crested flycatcher | *Myiarchus crinitus* | | ✓ |
| | Harris's sparrow | *Zonotrochia querula* | | ✓ |
| | Lapland longspur | *Calcarius lapponicus* | | ✓ |
| | LeConte's sparrow | *Ammodramus leconteii* | | ✓ |
| | Lincoln's sparrow | *Melospiza lincolnii* | | ✓ |
| | Loggerhead shrike | *Lanius ludovicianus* | | ✓ |
| | Nelson's sparrow | *Ammodramus nelsoni* | | ✓ |
| | Northern rough-winged swallow | *Stelgidopteryx serripennis* | | ✓ |
| | Northern waterthrush | *Parkesia noveboracensis* | | ✓ |
| | Pine grosbeak | *Pinicola enucleator* | | ✓ |
| | Pine siskin | *Spinus pinus* | | ✓ |
| | Purple martin | *Progne subis* | | ✓ |
| | Red-breasted nuthatch | *Sitta canadensis* | | ✓ |
| | Red-winged blackbird | *Agelaius phoeniceus* | ✓ | ✓ |
| | Rusty blackbird | *Euphagus carolinus* | | ✓ |
| | Savannah sparrow | *Passerculus sandwichensis* | | ✓ |
| | Snow bunting | *Plectrophenax nivalis* | | ✓ |
| | Song sparrow | *Melospiza melodia* | | ✓ |
| | Swamp sparrow | *Melospiza georgiana* | | ✓ |

*Continued on next page*

Supplementary Table A2 – *Continued from previous page*

| Group | Animal (common name) | Suspected species name | Camera | Recorder |
|---|---|---|---|---|
| Passerines (continued) | Tree swallow | *Tachycineta bicolor* | | ✓ |
| | Varied thrush | *Ixoreus naevius* | | ✓ |
| | Vesper sparrow | *Pooecetes gramineus* | | ✓ |
| | Western kingbird | *Tyrannus verticalis* | | ✓ |
| | White-breasted nuthatch | *Sitta carolinensis* | | ✓ |
| | White-winged crossbill | *Loxia leucoptera* | | ✓ |
| | Winter wren | *Troglodytes hiemalis* | | ✓ |
| | Yellow-headed blackbird | *Xanthocephalus xanthocephalus* | ✓ | ✓ |
| | Yellow-rumped warbler | *Setophaga coronata* | | ✓ |
| | Yellow warbler | *Setophaga petechia* | | ✓ |
| Pelicans | American white pelican | *Pelecanus erythrorhynchos* | ✓ | |
| Rails | American coot | *Fulica americana* | ✓ | ✓ |
| | Sora | *Porzana carolina* | | ✓ |
| | Virginia rail | *Rallus limicola* | | ✓ |
| | Yellow rail | *Coturnicops noveboracensis* | | ✓ |
| Raptors | American kestrel | *Falco sparverius* | | ✓ |
| | Bald eagle | *Haliaeetus leucocephalus* | | ✓ |
| | Boreal owl | *Aegolius funereus* | | ✓ |
| | Broad-winged hawk | *Buteo platypterus* | | ✓ |
| | Burrowing owl | *Athene cunicularia* | | ✓ |
| | Cooper's hawk | *Accipiter cooperii* | ✓ | |
| | Great horned owl | *Bubo virginianus* | | ✓ |
| | Long-eared owl | *Asio otus* | | ✓ |
| | Merlin | *Falco columbarius* | | ✓ |
| | Northern pygmy owl | *Glaucidium californicum* | | ✓ |
| | Northern saw-whet owl | *Aegolius acadicus* | | ✓ |
| | Osprey | *Pandion haliaetus* | | ✓ |
| | Peregrine falcon | *Falco peregrinus* | | ✓ |
| | Red-tailed hawk | *Buteo jamaicensis* | | ✓ |
| | Short-eared owl | *Asio flammeus* | | ✓ |
| | Swainson's hawk | *Buteo swainsoni* | | ✓ |
| Shorebirds | American avocet | *Recurvirostra americana* | ✓ | ✓ |
| | Dunlin | *Calidris alpina* | | ✓ |
| | Greater yellowlegs | *Tringa melanoleuca* | | ✓ |
| | Killdeer | *Charadrius vociferus* | | ✓ |
| | Least sandpiper | *Calidris minutilla* | | ✓ |
| | Lesser yellowlegs | *Tringa flavipes* | ✓ | ✓ |
| | Long-billed curlew | *Numenius americans* | | ✓ |
| | Long-billed dowitcher | *Limnodromus scolopaceus* | | ✓ |
| | Marbled godwit | *Limosa fedoa* | | ✓ |
| | Pectoral sandpiper | *Calidris melanotos* | | ✓ |

*Continued on next page*

Supplementary Table A2 – *Continued from previous page*

| Group | Animal (common name) | Suspected species name | Camera | Recorder |
|---|---|---|---|---|
| Shorebirds (continued) | Semipalmated plover | *Charadrius semipalmatus* | | ✓ |
| | Semipalmated sandpiper | *Calidris pusilla* | | ✓ |
| | Short-billed dowitcher | *Limnodromus griseus* | | ✓ |
| | Solitary sandpiper | *Tringa solitaria* | | ✓ |
| | Spotted sandpiper | *Actitius macularius* | ✓ | ✓ |
| | Upland sandpiper | *Bartramia longicauda* | | ✓ |
| | Willet | *Tringa semipalmata* | | ✓ |
| | Wilson's phalarope | *Phalaropus tricolor* | ✓ | ✓ |
| | Wilson's snipe | *Gallinago delicata* | | ✓ |
| Terns | Black tern | *Chlidonias niger* | ✓ | ✓ |
| | Caspian tern | *Hydroprogne caspia* | | ✓ |
| | Common tern | *Sterna hirundo* | ✓ | ✓ |
| | Forster's tern | *Sterna forsteri* | ✓ | ✓ |
| | | **Mammals** | | |
| Caniforms | Coyote | *Canis latrans* | | ✓ |
| | Gray wolf | *Canis lupus* | | ✓ |
| Rodents | Muskrat | *Ondatra zibethicus* | ✓ | |
| Ruminants | White-tailed deer | *Odocoileus virginianus* | ✓ | |
| | | **Reptiles** | | |
| Snakes | Garter snake | *Thamnophis sirtalis* | ✓ | |

Supplementary Table A3. Invertebrates detected at the wetland sites and Heritage Lake from benthic kick netting in 2022.

| Group | Animal (common name) | Species name |
|---|---|---|
| Amphipods | Freshwater shrimp | *Gammarus lacustris* |
| | Mexican freshwater shrimp | *Hyalella azteca* |
| Annelids | Sludge worm | *Tubifex* sp. |
| Aquatic mites | Mite | *Arrenurus* sp. |
| | Mite | *Eylais* sp. |
| | Mite | *Limnesia undulatoides* |
| | Mite | *Limnesia* sp. |
| | Mite | *Limnesiinae* sp. |
| Backswimmers | Backswimmer | *Notonecta undulata* |
| Beetles | Antenna agabus beetle | *Agabus cf. antennatus* |
| | Fall's crawling water beetle | *Haliplus falli* |
| | Clearneck crawling water beetle | *Haliplus immaculicollis* |
| | Diving beetle | *Hygrotus* sp. |
| | Lake Superior predaceous diving beetle | *Neoporus superioris* |
| | Satiny swimming beetle | *Rhantus sericans* |
| Caddisflies | External northern caddisfly | *Limnephalus externus* |
| | Caddisfly | *Mystacides* sp. |
| | Grey giant caddisfly | *Phryganea cinerea* |
| Chironomids | Midge | *Chaoboridae* sp. |
| | Midge | *Chironomus atrella* |
| | Midge | *Chironomidae* sp. |
| | Midge | *Cryptochironomus* sp. |
| Damselflies | Hagen's bluet | *Enallagma hangeni* |
| | American bluet | *Enallagma* sp. |
| | Damselfly | *Ischnura* sp. |
| Dragonflies | Variable darner | *Aeshna interrupta* |
| | Four-spotted skimmer | *Libellula quadrimaculata* |
| Leafhoppers | Silver leafhopper | *Athysanus argentarius* |
| Leeches | Leech | *Erpobdella obscura* |
| | Leech | *Erpobdella punctata* |
| | Leech | *Erpobdella* sp. |
| | Scutate snail leech | *Helobdella modesta* |
| | Leech | *Helobdella* sp. |
| | Leech | *Glossiphonia elegans* |
| | Leech | *Glossiphoniidae sp.* |
| | Leech | *Placobdella akahkway* |
| Mayfly | Common small square-gilled mayfly | *Caenis latipennis* |
| | Young's small square-gilled mayfly | *Caenis youngi* |
| | Red small minnow mayfly | *Callibaetis ferrugineus* |
| | Mayfly | *Callibaetis* sp. |

*Continued on next page*

Supplementary Table A3 – *Continued from previous page*

| Group | Animal (common name) | Species name |
|---|---|---|
| Water boatmen | Water boatman | *Callicorixa audeni* |
| | Water boatman | *Cymatia americana* |
| | Water boatman | *Sigara bicoloripennis* |
| | Water boatman | *Sigara decoratella* |
| | Water boatman | *Trichocorixa borealis* |
| | Water boatman | *Trichocorixa sexcincta* |

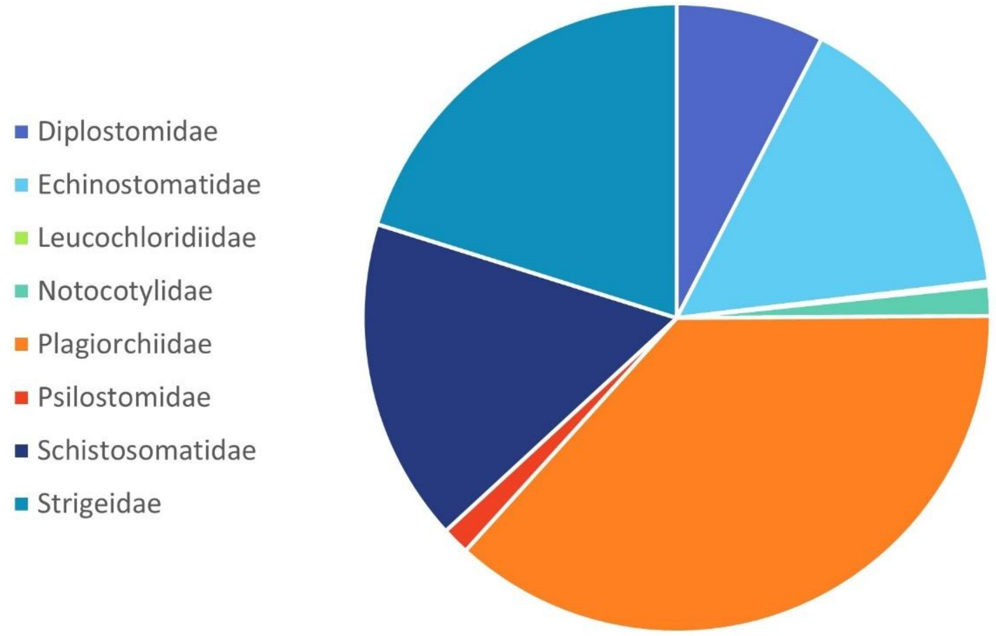


Supplementary Figure A1. Proportion of each trematode family based on DNA sequence identifications combined across all sites and years.

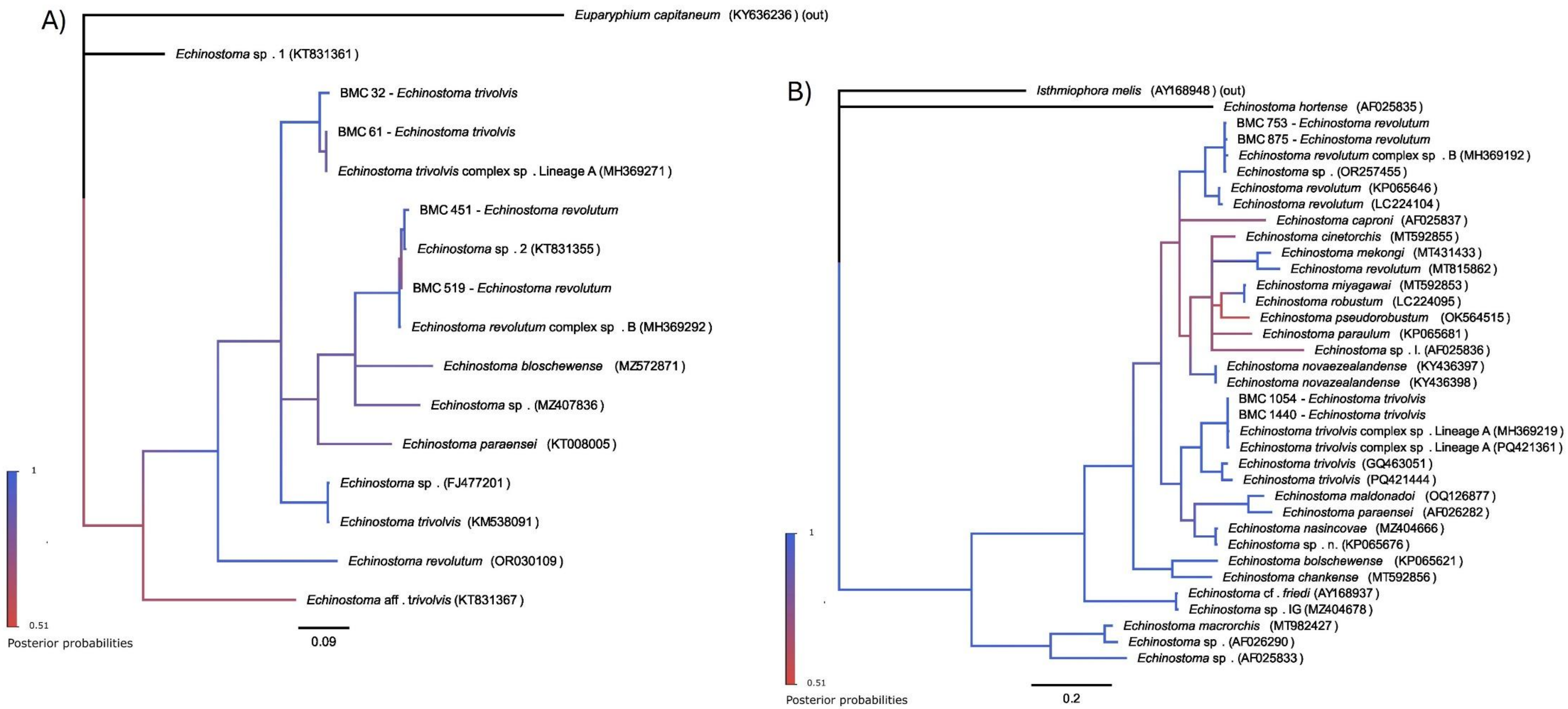


Supplementary Figure A2. Bayesian inference molecular phylogenies of the Echinostomatidae (genus: *Echinostoma*) based on A) *COI*, with substitution model HKY + G + I, and B) *nad1*, using substitution model HKY + G. Accession numbers follow species names. Samples with a "BMC" label indicate sequences obtained during this study.

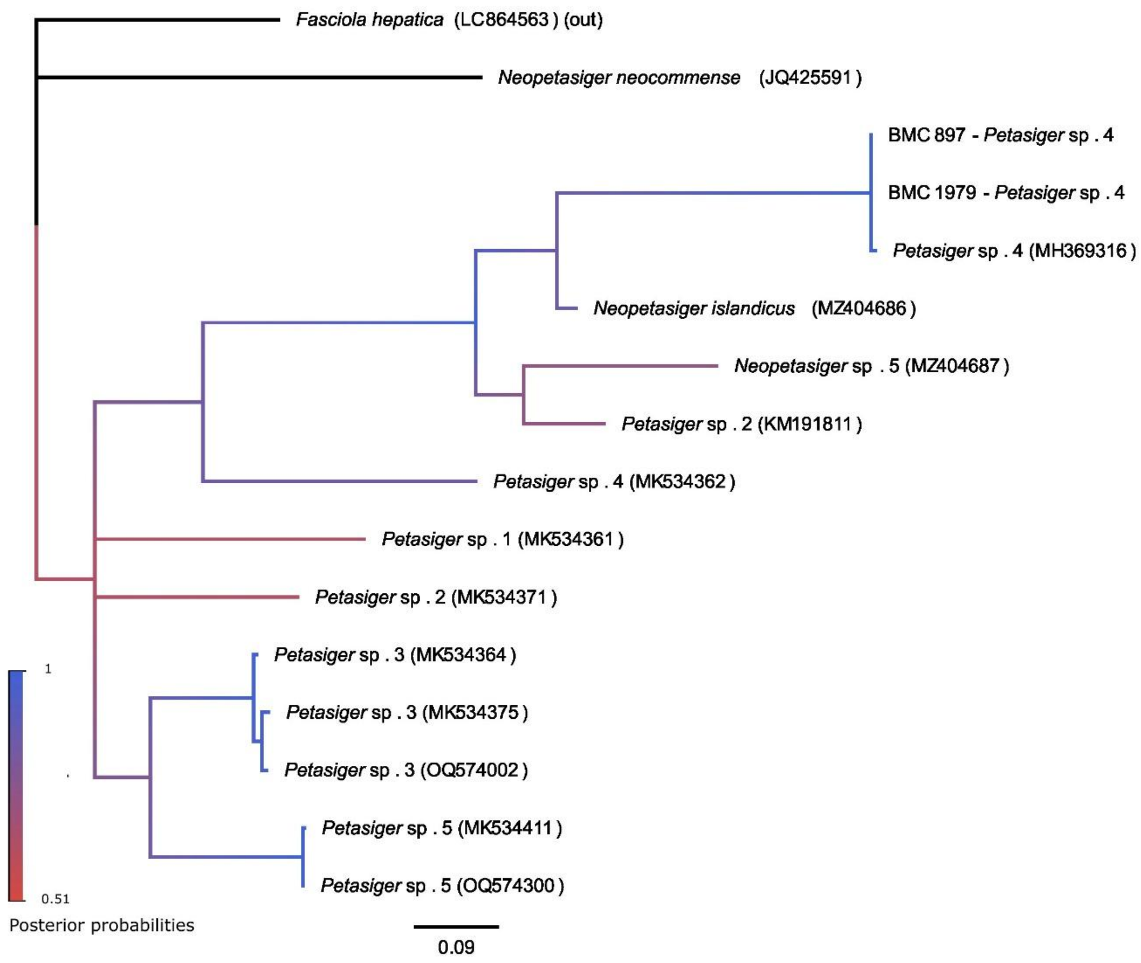


Supplementary Figure A3. Bayesian inference molecular phylogenies of the Echinostomatidae (genus: *Petasiger*) based on *nad1*, using substitution model HKY + G. Accession numbers follow species names. Samples with a "BMC" label indicate sequences obtained during this study.

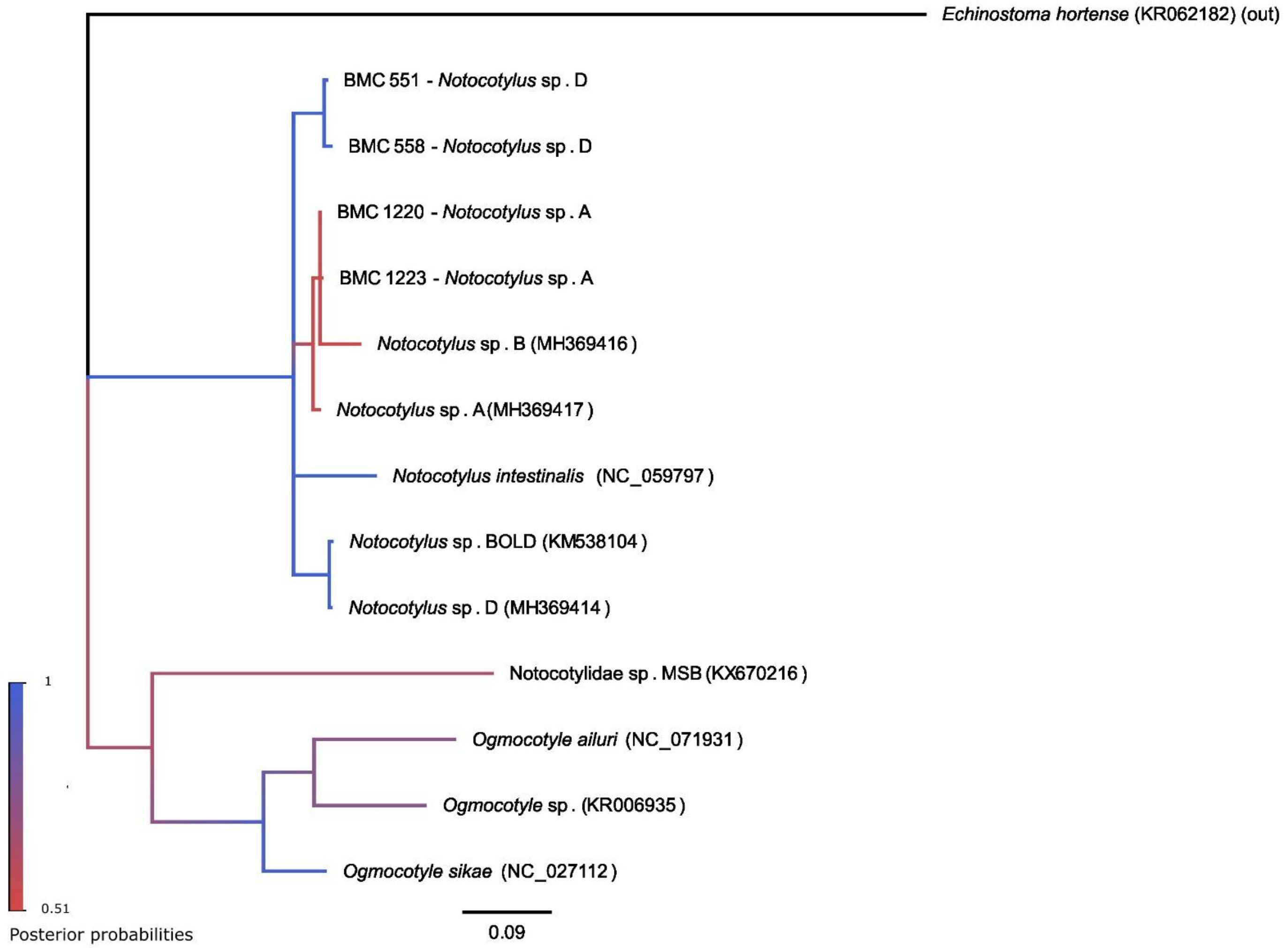


Supplementary Figure A4. Bayesian inference molecular phylogeny of the Notocotylidae based on *COI*, using substitution model HKY + G. Accession numbers follow species names. Samples with a "BMC" label indicate sequences obtained during this study.

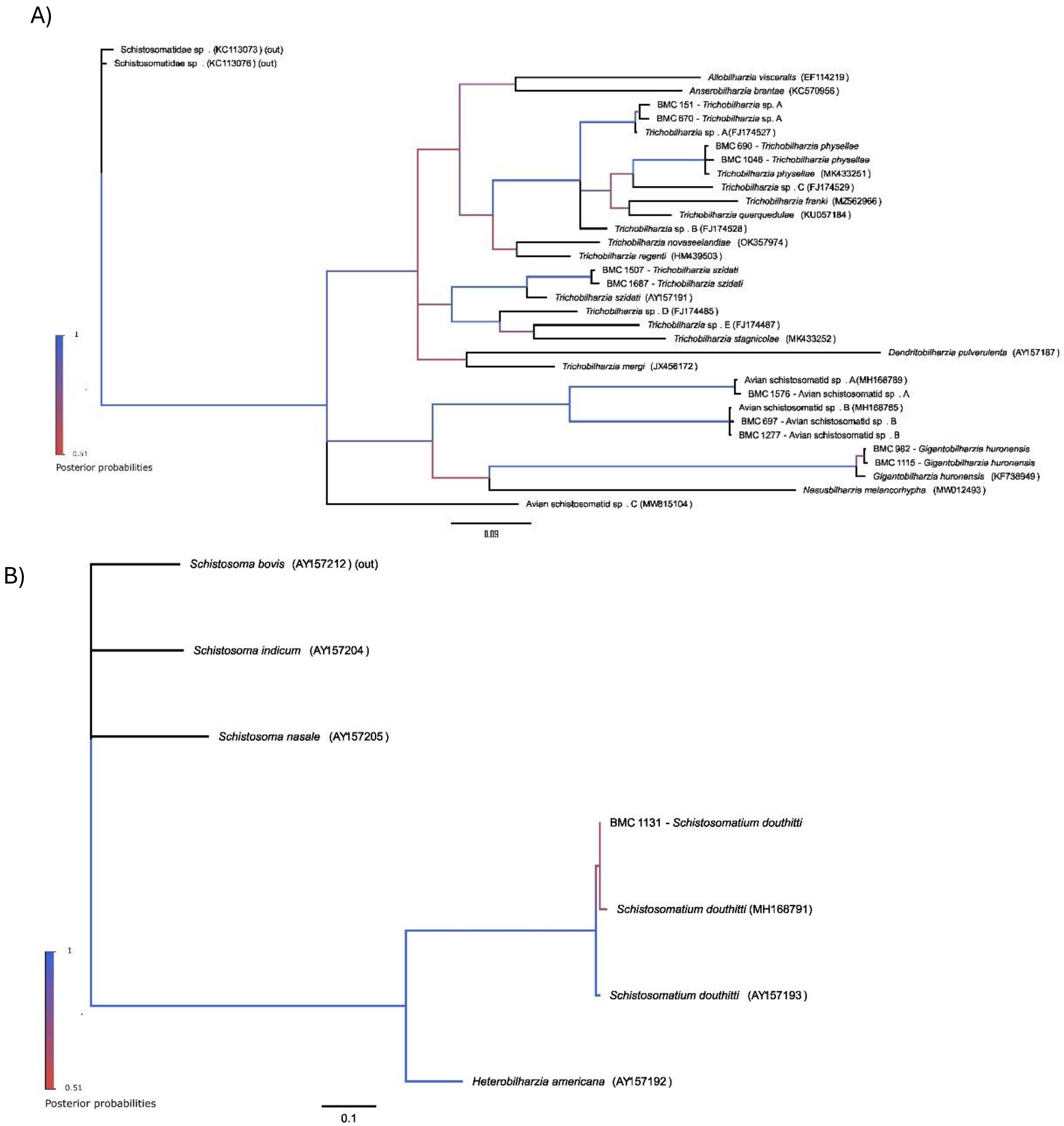


Supplementary Figure A5. Bayesian inference molecular phylogenies of the Schistosomatidae based on *COI*. A) Avian schistosomes using substitution model GTR + G. B) Mammalian schistosomes using substitution model HKY + G. Accession numbers follow species names. Samples with a “BMC” label indicate sequences obtained during this study.

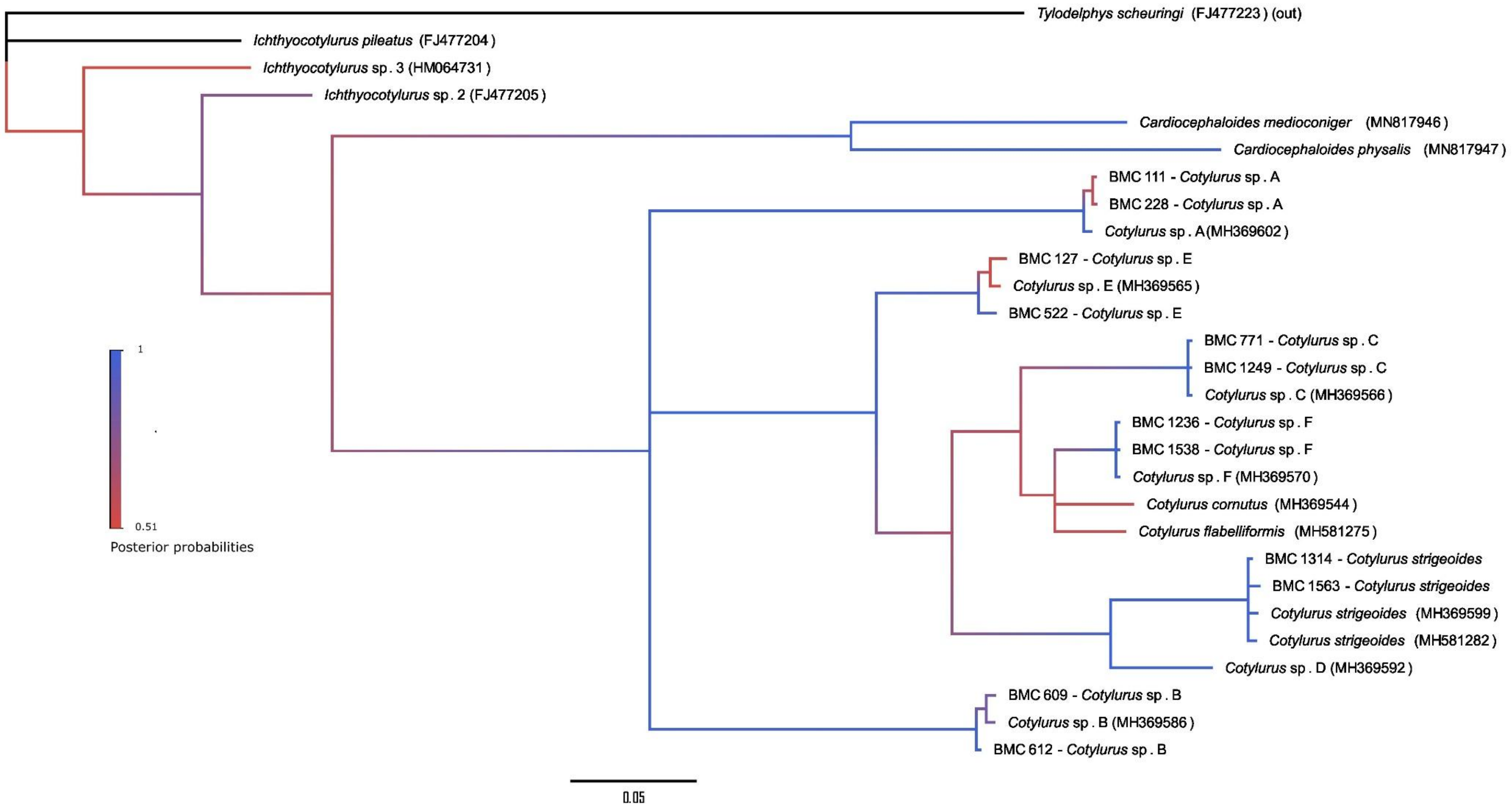


Supplementary Figure A6. Bayesian inference molecular phylogeny of the Strigeidae I based on *COI* using the HKY + G + I substitution model. Accession numbers follow species names. Samples with a “BMC” label indicate sequences obtained during this study.

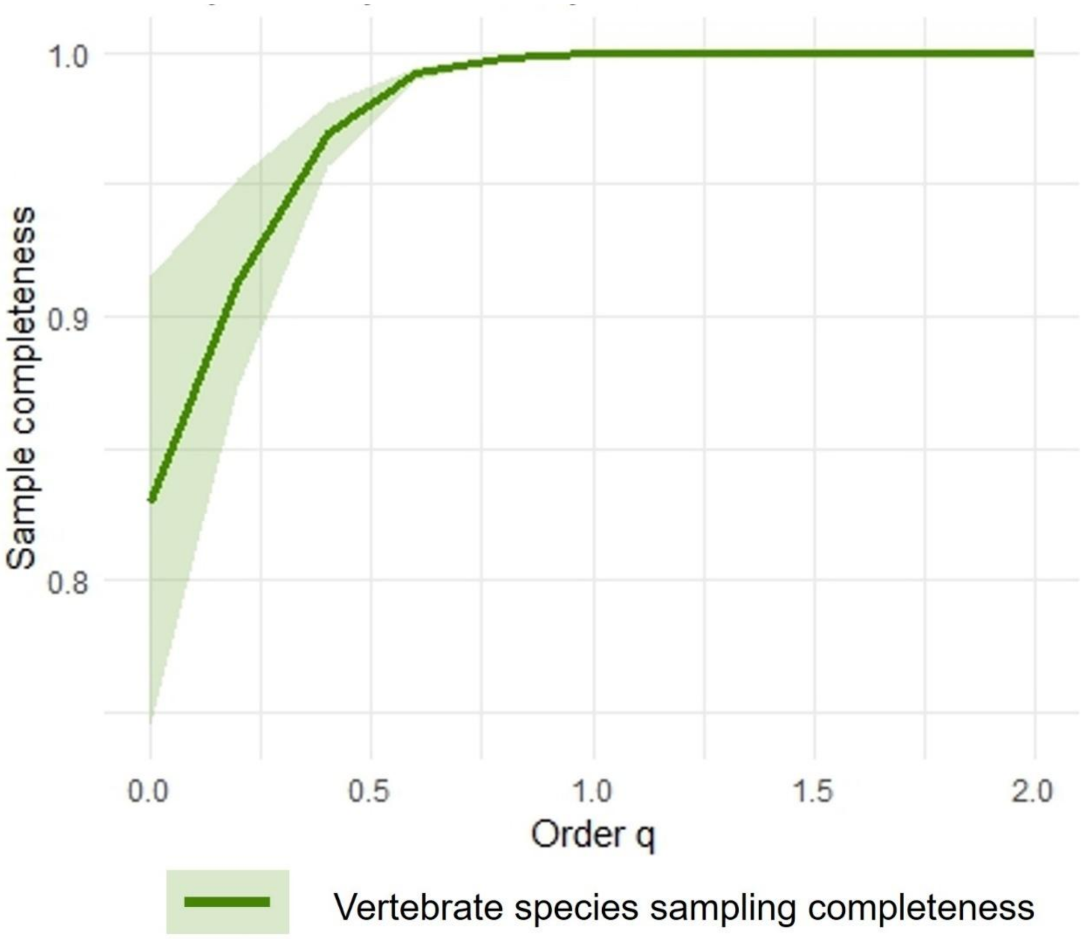


Supplementary Figure A7. Sampling completeness plot for vertebrates recorded from field cameras and bird song recorders at the eight wetland study sites combined for years 2020, 2021, and 2022.

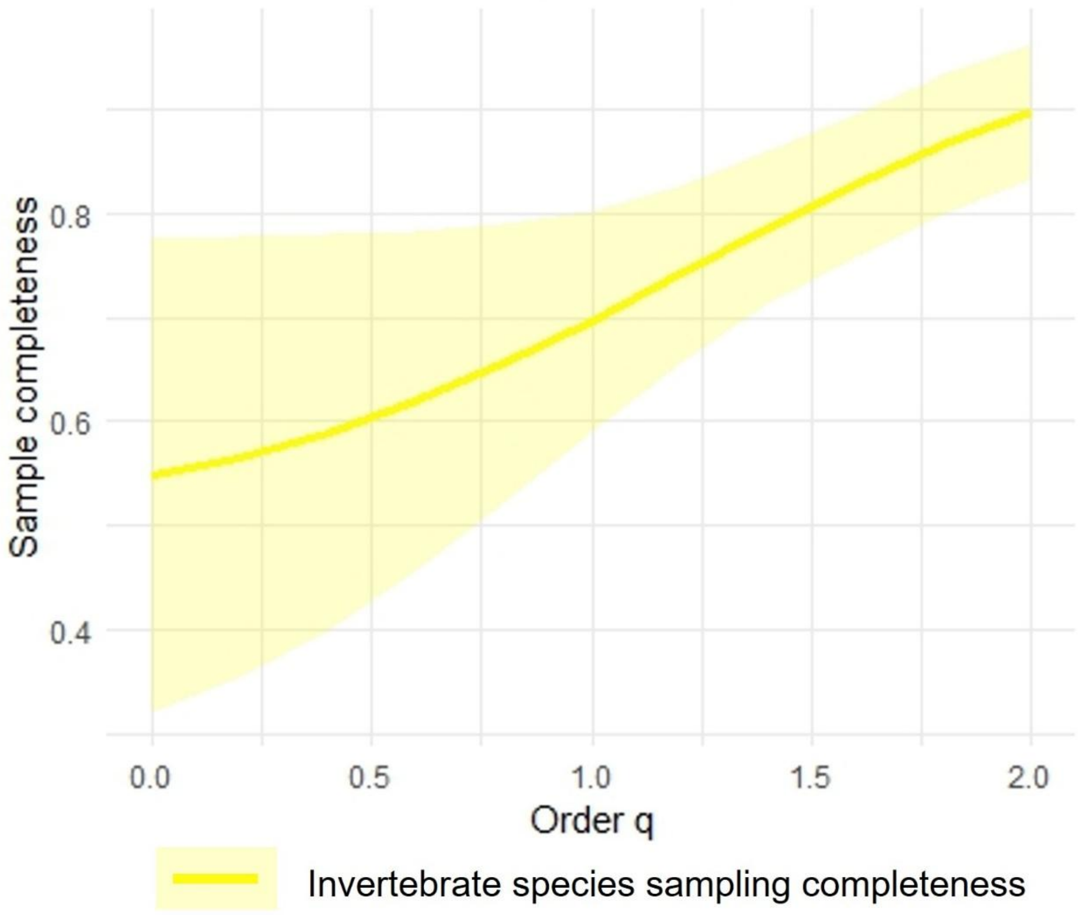


Supplementary Figure A8. Sampling completeness plot for invertebrates collected using benthic kick netting at the eight wetland sites and Heritage Lake in 2022.

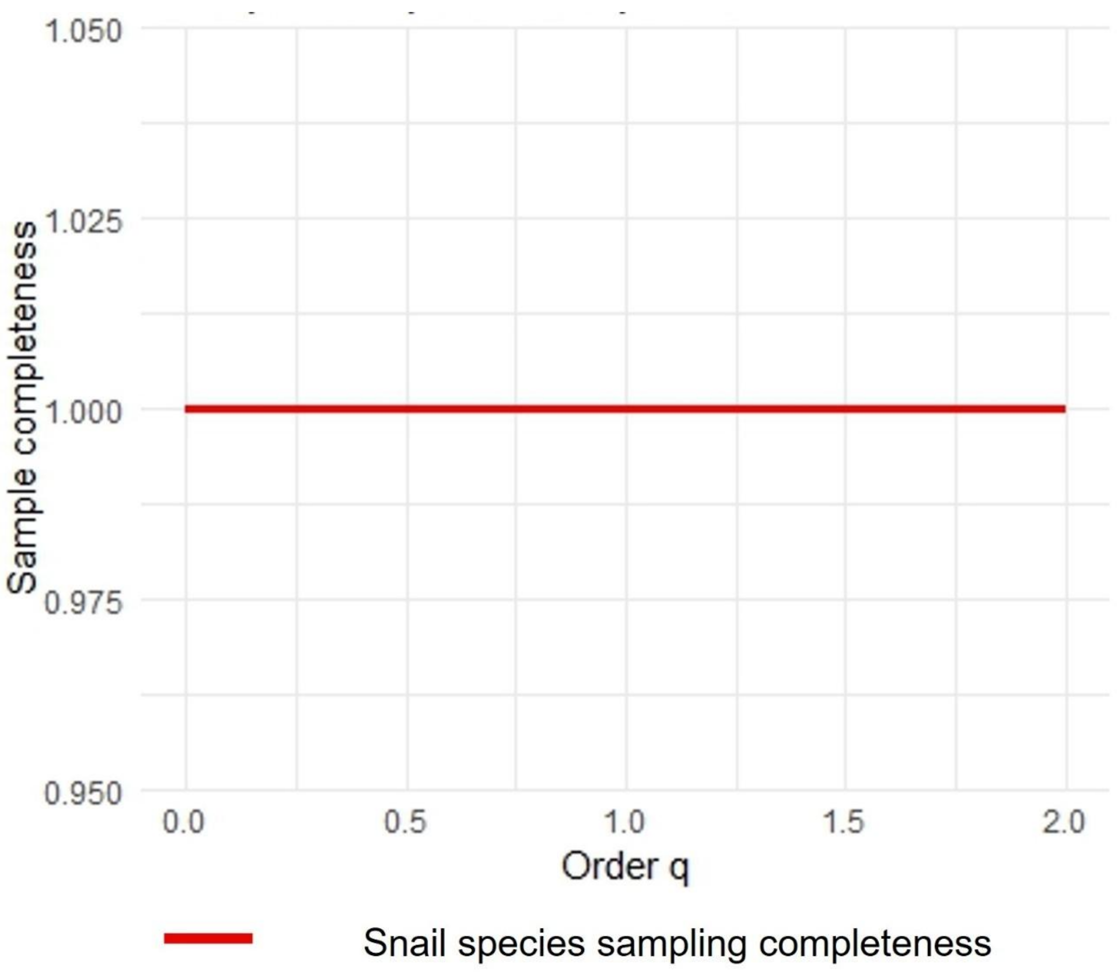


Supplementary Figure A9. Sampling completeness plot for snails collected at the eight wetland sites and Heritage Lake for 2019-2022.

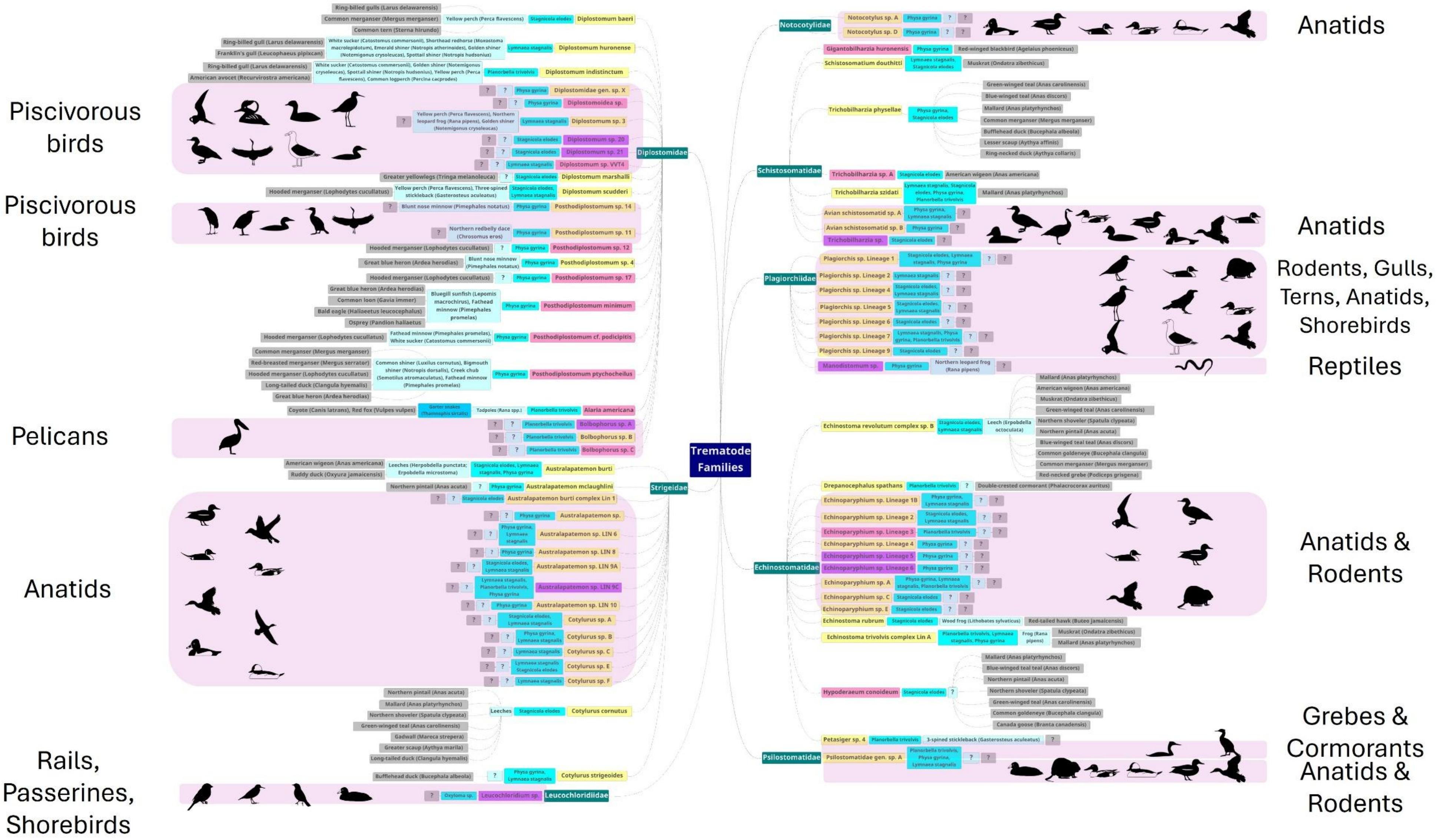
Piscivorous birds
Piscivorous birds
Pelicans
Anatids
Rails, Passerines, Shorebirds
Trematode Families
Diplostomidae
Strigeidae
Leucochloridiidae
Notocotylidae
Schistosomatidae
Plagiorchiidae
Echinostomatidae
Psilostomatidae
Anatids
Anatids
Rodents, Gulls, Terns, Anatids, Shorebirds
Reptiles
Anatids & Rodents
Grebes & Cormorants
Anatids & Rodents

Supplementary Figure A10. Host-parasite interactions observed during this study with inferred potential definitive hosts where needed. Trematode families are in green. First and second-intermediate hosts are in shades of blue, along with paratenic hosts. Trematode species in pink are those not previously reported in Alberta. Trematode species found during this study and in the previous study in Alberta are in yellow. Trematodes in purple indicate putative novel species identified during this study. Groups with inferred definitive host relationships are within transparent pink boxes and include inferred potential hosts based on species observed during the traditional biodiversity assessment. Definitive host images from PhyloPic (Palomo-Munoz, 2021; Price, 2023; Keesey, 2025).
