## Appendix B for "Snail-trematode dynamics in central Alberta wetlands: A longitudinal survey of infections and interactions"

Table B1. Averaged percent similarities of the species belonging to the Diplostomidae I. Standard error estimate(s) are shown above the diagonal and were obtained by a bootstrap procedure (500 replicates). Ambiguous positions were removed for each sequence pair. Evolutionary analyses were conducted in MEGA (Tamura et al., 2021).

| | 1 | 2 | 3 | 4 | 5 | 6 | 7 | 8 | 9 | 10 | 11 | 12 | 13 |
|---|---|---|---|---|---|---|---|---|---|---|---|---|---|
| 1. *Alaria americana* | - | 1.498 | 1.4456 | 1.6922 | 0.51 | 1.7735 | 2.156 | 2.0798 | 1.8782 | 1.8544 | 1.805 | 1.7281 | 1.7788 |
| 2. *Alaria* sp. 1 | 89.323 | - | 1.5015 | 1.735 | 1.4311 | 1.6551 | 2.1147 | 2.0459 | 1.7534 | 1.9001 | 1.8319 | 1.6831 | 1.7549 |
| 3. *Alaria* sp. 2 | 90.885 | 90.625 | - | 1.7542 | 1.4859 | 1.8821 | 2.1852 | 2.0689 | 1.8067 | 1.9103 | 1.8711 | 1.8327 | 1.8818 |
| 4. *Austrodiplostomum ostrowskiae* | 86.198 | 85.417 | 85.417 | - | 1.7509 | 1.7249 | 2.0087 | 2.0476 | 1.7087 | 1.7518 | 1.745 | 1.7424 | 1.7049 |
| 5. BMC - *Alaria americana* (2) | 99.089 | 89.453 | 90.495 | 85.938 | - | 1.7262 | 2.1239 | 1.9926 | 1.7933 | 1.898 | 1.8359 | 1.6617 | 1.7142 |
| 6. BMC - *Diplostomum indistinctum* | 85.417 | 87.240 | 85.156 | 84.115 | 85.547 | - | 1.9839 | 1.9081 | 1.6608 | 1.8809 | 1.7128 | 1.4781 | 1.6686 |
| 7. BMC - Diplostomidae gen. sp. X | 77.083 | 78.646 | 77.604 | 82.292 | 77.214 | 78.385 | - | 2.0235 | 2.0503 | 2.0322 | 2.046 | 1.9388 | 2.0937 |
| 8. BMC - *Diplostomum* sp. 20 | 81.510 | 82.031 | 80.729 | 82.292 | 82.422 | 83.333 | 79.427 | - | 1.8827 | 1.9087 | 1.8785 | 1.9092 | 1.8937 |
| 9. BMC - *Diplostomum scudderi* (2) | 85.156 | 86.068 | 86.068 | 85.417 | 85.547 | 88.411 | 79.557 | 84.245 | - | 1.7821 | 1.537 | 1.5742 | 1.4235 |
| 10. BMC - *Diplostomum* sp. 21 (2) | 81.901 | 83.464 | 83.984 | 84.245 | 81.641 | 82.682 | 79.036 | 83.464 | 85.156 | - | 1.6767 | 1.8468 | 1.7958 |
| 11. BMC - *Diplostomum marshalli* | 84.635 | 85.156 | 83.854 | 85.938 | 84.505 | 87.760 | 80.208 | 84.896 | 89.323 | 87.370 | - | 1.6626 | 1.6713 |
| 12. BMC - *Diplostomum* sp. 3 | 86.458 | 86.198 | 84.635 | 86.719 | 86.589 | 90.104 | 79.948 | 84.896 | 89.323 | 82.682 | 87.500 | - | 1.7107 |
| 13. BMC - *Diplostomum* sp. VVT4 | 85.677 | 86.719 | 84.896 | 84.635 | 85.807 | 87.760 | 77.865 | 84.896 | 90.495 | 83.984 | 87.500 | 86.458 | - |
| 14. BMC - *Diplostomum huronense* | 86.126 | 85.864 | 84.031 | 84.817 | 85.471 | 87.173 | 79.058 | 84.031 | 89.005 | 84.686 | 89.529 | 89.791 | 89.005 |
| 15. Diplostomidae gen. sp. O | 79.427 | 79.688 | 79.688 | 80.990 | 79.036 | 78.646 | 87.500 | 77.865 | 78.776 | 78.776 | 79.948 | 79.167 | 78.125 |
| 16. Diplostomidae gen. sp. X | 77.083 | 78.906 | 77.865 | 82.292 | 77.214 | 79.167 | 99.219 | 80.208 | 79.818 | 79.297 | 80.208 | 80.729 | 78.125 |
| 17. Diplostomoidea sp. | 81.462 | 82.768 | 83.551 | 81.723 | 80.809 | 81.462 | 79.373 | 77.807 | 82.115 | 79.765 | 83.029 | 81.984 | 81.201 |
| 18. D*iplostomum ardeae* | 88.021 | 87.760 | 86.719 | 87.500 | 87.891 | 87.240 | 80.208 | 83.073 | 88.932 | 86.328 | 88.542 | 87.500 | 88.021 |
| 19. *Diplostomum baeri* | 87.240 | 84.896 | 86.198 | 85.156 | 87.370 | 87.760 | 80.729 | 85.156 | 88.672 | 83.464 | 88.021 | 88.021 | 89.063 |
| 20. *Diplostomum gavium* | 86.198 | 86.198 | 84.635 | 86.719 | 86.328 | 89.844 | 80.208 | 84.896 | 89.063 | 82.682 | 87.240 | 99.219 | 86.198 |
| 21. *Diplostomum huronense* | 86.198 | 86.979 | 85.156 | 85.156 | 85.547 | 88.802 | 78.906 | 83.594 | 89.583 | 84.766 | 88.802 | 89.844 | 88.542 |
| 22. *Diplostomum indistinctum* | 85.417 | 87.240 | 85.156 | 83.854 | 85.547 | 98.958 | 78.385 | 84.115 | 88.151 | 83.203 | 87.500 | 90.104 | 87.760 |
| 23. *Diplostomum marshalli* | 84.635 | 85.938 | 84.375 | 86.458 | 84.505 | 88.281 | 80.208 | 85.417 | 89.063 | 87.370 | 98.958 | 87.500 | 87.500 |
| 24. *Diplostomum mergi* | 85.938 | 87.760 | 85.938 | 85.677 | 85.547 | 89.063 | 79.948 | 83.073 | 88.932 | 86.589 | 89.063 | 88.542 | 88.021 |
| 25. *Diplostomum pseudospathaceum* | 87.206 | 87.467 | 85.379 | 86.684 | 87.337 | 90.601 | 80.418 | 85.379 | 90.339 | 84.204 | 87.990 | 97.128 | 87.728 |

*Continued on next page*

Table B1 – *Continued from previous page*

| | 14 | 15 | 16 | 17 | 18 | 19 | 20 | 21 | 22 | 23 | 24 | 25 | 26 | 27 | 28 |
|---|---|---|---|---|---|---|---|---|---|---|---|---|---|---|---|
| 1. *Alaria americana* | 1.7395 | 2.0425 | 2.1555 | 1.8878 | 1.6121 | 1.723 | 1.753 | 1.757 | 1.8099 | 1.8132 | 1.8275 | 1.6784 | 1.8703 | 1.7205 | 1.6536 |
| 2. *Alaria* sp. 1 | 1.7169 | 2.0695 | 2.113 | 1.9612 | 1.6286 | 1.8741 | 1.6819 | 1.6776 | 1.7058 | 1.775 | 1.7523 | 1.6183 | 1.7631 | 1.7278 | 1.7737 |
| 3. *Alaria* sp. 2 | 1.908 | 2.0675 | 2.178 | 1.8566 | 1.711 | 1.8071 | 1.8248 | 1.8758 | 1.9113 | 1.8555 | 1.8548 | 1.7873 | 1.8193 | 1.8844 | 1.7857 |
| 4. *Austrodiplostomum ostrowskiae* | 1.7575 | 1.9692 | 2.0245 | 1.8415 | 1.6163 | 1.7616 | 1.7279 | 1.7361 | 1.7822 | 1.737 | 1.7332 | 1.667 | 1.7187 | 1.7456 | 1.5915 |
| 5. BMC - *Alaria americana* (2) | 1.833 | 2.0411 | 2.1261 | 1.9531 | 1.6482 | 1.7474 | 1.6883 | 1.8432 | 1.7662 | 1.8417 | 1.8936 | 1.6194 | 1.7866 | 1.8135 | 1.7039 |
| 6. BMC - *Diplostomum indistinctum* | 1.7548 | 2.0358 | 1.9561 | 2.0078 | 1.6454 | 1.7388 | 1.4981 | 1.6416 | 0.5138 | 1.6752 | 1.5733 | 1.446 | 1.6492 | 1.7431 | 1.6899 |
| 7. BMC - Diplostomidae gen. sp. X | 2.0094 | 1.7422 | 0.4278 | 2.0678 | 1.9576 | 2.025 | 1.9201 | 2.0044 | 1.9857 | 2.0174 | 1.9937 | 1.8648 | 2.0804 | 2.0017 | 2.0859 |
| 8. BMC - *Diplostomum* sp. 20 | 1.8695 | 2.1542 | 2.0025 | 2.0706 | 1.906 | 1.8715 | 1.9034 | 1.8709 | 1.8588 | 1.8726 | 1.9298 | 1.8345 | 1.884 | 1.8646 | 1.8722 |
| 9. BMC - *Diplostomum scudderi* (2) | 1.5477 | 2.0929 | 2.0403 | 1.9085 | 1.5842 | 1.627 | 1.6027 | 1.526 | 1.6994 | 1.5608 | 1.6232 | 1.4915 | 0.2811 | 1.5384 | 1.6531 |
| 10. BMC - *Diplostomum* sp. 21 (2) | 1.7931 | 2.076 | 2.0606 | 1.9153 | 1.6726 | 1.8208 | 1.8237 | 1.7374 | 1.8853 | 1.6319 | 1.7022 | 1.8177 | 1.7963 | 1.777 | 1.7522 |
| 11. BMC - *Diplostomum marshalli* | 1.6019 | 2.0532 | 2.0575 | 1.8857 | 1.5829 | 1.6042 | 1.6607 | 1.584 | 1.7328 | 0.5141 | 1.5853 | 1.652 | 1.5516 | 1.5871 | 1.5917 |
| 12. BMC - *Diplostomum* sp. 3 | 1.5552 | 1.963 | 1.9164 | 1.9083 | 1.5521 | 1.7037 | 0.452 | 1.5441 | 1.498 | 1.6852 | 1.6435 | 0.8496 | 1.5661 | 1.5348 | 1.6354 |
| 13. BMC - *Diplostomum* sp. VVT4 | 1.5122 | 2.0565 | 2.0976 | 1.9472 | 1.6094 | 1.5476 | 1.7259 | 1.5204 | 1.6814 | 1.6724 | 1.6168 | 1.6008 | 1.4272 | 1.488 | 1.4417 |
| 14. BMC - *Diplostomum huronense* | - | 2.0565 | 2.019 | 1.9131 | 1.6661 | 1.6551 | 1.5759 | 0.8019 | 1.7763 | 1.6118 | 1.6006 | 1.4915 | 1.5494 | 0.3068 | 1.5294 |
| 15. Diplostomidae gen. sp. O | 77.487 | - | 1.7268 | 2.0457 | 1.925 | 2.054 | 1.9518 | 2.0922 | 2.0258 | 2.0597 | 1.922 | 1.8972 | 2.0686 | 2.0564 | 2.0243 |
| 16. Diplostomidae gen. sp. X | 79.058 | 87.760 | - | 2.0708 | 1.9933 | 1.9891 | 1.8964 | 2.0167 | 1.9578 | 2.0265 | 2.0121 | 1.8692 | 2.0728 | 2.0099 | 2.0905 |
| 17. Diplostomoidea sp. | 81.890 | 80.679 | 79.112 | - | 1.8156 | 1.9201 | 1.8968 | 1.9045 | 2.0186 | 1.8718 | 1.8813 | 1.8942 | 1.9112 | 1.9201 | 1.799 |
| 18. D*iplostomum ardeae* | 87.435 | 81.510 | 80.469 | 83.290 | - | 1.5195 | 1.5594 | 1.6116 | 1.6549 | 1.5779 | 1.4704 | 1.5252 | 1.5516 | 1.6737 | 1.563 |
| 19. *Diplostomum baeri* | 87.696 | 80.469 | 81.510 | 82.768 | 88.542 | - | 1.734 | 1.672 | 1.7379 | 1.5899 | 1.5345 | 1.6793 | 1.6161 | 1.6583 | 1.4689 |
| 20. *Diplostomum gavium* | 89.529 | 79.427 | 80.990 | 81.723 | 87.240 | 87.760 | - | 1.5181 | 1.5125 | 1.6839 | 1.646 | 0.8938 | 1.5966 | 1.5534 | 1.6202 |
| 21. *Diplostomum huronense* | 97.382 | 76.823 | 78.906 | 82.507 | 86.979 | 87.240 | 90.104 | - | 1.6804 | 1.5572 | 1.4852 | 1.4934 | 1.5254 | 0.8061 | 1.5834 |
| 22. *Diplostomum indistinctum* | 86.911 | 78.646 | 79.167 | 81.201 | 87.240 | 87.760 | 89.844 | 88.542 | - | 1.6924 | 1.5941 | 1.4727 | 1.6841 | 1.7649 | 1.723 |
| 23. *Diplostomum marshalli* | 89.267 | 79.948 | 80.208 | 83.290 | 88.281 | 88.281 | 87.240 | 89.063 | 88.021 | - | 1.5598 | 1.6223 | 1.5705 | 1.5948 | 1.5896 |
| 24. *Diplostomum mergi* | 89.267 | 79.427 | 79.427 | 83.290 | 89.323 | 89.583 | 88.281 | 90.104 | 88.802 | 89.583 | - | 1.5476 | 1.613 | 1.589 | 1.5194 |
| 25. *Diplostomum pseudospathaceum* | 91.076 | 80.679 | 80.679 | 82.199 | 88.512 | 88.251 | 96.867 | 90.601 | 90.601 | 88.512 | 89.295 | - | 1.4898 | 1.455 | 1.6094 |

*Continued on next page*

Table B1 – *Continued from previous page*

| | 29 | 30 | 31 | 32 | 33 | 34 | 35 | 36 | 37 | 38 | 39 | 40 | 41 | 42 |
|---|---|---|---|---|---|---|---|---|---|---|---|---|---|---|
| 1. *Alaria americana* | 1.7506 | 1.8329 | 1.8945 | 1.7281 | 1.7645 | 1.7775 | 1.8423 | 1.8782 | 1.8977 | 1.756 | 1.7002 | 1.7827 | 1.7089 | 1.743 |
| 2. *Alaria* sp. 1 | 1.6686 | 1.7321 | 1.7547 | 1.6616 | 1.7981 | 1.7651 | 1.7761 | 1.7854 | 1.7665 | 1.587 | 1.7365 | 1.7312 | 1.7207 | 1.7966 |
| 3. *Alaria* sp. 2 | 1.8227 | 1.9387 | 1.8966 | 1.7954 | 1.838 | 1.8008 | 1.8668 | 1.8459 | 1.9312 | 1.6986 | 1.7706 | 1.8327 | 1.8178 | 1.8247 |
| 4. *Austrodiplostomum ostrowskiae* | 1.7238 | 1.7902 | 1.6865 | 1.6218 | 1.8061 | 1.6772 | 1.7369 | 1.7568 | 1.6636 | 1.6511 | 1.7962 | 1.6869 | 1.7015 | 1.6547 |
| 5. BMC - *Alaria americana* (2) | 1.6839 | 1.792 | 1.835 | 1.683 | 1.6785 | 1.7312 | 1.7501 | 1.8274 | 1.8594 | 1.6781 | 1.7299 | 1.6915 | 1.7059 | 1.7991 |
| 6. BMC - *Diplostomum indistinctum* | 1.4797 | 0.5948 | 1.6329 | 1.6776 | 1.6947 | 1.7319 | 1.7035 | 1.6424 | 1.4648 | 1.4752 | 1.4952 | 1.636 | 1.7263 | 1.577 |
| 7. BMC - Diplostomidae gen. sp. X | 1.9201 | 1.9951 | 2.0929 | 2.1263 | 2.0612 | 1.9601 | 2.059 | 2.0611 | 1.9071 | 2.0556 | 2.0575 | 1.944 | 2.0307 | 2.108 |
| 8. BMC - *Diplostomum* sp. 20 | 1.894 | 1.8545 | 1.9438 | 1.9614 | 1.9 | 1.7617 | 1.8365 | 1.88 | 2.0231 | 1.9156 | 1.8443 | 1.8817 | 2.0058 | 1.9572 |
| 9. BMC - *Diplostomum scudderi* (2) | 1.5898 | 1.7066 | 1.6538 | 1.6482 | 1.5402 | 1.6049 | 1.6118 | 0.4055 | 1.5648 | 1.5742 | 1.6286 | 1.5039 | 1.6745 | 1.7425 |
| 10. BMC - *Diplostomum* sp. 21 (2) | 1.8336 | 1.9062 | 1.8698 | 1.7848 | 1.8018 | 1.8612 | 1.8179 | 1.7672 | 1.874 | 1.7724 | 1.8079 | 1.7762 | 1.9018 | 1.8894 |
| 11. BMC - *Diplostomum marshalli* | 1.6654 | 1.7313 | 1.6694 | 1.6383 | 1.7717 | 1.7393 | 1.5519 | 1.5738 | 1.7374 | 1.6094 | 1.6144 | 1.6471 | 1.7479 | 1.6571 |
| 12. BMC - *Diplostomum* sp. 3 | 0.3717 | 1.498 | 1.7165 | 1.6314 | 1.6069 | 1.6781 | 1.7326 | 1.5912 | 1.6865 | 1.5112 | 1.5989 | 1.5788 | 1.645 | 1.5765 |
| 13. BMC - *Diplostomum* sp. VVT4 | 1.7163 | 1.6741 | 1.7009 | 1.5988 | 1.6402 | 1.6164 | 1.5878 | 1.3461 | 1.6911 | 1.5369 | 1.628 | 1.138 | 1.6365 | 1.5938 |
| 14. BMC - *Diplostomum huronense* | 1.5705 | 1.7892 | 1.686 | 1.5863 | 1.6835 | 1.6492 | 1.6827 | 1.5766 | 1.6316 | 1.6363 | 1.5473 | 1.5687 | 1.6239 | 1.5811 |
| 15. Diplostomidae gen. sp. O | 1.9518 | 2.0418 | 2.0831 | 2.0495 | 2.0218 | 2.0434 | 2.1249 | 2.1029 | 1.9761 | 2.0314 | 2.0718 | 2.0316 | 2.0343 | 2.0829 |
| 16. Diplostomidae gen. sp. X | 1.8964 | 1.9724 | 2.0869 | 2.1251 | 2.0651 | 1.9669 | 2.0465 | 2.0509 | 1.913 | 2.0374 | 2.0168 | 1.9446 | 2.0182 | 2.1043 |
| 17. Diplostomoidea sp. | 1.9043 | 2.038 | 1.857 | 1.8653 | 1.9882 | 1.9569 | 1.7778 | 1.9249 | 1.9268 | 1.8602 | 1.8567 | 1.8869 | 1.918 | 1.8583 |
| 18. D*iplostomum ardeae* | 1.557 | 1.6762 | 1.5076 | 1.4379 | 1.5777 | 1.5553 | 1.6169 | 1.6085 | 1.6637 | 1.489 | 1.6025 | 1.5489 | 1.6999 | 1.6267 |
| 19. *Diplostomum baeri* | 1.7333 | 1.7623 | 1.7162 | 1.6793 | 1.7427 | 1.7031 | 1.6673 | 1.604 | 1.6035 | 1.4736 | 1.5201 | 1.5903 | 1.6648 | 1.7447 |
| 20. *Diplostomum gavium* | 0.2653 | 1.5125 | 1.7036 | 1.6222 | 1.6149 | 1.6893 | 1.7319 | 1.6227 | 1.6894 | 1.5236 | 1.616 | 1.5837 | 1.6289 | 1.5348 |
| 21. *Diplostomum huronense* | 1.5125 | 1.6922 | 1.6776 | 1.634 | 1.6064 | 1.6321 | 1.6532 | 1.5529 | 1.53 | 1.5639 | 1.5605 | 1.5319 | 1.6737 | 1.5054 |
| 22. *Diplostomum indistinctum* | 1.4983 | 0.2604 | 1.6409 | 1.7059 | 1.6747 | 1.7868 | 1.736 | 1.6803 | 1.519 | 1.4731 | 1.4864 | 1.6505 | 1.7282 | 1.6109 |
| 23. *Diplostomum marshalli* | 1.6882 | 1.6914 | 1.6614 | 1.608 | 1.7618 | 1.7214 | 1.5454 | 1.5955 | 1.6902 | 1.5706 | 1.5992 | 1.6386 | 1.7317 | 1.6022 |
| 24. *Diplostomum mergi* | 1.646 | 1.6079 | 1.6079 | 1.5828 | 1.6635 | 1.5831 | 1.572 | 1.6327 | 1.6394 | 1.4631 | 1.5726 | 1.6042 | 1.7167 | 1.5494 |
| 25. *Diplostomum pseudospathaceum* | 0.8644 | 1.4727 | 1.7183 | 1.6254 | 1.5056 | 1.6331 | 1.7409 | 1.5165 | 1.573 | 1.4546 | 1.5862 | 1.5176 | 1.6247 | 1.5445 |

*Continued on next page*

Table B1 – *Continued from previous page*

| | 43 | 44 | 45 | 46 | 47 | 48 | 49 | 50 | 51 | 52 | 53 | 54 | 55 | 56 |
|---|---|---|---|---|---|---|---|---|---|---|---|---|---|---|
| 1. *Alaria americana* | 1.7641 | 1.8781 | 1.8552 | 1.8542 | 1.8564 | 1.7675 | 1.7303 | 1.6239 | 1.7683 | 2.014 | 1.8547 | 1.8608 | 1.7956 | 1.6664 |
| 2. *Alaria* sp. 1 | 1.8319 | 1.769 | 1.7478 | 1.8131 | 1.743 | 1.7551 | 1.8036 | 1.619 | 1.7188 | 2.0289 | 1.8544 | 1.8402 | 1.7852 | 1.7484 |
| 3. *Alaria* sp. 2 | 1.8622 | 1.8818 | 1.7772 | 1.7898 | 1.8251 | 1.8044 | 1.8511 | 1.7073 | 1.7909 | 2.0824 | 1.9191 | 1.996 | 1.8891 | 1.7098 |
| 4. *Austrodiplostomum ostrowskiae* | 1.7474 | 1.7582 | 1.6873 | 1.7581 | 1.667 | 1.6901 | 1.6836 | 1.6337 | 1.6701 | 1.9483 | 1.6573 | 1.6862 | 1.6099 | 1.6306 |
| 5. BMC - *Alaria americana* (2) | 1.8242 | 1.7989 | 1.768 | 1.7641 | 1.79 | 1.7396 | 1.7699 | 1.6612 | 1.6589 | 1.9798 | 1.9032 | 1.8016 | 1.7717 | 1.6756 |
| 6. BMC - *Diplostomum indistinctum* | 1.7128 | 1.591 | 1.6934 | 1.7327 | 1.5485 | 1.7055 | 1.7134 | 1.6522 | 1.4902 | 2.0197 | 1.838 | 1.7993 | 1.661 | 1.6858 |
| 7. BMC - Diplostomidae gen. sp. X | 2.046 | 1.9741 | 2.0707 | 2.0779 | 2.0407 | 1.9437 | 2.0891 | 1.9768 | 1.8764 | 1.7664 | 2.0411 | 1.8817 | 1.9346 | 2.1277 |
| 8. BMC - *Diplostomum* sp. 20 | 1.8909 | 1.7255 | 1.8825 | 1.9904 | 1.8798 | 1.7848 | 1.8896 | 1.8964 | 1.8918 | 2.0635 | 1.8712 | 1.9444 | 1.9209 | 1.9299 |
| 9. BMC - *Diplostomum scudderi* (2) | 1.5269 | 1.5806 | 0.4034 | 1.4129 | 1.569 | 1.6303 | 1.4448 | 1.5745 | 1.5871 | 1.9989 | 1.6323 | 1.8745 | 1.7009 | 1.7657 |
| 10. BMC - *Diplostomum* sp. 21 (2) | 1.6654 | 1.7462 | 1.785 | 1.8848 | 1.8361 | 1.8429 | 1.7888 | 1.7048 | 1.7744 | 1.9706 | 1.8723 | 1.8943 | 1.7529 | 1.8656 |
| 11. BMC - *Diplostomum marshalli* | 0.3807 | 1.7204 | 1.5697 | 1.8028 | 1.6788 | 1.7536 | 1.6329 | 1.6326 | 1.6274 | 2.117 | 1.7754 | 1.8414 | 1.763 | 1.7354 |
| 12. BMC - *Diplostomum* sp. 3 | 1.6543 | 1.5042 | 1.5862 | 1.6578 | 1.673 | 1.7112 | 1.756 | 1.5572 | 1.4374 | 2.0122 | 1.7229 | 1.7974 | 1.6604 | 1.8073 |
| 13. BMC - *Diplostomum* sp. VVT4 | 1.6713 | 1.6536 | 1.3917 | 1.5781 | 1.6199 | 1.6138 | 0.4333 | 1.5829 | 1.6411 | 2.1002 | 1.7326 | 1.8031 | 1.5751 | 1.6398 |
| 14. BMC - *Diplostomum huronense* | 1.5617 | 1.6574 | 1.5098 | 1.7743 | 1.7117 | 1.6253 | 1.4878 | 1.6389 | 1.5362 | 2.1111 | 1.7489 | 1.8882 | 1.5936 | 1.7797 |
| 15. Diplostomidae gen. sp. O | 2.0344 | 1.9821 | 2.084 | 2.0246 | 2.1142 | 2.0395 | 2.0287 | 1.95 | 1.8747 | 1.739 | 1.9875 | 1.9339 | 1.9851 | 2.0087 |
| 16. Diplostomidae gen. sp. X | 2.0575 | 1.9882 | 2.0607 | 2.0735 | 2.0369 | 1.948 | 2.094 | 2.0042 | 1.8845 | 1.791 | 2.0323 | 1.8867 | 1.9241 | 2.117 |
| 17. Diplostomoidea sp. | 1.9123 | 1.9057 | 1.9209 | 1.9671 | 1.9894 | 1.9393 | 1.8793 | 1.8135 | 1.8748 | 1.8987 | 1.7816 | 1.9352 | 1.9319 | 1.8584 |
| 18. Di*plostomum ardeae* | 1.5829 | 1.5051 | 1.5542 | 1.6623 | 1.679 | 1.572 | 1.5959 | 0.3704 | 1.54 | 1.9361 | 1.7217 | 1.7116 | 1.6081 | 1.6549 |
| 19. *Diplostomum baeri* | 1.5879 | 1.6534 | 1.6212 | 1.7419 | 1.6764 | 1.6763 | 1.5054 | 1.568 | 1.5788 | 1.9956 | 1.8967 | 1.8196 | 1.6917 | 1.6911 |
| 20. *Diplostomum gavium* | 1.6565 | 1.519 | 1.6155 | 1.6118 | 1.6856 | 1.7186 | 1.7678 | 1.5639 | 1.4577 | 2.0144 | 1.7311 | 1.8132 | 1.6355 | 1.8255 |
| 21. *Diplostomum huronense* | 1.5558 | 1.5899 | 1.4794 | 1.7383 | 1.6427 | 1.6244 | 1.4927 | 1.567 | 1.4358 | 2.0342 | 1.7509 | 1.8801 | 1.6245 | 1.7444 |
| 22. *Diplostomum indistinctum* | 1.7328 | 1.5754 | 1.734 | 1.7584 | 1.5905 | 1.7581 | 1.7201 | 1.657 | 1.4965 | 2.0052 | 1.8136 | 1.8091 | 1.6949 | 1.7147 |
| 23. *Diplostomum marshalli* | 0.5136 | 1.7078 | 1.593 | 1.8233 | 1.6691 | 1.7351 | 1.63 | 1.5981 | 1.6039 | 2.073 | 1.7478 | 1.7953 | 1.7625 | 1.7793 |
| 24. *Diplostomum mergi* | 1.5724 | 1.3658 | 1.6404 | 1.8649 | 1.6602 | 1.5709 | 1.587 | 1.5153 | 1.5724 | 1.9974 | 1.7534 | 1.6524 | 1.6718 | 1.7301 |
| 25. *Diplostomum pseudospathaceum* | 1.6436 | 1.4875 | 1.5251 | 1.5992 | 1.5599 | 1.6662 | 1.6517 | 1.5727 | 1.4039 | 1.9976 | 1.606 | 1.7293 | 1.5975 | 1.7615 |

*Continued on next page*

Table B1 – *Continued from previous page*

| | 1 | 2 | 3 | 4 | 5 | 6 | 7 | 8 | 9 | 10 | 11 | 12 | 13 |
|---|---|---|---|---|---|---|---|---|---|---|---|---|---|
| 26. *Diplostomum scudderi* | 85.417 | 85.938 | 85.938 | 85.156 | 85.807 | 88.802 | 79.167 | 84.115 | 99.609 | 85.026 | 89.063 | 89.583 | 90.365 |
| 27. *Diplostomum* sp. 1 (2) | 86.328 | 86.198 | 83.854 | 84.896 | 85.677 | 87.240 | 78.906 | 83.854 | 88.802 | 84.245 | 89.193 | 90.104 | 88.802 |
| 28. *Diplostomum* sp. 2 | 87.240 | 87.240 | 85.938 | 86.719 | 86.849 | 87.240 | 78.646 | 85.417 | 88.672 | 86.589 | 89.323 | 87.760 | 90.885 |
| 29. *Diplostomum* sp. 3 | 86.458 | 86.458 | 84.896 | 86.979 | 86.589 | 90.104 | 80.208 | 85.156 | 89.323 | 82.943 | 87.500 | 99.479 | 86.458 |
| 30. *Diplostomum* sp. 4 | 85.156 | 86.979 | 84.896 | 83.594 | 85.286 | 98.698 | 78.125 | 84.375 | 87.891 | 82.943 | 87.240 | 90.104 | 88.021 |
| 31. *Diplostomum* sp. 6 | 84.375 | 84.896 | 83.594 | 86.979 | 84.505 | 86.198 | 80.208 | 83.073 | 87.109 | 83.203 | 86.198 | 85.677 | 86.198 |
| 32. *Diplostomum* sp. 7 | 84.896 | 87.240 | 84.635 | 88.021 | 85.026 | 86.458 | 79.427 | 83.333 | 87.500 | 84.766 | 87.760 | 87.240 | 86.719 |
| 33. *Diplostomum* sp. 9 | 87.760 | 85.938 | 85.677 | 84.375 | 88.021 | 88.021 | 79.167 | 84.115 | 89.714 | 85.026 | 87.500 | 87.500 | 89.063 |
| 34. *Diplostomum* sp. 10 | 85.417 | 86.458 | 86.458 | 85.677 | 85.807 | 86.719 | 79.688 | 84.375 | 88.932 | 82.161 | 84.896 | 87.240 | 88.281 |
| 35. *Diplostomum* sp. 12 | 84.896 | 85.938 | 84.896 | 85.156 | 85.547 | 87.500 | 78.385 | 84.635 | 89.583 | 85.026 | 89.063 | 86.458 | 88.802 |
| 36. *Diplostomum* sp. 13 | 85.156 | 85.938 | 85.938 | 84.896 | 85.286 | 88.802 | 79.167 | 84.375 | 99.349 | 85.286 | 89.063 | 89.063 | 91.146 |
| 37. *Diplostomum* sp. 14 | 84.896 | 85.677 | 85.677 | 85.677 | 85.026 | 91.146 | 79.427 | 81.771 | 88.932 | 83.724 | 87.240 | 88.021 | 86.979 |
| 38. *Diplostomum* sp. 15 | 86.719 | 89.323 | 87.760 | 86.719 | 87.109 | 90.365 | 79.427 | 84.375 | 89.974 | 84.245 | 88.542 | 89.583 | 90.625 |
| 39. *Diplostomum* sp. 16 | 85.938 | 85.938 | 84.896 | 83.594 | 85.547 | 89.844 | 77.865 | 85.156 | 88.411 | 84.245 | 87.760 | 88.542 | 87.240 |
| 40. *Diplostomum* sp. 17 | 84.896 | 86.719 | 84.896 | 86.458 | 85.286 | 88.281 | 79.948 | 84.375 | 89.974 | 83.203 | 86.979 | 87.760 | 94.271 |
| 41. *Diplostomum* sp. 18 | 84.635 | 85.156 | 84.375 | 85.417 | 84.505 | 85.677 | 78.646 | 82.292 | 88.281 | 83.203 | 86.719 | 86.719 | 88.281 |
| 42. *Diplostomum* sp. 19 | 86.198 | 86.198 | 84.635 | 86.719 | 85.807 | 89.063 | 78.385 | 84.115 | 88.021 | 83.984 | 87.500 | 89.323 | 87.500 |
| 43. *Diplostomum* sp. A | 85.156 | 85.156 | 84.115 | 86.198 | 84.766 | 87.760 | 80.208 | 84.635 | 89.583 | 87.370 | 99.479 | 87.760 | 87.500 |
| 44. *Diplostomum* sp. B | 84.115 | 85.677 | 85.677 | 84.375 | 84.505 | 89.063 | 79.948 | 85.417 | 89.974 | 85.286 | 87.240 | 90.104 | 88.281 |
| 45. *Diplostomum* sp. C | 85.417 | 86.198 | 86.458 | 85.417 | 85.807 | 88.281 | 78.906 | 84.375 | 99.349 | 85.026 | 89.063 | 89.063 | 90.885 |
| 46. *Diplostomum* sp. VVT1 | 85.417 | 85.677 | 85.938 | 84.115 | 85.807 | 86.458 | 78.646 | 82.031 | 92.318 | 82.943 | 85.677 | 86.719 | 89.063 |
| 47. *Diplostomum* sp. VVT2 | 84.115 | 85.938 | 85.156 | 86.458 | 84.505 | 89.583 | 78.646 | 84.375 | 88.932 | 84.245 | 87.240 | 87.500 | 86.719 |
| 48. *Diplostomum* sp. VVT3 | 85.417 | 86.198 | 86.198 | 85.677 | 85.547 | 86.979 | 79.688 | 83.854 | 88.672 | 82.422 | 84.896 | 86.458 | 88.281 |
| 49. *Diplostomum* sp. VVT4 | 85.677 | 86.198 | 84.896 | 84.635 | 85.286 | 87.240 | 77.865 | 84.896 | 90.495 | 83.984 | 88.021 | 85.938 | 99.219 |
| 50. *Diplostomum* sp. VVT5 | 87.760 | 88.021 | 86.979 | 87.500 | 87.630 | 87.240 | 80.208 | 83.333 | 88.932 | 85.807 | 88.021 | 87.500 | 88.021 |
| 51. *Diplostomum spathaceum* | 85.938 | 85.938 | 85.677 | 85.156 | 86.328 | 90.365 | 80.729 | 83.073 | 88.411 | 83.984 | 89.063 | 90.104 | 88.021 |
| 52. *Ornithodiplostomum scardinii* (out) | 79.167 | 80.208 | 78.906 | 81.250 | 79.557 | 80.990 | 87.240 | 80.469 | 81.641 | 80.599 | 80.469 | 80.990 | 78.906 |

*Continued on next page*

Table B1 – *Continued from previous page*

| | 14 | 15 | 16 | 17 | 18 | 19 | 20 | 21 | 22 | 23 | 24 | 25 | 26 | 27 | 28 |
|---|---|---|---|---|---|---|---|---|---|---|---|---|---|---|---|
| 26. *Diplostomum scudderi* | 89.267 | 79.167 | 79.427 | 81.984 | 89.323 | 89.063 | 89.323 | 89.844 | 88.542 | 88.802 | 89.323 | 90.601 | - | 1.5422 | 1.6339 |
| 27. *Diplostomum* sp. 1 (2) | 99.476 | 77.344 | 78.906 | 81.984 | 86.979 | 87.500 | 89.844 | 97.396 | 86.979 | 88.932 | 89.323 | 91.384 | 89.063 | - | 1.5371 |
| 28. *Diplostomum* sp. 2 | 89.791 | 80.208 | 78.906 | 83.551 | 88.802 | 89.844 | 87.760 | 88.802 | 86.979 | 89.323 | 89.583 | 89.034 | 89.063 | 89.323 | - |
| 29. *Diplostomum* sp. 3 | 89.791 | 79.427 | 80.990 | 81.984 | 87.500 | 88.021 | 99.740 | 90.365 | 90.104 | 87.500 | 88.542 | 97.128 | 89.583 | 90.104 | 88.021 |
| 30. *Diplostomum* sp. 4 | 86.649 | 78.385 | 78.906 | 80.940 | 86.979 | 87.500 | 89.844 | 88.281 | 99.740 | 87.760 | 88.542 | 90.601 | 88.281 | 86.719 | 86.719 |
| 31. *Diplostomum* sp. 6 | 85.602 | 78.906 | 80.469 | 82.245 | 88.281 | 86.198 | 85.938 | 86.198 | 86.458 | 86.458 | 86.458 | 85.901 | 87.500 | 85.677 | 89.063 |
| 32. *Diplostomum* sp. 7 | 87.173 | 79.427 | 79.688 | 81.723 | 89.323 | 86.458 | 87.240 | 86.979 | 86.458 | 88.542 | 87.240 | 88.512 | 87.760 | 86.979 | 90.365 |
| 33. *Diplostomum* sp. 9 | 87.696 | 80.469 | 79.167 | 80.940 | 88.281 | 87.760 | 87.240 | 87.760 | 88.542 | 87.760 | 88.542 | 89.034 | 90.104 | 87.500 | 89.063 |
| 34. *Diplostomum* sp. 10 | 87.435 | 79.427 | 79.948 | 81.984 | 88.021 | 86.458 | 87.240 | 88.021 | 86.458 | 85.677 | 88.021 | 87.990 | 88.802 | 86.979 | 88.802 |
| 35. *Diplostomum* sp. 12 | 87.435 | 79.427 | 78.646 | 83.812 | 87.500 | 86.979 | 86.198 | 87.500 | 87.240 | 89.323 | 88.281 | 86.945 | 89.323 | 87.370 | 92.188 |
| 36. *Diplostomum* sp. 13 | 88.743 | 78.646 | 79.427 | 81.984 | 88.802 | 89.063 | 88.802 | 89.323 | 88.542 | 88.802 | 89.063 | 90.078 | 99.219 | 88.542 | 88.542 |
| 37. *Diplostomum* sp. 14 | 88.482 | 78.646 | 79.688 | 82.507 | 86.719 | 88.542 | 87.760 | 90.104 | 90.885 | 87.760 | 88.802 | 89.034 | 89.323 | 88.281 | 88.021 |
| 38. *Diplostomum* sp. 15 | 88.482 | 78.906 | 79.688 | 84.334 | 89.844 | 90.885 | 89.323 | 89.583 | 90.365 | 89.583 | 92.188 | 90.078 | 89.844 | 88.542 | 90.104 |
| 39. *Diplostomum* sp. 16 | 87.958 | 78.906 | 78.646 | 82.507 | 86.979 | 89.583 | 88.281 | 88.281 | 89.844 | 88.281 | 88.542 | 89.034 | 88.802 | 88.021 | 88.542 |
| 40. *Diplostomum* sp. 17 | 88.482 | 78.646 | 80.208 | 82.507 | 88.542 | 88.281 | 87.500 | 89.063 | 88.021 | 87.500 | 88.281 | 88.512 | 90.365 | 88.281 | 89.063 |
| 41. *Diplostomum* sp. 18 | 87.958 | 79.427 | 78.906 | 81.723 | 86.198 | 86.979 | 86.458 | 87.500 | 85.677 | 87.240 | 87.240 | 87.206 | 88.542 | 87.500 | 90.365 |
| 42. *Diplostomum* sp. 19 | 89.005 | 77.865 | 78.646 | 81.723 | 86.198 | 86.458 | 89.583 | 91.146 | 88.802 | 88.281 | 88.281 | 90.078 | 88.281 | 89.453 | 91.927 |
| 43. *Diplostomum* sp. A | 89.791 | 80.208 | 80.208 | 83.029 | 88.542 | 88.281 | 87.500 | 89.063 | 87.500 | 98.958 | 89.323 | 88.251 | 89.323 | 89.453 | 89.063 |
| 44. *Diplostomum* sp. B | 88.220 | 80.208 | 80.208 | 82.245 | 88.802 | 88.281 | 89.844 | 89.583 | 89.323 | 87.500 | 91.927 | 90.601 | 90.365 | 88.281 | 88.542 |
| 45. *Diplostomum* sp. C | 89.529 | 78.906 | 79.167 | 81.723 | 89.063 | 88.802 | 88.802 | 90.104 | 88.021 | 88.802 | 88.802 | 89.817 | 99.219 | 89.323 | 88.802 |
| 46. *Diplostomum* sp. VVT1 | 86.649 | 79.427 | 78.906 | 80.679 | 87.240 | 87.500 | 86.979 | 87.240 | 86.458 | 85.156 | 86.198 | 87.467 | 92.708 | 86.458 | 89.063 |
| 47. *Diplostomum* sp. VVT2 | 86.387 | 77.604 | 78.906 | 80.679 | 86.719 | 87.240 | 87.240 | 87.240 | 89.323 | 87.760 | 87.240 | 89.034 | 89.323 | 86.458 | 87.240 |
| 48. *Diplostomum* sp. VVT3 | 87.435 | 79.427 | 79.948 | 82.245 | 87.760 | 86.719 | 86.458 | 88.021 | 86.719 | 85.677 | 88.281 | 87.206 | 88.542 | 86.979 | 88.802 |
| 49. *Diplostomum* sp. VVT4 | 89.267 | 78.646 | 78.125 | 81.462 | 87.760 | 89.583 | 85.677 | 88.802 | 87.240 | 88.021 | 88.281 | 87.206 | 90.365 | 89.063 | 90.885 |
| 50. *Diplostomum* sp. VVT5 | 87.435 | 81.250 | 80.469 | 83.551 | 99.479 | 88.021 | 87.240 | 87.500 | 87.240 | 88.281 | 89.323 | 87.990 | 89.323 | 86.979 | 88.281 |
| 51. *Diplostomum spathaceum* | 90.576 | 80.990 | 80.729 | 83.290 | 88.542 | 88.542 | 90.365 | 91.667 | 90.365 | 89.323 | 89.583 | 91.123 | 88.802 | 90.365 | 89.063 |
| 52. *Ornithodiplostomum scardinii* (out) | 79.058 | 87.760 | 86.979 | 81.201 | 81.771 | 81.250 | 80.990 | 80.469 | 81.250 | 81.250 | 82.031 | 81.462 | 82.031 | 78.906 | 81.510 |

*Continued on next page*

Table B1 – *Continued from previous page*

| | 29 | 30 | 31 | 32 | 33 | 34 | 35 | 36 | 37 | 38 | 39 | 40 | 41 | 42 |
|---|---|---|---|---|---|---|---|---|---|---|---|---|---|---|
| 26. *Diplostomum scudderi* | 1.5853 | 1.695 | 1.6247 | 1.6278 | 1.5213 | 1.618 | 1.6216 | 0.467 | 1.547 | 1.5819 | 1.63 | 1.4998 | 1.6658 | 1.7277 |
| 27. *Diplostomum* sp. 1 (2) | 1.5474 | 1.7784 | 1.6775 | 1.5835 | 1.6579 | 1.6405 | 1.6647 | 1.5657 | 1.643 | 1.6263 | 1.5388 | 1.5417 | 1.6277 | 1.5729 |
| 28. *Diplostomum* sp. 2 | 1.6309 | 1.7454 | 1.4873 | 1.4248 | 1.6575 | 1.5715 | 1.2874 | 1.6927 | 1.6707 | 1.5191 | 1.5648 | 1.5779 | 1.4929 | 1.3412 |
| 29. *Diplostomum* sp. 3 | - | 1.4983 | 1.7029 | 1.6188 | 1.5987 | 1.6796 | 1.7453 | 1.6123 | 1.6918 | 1.5135 | 1.6079 | 1.5738 | 1.6459 | 1.5427 |
| 30. *Diplostomum* sp. 4 | 90.104 | - | 1.6613 | 1.7328 | 1.7043 | 1.7868 | 1.7604 | 1.6882 | 1.537 | 1.4872 | 1.5305 | 1.6354 | 1.7498 | 1.6351 |
| 31. *Diplostomum* sp. 6 | 86.198 | 86.198 | - | 1.4175 | 1.7377 | 1.5572 | 1.4329 | 1.7061 | 1.679 | 1.5542 | 1.6489 | 1.6086 | 1.6295 | 1.5371 |
| 32. *Diplostomum* sp. 7 | 87.500 | 86.198 | 91.667 | - | 1.579 | 1.607 | 1.5988 | 1.6641 | 1.6768 | 1.5663 | 1.6105 | 1.5819 | 1.5155 | 1.5788 |
| 33. *Diplostomum* sp. 9 | 87.500 | 88.281 | 84.896 | 88.021 | - | 1.7158 | 1.7821 | 1.5636 | 1.6746 | 1.607 | 1.6398 | 1.6506 | 1.6894 | 1.6864 |
| 34. *Diplostomum* sp. 10 | 87.500 | 86.458 | 88.021 | 87.500 | 86.719 | - | 1.4846 | 1.6382 | 1.688 | 1.433 | 1.7463 | 1.5813 | 1.6411 | 1.6016 |
| 35. *Diplostomum* sp. 12 | 86.458 | 86.979 | 90.365 | 89.063 | 86.979 | 88.802 | - | 1.648 | 1.6649 | 1.5456 | 1.6002 | 1.5929 | 1.4549 | 1.3479 |
| 36. *Diplostomum* sp. 13 | 89.063 | 88.281 | 86.719 | 87.240 | 89.583 | 88.542 | 89.323 | - | 1.5811 | 1.5834 | 1.6296 | 1.5247 | 1.6745 | 1.7669 |
| 37. *Diplostomum* sp. 14 | 88.021 | 90.625 | 85.938 | 86.198 | 87.500 | 86.458 | 87.500 | 88.802 | - | 1.4724 | 1.5704 | 1.6275 | 1.701 | 1.6674 |
| 38. *Diplostomum* sp. 15 | 89.583 | 90.104 | 88.542 | 89.323 | 88.542 | 90.365 | 89.844 | 89.844 | 90.365 | - | 1.4369 | 1.4587 | 1.6034 | 1.5598 |
| 39. *Diplostomum* sp. 16 | 88.542 | 89.583 | 85.938 | 86.979 | 88.542 | 84.375 | 88.802 | 88.542 | 88.802 | 89.583 | - | 1.6635 | 1.632 | 1.4515 |
| 40. *Diplostomum* sp. 17 | 87.760 | 88.281 | 87.500 | 88.021 | 88.281 | 88.021 | 88.281 | 89.844 | 87.240 | 90.625 | 87.500 | - | 1.5902 | 1.6014 |
| 41. *Diplostomum* sp. 18 | 86.719 | 85.417 | 86.979 | 88.802 | 86.719 | 86.719 | 91.667 | 88.281 | 86.458 | 88.802 | 87.760 | 88.802 | - | 1.5135 |
| 42. *Diplostomum* sp. 19 | 89.844 | 88.542 | 90.365 | 89.063 | 87.500 | 87.760 | 91.927 | 87.760 | 88.021 | 89.323 | 90.104 | 88.281 | 90.365 | - |
| 43. *Diplostomum* sp. A | 87.760 | 87.240 | 86.198 | 87.760 | 87.500 | 85.156 | 88.802 | 89.323 | 86.979 | 88.802 | 87.500 | 86.979 | 87.240 | 87.240 |
| 44. *Diplostomum* sp. B | 90.104 | 89.063 | 88.281 | 86.458 | 86.719 | 88.281 | 88.021 | 89.844 | 88.281 | 91.667 | 88.542 | 88.542 | 86.198 | 88.542 |
| 45. *Diplostomum* sp. C | 89.063 | 87.760 | 86.979 | 87.500 | 90.104 | 89.583 | 89.583 | 98.958 | 88.802 | 90.365 | 88.281 | 90.365 | 88.281 | 88.281 |
| 46. *Diplostomum* sp. VVT1 | 87.240 | 86.198 | 87.500 | 87.240 | 90.104 | 88.802 | 87.500 | 91.927 | 87.240 | 88.542 | 88.542 | 89.583 | 89.063 | 89.063 |
| 47. *Diplostomum* sp. VVT2 | 87.500 | 89.323 | 89.323 | 90.104 | 85.417 | 87.760 | 87.760 | 88.542 | 88.021 | 87.500 | 88.542 | 88.021 | 86.458 | 88.281 |
| 48. *Diplostomum* sp. VVT3 | 86.719 | 86.458 | 87.760 | 87.500 | 86.458 | 99.219 | 88.802 | 88.802 | 86.458 | 90.104 | 84.375 | 87.500 | 86.719 | 87.760 |
| 49. *Diplostomum* sp. VVT4 | 85.938 | 87.500 | 85.677 | 86.198 | 88.802 | 87.760 | 88.281 | 91.146 | 86.979 | 90.104 | 87.500 | 94.010 | 88.021 | 87.760 |
| 50. *Diplostomum* sp. VVT5 | 87.500 | 86.979 | 88.281 | 89.323 | 87.760 | 88.021 | 87.500 | 88.802 | 86.719 | 89.844 | 86.979 | 88.542 | 86.198 | 86.198 |
| 51. *Diplostomum spathaceum* | 90.104 | 90.104 | 86.979 | 87.240 | 90.365 | 88.021 | 88.021 | 88.281 | 91.927 | 90.885 | 90.365 | 88.281 | 87.240 | 89.063 |
| 52. *Ornithodiplostomum scardinii* (out) | 81.250 | 80.990 | 81.771 | 81.771 | 81.771 | 82.292 | 81.510 | 81.250 | 82.552 | 82.031 | 81.250 | 80.469 | 79.948 | 80.729 |

*Continued on next page*

Table B1 – *Continued from previous page*

| | 43 | 44 | 45 | 46 | 47 | 48 | 49 | 50 | 51 | 52 | 53 | 54 | 55 | 56 |
|---|---|---|---|---|---|---|---|---|---|---|---|---|---|---|
| 26. *Diplostomum scudderi* | 1.5397 | 1.5587 | 0.4584 | 1.3856 | 1.5707 | 1.6401 | 1.4499 | 1.544 | 1.5769 | 1.9673 | 1.6302 | 1.8568 | 1.6895 | 1.7687 |
| 27. *Diplostomum* sp. 1 (2) | 1.5541 | 1.6467 | 1.5003 | 1.7513 | 1.7005 | 1.623 | 1.4664 | 1.6475 | 1.5123 | 2.0976 | 1.7251 | 1.8566 | 1.5915 | 1.7603 |
| 28. *Diplostomum* sp. 2 | 1.6246 | 1.5786 | 1.6516 | 1.6158 | 1.6698 | 1.5663 | 1.4147 | 1.6075 | 1.5114 | 1.9891 | 1.7016 | 1.7724 | 1.6828 | 1.6872 |
| 29. *Diplostomum* sp. 3 | 1.6622 | 1.5128 | 1.6006 | 1.6199 | 1.6832 | 1.7146 | 1.7602 | 1.5592 | 1.4517 | 2.0124 | 1.7247 | 1.791 | 1.6325 | 1.8139 |
| 30. *Diplostomum* sp. 4 | 1.7313 | 1.5982 | 1.7389 | 1.7837 | 1.5905 | 1.7768 | 1.7122 | 1.6755 | 1.5204 | 2.0326 | 1.8446 | 1.826 | 1.7333 | 1.7388 |
| 31. *Diplostomum* sp. 6 | 1.6694 | 1.5184 | 1.6733 | 1.654 | 1.5732 | 1.5919 | 1.7323 | 1.4995 | 1.5972 | 1.9371 | 1.7683 | 1.8078 | 1.6509 | 1.7018 |
| 32. *Diplostomum* sp. 7 | 1.6383 | 1.6937 | 1.6246 | 1.5261 | 1.5491 | 1.5958 | 1.6161 | 1.454 | 1.6209 | 2.0076 | 1.7044 | 1.7493 | 1.6625 | 1.6853 |
| 33. *Diplostomum* sp. 9 | 1.7717 | 1.6631 | 1.5108 | 1.5065 | 1.7824 | 1.7076 | 1.6803 | 1.6205 | 1.4926 | 1.9196 | 1.7469 | 1.9545 | 1.7336 | 1.7903 |
| 34. *Diplostomum* sp. 10 | 1.7412 | 1.5905 | 1.5504 | 1.6727 | 1.555 | 0.4654 | 1.6449 | 1.5699 | 1.6365 | 1.9544 | 1.8193 | 1.7702 | 1.6645 | 1.776 |
| 35. *Diplostomum* sp. 12 | 1.5947 | 1.5665 | 1.6118 | 1.738 | 1.5914 | 1.4839 | 1.6262 | 1.6249 | 1.5928 | 1.9942 | 1.7072 | 1.7974 | 1.7084 | 1.7245 |
| 36. *Diplostomum* sp. 13 | 1.5628 | 1.577 | 0.5307 | 1.4634 | 1.6227 | 1.6293 | 1.3728 | 1.5943 | 1.603 | 2.0224 | 1.6485 | 1.8596 | 1.6994 | 1.798 |
| 37. *Diplostomum* sp. 14 | 1.765 | 1.5976 | 1.5825 | 1.653 | 1.6176 | 1.6827 | 1.7083 | 1.6948 | 1.389 | 1.8459 | 1.7713 | 1.7853 | 1.7268 | 1.7109 |
| 38. *Diplostomum* sp. 15 | 1.6103 | 1.3892 | 1.5614 | 1.614 | 1.6234 | 1.4492 | 1.5934 | 1.5163 | 1.3674 | 1.9545 | 1.7105 | 1.6765 | 1.6025 | 1.5862 |
| 39. *Diplostomum* sp. 16 | 1.634 | 1.6289 | 1.6589 | 1.5791 | 1.5627 | 1.7433 | 1.5929 | 1.6131 | 1.4764 | 1.9614 | 1.7144 | 1.8407 | 1.6868 | 1.5299 |
| 40. *Diplostomum* sp. 17 | 1.6471 | 1.6481 | 1.4899 | 1.5306 | 1.5601 | 1.5841 | 1.2098 | 1.5624 | 1.5939 | 1.9765 | 1.7156 | 1.7161 | 1.5723 | 1.6196 |
| 41. *Diplostomum* sp. 18 | 1.6872 | 1.8033 | 1.6781 | 1.5431 | 1.7281 | 1.6219 | 1.6601 | 1.7326 | 1.5819 | 2.0114 | 1.7854 | 1.7484 | 1.7293 | 1.7651 |
| 42. *Diplostomum* sp. 19 | 1.6943 | 1.6422 | 1.7196 | 1.5719 | 1.6459 | 1.6165 | 1.5454 | 1.6436 | 1.5872 | 2.027 | 1.7052 | 1.8625 | 1.7963 | 1.7658 |
| 43. *Diplostomum* sp. A | - | 1.7124 | 1.5618 | 1.8028 | 1.6788 | 1.7538 | 1.6329 | 1.6326 | 1.6274 | 2.1023 | 1.7836 | 1.8304 | 1.7457 | 1.7354 |
| 44. *Diplostomum* sp. B | 87.500 | - | 1.5995 | 1.7676 | 1.6469 | 1.6023 | 1.6641 | 1.5171 | 1.4939 | 1.9363 | 1.7568 | 1.7178 | 1.6157 | 1.6623 |
| 45. *Diplostomum* sp. C | 89.323 | 89.844 | - | 1.3999 | 1.5549 | 1.5676 | 1.4184 | 1.5423 | 1.593 | 2.0104 | 1.6592 | 1.8882 | 1.7179 | 1.7672 |
| 46. *Diplostomum* sp. VVT1 | 85.677 | 87.240 | 92.448 | - | 1.6912 | 1.6791 | 1.6297 | 1.7124 | 1.5131 | 1.9539 | 1.6899 | 1.8856 | 1.7298 | 1.7151 |
| 47. *Diplostomum* sp. VVT2 | 87.240 | 87.760 | 89.063 | 86.979 | - | 1.6021 | 1.6485 | 1.6871 | 1.6729 | 2.0329 | 1.7065 | 1.7507 | 1.6391 | 1.738 |
| 48. *Diplostomum* sp. VVT3 | 85.156 | 88.021 | 89.323 | 88.542 | 87.240 | - | 1.6393 | 1.5949 | 1.6353 | 1.9341 | 1.8107 | 1.7815 | 1.6566 | 1.798 |
| 49. *Diplostomum* sp. VVT4 | 88.021 | 88.281 | 90.885 | 88.802 | 86.719 | 87.760 | - | 1.5711 | 1.674 | 2.1171 | 1.706 | 1.8327 | 1.6064 | 1.6121 |
| 50. *Diplostomum* sp. VVT5 | 88.021 | 88.802 | 89.063 | 86.719 | 86.719 | 87.760 | 87.760 | - | 1.5696 | 1.9441 | 1.7045 | 1.7199 | 1.6303 | 1.6718 |
| 51. *Diplostomum spathaceum* | 89.063 | 90.104 | 88.542 | 88.542 | 86.719 | 88.021 | 87.760 | 88.542 | - | 1.9155 | 1.692 | 1.7792 | 1.5295 | 1.7095 |
| 52. *Ornithodiplostomum scardinii* (out) | 80.729 | 81.771 | 81.250 | 80.729 | 79.948 | 82.292 | 78.646 | 81.771 | 83.333 | - | 2.0224 | 1.9355 | 1.9255 | 1.9431 |

*Continued on next page*

Table B1 – *Continued from previous page*

| | 1 | 2 | 3 | 4 | 5 | 6 | 7 | 8 | 9 | 10 | 11 | 12 | 13 |
|---|---|---|---|---|---|---|---|---|---|---|---|---|---|
| *53. Tylodelphys aztecae* | 84.375 | 82.292 | 82.552 | 84.896 | 83.724 | 85.417 | 78.125 | 82.813 | 87.500 | 82.422 | 84.375 | 86.198 | 85.417 |
| *54. Tylodelphys clavata* | 83.333 | 84.375 | 83.333 | 85.156 | 83.464 | 83.073 | 80.990 | 82.813 | 83.594 | 82.682 | 84.896 | 83.333 | 85.417 |
| *55. Tylodelphys excavata* | 85.677 | 84.896 | 84.635 | 88.281 | 85.547 | 85.156 | 81.510 | 82.292 | 86.068 | 85.286 | 84.896 | 86.458 | 85.677 |
| *56. Tylodelphys immer* | 87.240 | 85.417 | 86.198 | 86.979 | 87.109 | 86.979 | 77.083 | 82.813 | 85.286 | 82.161 | 86.198 | 84.896 | 87.240 |

| | 14 | 15 | 16 | 17 | 18 | 19 | 20 | 21 | 22 | 23 | 24 | 25 | 26 | 27 | 28 |
|---|---|---|---|---|---|---|---|---|---|---|---|---|---|---|---|
| *53. Tylodelphys aztecae* | 84.817 | 79.167 | 78.385 | 80.940 | 87.240 | 84.375 | 85.938 | 85.417 | 85.677 | 84.896 | 84.375 | 87.990 | 87.760 | 85.156 | 85.677 |
| *54. Tylodelphys clavata* | 84.031 | 80.729 | 81.250 | 82.245 | 86.719 | 84.115 | 83.333 | 83.854 | 83.073 | 85.938 | 86.198 | 84.856 | 83.854 | 83.984 | 85.156 |
| *55. Tylodelphys excavata* | 87.435 | 79.688 | 81.771 | 81.462 | 87.760 | 86.198 | 86.458 | 87.240 | 85.156 | 84.635 | 86.719 | 87.990 | 86.458 | 87.891 | 85.677 |
| *56. Tylodelphys immer* | 85.864 | 78.385 | 77.083 | 83.290 | 87.240 | 86.719 | 84.635 | 85.677 | 86.719 | 85.677 | 86.458 | 85.117 | 85.156 | 86.068 | 85.677 |

| | 29 | 30 | 31 | 32 | 33 | 34 | 35 | 36 | 37 | 38 | 39 | 40 | 41 | 42 |
|---|---|---|---|---|---|---|---|---|---|---|---|---|---|---|
| *53. Tylodelphys aztecae* | 86.198 | 85.417 | 84.896 | 86.458 | 85.156 | 84.896 | 85.938 | 87.240 | 85.156 | 84.896 | 85.938 | 85.156 | 83.333 | 86.198 |
| *54. Tylodelphys clavata* | 83.594 | 82.813 | 84.115 | 85.156 | 82.552 | 85.417 | 83.854 | 83.854 | 83.854 | 86.979 | 82.552 | 86.198 | 84.375 | 83.594 |
| *55. Tylodelphys excavata* | 86.719 | 84.896 | 86.198 | 86.198 | 84.635 | 87.240 | 85.156 | 86.198 | 85.417 | 87.240 | 84.896 | 86.198 | 84.635 | 85.417 |
| *56. Tylodelphys immer* | 84.896 | 86.458 | 85.156 | 84.115 | 85.156 | 85.156 | 85.417 | 84.896 | 86.458 | 88.021 | 88.021 | 86.458 | 83.333 | 84.115 |

| | 43 | 44 | 45 | 46 | 47 | 48 | 49 | 50 | 51 | 52 | 53 | 54 | 55 | 56 |
|---|---|---|---|---|---|---|---|---|---|---|---|---|---|---|
| *53. Tylodelphys aztecae* | 84.115 | 85.677 | 87.240 | 85.938 | 86.719 | 85.156 | 85.417 | 87.500 | 86.458 | 80.208 | - | 1.7305 | 1.7248 | 1.6603 |
| *54. Tylodelphys clavata* | 85.156 | 86.198 | 83.333 | 82.031 | 83.594 | 85.938 | 85.156 | 86.719 | 84.896 | 80.729 | 84.635 | - | 1.5465 | 1.834 |
| *55. Tylodelphys excavata* | 85.156 | 87.500 | 85.938 | 84.896 | 86.719 | 87.500 | 85.417 | 87.240 | 86.979 | 80.990 | 86.458 | 88.802 | - | 1.7329 |
| *56. Tylodelphys immer* | 86.198 | 85.677 | 85.417 | 85.156 | 85.417 | 85.156 | 87.240 | 86.979 | 86.198 | 79.427 | 86.719 | 85.156 | 86.979 | - |

Table B2. Averaged percent similarities of the species belonging to the Diplostomidae II. Standard error estimate(s) are shown above the diagonal and were obtained by a bootstrap procedure (500 replicates). Ambiguous positions were removed for each sequence pair. Evolutionary analyses were conducted in MEGA (Tamura et al., 2021).

| | 1 | 2 | 3 | 4 | 5 | 6 | 7 | 8 | 9 | 10 | 11 | 12 |
|---|---|---|---|---|---|---|---|---|---|---|---|---|
| 1. BMC - *Bolbophorus* sp. A | - | 2.344 | 2.364 | 2.306 | 2.380 | 2.437 | 2.368 | 2.339 | 2.322 | 1.464 | 2.319 | 2.232 |
| 2. BMC - *Posthodiplostomum* sp. 4 | 78.146 | - | 1.937 | 1.886 | 2.064 | 2.075 | 2.087 | 1.952 | 2.015 | 2.108 | 2.265 | 2.372 |
| 3. BMC - *Posthodiplostomum* sp. 11 | 77.409 | 86.622 | - | 0.767 | 0.576 | 1.675 | 1.803 | 1.536 | 1.913 | 2.181 | 2.101 | 2.353 |
| 4. BMC - *Posthodiplostomum* sp. 12 (2) | 76.68 | 85.125 | 96.06 | - | 0.920 | 1.434 | 1.689 | 1.414 | 1.791 | 2.106 | 2.129 | 2.227 |
| 5. BMC - *Posthodiplostomum* sp. 11 | 75.908 | 85.382 | 99.003 | 95.25 | - | 1.761 | 1.865 | 1.606 | 1.967 | 2.227 | 2.162 | 2.363 |
| 6. BMC - Diplostomidae gen. sp. X | 75.908 | 85.05 | 91.362 | 91.284 | 90.429 | - | 1.817 | 1.677 | 2.086 | 2.246 | 2.327 | 2.294 |
| 7. BMC - *Posthodiplostomum* cf. *podicipitis* (2) | 77.815 | 84.385 | 88.333 | 88.052 | 87.417 | 88.079 | - | 1.744 | 1.954 | 2.165 | 2.221 | 2.262 |
| 8. BMC - *Posthodiplostomum ptychocheilus* | 75.908 | 85.714 | 92.691 | 91.096 | 91.419 | 90.759 | 89.073 | - | 1.887 | 2.160 | 2.170 | 2.310 |
| 9. BMC - *Posthodiplostomum* sp. 14 | 75.908 | 85.382 | 87.708 | 87.08 | 86.799 | 85.149 | 87.086 | 89.769 | - | 2.200 | 2.339 | 2.431 |
| 10. BMC - *Bolbophorus* sp. B (2) | 92.244 | 83.779 | 82.107 | 81.164 | 81.063 | 80.399 | 82.333 | 81.063 | 80.565 | - | 2.007 | 1.917 |
| 11. BMC - *Bolbophorus* sp. C | 80.602 | 81.544 | 82.77 | 80.355 | 81.879 | 79.195 | 81.481 | 82.215 | 79.195 | 85.642 | - | 1.953 |
| 12. *Bolbophorus damnificus* | 81.188 | 80.464 | 80.333 | 79.969 | 80.132 | 79.47 | 81.395 | 80.464 | 79.47 | 86.833 | 87.625 | - |
| 13. *Bolbophorus* sp. KM538081 | 81.188 | 81.457 | 82 | 79.61 | 81.126 | 79.139 | 81.063 | 81.457 | 78.808 | 86.5 | 98.328 | 87.129 |
| 14. *Bolbophorus* sp. KT831373 | 93.069 | 83.946 | 81.94 | 81.253 | 80.731 | 80.731 | 82.667 | 81.063 | 80.731 | 98.845 | 85.473 | 86.667 |
| 15. *Bolbophorus* sp. KU707938 | 89.439 | 82.274 | 82.943 | 82.095 | 81.728 | 80.399 | 83 | 81.728 | 81.395 | 95.847 | 85.135 | 86.667 |
| 16. *Bolbophorus* sp. MH368809 | 93.069 | 83.946 | 81.94 | 81.253 | 80.731 | 80.731 | 82.667 | 81.063 | 80.731 | 98.845 | 85.473 | 86.667 |
| 17. *Crocodillicola pseudostoma* (out) | 68.317 | 70.234 | 71.141 | 71.374 | 70 | 69.667 | 70.234 | 71.333 | 73.667 | 71.333 | 69.799 | 69.667 |
| 18. Diplostomidae gen. sp. O | 76.898 | 83.775 | 89.333 | 88.378 | 88.411 | 86.093 | 86.047 | 89.073 | 87.417 | 81.333 | 78.595 | 79.208 |
| 19. Diplostomidae gen. sp. X | 76.238 | 85.382 | 91.694 | 91.278 | 90.759 | 99.67 | 88.411 | 90.429 | 85.479 | 80.731 | 79.53 | 79.801 |
| 20. Diplostomoidea sp. | 77.483 | 80.066 | 80.602 | 79.042 | 80.399 | 79.734 | 79.402 | 82.06 | 80.066 | 81.438 | 81.818 | 82.06 |
| 21. *Neodiplostomum americanum* | 76.238 | 78.146 | 76 | 75.431 | 74.834 | 76.49 | 78.405 | 77.152 | 77.483 | 80.5 | 81.94 | 82.178 |
| 22. *Ornithodiplostomum scardinii* | 77.888 | 83.056 | 88.372 | 87.922 | 87.129 | 87.789 | 89.404 | 91.419 | 86.139 | 82.558 | 80.537 | 80.464 |
| 23. *Ornithodiplostomum* sp. 1 | 74.917 | 84.718 | 86.622 | 86.322 | 85.714 | 86.047 | 87 | 87.708 | 86.379 | 79.333 | 79.195 | 77.152 |
| 24. *Ornithodiplostomum* sp. 2 | 77.558 | 85.43 | 97.667 | 94.908 | 97.02 | 91.391 | 88.04 | 90.728 | 86.755 | 81.333 | 82.274 | 79.538 |
| 25. *Ornithodiplostomum* sp. 3 | 76.238 | 85.05 | 92.691 | 95.181 | 92.079 | 91.089 | 87.086 | 89.769 | 87.459 | 80.565 | 78.523 | 80.464 |

*Continued on next page*

Table B2 – *Continued from previous page*

| | 13 | 14 | 15 | 16 | 17 | 18 | 19 | 20 | 21 | 22 | 23 | 24 | 25 | 26 |
|---|---|---|---|---|---|---|---|---|---|---|---|---|---|---|
| 1. BMC - *Bolbophorus* sp. A | 2.208 | 1.448 | 1.793 | 1.448 | 2.554 | 2.329 | 2.445 | 2.298 | 2.385 | 2.360 | 2.412 | 2.379 | 2.403 | 2.307 |
| 2. BMC - *Posthodiplostomum* sp. 4 | 2.299 | 2.135 | 2.244 | 2.135 | 2.624 | 2.154 | 2.051 | 2.333 | 2.390 | 2.241 | 2.058 | 2.009 | 2.076 | 1.971 |
| 3. BMC - *Posthodiplostomum* sp. 11 | 2.172 | 2.227 | 2.151 | 2.227 | 2.706 | 1.893 | 1.644 | 2.346 | 2.486 | 1.884 | 1.848 | 0.856 | 1.569 | 1.876 |
| 4. BMC - *Posthodiplostomum* sp. 12 (2) | 2.233 | 2.149 | 2.044 | 2.149 | 2.602 | 1.816 | 1.435 | 2.293 | 2.420 | 1.755 | 1.711 | 0.961 | 1.005 | 1.764 |
| 5. BMC - *Posthodiplostomum* sp. 11 | 2.225 | 2.272 | 2.197 | 2.272 | 2.715 | 1.974 | 1.735 | 2.365 | 2.544 | 1.963 | 1.911 | 1.008 | 1.622 | 1.929 |
| 6. BMC - Diplostomidae gen. sp. X | 2.383 | 2.276 | 2.219 | 2.276 | 2.691 | 2.085 | 0.336 | 2.389 | 2.447 | 1.921 | 1.912 | 1.623 | 1.626 | 2.048 |
| 7. BMC - *Posthodiplostomum* cf. *podicipitis* (2) | 2.256 | 2.198 | 2.175 | 2.198 | 2.767 | 1.987 | 1.787 | 2.341 | 2.431 | 1.738 | 1.831 | 1.821 | 1.943 | 1.904 |
| 8. BMC - *Posthodiplostomum ptychocheilus* | 2.202 | 2.187 | 2.175 | 2.187 | 2.659 | 1.909 | 1.701 | 2.260 | 2.420 | 1.615 | 1.953 | 1.617 | 1.773 | 1.818 |
| 9. BMC - *Posthodiplostomum* sp. 14 | 2.359 | 2.230 | 2.189 | 2.230 | 2.607 | 1.937 | 2.066 | 2.300 | 2.527 | 1.980 | 2.003 | 1.900 | 1.890 | 0.459 |
| 10. BMC - *Bolbophorus* sp. B (2) | 1.942 | 0.485 | 1.149 | 0.485 | 2.532 | 2.190 | 2.256 | 2.099 | 2.252 | 2.141 | 2.283 | 2.234 | 2.236 | 2.169 |
| 11. BMC - *Bolbophorus* sp. C | 0.709 | 2.021 | 1.996 | 2.021 | 2.701 | 2.412 | 2.332 | 2.160 | 2.101 | 2.257 | 2.348 | 2.107 | 2.311 | 2.314 |
| 12. *Bolbophorus damnificus* | 2.003 | 1.959 | 1.932 | 1.959 | 2.661 | 2.418 | 2.303 | 2.224 | 2.189 | 2.310 | 2.429 | 2.334 | 2.337 | 2.400 |
| 13. *Bolbophorus* sp. KM538081 | - | 1.965 | 2.075 | 1.965 | 2.703 | 2.441 | 2.392 | 2.105 | 2.157 | 2.296 | 2.392 | 2.180 | 2.390 | 2.330 |
| 14. *Bolbophorus* sp. KT831373 | 86.667 | - | 1.132 | 0.000 | 2.525 | 2.229 | 2.286 | 2.098 | 2.263 | 2.181 | 2.306 | 2.266 | 2.274 | 2.204 |
| 15. *Bolbophorus* sp. KU707938 | 84.667 | 96.346 | - | 1.132 | 2.531 | 2.152 | 2.220 | 2.215 | 2.224 | 2.158 | 2.239 | 2.208 | 2.176 | 2.160 |
| 16. *Bolbophorus* sp. MH368809 | 86.667 | 100 | 96.346 | - | 2.525 | 2.229 | 2.286 | 2.098 | 2.263 | 2.181 | 2.306 | 2.266 | 2.274 | 2.204 |
| 17. *Crocodillicola pseudostoma* (out) | 69.667 | 71.667 | 71.096 | 71.667 | - | 2.744 | 2.680 | 2.614 | 2.555 | 2.713 | 2.659 | 2.672 | 2.603 | 2.613 |
| 18. Diplostomidae gen. sp. O | 78.218 | 81.333 | 82.333 | 81.333 | 70.667 | - | 2.054 | 2.250 | 2.549 | 1.762 | 1.985 | 1.926 | 1.963 | 1.921 |
| 19. Diplostomidae gen. sp. X | 79.47 | 81.063 | 80.731 | 81.063 | 70 | 86.424 | - | 2.394 | 2.431 | 1.950 | 1.945 | 1.598 | 1.649 | 2.028 |
| 20. Diplostomoidea sp. | 82.392 | 81.605 | 80.268 | 81.605 | 69.231 | 81.063 | 80.066 | - | 2.388 | 2.312 | 2.323 | 2.310 | 2.335 | 2.270 |
| 21. *Neodiplostomum americanum* | 81.848 | 80.667 | 81 | 80.667 | 72.667 | 75.908 | 76.821 | 78.738 | - | 2.419 | 2.455 | 2.437 | 2.529 | 2.504 |
| 22. *Ornithodiplostomum scardinii* | 80.795 | 82.724 | 82.724 | 82.724 | 70.667 | 88.079 | 87.459 | 81.063 | 78.477 | - | 2.054 | 1.937 | 1.908 | 1.955 |
| 23. *Ornithodiplostomum* sp. 1 | 78.808 | 79.333 | 81 | 79.333 | 71.333 | 86.093 | 85.714 | 77.667 | 76.821 | 86.379 | - | 1.837 | 1.857 | 2.000 |
| 24. *Ornithodiplostomum* sp. 2 | 81.518 | 81.667 | 81.667 | 81.667 | 72.333 | 88.119 | 91.722 | 80.066 | 76.238 | 87.086 | 86.755 | - | 1.473 | 1.859 |
| 25. *Ornithodiplostomum* sp. 3 | 78.146 | 80.731 | 81.728 | 80.731 | 73 | 88.411 | 90.759 | 79.402 | 75.828 | 87.459 | 86.047 | 92.384 | - | 1.878 |

*Continued on next page*

Table B2 – *Continued from previous page*

| | 27 | 28 | 29 | 30 | 31 | 32 | 33 | 34 | 35 | 36 | 37 | 38 | 39 |
|---|---|---|---|---|---|---|---|---|---|---|---|---|---|
| 1. BMC - *Bolbophorus* sp. A | 2.441 | 2.377 | 2.450 | 2.367 | 2.347 | 2.393 | 2.404 | 2.428 | 2.283 | 2.595 | 2.413 | 2.434 | 2.367 |
| 2. BMC - *Posthodiplostomum* sp. 4 | 2.045 | 2.099 | 2.106 | 1.239 | 1.959 | 2.085 | 2.086 | 2.048 | 0.662 | 2.126 | 2.272 | 2.036 | 1.984 |
| 3. BMC - *Posthodiplostomum* sp. 11 | 2.140 | 1.835 | 2.149 | 1.978 | 1.577 | 2.068 | 2.036 | 2.139 | 1.891 | 2.190 | 2.114 | 2.137 | 0.440 |
| 4. BMC - *Posthodiplostomum* sp. 12 (2) | 2.002 | 1.719 | 2.074 | 1.924 | 1.465 | 2.046 | 2.031 | 2.008 | 1.863 | 2.124 | 1.984 | 2.064 | 0.729 |
| 5. BMC - *Posthodiplostomum* sp. 11 | 2.226 | 1.895 | 2.223 | 2.054 | 1.649 | 2.161 | 2.120 | 2.231 | 1.998 | 2.242 | 2.191 | 2.184 | 0.747 |
| 6. BMC - Diplostomidae gen. sp. X | 2.208 | 1.853 | 2.127 | 1.977 | 1.726 | 2.267 | 2.234 | 2.218 | 1.963 | 2.316 | 2.188 | 2.265 | 1.691 |
| 7. BMC - *Posthodiplostomum* cf. *podicipitis* (2) | 2.206 | 0.225 | 2.168 | 2.047 | 1.765 | 2.130 | 2.083 | 2.212 | 1.974 | 2.136 | 2.234 | 2.176 | 1.801 |
| 8. BMC - *Posthodiplostomum ptychocheilus* | 2.090 | 1.758 | 2.051 | 2.023 | 0.328 | 2.140 | 2.143 | 2.095 | 1.912 | 2.219 | 2.214 | 2.092 | 1.561 |
| 9. BMC - *Posthodiplostomum* sp. 14 | 2.119 | 1.958 | 2.060 | 2.013 | 1.898 | 2.062 | 2.007 | 2.114 | 1.946 | 2.134 | 2.144 | 1.998 | 1.912 |
| 10. BMC - *Bolbophorus* sp. B (2) | 2.237 | 2.179 | 2.218 | 2.091 | 2.165 | 2.241 | 2.215 | 2.225 | 2.026 | 2.419 | 2.231 | 2.284 | 2.193 |
| 11. BMC - *Bolbophorus* sp. C | 2.168 | 2.222 | 2.219 | 2.251 | 2.181 | 2.289 | 2.256 | 2.185 | 2.206 | 2.494 | 2.358 | 2.352 | 2.152 |
| 12. *Bolbophorus damnificus* | 2.270 | 2.249 | 2.343 | 2.364 | 2.318 | 2.348 | 2.356 | 2.270 | 2.322 | 2.609 | 2.262 | 2.386 | 2.344 |
| 13. *Bolbophorus* sp. KM538081 | 2.235 | 2.257 | 2.287 | 2.233 | 2.213 | 2.309 | 2.271 | 2.244 | 2.207 | 2.539 | 2.400 | 2.356 | 2.231 |
| 14. *Bolbophorus* sp. KT831373 | 2.285 | 2.214 | 2.228 | 2.125 | 2.194 | 2.266 | 2.223 | 2.271 | 2.056 | 2.477 | 2.254 | 2.323 | 2.241 |
| 15. *Bolbophorus* sp. KU707938 | 2.318 | 2.186 | 2.226 | 2.187 | 2.179 | 2.223 | 2.182 | 2.293 | 2.149 | 2.429 | 2.141 | 2.298 | 2.160 |
| 16. *Bolbophorus* sp. MH368809 | 2.285 | 2.214 | 2.228 | 2.125 | 2.194 | 2.266 | 2.223 | 2.271 | 2.056 | 2.477 | 2.254 | 2.323 | 2.241 |
| 17. *Crocodillicola pseudostoma* (out) | 2.439 | 2.775 | 2.610 | 2.650 | 2.673 | 2.649 | 2.567 | 2.439 | 2.583 | 2.726 | 2.594 | 2.689 | 2.701 |
| 18. Diplostomidae gen. sp. O | 2.173 | 1.976 | 2.089 | 2.125 | 1.939 | 2.063 | 2.071 | 2.199 | 2.068 | 2.228 | 2.218 | 2.132 | 1.883 |
| 19. Diplostomidae gen. sp. X | 2.198 | 1.826 | 2.097 | 1.956 | 1.749 | 2.245 | 2.209 | 2.206 | 1.936 | 2.289 | 2.166 | 2.232 | 1.659 |
| 20. Diplostomoidea sp. | 2.199 | 2.340 | 2.323 | 2.213 | 2.287 | 2.207 | 2.162 | 2.216 | 2.200 | 2.528 | 2.311 | 2.403 | 2.357 |
| 21. *Neodiplostomum americanum* | 2.411 | 2.421 | 2.394 | 2.382 | 2.421 | 2.377 | 2.342 | 2.421 | 2.318 | 2.632 | 2.545 | 2.335 | 2.502 |
| 22. *Ornithodiplostomum scardinii* | 2.051 | 1.749 | 2.223 | 2.129 | 1.679 | 2.174 | 2.237 | 2.060 | 2.124 | 2.279 | 2.207 | 2.244 | 1.906 |
| 23. *Ornithodiplostomum* sp. 1 | 2.158 | 1.844 | 2.294 | 2.186 | 1.966 | 2.304 | 2.240 | 2.160 | 2.024 | 2.061 | 2.269 | 2.252 | 1.869 |
| 24. *Ornithodiplostomum* sp. 2 | 2.170 | 1.853 | 2.114 | 1.986 | 1.658 | 2.150 | 2.094 | 2.164 | 1.945 | 2.129 | 2.217 | 2.160 | 0.955 |
| 25. *Ornithodiplostomum* sp. 3 | 2.103 | 1.966 | 2.181 | 2.095 | 1.812 | 2.194 | 2.178 | 2.119 | 2.015 | 2.156 | 2.138 | 2.169 | 1.548 |

*Continued on next page*

Table B2 – *Continued from previous page*

| | 40 | 41 | 42 | 43 | 44 | 45 | 46 | 47 | 48 | 49 |
|---|---|---|---|---|---|---|---|---|---|---|
| 1. BMC - *Bolbophorus* sp. A | 2.325 | 2.430 | 2.331 | 2.448 | 2.504 | 2.453 | 2.443 | 2.423 | 2.356 | 2.554 |
| 2. BMC - *Posthodiplostomum* sp. 4 | 1.962 | 2.156 | 1.959 | 2.190 | 2.102 | 2.058 | 2.091 | 1.983 | 2.175 | 2.004 |
| 3. BMC - *Posthodiplostomum* sp. 11 | 1.937 | 1.891 | 1.874 | 1.700 | 2.129 | 2.034 | 1.927 | 1.997 | 2.089 | 2.073 |
| 4. BMC - *Posthodiplostomum* sp. 12 (2) | 1.869 | 1.810 | 1.707 | 1.677 | 2.055 | 1.921 | 1.858 | 1.888 | 1.949 | 1.969 |
| 5. BMC - *Posthodiplostomum* sp. 11 | 1.986 | 1.977 | 1.914 | 1.802 | 2.190 | 2.098 | 1.996 | 2.065 | 2.182 | 2.143 |
| 6. BMC - Diplostomidae gen. sp. X | 1.929 | 2.069 | 2.058 | 1.893 | 2.142 | 2.121 | 1.975 | 2.144 | 2.205 | 2.095 |
| 7. BMC - *Posthodiplostomum* cf. *podicipitis* (2) | 1.925 | 2.013 | 1.947 | 1.909 | 2.123 | 2.063 | 2.062 | 2.012 | 1.995 | 2.051 |
| 8. BMC - *Posthodiplostomum ptychocheilus* | 1.949 | 2.012 | 1.724 | 1.986 | 2.069 | 2.123 | 1.999 | 1.913 | 2.218 | 1.982 |
| 9. BMC - *Posthodiplostomum* sp. 14 | 1.939 | 1.863 | 1.713 | 2.033 | 1.989 | 1.873 | 1.931 | 1.937 | 2.090 | 1.968 |
| 10. BMC - *Bolbophorus* sp. B (2) | 2.111 | 2.171 | 2.204 | 2.289 | 2.186 | 2.155 | 2.249 | 2.174 | 2.150 | 2.269 |
| 11. BMC - *Bolbophorus* sp. C | 2.316 | 2.262 | 2.357 | 2.319 | 2.254 | 2.164 | 2.219 | 2.359 | 2.347 | 2.266 |
| 12. *Bolbophorus damnificus* | 2.264 | 2.346 | 2.378 | 2.361 | 2.320 | 2.287 | 2.327 | 2.382 | 2.321 | 2.346 |
| 13. *Bolbophorus* sp. KM538081 | 2.362 | 2.362 | 2.423 | 2.360 | 2.288 | 2.278 | 2.296 | 2.401 | 2.380 | 2.348 |
| 14. *Bolbophorus* sp. KT831373 | 2.129 | 2.215 | 2.237 | 2.304 | 2.236 | 2.180 | 2.283 | 2.204 | 2.159 | 2.307 |
| 15. *Bolbophorus* sp. KU707938 | 2.151 | 2.184 | 2.161 | 2.289 | 2.150 | 2.078 | 2.269 | 2.153 | 2.124 | 2.240 |
| 16. *Bolbophorus* sp. MH368809 | 2.129 | 2.215 | 2.237 | 2.304 | 2.236 | 2.180 | 2.283 | 2.204 | 2.159 | 2.307 |
| 17. *Crocodillicola pseudostoma* (out) | 2.689 | 2.685 | 2.760 | 2.688 | 2.681 | 2.650 | 2.543 | 2.563 | 2.567 | 2.611 |
| 18. Diplostomidae gen. sp. O | 2.168 | 1.960 | 1.941 | 1.927 | 2.105 | 1.921 | 2.282 | 2.219 | 2.152 | 2.102 |
| 19. Diplostomidae gen. sp. X | 1.921 | 2.052 | 2.031 | 1.867 | 2.114 | 2.086 | 1.964 | 2.109 | 2.172 | 2.072 |
| 20. Diplostomoidea sp. | 2.344 | 2.328 | 2.354 | 2.507 | 2.264 | 2.196 | 2.446 | 2.219 | 2.460 | 2.297 |
| 21. *Neodiplostomum americanum* | 2.331 | 2.350 | 2.486 | 2.456 | 2.399 | 2.347 | 2.465 | 2.519 | 2.357 | 2.377 |
| 22. *Ornithodiplostomum scardinii* | 2.216 | 2.004 | 2.023 | 1.707 | 2.004 | 2.141 | 2.104 | 2.023 | 2.127 | 1.923 |
| 23. *Ornithodiplostomum* sp. 1 | 1.994 | 2.082 | 2.067 | 2.035 | 2.185 | 2.081 | 2.173 | 2.111 | 2.105 | 2.180 |
| 24. *Ornithodiplostomum* sp. 2 | 1.986 | 1.895 | 1.885 | 1.681 | 2.100 | 2.003 | 2.055 | 2.035 | 2.131 | 2.070 |
| 25. *Ornithodiplostomum* sp. 3 | 2.023 | 1.893 | 1.789 | 1.804 | 2.102 | 1.990 | 2.086 | 2.140 | 2.126 | 2.116 |

*Continued on next page*

Table B2 – *Continued from previous page*

| | 1 | 2 | 3 | 4 | 5 | 6 | 7 | 8 | 9 | 10 | 11 | 12 |
|---|---|---|---|---|---|---|---|---|---|---|---|---|
| 26. *Ornithodiplostomum* sp. 8 | 76.568 | 86.047 | 88.372 | 87.41 | 87.459 | 85.809 | 87.748 | 90.429 | 99.34 | 81.229 | 79.866 | 79.47 |
| 27. *Posthodiplostomum centrarchi* | 76.238 | 84.053 | 83.721 | 83.052 | 82.508 | 81.848 | 82.119 | 84.158 | 82.178 | 81.395 | 81.208 | 78.477 |
| 28. *Posthodiplostomum* cf. *podicipitis* | 77.815 | 84.385 | 88.333 | 88.052 | 87.417 | 88.079 | 99.67 | 89.073 | 87.086 | 82.333 | 81.481 | 81.728 |
| 29. *Posthodiplostomum cuticola* | 76.08 | 84.281 | 83.165 | 82.314 | 81.94 | 83.278 | 83.221 | 83.946 | 83.278 | 81.879 | 81 | 79.667 |
| 30. *Posthodiplostomum minimum* | 78.548 | 95.364 | 86.667 | 85.004 | 85.43 | 85.43 | 85.382 | 85.762 | 85.762 | 83.833 | 80.936 | 80.198 |
| 31. *Posthodiplostomum ptychocheilus* | 75.578 | 85.382 | 92.359 | 90.761 | 91.089 | 90.429 | 88.742 | 99.67 | 89.439 | 80.731 | 81.879 | 80.132 |
| 32. *Posthodiplostomum* sp. 1 | 76 | 83.221 | 84.564 | 83.218 | 83.333 | 82.667 | 83.946 | 83.667 | 84.667 | 81.376 | 81.017 | 77.926 |
| 33. *Posthodiplostomum* sp. 2 | 76.568 | 84.053 | 85.382 | 84.059 | 84.158 | 83.498 | 84.437 | 83.828 | 85.149 | 82.226 | 82.215 | 78.808 |
| 34. *Posthodiplostomum* sp. 3 | 75.908 | 84.385 | 84.053 | 83.217 | 82.838 | 82.178 | 82.45 | 84.488 | 82.508 | 81.063 | 81.544 | 78.808 |
| 35. *Posthodiplostomum* sp. 4 (2) | 78.053 | 97.185 | 86.333 | 84.753 | 85.099 | 84.934 | 84.718 | 85.43 | 85.265 | 83.5 | 80.769 | 80.033 |
| 36. *Posthodiplostomum* sp. 5 | 72.517 | 85.05 | 81.94 | 81.253 | 81.395 | 81.395 | 82.667 | 81.728 | 82.392 | 78.261 | 78.523 | 75.497 |
| 37. *Posthodiplostomum* sp. 7 | 75.578 | 81.728 | 84.718 | 83.558 | 83.828 | 83.498 | 82.119 | 83.498 | 83.168 | 81.063 | 79.53 | 80.464 |
| 38. *Posthodiplostomum* sp. 8 | 74.503 | 83.444 | 81.271 | 81.425 | 80.399 | 80.066 | 82.45 | 81.395 | 83.721 | 80.268 | 75.503 | 79.139 |
| 39. *Posthodiplostomum* sp. 11 | 76.568 | 85.714 | 99.336 | 96.246 | 98.35 | 90.759 | 87.748 | 91.749 | 87.129 | 81.229 | 81.544 | 79.801 |
| 40. *Posthodiplostomum* sp. 17 | 78.878 | 85.714 | 85.714 | 84.554 | 84.818 | 85.149 | 84.768 | 85.809 | 85.809 | 84.385 | 80.201 | 80.132 |
| 41. *Posthodiplostomum* sp. 18 | 76.238 | 83.721 | 88.04 | 87.239 | 87.129 | 85.149 | 86.093 | 86.469 | 86.139 | 81.063 | 80.537 | 79.801 |
| 42. *Posthodiplostomum* sp. 19 | 76.49 | 85.382 | 88.333 | 87.881 | 87.417 | 85.762 | 86.469 | 89.735 | 90.066 | 81.333 | 79.125 | 79.734 |
| 43. *Posthodiplostomum* sp. 20 | 76.159 | 83.721 | 90.333 | 88.543 | 89.073 | 88.079 | 88.119 | 87.086 | 85.43 | 80.167 | 79.798 | 79.734 |
| 44. *Posthodiplostomum* sp. 21 | 78.218 | 85.382 | 83.721 | 82.545 | 82.838 | 83.168 | 83.775 | 83.828 | 84.158 | 84.219 | 81.879 | 82.119 |
| 45. *Posthodiplostomum* sp. 22 | 78.218 | 86.047 | 86.047 | 85.407 | 85.149 | 85.479 | 86.093 | 85.149 | 86.139 | 83.555 | 82.886 | 81.457 |
| 46. *Posthodiplostomum* sp. 23 | 74.257 | 83.775 | 87 | 86.54 | 85.762 | 86.424 | 85.05 | 85.43 | 85.43 | 80.167 | 80.936 | 80.198 |
| 47. *Posthodiplostomum* sp. 24 | 77.558 | 86.379 | 85.05 | 83.888 | 83.828 | 83.498 | 84.437 | 86.799 | 86.469 | 82.558 | 79.53 | 79.47 |
| 48. *Posthodiplostomum* sp. 25 | 77.558 | 85.099 | 86 | 85.192 | 84.768 | 84.768 | 86.379 | 83.113 | 83.775 | 83 | 79.599 | 80.528 |
| 49. *Posthodiplostomum* sp. 26 | 77.558 | 86.379 | 85.714 | 84.901 | 84.488 | 86.139 | 84.768 | 85.809 | 84.818 | 83.056 | 80.872 | 80.132 |

*Continued on next page*

Table B2 – *Continued from previous page*

| | 13 | 14 | 15 | 16 | 17 | 18 | 19 | 20 | 21 | 22 | 23 | 24 | 25 | 26 |
|---|---|---|---|---|---|---|---|---|---|---|---|---|---|---|
| 26. *Ornithodiplostomum* sp. 8 | 79.47 | 81.395 | 82.06 | 81.395 | 73.667 | 87.417 | 86.139 | 80.066 | 77.483 | 86.799 | 86.379 | 87.417 | 87.459 | - |
| 27. *Posthodiplostomum centrarchi* | 81.126 | 81.728 | 80.066 | 81.728 | 74 | 82.119 | 82.178 | 80.066 | 78.808 | 83.498 | 84.053 | 83.113 | 82.838 | 82.178 |
| 28. *Posthodiplostomum* cf. *podicipitis* | 81.063 | 82.667 | 83 | 82.667 | 70.234 | 86.379 | 88.411 | 79.734 | 78.738 | 89.404 | 87 | 88.04 | 87.086 | 87.748 |
| 29. *Posthodiplostomum cuticola* | 81 | 82.215 | 81.879 | 82.215 | 69.333 | 82 | 83.612 | 80.537 | 79 | 81.94 | 82 | 83 | 82.274 | 83.946 |
| 30. *Posthodiplostomum minimum* | 81.848 | 84 | 82 | 84 | 72 | 85.149 | 85.762 | 80.399 | 77.888 | 83.775 | 84.768 | 85.479 | 85.099 | 86.424 |
| 31. *Posthodiplostomum ptychocheilus* | 81.126 | 80.731 | 81.395 | 80.731 | 71 | 88.742 | 90.099 | 81.728 | 76.821 | 91.089 | 87.375 | 90.397 | 89.439 | 90.099 |
| 32. *Posthodiplostomum* sp. 1 | 80.602 | 81.879 | 82.55 | 81.879 | 71.717 | 82.943 | 83 | 79.195 | 78.595 | 80.333 | 80.872 | 83.612 | 82 | 85.333 |
| 33. *Posthodiplostomum* sp. 2 | 81.788 | 82.724 | 83.389 | 82.724 | 71.667 | 83.775 | 83.828 | 80.066 | 79.801 | 80.528 | 82.06 | 84.437 | 83.168 | 85.809 |
| 34. *Posthodiplostomum* sp. 3 | 81.457 | 81.395 | 79.734 | 81.395 | 74 | 82.45 | 82.508 | 80.399 | 79.139 | 83.828 | 84.385 | 83.444 | 83.168 | 82.508 |
| 35. *Posthodiplostomum* sp. 4 (2) | 81.188 | 83.667 | 81.833 | 83.667 | 71 | 84.158 | 85.265 | 80.066 | 77.888 | 83.113 | 84.768 | 85.149 | 84.768 | 85.927 |
| 36. *Posthodiplostomum* sp. 5 | 78.146 | 78.595 | 78.595 | 78.595 | 72.241 | 81.788 | 81.728 | 75.333 | 73.841 | 80.731 | 84.385 | 82.781 | 81.395 | 82.392 |
| 37. *Posthodiplostomum* sp. 7 | 79.47 | 81.063 | 82.392 | 81.063 | 69.667 | 81.457 | 83.828 | 79.402 | 76.159 | 82.838 | 79.402 | 82.781 | 83.168 | 83.168 |
| 38. *Posthodiplostomum* sp. 8 | 75.497 | 80.602 | 80.602 | 80.602 | 71.237 | 80.464 | 80.399 | 76.744 | 76.49 | 80.399 | 78.738 | 80.795 | 82.724 | 84.385 |
| 39. *Posthodiplostomum* sp. 11 | 80.795 | 81.063 | 82.06 | 81.063 | 70.333 | 88.742 | 91.089 | 79.734 | 74.834 | 87.789 | 86.047 | 97.351 | 92.409 | 87.789 |
| 40. *Posthodiplostomum* sp. 17 | 80.464 | 84.385 | 83.721 | 84.385 | 70.333 | 83.444 | 85.479 | 79.07 | 79.139 | 82.178 | 85.05 | 85.099 | 84.158 | 86.469 |
| 41. *Posthodiplostomum* sp. 18 | 79.47 | 81.063 | 81.395 | 81.063 | 71 | 87.748 | 85.479 | 79.402 | 78.477 | 85.809 | 85.05 | 87.417 | 87.459 | 86.799 |
| 42. *Posthodiplostomum* sp. 19 | 78.405 | 81.333 | 83 | 81.333 | 71.906 | 88.04 | 86.093 | 79.07 | 77.409 | 86.424 | 84.667 | 87.043 | 88.411 | 90.728 |
| 43. *Posthodiplostomum* sp. 20 | 79.07 | 81 | 81 | 81 | 70.903 | 87.043 | 88.411 | 78.405 | 78.073 | 89.404 | 84.667 | 90.033 | 88.742 | 85.43 |
| 44. *Posthodiplostomum* sp. 21 | 81.788 | 84.385 | 85.382 | 84.385 | 71.333 | 83.444 | 83.498 | 81.728 | 80.132 | 84.818 | 83.721 | 83.775 | 83.168 | 84.818 |
| 45. *Posthodiplostomum* sp. 22 | 82.45 | 84.053 | 84.385 | 84.053 | 71.667 | 87.417 | 85.809 | 81.728 | 80.464 | 84.818 | 84.053 | 86.093 | 86.139 | 86.799 |
| 46. *Posthodiplostomum* sp. 23 | 80.198 | 80.333 | 81.667 | 80.333 | 73.333 | 82.838 | 86.755 | 78.073 | 77.228 | 84.106 | 82.781 | 85.479 | 86.424 | 85.43 |
| 47. *Posthodiplostomum* sp. 24 | 79.801 | 82.724 | 83.056 | 82.724 | 74 | 82.781 | 83.828 | 81.728 | 77.483 | 85.149 | 83.721 | 83.775 | 82.838 | 86.469 |
| 48. *Posthodiplostomum* sp. 25 | 80.198 | 83.333 | 83 | 83.333 | 73.667 | 83.498 | 85.099 | 77.741 | 78.548 | 84.437 | 81.788 | 85.149 | 85.099 | 83.113 |
| 49. *Posthodiplostomum* sp. 26 | 80.795 | 83.389 | 83.389 | 83.389 | 73.333 | 84.437 | 86.469 | 80.731 | 77.815 | 86.139 | 84.053 | 85.099 | 84.488 | 85.479 |

*Continued on next page*

Table B2 – *Continued from previous page*

| | 27 | 28 | 29 | 30 | 31 | 32 | 33 | 34 | 35 | 36 | 37 | 38 | 39 |
|---|---|---|---|---|---|---|---|---|---|---|---|---|---|
| 26. *Ornithodiplostomum* sp. 8 | 2.116 | 1.910 | 2.013 | 1.979 | 1.831 | 2.030 | 1.973 | 2.112 | 1.905 | 2.128 | 2.147 | 1.964 | 1.887 |
| 27. *Posthodiplostomum centrarchi* | - | 2.212 | 2.238 | 2.041 | 2.113 | 2.127 | 2.098 | 0.324 | 1.963 | 2.143 | 2.106 | 2.172 | 2.171 |
| 28. *Posthodiplostomum* cf. *podicipitis* | 82.119 | - | 2.163 | 2.070 | 1.778 | 2.118 | 2.069 | 2.219 | 1.994 | 2.136 | 2.246 | 2.174 | 1.831 |
| 29. *Posthodiplostomum cuticola* | 80.268 | 83.557 | - | 2.039 | 2.047 | 1.976 | 1.953 | 2.254 | 2.002 | 2.285 | 2.249 | 2.084 | 2.188 |
| 30. *Posthodiplostomum minimum* | 84.768 | 85.382 | 85.333 | - | 2.032 | 1.982 | 1.934 | 2.055 | 0.660 | 2.076 | 2.291 | 2.123 | 2.002 |
| 31. *Posthodiplostomum ptychocheilus* | 83.828 | 88.742 | 83.612 | 85.43 | - | 2.129 | 2.118 | 2.115 | 1.914 | 2.231 | 2.233 | 2.087 | 1.596 |
| 32. *Posthodiplostomum* sp. 1 | 83 | 84.281 | 84.797 | 83.612 | 83.333 | - | 0.956 | 2.147 | 1.943 | 2.003 | 2.175 | 1.969 | 2.145 |
| 33. *Posthodiplostomum* sp. 2 | 83.498 | 84.768 | 86.288 | 84.768 | 83.498 | 97 | - | 2.121 | 1.903 | 2.091 | 2.257 | 1.975 | 2.121 |
| 34. *Posthodiplostomum* sp. 3 | 99.67 | 82.45 | 80.602 | 85.099 | 84.158 | 83.333 | 83.828 | - | 1.972 | 2.143 | 2.093 | 2.169 | 2.165 |
| 35. *Posthodiplostomum* sp. 4 (2) | 83.94 | 84.718 | 84.833 | 97.195 | 85.099 | 83.278 | 84.437 | 84.272 | - | 2.031 | 2.211 | 1.968 | 1.934 |
| 36. *Posthodiplostomum* sp. 5 | 83.056 | 82.667 | 79.264 | 84.437 | 81.395 | 83.893 | 82.06 | 83.056 | 84.603 | - | 2.239 | 2.103 | 2.209 |
| 37. *Posthodiplostomum* sp. 7 | 82.178 | 82.119 | 80.936 | 81.126 | 83.168 | 82.667 | 81.848 | 82.508 | 81.291 | 82.06 | - | 2.162 | 2.147 |
| 38. *Posthodiplostomum* sp. 8 | 82.392 | 82.781 | 81.94 | 82.781 | 81.063 | 84.899 | 84.385 | 82.06 | 83.609 | 81.395 | 80.731 | - | 2.140 |
| 39. *Posthodiplostomum* sp. 11 | 83.168 | 87.748 | 82.274 | 85.762 | 91.419 | 83.333 | 84.158 | 83.498 | 85.43 | 81.063 | 83.498 | 81.063 | - |
| 40. *Posthodiplostomum* sp. 17 | 81.518 | 84.768 | 80.602 | 84.768 | 86.139 | 82.333 | 83.498 | 81.188 | 85.099 | 82.392 | 82.178 | 81.728 | 85.149 |
| 41. *Posthodiplostomum* sp. 18 | 84.818 | 86.093 | 81.94 | 85.43 | 86.139 | 83.333 | 84.488 | 85.149 | 84.272 | 81.395 | 82.178 | 81.063 | 87.789 |
| 42. *Posthodiplostomum* sp. 19 | 82.45 | 86.469 | 82.55 | 86.711 | 89.404 | 83.278 | 85.099 | 82.119 | 86.047 | 80.667 | 82.781 | 84.437 | 87.748 |
| 43. *Posthodiplostomum* sp. 20 | 82.119 | 88.119 | 82.215 | 83.056 | 86.755 | 82.274 | 83.444 | 82.45 | 83.223 | 81 | 83.444 | 81.126 | 89.073 |
| 44. *Posthodiplostomum* sp. 21 | 83.498 | 84.106 | 82.609 | 84.437 | 83.498 | 84 | 85.149 | 83.828 | 84.934 | 83.721 | 86.139 | 84.053 | 82.838 |
| 45. *Posthodiplostomum* sp. 22 | 84.158 | 86.093 | 83.278 | 85.762 | 84.818 | 85.667 | 87.129 | 84.488 | 85.762 | 84.718 | 85.479 | 83.389 | 85.149 |
| 46. *Posthodiplostomum* sp. 23 | 82.119 | 85.05 | 82 | 83.498 | 85.099 | 85.284 | 87.417 | 82.45 | 83.333 | 81.457 | 83.113 | 80.795 | 86.093 |
| 47. *Posthodiplostomum* sp. 24 | 83.828 | 84.437 | 80.936 | 86.755 | 86.469 | 84 | 83.828 | 84.158 | 86.424 | 84.385 | 85.149 | 82.392 | 84.158 |
| 48. *Posthodiplostomum* sp. 25 | 81.457 | 86.379 | 82 | 83.828 | 82.781 | 81.271 | 81.788 | 81.788 | 84.653 | 83.113 | 86.755 | 83.113 | 84.768 |
| 49. *Posthodiplostomum* sp. 26 | 82.178 | 84.768 | 82.609 | 87.417 | 85.479 | 83.333 | 83.168 | 82.508 | 86.921 | 83.389 | 83.828 | 83.721 | 84.818 |

*Continued on next page*

Table B2 – *Continued from previous page*

| | 40 | 41 | 42 | 43 | 44 | 45 | 46 | 47 | 48 | 49 |
|---|---|---|---|---|---|---|---|---|---|---|
| 26. *Ornithodiplostomum* sp. 8 | 1.921 | 1.856 | 1.658 | 2.024 | 1.968 | 1.862 | 1.945 | 1.948 | 2.132 | 1.946 |
| 27. *Posthodiplostomum centrarchi* | 2.191 | 2.119 | 2.211 | 2.108 | 2.095 | 2.108 | 2.096 | 2.130 | 2.157 | 2.217 |
| 28. *Posthodiplostomum* cf. *podicipitis* | 1.933 | 2.028 | 1.963 | 1.922 | 2.104 | 2.080 | 2.075 | 2.024 | 2.009 | 2.079 |
| 29. *Posthodiplostomum cuticola* | 2.272 | 2.162 | 2.070 | 2.280 | 2.240 | 2.165 | 2.202 | 2.170 | 2.218 | 2.170 |
| 30. *Posthodiplostomum minimum* | 2.018 | 2.058 | 1.971 | 2.221 | 2.128 | 2.018 | 2.123 | 1.914 | 2.148 | 1.924 |
| 31. *Posthodiplostomum ptychocheilus* | 1.954 | 2.040 | 1.717 | 2.034 | 2.086 | 2.157 | 2.010 | 1.914 | 2.220 | 1.999 |
| 32. *Posthodiplostomum* sp. 1 | 2.177 | 2.026 | 2.006 | 2.186 | 2.030 | 1.880 | 2.008 | 2.025 | 2.256 | 2.018 |
| 33. *Posthodiplostomum* sp. 2 | 2.142 | 2.012 | 1.976 | 2.177 | 2.033 | 1.846 | 1.913 | 2.034 | 2.264 | 2.059 |
| 34. *Posthodiplostomum* sp. 3 | 2.187 | 2.119 | 2.197 | 2.126 | 2.097 | 2.112 | 2.122 | 2.138 | 2.148 | 2.232 |
| 35. *Posthodiplostomum* sp. 4 (2) | 1.895 | 2.027 | 1.887 | 2.130 | 2.055 | 1.955 | 2.056 | 1.848 | 2.048 | 1.897 |
| 36. *Posthodiplostomum* sp. 5 | 2.131 | 2.289 | 2.212 | 2.213 | 2.056 | 2.109 | 2.152 | 2.130 | 2.207 | 2.196 |
| 37. *Posthodiplostomum* sp. 7 | 2.100 | 2.193 | 2.170 | 2.153 | 1.950 | 2.015 | 2.225 | 2.007 | 2.016 | 2.153 |
| 38. *Posthodiplostomum* sp. 8 | 2.036 | 2.121 | 1.926 | 2.220 | 2.047 | 2.028 | 2.219 | 2.023 | 2.195 | 2.083 |
| 39. *Posthodiplostomum* sp. 11 | 1.957 | 1.893 | 1.869 | 1.744 | 2.148 | 2.050 | 1.981 | 2.004 | 2.136 | 2.115 |
| 40. *Posthodiplostomum* sp. 17 | - | 2.112 | 1.902 | 2.202 | 2.139 | 2.049 | 2.134 | 2.104 | 2.183 | 2.014 |
| 41. *Posthodiplostomum* sp. 18 | 83.168 | - | 1.858 | 1.892 | 2.079 | 1.760 | 2.176 | 2.184 | 2.066 | 2.026 |
| 42. *Posthodiplostomum* sp. 19 | 86.755 | 88.079 | - | 1.941 | 2.093 | 1.964 | 2.032 | 2.118 | 2.219 | 2.109 |
| 43. *Posthodiplostomum* sp. 20 | 83.113 | 87.748 | 87.129 | - | 2.057 | 2.125 | 2.079 | 2.231 | 2.008 | 2.130 |
| 44. *Posthodiplostomum* sp. 21 | 84.818 | 83.828 | 83.113 | 84.106 | - | 1.794 | 2.138 | 1.917 | 1.863 | 1.732 |
| 45. *Posthodiplostomum* sp. 22 | 84.818 | 88.119 | 86.424 | 85.099 | 89.439 | - | 2.073 | 1.918 | 2.052 | 1.857 |
| 46. *Posthodiplostomum* sp. 23 | 83.113 | 84.768 | 85.382 | 86.711 | 83.113 | 85.099 | - | 1.994 | 2.163 | 2.168 |
| 47. *Posthodiplostomum* sp. 24 | 82.838 | 82.508 | 84.768 | 82.781 | 87.789 | 86.799 | 83.444 | - | 1.994 | 1.793 |
| 48. *Posthodiplostomum* sp. 25 | 81.126 | 84.106 | 82.392 | 86.711 | 88.411 | 87.086 | 84.158 | 86.755 | - | 1.985 |
| 49. *Posthodiplostomum* sp. 26 | 85.479 | 83.828 | 83.444 | 83.444 | 90.759 | 88.779 | 82.119 | 89.439 | 87.748 | - |

Table B3. Averaged percent similarities between *Echinoparyphium, Hypoderaeum* and *Drepanocephalus* species (*COI* gene). Standard error estimate(s) are shown above the diagonal and were obtained by a bootstrap procedure (500 replicates). Ambiguous positions were removed for each sequence pair. Evolutionary analyses were conducted in MEGA (Tamura et al., 2021).

| | 1 | 2 | 3 | 4 | 5 | 6 | 7 | 8 | 9 | 10 | 11 | 12 |
|---|---|---|---|---|---|---|---|---|---|---|---|---|
| 1. BMC - *Echinoparyphium* sp. A (2) | - | 1.514 | 1.660 | 1.431 | 1.490 | 1.699 | 1.714 | 1.756 | 1.476 | 1.852 | 1.852 | 1.914 |
| 2. BMC - *Echinoparyphium* sp. Lin 1B (3) | 85.553 | - | 1.761 | 1.668 | 1.588 | 1.796 | 1.616 | 1.711 | 1.694 | 1.905 | 1.905 | 1.987 |
| 3. BMC - *Echinoparyphium* sp. C | 80.943 | 80.464 | - | 1.753 | 1.623 | 1.788 | 1.610 | 1.704 | 1.626 | 1.887 | 1.887 | 1.838 |
| 4. BMC - *Echinoparyphium* sp. Lin 2 (2) | 87.193 | 86.612 | 81.352 | - | 1.496 | 1.539 | 1.789 | 1.765 | 1.518 | 1.894 | 1.894 | 1.943 |
| 5. BMC - *Echinoparyphium* sp. Lin 3 | 88.217 | 84.836 | 81.352 | 88.115 | - | 1.672 | 1.725 | 1.741 | 1.522 | 1.921 | 1.921 | 1.915 |
| 6. BMC - *Echinoparyphium* sp. Lin 5 | 82.889 | 82.309 | 81.557 | 85.656 | 83.402 | - | 1.728 | 1.817 | 1.625 | 1.891 | 1.891 | 1.886 |
| 7. BMC - *Echinoparyphium* sp. E | 82.992 | 82.787 | 84.631 | 81.557 | 83.607 | 80.533 | - | 1.663 | 1.705 | 1.926 | 1.926 | 1.958 |
| 8. BMC - *Hypoderaeum conoideum* | 81.557 | 82.377 | 80.328 | 81.352 | 81.762 | 79.918 | 82.582 | - | 1.692 | 1.832 | 1.832 | 1.936 |
| 9. BMC - *Echinoparyphium* sp. Lin 6 | 86.783 | 82.855 | 82.172 | 86.885 | 87.09 | 84.426 | 82.992 | 83.197 | - | 1.940 | 1.940 | 1.936 |
| 10. *Drepanocephalus mexicanus* | 75.462 | 73.922 | 75.359 | 75.154 | 74.538 | 74.127 | 74.538 | 72.951 | 72.485 | - | 0.285 | 1.489 |
| 11. *Drepanocephalus* sp. | 75.462 | 73.922 | 75.359 | 75.154 | 74.538 | 74.127 | 74.538 | 72.951 | 72.485 | 99.59 | - | 1.508 |
| 12. *Drepanocephalus spathans* | 75.873 | 72.553 | 75.359 | 74.743 | 74.949 | 75.154 | 74.538 | 73.156 | 73.717 | 87.09 | 87.295 | - |
| 13. *Echinoparyphium aconiatum* | 85.451 | 83.88 | 82.787 | 83.607 | 83.607 | 82.172 | 83.607 | 81.148 | 82.172 | 75.154 | 75.154 | 76.797 |
| 14. *Echinoparyphium* sp. A2 | 80.635 | 80.874 | 83.607 | 78.484 | 81.148 | 79.508 | 82.377 | 79.303 | 81.967 | 75.77 | 75.77 | 76.591 |
| 15. *Echinoparyphium* sp. A | 99.59 | 85.451 | 80.943 | 87.411 | 88.32 | 82.992 | 82.992 | 81.352 | 86.885 | 75.565 | 75.565 | 75.77 |
| 16. *Echinoparyphium* sp. C | 80.533 | 80.464 | 99.59 | 80.943 | 81.352 | 81.148 | 84.221 | 79.918 | 81.762 | 75.359 | 75.359 | 74.949 |
| 17. *Echinoparyphium* sp. E | 82.992 | 82.104 | 84.631 | 81.557 | 83.811 | 80.533 | 99.18 | 82.172 | 82.787 | 74.743 | 74.743 | 74.743 |
| 18. *Echinoparyphium* sp. Lin 3 | 86.783 | 84.221 | 80.738 | 88.525 | 96.311 | 82.992 | 83.402 | 81.762 | 86.27 | 75.359 | 75.359 | 75.565 |
| 19. *Echinoparyphium* sp. Lin 1A | 85.963 | 89.208 | 79.303 | 88.73 | 86.885 | 83.607 | 81.762 | 80.943 | 86.066 | 75.154 | 75.154 | 75.154 |
| 20. *Echinoparyphium* sp. Lin 2 | 87.602 | 86.612 | 81.148 | 99.59 | 88.32 | 85.656 | 81.352 | 81.557 | 86.885 | 74.949 | 74.949 | 74.538 |
| 21. *Euparyphium capitaneum* (out) (2) | 78.337 | 76.42 | 75.359 | 76.694 | 77.926 | 77.413 | 77.002 | 73.463 | 76.489 | 77.459 | 77.869 | 78.689 |
| 22. *Hypoderaeum conoideum* | 81.557 | 82.377 | 80.328 | 81.352 | 81.762 | 79.918 | 82.582 | 100 | 83.197 | 72.895 | 72.895 | 73.101 |
| 23. *Hypoderaeum* sp. | 87.193 | 86.202 | 81.148 | 99.59 | 88.32 | 85.656 | 81.352 | 81.557 | 86.475 | 75.359 | 75.359 | 74.949 |

*Continued on next page*

Table B3 – *Continued from previous page*

| | 13 | 14 | 15 | 16 | 17 | 18 | 19 | 20 | 21 | 22 | 23 |
|---|---|---|---|---|---|---|---|---|---|---|---|
| 1. BMC - *Echinoparyphium* sp. A (2) | 1.589 | 1.767 | 0.199 | 1.660 | 1.711 | 1.501 | 1.535 | 1.429 | 1.775 | 1.756 | 1.452 |
| 2. BMC - *Echinoparyphium* sp. Lin 1B (3) | 1.717 | 1.652 | 1.534 | 1.781 | 1.624 | 1.622 | 1.360 | 1.648 | 1.874 | 1.711 | 1.690 |
| 3. BMC - *Echinoparyphium* sp. C | 1.619 | 1.628 | 1.662 | 0.286 | 1.603 | 1.627 | 1.768 | 1.754 | 1.828 | 1.704 | 1.777 |
| 4. BMC - *Echinoparyphium* sp. Lin 2 (2) | 1.683 | 1.852 | 1.440 | 1.762 | 1.760 | 1.464 | 1.429 | 0.255 | 1.853 | 1.765 | 0.255 |
| 5. BMC - *Echinoparyphium* sp. Lin 3 | 1.715 | 1.688 | 1.503 | 1.616 | 1.702 | 0.804 | 1.531 | 1.488 | 1.836 | 1.741 | 1.495 |
| 6. BMC - *Echinoparyphium* sp. Lin 5 | 1.770 | 1.850 | 1.719 | 1.778 | 1.718 | 1.668 | 1.657 | 1.556 | 1.795 | 1.817 | 1.535 |
| 7. BMC - *Echinoparyphium* sp. E | 1.648 | 1.682 | 1.713 | 1.628 | 0.394 | 1.646 | 1.786 | 1.789 | 1.883 | 1.663 | 1.799 |
| 8. BMC - *Hypoderaeum conoideum* | 1.801 | 1.812 | 1.770 | 1.692 | 1.689 | 1.726 | 1.754 | 1.786 | 1.817 | 0.000 | 1.767 |
| 9. BMC - *Echinoparyphium* sp. Lin 6 | 1.764 | 1.669 | 1.490 | 1.625 | 1.690 | 1.487 | 1.503 | 1.516 | 1.753 | 1.692 | 1.552 |
| 10. *Drepanocephalus mexicanus* | 1.909 | 1.900 | 1.849 | 1.901 | 1.899 | 1.859 | 1.910 | 1.899 | 1.820 | 1.833 | 1.900 |
| 11. *Drepanocephalus* sp. | 1.909 | 1.900 | 1.849 | 1.901 | 1.899 | 1.859 | 1.910 | 1.899 | 1.820 | 1.833 | 1.900 |
| 12. *Drepanocephalus spathans* | 1.895 | 1.895 | 1.915 | 1.818 | 1.934 | 1.847 | 1.901 | 1.947 | 1.881 | 1.935 | 1.950 |
| 13. *Echinoparyphium aconiatum* | - | 1.811 | 1.607 | 1.617 | 1.664 | 1.763 | 1.782 | 1.668 | 1.775 | 1.801 | 1.704 |
| 14. *Echinoparyphium* sp. A2 | 81.557 | - | 1.789 | 1.626 | 1.652 | 1.638 | 1.721 | 1.850 | 1.885 | 1.812 | 1.864 |
| 15. *Echinoparyphium* sp. A | 85.451 | 80.328 | - | 1.664 | 1.705 | 1.515 | 1.551 | 1.438 | 1.794 | 1.770 | 1.458 |
| 16. *Echinoparyphium* sp. C | 82.377 | 83.402 | 80.533 | - | 1.624 | 1.618 | 1.757 | 1.763 | 1.828 | 1.692 | 1.785 |
| 17. *Echinoparyphium* sp. E | 83.607 | 82.582 | 82.992 | 84.221 | - | 1.618 | 1.748 | 1.759 | 1.909 | 1.689 | 1.770 |
| 18. *Echinoparyphium* sp. Lin 3 | 82.172 | 81.352 | 86.885 | 80.738 | 83.607 | - | 1.469 | 1.460 | 1.831 | 1.726 | 1.460 |
| 19. *Echinoparyphium* sp. Lin 1A | 81.762 | 80.943 | 86.066 | 79.303 | 82.172 | 87.09 | - | 1.451 | 1.871 | 1.754 | 1.438 |
| 20. *Echinoparyphium* sp. Lin 2 | 83.607 | 78.484 | 87.91 | 80.738 | 81.352 | 88.73 | 88.525 | - | 1.857 | 1.786 | 0.290 |
| 21. *Euparyphium capitaneum* (out) (2) | 79.877 | 77.002 | 78.439 | 75.154 | 77.002 | 77.515 | 76.283 | 76.899 | - | 1.817 | 1.861 |
| 22. *Hypoderaeum conoideum* | 81.148 | 79.303 | 81.352 | 79.918 | 82.172 | 81.762 | 80.943 | 81.557 | 73.409 | - | 1.767 |
| 23. *Hypoderaeum* sp. | 83.197 | 78.484 | 87.5 | 80.738 | 81.352 | 88.73 | 88.525 | 99.59 | 76.489 | 81.557 | - |

Table B4. Averaged percent similarities between *Echinoparyphium, Hypoderaeum* and *Drepanocephalus* species (*nad1* gene). Standard error estimate(s) are shown above the diagonal and were obtained by a bootstrap procedure (500 replicates). Ambiguous positions were removed for each sequence pair. Evolutionary analyses were conducted in MEGA (Tamura et al., 2021).

| | 1 | 2 | 3 | 4 | 5 | 6 | 7 | 8 | 9 | 10 | 11 |
|---|---|---|---|---|---|---|---|---|---|---|---|
| 1. BMC - *Drepanocephalus spathans* (2) | - | 2.5685 | 2.4202 | 2.4984 | 2.5570 | 2.4845 | 2.5906 | 2.4610 | 1.8439 | 1.8248 | 0.0000 |
| 2. BMC - *Echinoparyphium* sp. Lin 2 (2) | 71.0145 | - | 2.0594 | 1.8853 | 0.2696 | 2.2762 | 2.0789 | 2.0606 | 2.5969 | 2.5778 | 2.5685 |
| 3. BMC - *Echinoparyphium* sp. Lin 4 | 70.1449 | 82.7089 | - | 1.9813 | 2.0541 | 2.0347 | 2.2070 | 0.5997 | 2.4625 | 2.4681 | 2.4202 |
| 4. BMC - *Echinoparyphium* sp. Lin 5 | 71.8023 | 84.9711 | 82.0809 | - | 1.8853 | 2.0569 | 2.0383 | 1.9622 | 2.5091 | 2.5084 | 2.4984 |
| 5. BMC - *Echinoparyphium rubrum* | 71.3043 | 99.7118 | 82.9971 | 84.9711 | - | 2.2625 | 2.0658 | 2.0529 | 2.5804 | 2.5611 | 2.5570 |
| 6. BMC - *Echinoparyphium* sp. Lin 6 | 69.7674 | 78.3237 | 80.9249 | 81.5029 | 78.6127 | - | 2.1997 | 2.0135 | 2.5162 | 2.5303 | 2.4845 |
| 7. BMC - *Echinoparyphium* sp. Lin 1B (2) | 69.4767 | 82.6590 | 79.7688 | 83.8150 | 82.9480 | 81.2680 | - | 2.1648 | 2.5050 | 2.5673 | 2.5906 |
| 8. BMC - *Echinoparyphium* sp. A (2) | 68.8406 | 82.4207 | 98.4150 | 81.6474 | 82.7089 | 80.4913 | 79.7688 | - | 2.4724 | 2.4825 | 2.4610 |
| 9. *Drepanocephalus mexicanus* | 85.8790 | 66.6667 | 67.2464 | 68.6047 | 66.9565 | 66.5698 | 66.8605 | 66.5217 | - | 0.6572 | 1.8439 |
| 10. *Drepanocephalus* sp. | 86.4553 | 67.2464 | 67.5362 | 68.6047 | 67.5362 | 66.8605 | 66.5698 | 66.8116 | 98.5591 | - | 1.8248 |
| 11. *Drepanocephalus spathans* | 100 | 71.0145 | 70.1449 | 71.8023 | 71.3043 | 69.7674 | 69.4767 | 68.8406 | 85.8790 | 86.4553 | - |
| 12. *Echinoparyphium aconiatum* | 70.6395 | 78.9017 | 78.6127 | 80.0578 | 79.1908 | 75.7225 | 76.0116 | 78.9017 | 69.1860 | 69.1860 | 70.6395 |
| 13. *Echinoparyphium ellisi* | 68.6957 | 84.7262 | 81.2680 | 84.3931 | 85.0144 | 83.2370 | 82.3699 | 81.1239 | 65.5072 | 65.5072 | 68.6957 |
| 14. *Echinoparyphium poulini* | 69.8551 | 85.5908 | 81.8444 | 84.6821 | 85.5908 | 78.6127 | 81.7919 | 81.7003 | 62.6087 | 63.7681 | 69.8551 |
| 15. *Echinoparyphium recurvatum* | 68.9855 | 82.4207 | 76.3689 | 82.3699 | 82.1326 | 78.3237 | 78.9017 | 76.5130 | 64.0580 | 64.9275 | 68.9855 |
| 16. *Echinoparyphium rubrum* | 71.3043 | 99.7118 | 82.9971 | 84.9711 | 100.0000 | 78.6127 | 82.9480 | 82.7089 | 66.9565 | 67.5362 | 71.3043 |
| 17. *Echinoparyphium* sp. 1 | 71.0145 | 80.4035 | 78.0980 | 76.8786 | 80.6916 | 74.8555 | 76.0116 | 78.0980 | 67.2464 | 67.5362 | 71.0145 |
| 18. *Echinoparyphium* sp. 2 | 68.9855 | 82.4207 | 98.8473 | 81.7919 | 82.7089 | 81.2139 | 79.4798 | 98.9914 | 66.6667 | 66.9565 | 68.9855 |
| 19. *Echinoparyphium* sp. LC599763 | 70.1449 | 84.4380 | 79.5389 | 81.7919 | 84.7262 | 78.0347 | 79.4798 | 79.9712 | 66.0870 | 67.2464 | 70.1449 |
| 20. *Echinoparyphium* sp. (3) | 69.4767 | 81.8882 | 78.9017 | 82.6590 | 81.8882 | 77.5530 | 83.0443 | 78.6127 | 67.2481 | 67.2481 | 69.4767 |
| 21. *Echinoparyphium* sp. PQ421432 | 68.9855 | 84.4380 | 83.5735 | 85.5491 | 84.7262 | 80.9249 | 90.4624 | 83.4294 | 67.2464 | 67.2464 | 68.9855 |
| 22. *Echinoparyphium* sp. A2 | 70.6395 | 81.7919 | 83.2370 | 81.2139 | 82.0809 | 81.2139 | 80.9249 | 83.2370 | 67.7326 | 67.7326 | 70.6395 |
| 23. *Echinoparyphium* sp. A | 69.5652 | 82.9971 | 97.1182 | 84.1040 | 83.2853 | 82.0809 | 80.3468 | 97.2622 | 67.2464 | 67.5362 | 69.5652 |

*Continued on next page*

Table B4 – *Continued from previous page*

| | 12 | 13 | 14 | 15 | 16 | 17 | 18 | 19 | 20 | 21 | 22 | 23 |
|---|---|---|---|---|---|---|---|---|---|---|---|---|
| 1. BMC - *Drepanocephalus spathans* (2) | 2.4283 | 2.5576 | 2.4515 | 2.5130 | 2.5570 | 2.4413 | 2.4448 | 2.4290 | 2.3910 | 2.6180 | 2.3852 | 2.4879 |
| 2. BMC - *Echinoparyphium* sp. Lin 2 (2) | 2.1305 | 2.0545 | 1.8775 | 2.0534 | 0.2696 | 2.1251 | 2.0802 | 1.9229 | 2.0314 | 1.9592 | 2.0069 | 2.0467 |
| 3. BMC - *Echinoparyphium* sp. Lin 4 | 2.0953 | 2.1282 | 2.0482 | 2.2623 | 2.0541 | 2.1760 | 0.5711 | 2.2170 | 2.1472 | 1.9881 | 2.0107 | 0.9165 |
| 4. BMC - *Echinoparyphium* sp. Lin 5 | 2.1253 | 1.9843 | 1.9622 | 2.1802 | 1.8853 | 2.1607 | 1.9860 | 2.0898 | 2.0562 | 1.9271 | 2.0326 | 1.8948 |
| 5. BMC - *Echinoparyphium rubrum* | 2.1192 | 2.0366 | 1.8775 | 2.0689 | 0.0000 | 2.1149 | 2.0690 | 1.9130 | 2.0314 | 1.9471 | 2.0046 | 2.0427 |
| 6. BMC - *Echinoparyphium* sp. Lin 6 | 2.2645 | 2.0163 | 2.1145 | 2.1877 | 2.2625 | 2.3340 | 2.0304 | 2.2296 | 2.2946 | 2.1220 | 1.9960 | 1.9731 |
| 7. BMC - *Echinoparyphium* sp. Lin 1B (2) | 2.3543 | 2.1664 | 2.0864 | 2.2142 | 2.0658 | 2.2954 | 2.2070 | 2.1476 | 1.8944 | 1.4803 | 2.0589 | 2.1591 |
| 8. BMC - *Echinoparyphium* sp. A (2) | 2.0282 | 2.0976 | 2.0195 | 2.2303 | 2.0529 | 2.1931 | 0.4288 | 2.1397 | 2.1477 | 1.9590 | 1.9739 | 0.8246 |
| 9. *Drepanocephalus mexicanus* | 2.4982 | 2.5847 | 2.6147 | 2.5558 | 2.5804 | 2.5519 | 2.4757 | 2.5522 | 2.4174 | 2.5279 | 2.4851 | 2.4975 |
| 10. *Drepanocephalus* sp. | 2.4909 | 2.6008 | 2.5905 | 2.5348 | 2.5611 | 2.5363 | 2.4824 | 2.5276 | 2.4589 | 2.5663 | 2.4655 | 2.4999 |
| 11. *Drepanocephalus spathans* | 2.4283 | 2.5576 | 2.4515 | 2.5130 | 2.5570 | 2.4413 | 2.4448 | 2.4290 | 2.3910 | 2.6180 | 2.3852 | 2.4879 |
| 12. *Echinoparyphium aconiatum* | - | 2.2072 | 2.1294 | 2.2826 | 2.1192 | 2.1569 | 2.0686 | 2.1609 | 2.2649 | 2.3587 | 2.1547 | 2.0303 |
| 13. *Echinoparyphium ellisi* | 77.7457 | - | 2.0781 | 2.1822 | 2.0366 | 2.2006 | 2.1273 | 2.0527 | 2.1369 | 2.0571 | 2.1080 | 2.0252 |
| 14. *Echinoparyphium poulini* | 77.4566 | 82.1326 | - | 2.1092 | 1.8775 | 2.1639 | 2.0467 | 2.0411 | 2.0978 | 2.0577 | 1.9874 | 2.0111 |
| 15. *Echinoparyphium recurvatum* | 75.1445 | 80.1153 | 79.8271 | - | 2.0689 | 2.3773 | 2.2932 | 1.6482 | 2.3589 | 2.2204 | 2.1444 | 2.2453 |
| 16. *Echinoparyphium rubrum* | 79.1908 | 85.0144 | 85.5908 | 82.1326 | - | 2.1149 | 2.0690 | 1.9130 | 2.0314 | 1.9471 | 2.0046 | 2.0427 |
| 17. *Echinoparyphium* sp. 1 | 80.6358 | 76.3689 | 77.5216 | 74.0634 | 80.6916 | - | 2.2022 | 2.3055 | 2.2219 | 2.2957 | 2.1889 | 2.1905 |
| 18. *Echinoparyphium* sp. 2 | 78.6127 | 80.9798 | 81.5562 | 76.0807 | 82.7089 | 78.0980 | - | 2.2018 | 2.1197 | 1.9917 | 2.0222 | 0.8147 |
| 19. *Echinoparyphium* sp. LC599763 | 77.1676 | 81.2680 | 82.1326 | 89.3372 | 84.7262 | 75.7925 | 79.8271 | - | 2.1844 | 2.2054 | 2.1189 | 2.1213 |
| 20. *Echinoparyphium* sp. (3) | 75.5299 | 81.3102 | 80.7322 | 76.0116 | 81.8882 | 76.9750 | 79.1908 | 79.2871 | - | 2.0425 | 2.2204 | 2.0936 |
| 21. *Echinoparyphium* sp. PQ421432 | 76.8786 | 83.8617 | 82.9971 | 78.6744 | 84.7262 | 74.6398 | 83.2853 | 78.6744 | 80.9249 | - | 1.9661 | 2.0799 |
| 22. *Echinoparyphium* sp. A2 | 78.0347 | 81.5029 | 82.0809 | 79.1908 | 82.0809 | 79.7688 | 82.9480 | 80.0578 | 78.9981 | 82.9480 | - | 1.9927 |
| 23. *Echinoparyphium* sp. A | 80.3468 | 82.9971 | 82.4207 | 77.8098 | 83.2853 | 78.9625 | 97.6945 | 80.1153 | 80.6358 | 82.4207 | 83.5260 | - |

*Continued on next page*

Table B4 – *Continued from previous page*

| | 24 | 25 | 26 | 27 | 28 | 29 | 30 | 31 | 32 | 33 | 34 | 35 |
|---|---|---|---|---|---|---|---|---|---|---|---|---|
| 1. BMC - *Drepanocephalus spathans* (2) | 2.3873 | 2.1548 | 2.3440 | 2.5865 | 2.6109 | 2.5906 | 2.5869 | 2.2785 | 2.4547 | 2.6137 | 2.5431 | 2.4393 |
| 2. BMC - *Echinoparyphium* sp. Lin 2 (2) | 2.4386 | 2.4560 | 2.3445 | 2.0992 | 2.0398 | 2.0789 | 1.0424 | 1.8451 | 2.0289 | 2.3273 | 2.2613 | 2.5828 |
| 3. BMC - *Echinoparyphium* sp. Lin 4 | 2.3611 | 2.2545 | 2.2836 | 2.2195 | 2.0915 | 2.2070 | 2.1079 | 1.7304 | 0.5778 | 2.2398 | 2.1690 | 2.5388 |
| 4. BMC - *Echinoparyphium* sp. Lin 5 | 2.3547 | 2.4158 | 2.3217 | 2.1157 | 2.0888 | 2.0383 | 1.9149 | 1.8953 | 1.9890 | 2.2143 | 2.2010 | 2.6095 |
| 5. BMC - *Echinoparyphium rubrum* | 2.4363 | 2.4495 | 2.3150 | 2.0760 | 2.0227 | 2.0658 | 1.0701 | 1.8336 | 2.0180 | 2.3273 | 2.2613 | 2.5632 |
| 6. BMC - *Echinoparyphium* sp. Lin 6 | 2.3450 | 2.3787 | 2.2299 | 2.1291 | 2.0830 | 2.1997 | 2.2684 | 1.8068 | 2.0547 | 2.1726 | 2.2025 | 2.5907 |
| 7. BMC - *Echinoparyphium* sp. Lin 1B (2) | 2.3555 | 2.3729 | 2.3690 | 1.6973 | 1.7299 | 0.0000 | 2.0829 | 2.0078 | 2.1952 | 2.1724 | 2.1743 | 2.5704 |
| 8. BMC - *Echinoparyphium* sp. A (2) | 2.3064 | 2.3106 | 2.2336 | 2.1661 | 2.0652 | 2.1648 | 2.0998 | 1.7128 | 0.4290 | 2.2600 | 2.2059 | 2.5140 |
| 9. *Drepanocephalus mexicanus* | 2.5269 | 2.4698 | 2.3988 | 2.5327 | 2.5260 | 2.5050 | 2.6204 | 2.3662 | 2.4687 | 2.6376 | 2.5487 | 2.5267 |
| 10. *Drepanocephalus* sp. | 2.4906 | 2.4582 | 2.3582 | 2.5815 | 2.5784 | 2.5673 | 2.6043 | 2.3473 | 2.4735 | 2.5721 | 2.4775 | 2.4362 |
| 11. *Drepanocephalus spathans* | 2.3873 | 2.1548 | 2.3440 | 2.5865 | 2.6109 | 2.5906 | 2.5869 | 2.2785 | 2.4547 | 2.6137 | 2.5431 | 2.4393 |
| 12. *Echinoparyphium aconiatum* | 2.1808 | 2.4137 | 2.2053 | 2.3480 | 2.3692 | 2.3543 | 2.1101 | 2.0004 | 2.0324 | 2.1643 | 2.2635 | 2.6171 |
| 13. *Echinoparyphium ellisi* | 2.3422 | 2.3157 | 2.3668 | 1.8792 | 2.0190 | 2.1664 | 2.0139 | 1.9061 | 2.1286 | 2.4129 | 2.3852 | 2.4481 |
| 14. *Echinoparyphium poulini* | 2.3994 | 2.4409 | 2.3474 | 2.0919 | 2.1415 | 2.0864 | 1.8865 | 1.8456 | 2.0264 | 2.3617 | 2.2798 | 2.6486 |
| 15. *Echinoparyphium recurvatum* | 2.4187 | 2.3345 | 2.4560 | 2.2037 | 2.3893 | 2.2142 | 2.0515 | 2.0433 | 2.2534 | 2.4143 | 2.2506 | 2.4892 |
| 16. *Echinoparyphium rubrum* | 2.4363 | 2.4495 | 2.3150 | 2.0760 | 2.0227 | 2.0658 | 1.0701 | 1.8336 | 2.0180 | 2.3273 | 2.2613 | 2.5632 |
| 17. *Echinoparyphium* sp. 1 | 2.1810 | 2.3242 | 2.1978 | 2.2318 | 2.2593 | 2.2954 | 2.1847 | 1.9265 | 2.2141 | 2.1422 | 2.2134 | 2.5040 |
| 18. *Echinoparyphium* sp. 2 | 2.3273 | 2.3114 | 2.2456 | 2.2086 | 2.0807 | 2.2070 | 2.1146 | 1.7447 | 0.4222 | 2.2649 | 2.2012 | 2.5560 |
| 19. *Echinoparyphium* sp. LC599763 | 2.3591 | 2.2942 | 2.3515 | 2.1326 | 2.2802 | 2.1476 | 1.9384 | 1.9371 | 2.1456 | 2.5633 | 2.3844 | 2.4988 |
| 20. *Echinoparyphium* sp. (3) | 2.4087 | 2.4328 | 2.2682 | 2.0404 | 2.1456 | 1.8944 | 2.0617 | 2.0651 | 2.1403 | 2.2545 | 2.2652 | 2.5623 |
| 21. *Echinoparyphium* sp. PQ421432 | 2.4626 | 2.3714 | 2.2361 | 1.5160 | 1.3808 | 1.4803 | 2.0294 | 1.8906 | 1.9806 | 2.1727 | 2.1938 | 2.5967 |
| 22. *Echinoparyphium* sp. A2 | 2.3529 | 2.3711 | 2.3265 | 2.0335 | 2.1066 | 2.0589 | 2.0219 | 1.1814 | 2.0161 | 2.1743 | 2.2582 | 2.5216 |
| 23. *Echinoparyphium* sp. A | 2.2715 | 2.3677 | 2.2688 | 2.1186 | 2.1200 | 2.1591 | 2.0707 | 1.7171 | 0.8233 | 2.2000 | 2.1620 | 2.5175 |

*Continued on next page*

Table B4 – *Continued from previous page*

| | 1 | 2 | 3 | 4 | 5 | 6 | 7 | 8 | 9 | 10 | 11 |
|---|---|---|---|---|---|---|---|---|---|---|---|
| 24. *Echinoparyphium* sp. C | 70.0581 | 74.8555 | 76.0116 | 76.0116 | 75.1445 | 75.1445 | 74.5665 | 75.8671 | 66.8605 | 67.4419 | 70.0581 |
| 25. *Echinoparyphium* sp. D | 75.8721 | 75.1445 | 76.8786 | 73.6994 | 75.4335 | 71.3873 | 73.1214 | 75.7225 | 70.3488 | 70.6395 | 75.8721 |
| 26. *Echinoparyphium* sp. E | 75.5814 | 77.4566 | 76.3006 | 76.5896 | 77.7457 | 75.4335 | 76.0116 | 75.5780 | 72.6744 | 72.9651 | 75.5814 |
| 27. *Echinoparyphium* sp. Lin 1 | 68.4058 | 82.7089 | 80.9798 | 83.2370 | 82.9971 | 81.2139 | 87.5723 | 81.1239 | 66.6667 | 66.0870 | 68.4058 |
| 28. *Echinoparyphium* sp. Lin 1A | 69.8551 | 82.4207 | 80.9798 | 81.7919 | 82.7089 | 78.9017 | 86.4162 | 80.8357 | 67.5362 | 67.5362 | 69.8551 |
| 29. *Echinoparyphium* sp. Lin 1B | 69.4767 | 82.6590 | 79.7688 | 83.8150 | 82.9480 | 81.2680 | 100.0000 | 79.7688 | 66.8605 | 66.5698 | 69.4767 |
| 30. *Echinoparyphium* sp. Lin 2 | 70.1449 | 95.9654 | 80.9798 | 85.2601 | 95.6772 | 78.9017 | 82.6590 | 80.6916 | 67.2464 | 67.8261 | 70.1449 |
| 31. *Echinoparyphium* sp. Lin 3 (2) | 70.2490 | 81.0989 | 83.8379 | 80.7803 | 81.3875 | 80.6358 | 79.4798 | 83.7659 | 67.9247 | 68.6502 | 70.2490 |
| 32. *Echinoparyphium* sp. Lin 4 | 69.2754 | 82.9971 | 98.8473 | 81.7919 | 83.2853 | 80.6358 | 79.7688 | 98.9914 | 67.2464 | 67.5362 | 69.2754 |
| 33. *Hypoderaeum conoideum* | 70.3488 | 78.6127 | 78.3237 | 78.6127 | 78.6127 | 78.3862 | 79.2507 | 77.8902 | 70.3488 | 71.5116 | 70.3488 |
| 34. *Hypoderaeum* sp. Lin 1 (2) | 69.7674 | 77.4566 | 77.8902 | 78.4682 | 77.4566 | 77.3450 | 78.6444 | 77.1676 | 69.4767 | 70.6395 | 69.7674 |
| 35. *Isthmiophora melis* (out) | 73.2558 | 68.4971 | 68.7861 | 68.4971 | 68.7861 | 66.7630 | 68.7861 | 68.4971 | 69.1860 | 70.3488 | 73.2558 |

| | 12 | 13 | 14 | 15 | 16 | 17 | 18 | 19 | 20 | 21 | 22 | 23 |
|---|---|---|---|---|---|---|---|---|---|---|---|---|
| 24. *Echinoparyphium* sp. C | 79.7688 | 75.4335 | 74.5665 | 73.9884 | 75.1445 | 77.1676 | 76.0116 | 76.0116 | 73.5067 | 72.2543 | 74.8555 | 77.7457 |
| 25. *Echinoparyphium* sp. D | 75.4335 | 74.5665 | 73.9884 | 73.6994 | 75.4335 | 75.7225 | 75.7225 | 75.1445 | 74.1811 | 73.9884 | 75.7225 | 75.7225 |
| 26. *Echinoparyphium* sp. E | 79.1908 | 74.2775 | 74.8555 | 73.1214 | 77.7457 | 79.1908 | 75.7225 | 73.9884 | 75.8886 | 76.3006 | 76.5896 | 75.7225 |
| 27. *Echinoparyphium* sp. Lin 1 | 78.0347 | 86.1671 | 82.1326 | 79.5389 | 82.9971 | 75.7925 | 80.6916 | 79.2507 | 81.5029 | 92.2190 | 81.7919 | 81.8444 |
| 28. *Echinoparyphium* sp. Lin 1A | 73.6994 | 83.5735 | 80.4035 | 76.3689 | 82.7089 | 73.4870 | 80.6916 | 76.9452 | 78.6127 | 92.7954 | 80.3468 | 80.6916 |
| 29. *Echinoparyphium* sp. Lin 1B | 76.0116 | 82.3699 | 81.7919 | 78.9017 | 82.9480 | 76.0116 | 79.4798 | 79.4798 | 83.0443 | 90.4624 | 80.9249 | 80.3468 |
| 30. *Echinoparyphium* sp. Lin 2 | 79.4798 | 85.3026 | 85.3026 | 82.4207 | 95.6772 | 79.2507 | 80.6916 | 84.4380 | 81.3102 | 84.4380 | 82.6590 | 81.8444 |
| 31. *Echinoparyphium* sp. Lin 3 (2) | 78.0347 | 81.2426 | 80.9548 | 78.3570 | 81.3875 | 79.6547 | 83.5493 | 80.3760 | 77.8902 | 80.6654 | 91.3295 | 84.1265 |
| 32. *Echinoparyphium* sp. Lin 4 | 79.1908 | 80.9798 | 81.8444 | 76.6571 | 83.2853 | 78.0980 | 99.4236 | 80.4035 | 78.9017 | 83.5735 | 82.9480 | 97.6945 |
| 33. *Hypoderaeum conoideum* | 77.7457 | 77.1676 | 75.1445 | 74.2775 | 78.6127 | 79.4798 | 78.3237 | 73.9884 | 78.4200 | 78.9017 | 77.7457 | 79.1908 |
| 34. *Hypoderaeum* sp. Lin 1 (2) | 76.9115 | 77.0231 | 74.8555 | 76.0116 | 77.4566 | 77.4566 | 77.3121 | 73.8439 | 76.3969 | 77.7457 | 75.4335 | 79.0462 |
| 35. *Isthmiophora melis* (out) | 67.7233 | 68.7861 | 68.7861 | 69.3642 | 68.7861 | 70.2312 | 68.2081 | 69.3642 | 67.8227 | 67.9191 | 68.2081 | 69.3642 |

*Continued on next page*

Table B4 – *Continued from previous page*

| | 24 | 25 | 26 | 27 | 28 | 29 | 30 | 31 | 32 | 33 | 34 | 35 |
|---|---|---|---|---|---|---|---|---|---|---|---|---|
| 24. *Echinoparyphium* sp. C | - | 2.1955 | 2.1995 | 2.4075 | 2.3852 | 2.3555 | 2.4922 | 2.1294 | 2.3344 | 2.4241 | 2.3903 | 2.3991 |
| 25. *Echinoparyphium* sp. D | 78.3237 | - | 2.2871 | 2.4392 | 2.4323 | 2.3729 | 2.4970 | 2.1772 | 2.3279 | 2.4524 | 2.3783 | 2.5675 |
| 26. *Echinoparyphium* sp. E | 79.7688 | 78.6127 | - | 2.2811 | 2.2105 | 2.3690 | 2.2971 | 2.0265 | 2.2422 | 2.3394 | 2.3801 | 2.6352 |
| 27. *Echinoparyphium* sp. Lin 1 | 74.2775 | 73.6994 | 76.8786 | - | 1.6571 | 1.6973 | 2.0572 | 1.9058 | 2.1963 | 2.2704 | 2.2604 | 2.4414 |
| 28. *Echinoparyphium* sp. Lin 1A | 71.9653 | 72.2543 | 76.5896 | 89.9135 | - | 1.7299 | 2.0868 | 1.9635 | 2.0779 | 2.3016 | 2.2451 | 2.4968 |
| 29. *Echinoparyphium* sp. Lin 1B | 74.5665 | 73.1214 | 76.0116 | 87.5723 | 86.4162 | - | 2.0829 | 2.0078 | 2.1952 | 2.1724 | 2.1743 | 2.5704 |
| 30. *Echinoparyphium* sp. Lin 2 | 73.4104 | 75.1445 | 76.8786 | 84.1499 | 82.4207 | 82.6590 | - | 1.8409 | 2.0738 | 2.3638 | 2.3260 | 2.6291 |
| 31. *Echinoparyphium* sp. Lin 3 (2) | 75.0000 | 75.0346 | 76.1561 | 80.6642 | 79.0771 | 79.4798 | 81.9647 | - | 1.7544 | 2.1149 | 2.1749 | 2.3725 |
| 32. *Echinoparyphium* sp. Lin 4 | 76.0116 | 75.7225 | 76.0116 | 80.9798 | 80.9798 | 79.7688 | 81.2680 | 83.5493 | - | 2.2566 | 2.2026 | 2.5382 |
| 33. *Hypoderaeum conoideum* | 75.4335 | 73.4104 | 76.0116 | 77.7457 | 77.4566 | 79.2507 | 78.9017 | 77.3121 | 78.6127 | - | 1.4744 | 2.5155 |
| 34. *Hypoderaeum* sp. Lin 1 (2) | 74.8555 | 73.4104 | 73.6994 | 77.1676 | 76.0116 | 78.6444 | 77.3121 | 75.8671 | 77.6012 | 91.1979 | - | 2.4366 |
| 35. *Isthmiophora melis* (out) | 69.9422 | 69.6532 | 66.7630 | 70.5202 | 68.4971 | 68.7861 | 67.6301 | 68.2081 | 68.4971 | 69.3642 | 68.9756 | - |

Table B5. Averaged percent similarities between *Echinostoma* species (*COI* gene). Standard error estimate(s) are shown above the diagonal and were obtained by a bootstrap procedure (500 replicates). Ambiguous positions were removed for each sequence pair. Evolutionary analyses were conducted in MEGA (Tamura et al., 2021).

| | 1 | 2 | 3 | 4 | 5 | 6 | 7 | 8 | 9 | 10 | 11 | 12 | 13 | 14 |
|---|---|---|---|---|---|---|---|---|---|---|---|---|---|---|
| 1. BMC - *Echinostoma trivolvis* complex Lin A (2) | - | 1.504 | 1.679 | 1.539 | 1.546 | 1.582 | 1.543 | 1.670 | 1.506 | 1.320 | 1.539 | 1.320 | 0.334 | 1.930 |
| 2. BMC - *Echinostoma revolutum* complex sp. B (2) | 87.227 | - | 1.721 | 1.466 | 1.586 | 1.698 | 0.312 | 1.770 | 0.302 | 1.599 | 1.439 | 1.599 | 1.522 | 1.991 |
| 3. *Echinostoma* aff. *trivolvis* | 80.786 | 80.240 | - | 1.851 | 1.705 | 1.832 | 1.711 | 1.814 | 1.749 | 1.634 | 1.802 | 1.633 | 1.699 | 1.931 |
| 4. *Echinostoma bolschewense* | 87.991 | 87.664 | 80.349 | - | 1.648 | 1.622 | 1.464 | 1.701 | 1.492 | 1.582 | 1.519 | 1.580 | 1.551 | 1.990 |
| 5. *Echinostoma paraensei* | 87.227 | 87.227 | 80.568 | 85.371 | - | 1.722 | 1.597 | 1.837 | 1.593 | 1.567 | 1.507 | 1.568 | 1.580 | 1.930 |
| 6. *Echinostoma revolutum* | 85.371 | 83.515 | 80.786 | 85.371 | 83.843 | - | 1.717 | 1.715 | 1.699 | 1.619 | 1.702 | 1.619 | 1.613 | 1.984 |
| 7. *Echinostoma revolutum* complex sp. B | 87.227 | 99.236 | 80.349 | 87.991 | 87.555 | 83.624 | - | 1.787 | 0.418 | 1.611 | 1.473 | 1.611 | 1.571 | 2.009 |
| 8. *Echinostoma* sp. 1 | 82.642 | 81.769 | 81.441 | 82.533 | 81.878 | 82.751 | 82.096 | - | 1.781 | 1.684 | 1.742 | 1.683 | 1.680 | 1.972 |
| 9. *Echinostoma* sp. 2 | 87.227 | 99.236 | 79.913 | 87.555 | 87.118 | 83.624 | 99.127 | 81.659 | - | 1.615 | 1.477 | 1.615 | 1.532 | 1.990 |
| 10. *Echinostoma* sp. FJ477201 | 90.939 | 85.917 | 81.659 | 86.026 | 87.336 | 84.498 | 86.026 | 83.624 | 86.026 | - | 1.563 | 0.213 | 1.320 | 1.917 |
| 11. *Echinostoma* sp. MZ407836 | 86.681 | 88.428 | 79.913 | 87.991 | 87.555 | 84.061 | 88.428 | 81.004 | 88.210 | 86.026 | - | 1.563 | 1.553 | 1.940 |
| 12. *Echinostoma trivolvis* | 90.939 | 85.917 | 81.878 | 86.245 | 87.555 | 84.498 | 86.026 | 83.843 | 86.026 | 99.782 | 86.026 | - | 1.320 | 1.917 |
| 13. *Echinostoma trivolvis* complex sp. Lineage A | 99.017 | 87.227 | 80.786 | 87.991 | 87.336 | 85.153 | 87.118 | 82.751 | 87.118 | 91.266 | 86.900 | 91.266 | - | 1.935 |
| 14 *Euparyphium capitaneum* (out) | 76.310 | 76.638 | 76.856 | 75.546 | 77.074 | 74.891 | 76.419 | 76.638 | 76.856 | 76.856 | 78.384 | 76.856 | 76.638 | - |

Table B6. Averaged percent similarities between *Echinostoma* species (*nad1* gene). Standard error estimate(s) are shown above the diagonal and were obtained by a bootstrap procedure (500 replicates). Ambiguous positions were removed for each sequence pair. Evolutionary analyses were conducted in MEGA (Tamura et al., 2021).

| | 1 | 2 | 3 | 4 | 5 | 6 | 7 | 8 | 9 | 10 | 11 | 12 | 13 |
|---|---|---|---|---|---|---|---|---|---|---|---|---|---|
| 1. BMC - *Echinostoma revolutum* complex sp. B (2) | - | 1.626 | 1.725 | 1.680 | 1.930 | 1.716 | 1.545 | 2.310 | 1.938 | 1.696 | 1.721 | 1.515 | 1.618 |
| 2. BMC - *Echinostoma trivolvis* complex Lin A (2) | 86.374 | - | 1.757 | 1.663 | 1.855 | 1.790 | 1.736 | 2.168 | 1.922 | 1.679 | 1.735 | 1.707 | 1.516 |
| 3. *Echinostoma bolschewense* | 84.360 | 84.597 | - | 1.830 | 1.926 | 1.548 | 1.795 | 2.300 | 1.969 | 1.797 | 1.843 | 1.767 | 1.807 |
| 4. *Echinostoma caproni* | 85.664 | 86.019 | 81.754 | - | 1.859 | 1.868 | 1.723 | 2.268 | 2.008 | 1.652 | 1.757 | 1.788 | 1.748 |
| 5. *Echinostoma* cf. *friedi* | 80.569 | 81.043 | 80.806 | 80.332 | - | 1.830 | 2.007 | 2.188 | 2.065 | 1.979 | 1.977 | 1.949 | 1.890 |
| 6. *Echinostoma chankense* | 85.427 | 82.701 | 88.626 | 80.806 | 81.043 | - | 1.782 | 2.293 | 1.998 | 1.851 | 1.830 | 1.746 | 1.786 |
| 7. *Echinostoma cinetorchis* | 88.507 | 86.730 | 82.464 | 86.019 | 79.621 | 83.649 | - | 2.278 | 1.931 | 1.708 | 1.466 | 1.523 | 1.692 |
| 8. *Echinostoma hortense* | 66.627 | 66.508 | 66.033 | 65.321 | 65.558 | 65.796 | 66.033 | - | 2.220 | 2.275 | 2.282 | 2.339 | 2.263 |
| 9. *Echinostoma macrorchis* | 75.653 | 76.722 | 76.485 | 76.485 | 75.059 | 76.960 | 75.772 | 67.299 | - | 2.040 | 1.998 | 1.993 | 1.927 |
| 10. *Echinostoma maldonadoi* | 84.716 | 86.967 | 82.464 | 85.071 | 79.384 | 84.123 | 85.782 | 66.983 | 76.247 | - | 1.775 | 1.781 | 1.712 |
| 11. *Echinostoma mekongi* | 86.137 | 86.256 | 81.754 | 84.123 | 78.910 | 82.464 | 89.573 | 65.796 | 76.247 | 85.071 | - | 1.637 | 1.879 |
| 12. *Echinostoma miyagawai* | 88.863 | 85.782 | 83.175 | 85.308 | 81.043 | 84.597 | 89.573 | 65.083 | 76.247 | 83.412 | 87.204 | - | 1.727 |
| 13. *Echinostoma nasincovae* | 87.085 | 88.863 | 83.649 | 84.834 | 80.332 | 84.360 | 85.545 | 65.321 | 76.960 | 86.967 | 82.701 | 85.545 | - |
| 14. *Echinostoma novazealandense* (2) | 89.336 | 87.678 | 85.545 | 87.441 | 80.569 | 84.834 | 88.863 | 66.983 | 75.297 | 85.071 | 86.967 | 90.047 | 87.441 |
| 15. *Echinostoma paraensei* | 83.768 | 86.256 | 82.464 | 85.308 | 79.621 | 84.360 | 86.493 | 66.508 | 75.059 | 93.602 | 83.412 | 83.412 | 86.967 |
| 16. *Echinostoma paraulum* | 87.085 | 85.782 | 83.649 | 85.545 | 80.332 | 84.360 | 90.284 | 66.508 | 77.435 | 86.493 | 88.626 | 88.626 | 84.123 |
| 17. *Echinostoma pseudorobustum* | 86.848 | 87.204 | 83.175 | 84.597 | 80.569 | 84.597 | 90.521 | 66.508 | 76.247 | 85.308 | 87.441 | 91.706 | 86.967 |
| 18. *Echinostoma revolutum* (3) | 91.351 | 87.125 | 84.123 | 84.518 | 79.700 | 84.439 | 88.626 | 66.508 | 76.168 | 84.834 | 88.863 | 88.073 | 86.256 |
| 19. *Echinostoma revolutum* complex sp. B | 98.934 | 86.256 | 84.360 | 86.256 | 80.569 | 84.597 | 88.626 | 66.746 | 76.010 | 84.834 | 86.256 | 88.389 | 87.204 |

*Continued on next page*

Table B6 – *Continued from previous page*

| | 14 | 15 | 16 | 17 | 18 | 19 | 20 | 21 | 22 | 23 | 24 | 25 | 26 | 27 | 28 | 29 |
|---|---|---|---|---|---|---|---|---|---|---|---|---|---|---|---|---|
| 1. BMC - *Echinostoma revolutum* complex sp. B (2) | 1.507 | 1.727 | 1.625 | 1.605 | 1.045 | 0.464 | 1.490 | 1.999 | 1.928 | 0.325 | 1.694 | 1.921 | 1.574 | 1.568 | 1.612 | 2.200 |
| 2. BMC - *Echinostoma trivolvis* complex Lin A (2) | 1.636 | 1.778 | 1.731 | 1.651 | 1.371 | 1.642 | 1.695 | 2.082 | 1.924 | 1.665 | 1.822 | 1.869 | 1.520 | 1.254 | 0.171 | 2.213 |
| 3. *Echinostoma bolschewense* | 1.673 | 1.835 | 1.814 | 1.839 | 1.503 | 1.749 | 1.767 | 1.981 | 1.938 | 1.749 | 1.700 | 1.933 | 1.804 | 1.769 | 1.752 | 2.182 |
| 4. *Echinostoma caproni* | 1.677 | 1.713 | 1.774 | 1.798 | 1.533 | 1.644 | 1.789 | 2.132 | 1.985 | 1.669 | 1.776 | 1.871 | 1.758 | 1.711 | 1.664 | 2.247 |
| 5. *Echinostoma* cf. *friedi* | 1.947 | 1.930 | 1.920 | 1.921 | 1.775 | 1.941 | 1.961 | 2.044 | 2.068 | 1.942 | 2.045 | 0.386 | 1.887 | 1.915 | 1.845 | 2.215 |
| 6. *Echinostoma chankense* | 1.725 | 1.741 | 1.736 | 1.769 | 1.532 | 1.790 | 1.743 | 2.010 | 2.042 | 1.721 | 1.788 | 1.799 | 1.776 | 1.786 | 1.782 | 2.192 |
| 7. *Echinostoma cinetorchis* | 1.612 | 1.679 | 1.492 | 1.407 | 1.273 | 1.520 | 1.515 | 2.022 | 1.966 | 1.543 | 1.692 | 1.996 | 1.675 | 1.689 | 1.726 | 2.205 |
| 8. *Echinostoma hortense* | 2.194 | 2.267 | 2.228 | 2.304 | 2.119 | 2.343 | 2.340 | 2.242 | 2.248 | 2.305 | 2.291 | 2.190 | 2.250 | 2.188 | 2.169 | 2.273 |
| 9. *Echinostoma macrorchis* | 2.025 | 2.024 | 1.885 | 1.964 | 1.780 | 1.972 | 1.987 | 1.720 | 0.944 | 1.962 | 2.026 | 2.079 | 1.914 | 1.962 | 1.926 | 2.096 |
| 10. *Echinostoma maldonadoi* | 1.689 | 1.193 | 1.722 | 1.732 | 1.455 | 1.688 | 1.773 | 2.058 | 2.022 | 1.705 | 1.864 | 1.984 | 1.692 | 1.593 | 1.664 | 2.209 |
| 11. *Echinostoma mekongi* | 1.672 | 1.879 | 1.557 | 1.557 | 1.229 | 1.722 | 1.626 | 2.095 | 2.004 | 1.716 | 1.671 | 1.973 | 1.865 | 1.787 | 1.723 | 2.237 |
| 12. *Echinostoma miyagawai* | 1.500 | 1.792 | 1.534 | 1.342 | 1.325 | 1.560 | 0.235 | 2.092 | 2.026 | 1.537 | 1.713 | 1.939 | 1.756 | 1.747 | 1.697 | 2.292 |
| 13. *Echinostoma nasincovae* | 1.603 | 1.725 | 1.753 | 1.618 | 1.427 | 1.599 | 1.748 | 2.047 | 2.018 | 1.628 | 1.882 | 1.881 | 0.413 | 1.455 | 1.523 | 2.209 |
| 14. *Echinostoma novazealandense* (2) | - | 1.704 | 1.589 | 1.500 | 1.207 | 1.499 | 1.486 | 2.066 | 2.054 | 1.500 | 1.733 | 1.938 | 1.589 | 1.593 | 1.644 | 2.240 |
| 15. *Echinostoma paraensei* | 85.782 | - | 1.755 | 1.772 | 1.541 | 1.729 | 1.778 | 2.104 | 1.979 | 1.740 | 1.969 | 1.934 | 1.709 | 1.661 | 1.764 | 2.270 |
| 16. *Echinostoma paraulum* | 88.389 | 85.071 | - | 1.536 | 1.423 | 1.653 | 1.517 | 1.961 | 1.854 | 1.633 | 1.706 | 1.915 | 1.735 | 1.762 | 1.722 | 2.185 |
| 17. *Echinostoma pseudorobustum* | 89.810 | 84.834 | 89.100 | - | 1.322 | 1.637 | 1.314 | 2.091 | 2.055 | 1.589 | 1.805 | 1.899 | 1.606 | 1.588 | 1.656 | 2.238 |
| 18. *Echinostoma revolutum* (3) | 89.731 | 84.044 | 86.967 | 87.757 | - | 1.020 | 1.306 | 1.867 | 1.784 | 1.049 | 1.478 | 1.771 | 1.417 | 1.407 | 1.364 | 2.019 |
| 19. *Echinostoma revolutum* complex sp. B | 89.336 | 83.886 | 86.967 | 86.493 | 91.469 | - | 1.536 | 2.027 | 1.964 | 0.494 | 1.677 | 1.931 | 1.562 | 1.614 | 1.626 | 2.201 |

*Continued on next page*

Table B6 – *Continued from previous page*

| | 1 | 2 | 3 | 4 | 5 | 6 | 7 | 8 | 9 | 10 | 11 | 12 | 13 |
|---|---|---|---|---|---|---|---|---|---|---|---|---|---|
| 20. *Echinostoma robustum* | 89.100 | 86.019 | 83.412 | 85.071 | 80.806 | 84.834 | 89.810 | 65.321 | 76.485 | 83.649 | 87.441 | 99.763 | 85.308 |
| 21. *Echinostoma* sp. AF025833 | 76.722 | 75.772 | 76.960 | 75.534 | 76.247 | 77.435 | 76.010 | 65.166 | 84.597 | 75.534 | 75.772 | 74.822 | 76.960 |
| 22. *Echinostoma* sp. AF026290 | 75.891 | 77.197 | 76.722 | 76.722 | 75.059 | 76.960 | 76.010 | 67.062 | 96.209 | 76.722 | 75.772 | 76.010 | 76.247 |
| 23. *Echinostoma* sp. OR257455 | 99.406 | 86.223 | 84.323 | 85.748 | 80.523 | 85.511 | 88.836 | 66.667 | 75.714 | 84.561 | 86.461 | 88.836 | 87.173 |
| 24. *Echinostoma* sp. I | 85.427 | 84.123 | 84.834 | 83.412 | 78.436 | 84.123 | 85.308 | 65.796 | 76.247 | 82.464 | 86.493 | 84.360 | 83.649 |
| 25. *Echinostoma* sp. IG | 80.332 | 81.280 | 80.332 | 79.621 | 99.289 | 80.806 | 79.384 | 65.796 | 74.822 | 79.621 | 79.621 | 80.332 | 79.621 |
| 26. *Echinostoma* sp. n. | 87.559 | 88.863 | 83.649 | 84.360 | 80.095 | 84.360 | 85.782 | 65.796 | 76.960 | 86.967 | 82.938 | 85.071 | 99.289 |
| 27. *Echinostoma trivolvis* (2) | 85.782 | 91.943 | 85.071 | 83.768 | 79.739 | 84.242 | 85.782 | 65.558 | 76.366 | 86.493 | 84.716 | 84.953 | 88.981 |
| 28. *Echinostoma trivolvis* complex Lin A (2) | 86.374 | 99.763 | 84.360 | 85.900 | 81.161 | 82.701 | 86.730 | 66.390 | 76.603 | 87.085 | 86.374 | 85.782 | 88.744 |
| 29. *Isthmiophora melis* (out) | 72.565 | 70.546 | 71.971 | 68.884 | 70.309 | 69.834 | 72.684 | 68.884 | 70.784 | 71.971 | 72.209 | 71.734 | 71.259 |

*Continued on next page*

Table B6 – *Continued from previous page*

| | 14 | 15 | 16 | 17 | 18 | 19 | 20 | 21 | 22 | 23 | 24 | 25 | 26 | 27 | 28 | 29 |
|---|---|---|---|---|---|---|---|---|---|---|---|---|---|---|---|---|
| 20. *Echinostoma robustum* | 90.284 | 83.649 | 88.863 | 91.943 | 88.310 | 88.626 | - | 2.099 | 2.031 | 1.510 | 1.704 | 1.955 | 1.733 | 1.748 | 1.685 | 2.286 |
| 21. *Echinostoma* sp. AF025833 | 76.010 | 75.534 | 76.247 | 75.059 | 75.693 | 76.485 | 75.059 | - | 1.607 | 2.022 | 2.143 | 2.056 | 2.059 | 2.044 | 2.070 | 2.237 |
| 22. *Echinostoma* sp. AF026290 | 75.534 | 76.010 | 77.672 | 75.534 | 76.010 | 76.010 | 76.247 | 84.834 | - | 1.963 | 2.036 | 2.060 | 2.000 | 1.982 | 1.932 | 2.187 |
| 23. *Echinostoma* sp. OR257455 | 89.549 | 83.848 | 87.173 | 87.173 | 91.528 | 99.050 | 89.074 | 76.667 | 75.714 | - | 1.707 | 1.940 | 1.583 | 1.562 | 1.651 | 2.205 |
| 24. *Echinostoma* sp. I | 85.782 | 81.754 | 85.071 | 83.175 | 85.861 | 85.545 | 84.597 | 74.109 | 76.960 | 85.511 | - | 2.067 | 1.874 | 1.895 | 1.824 | 2.267 |
| 25. *Echinostoma* sp. IG | 80.332 | 79.858 | 80.095 | 80.332 | 79.779 | 80.332 | 80.095 | 76.010 | 74.822 | 80.285 | 78.673 | - | 1.870 | 1.910 | 1.859 | 2.213 |
| 26. *Echinostoma* sp. n. | 87.441 | 87.204 | 84.123 | 86.967 | 86.335 | 87.678 | 85.308 | 77.435 | 76.247 | 87.648 | 83.649 | 79.858 | - | 1.443 | 1.527 | 2.196 |
| 27. *Echinostoma trivolvis* (2) | 87.204 | 85.308 | 84.716 | 87.678 | 85.861 | 85.427 | 85.190 | 77.435 | 76.366 | 86.105 | 81.991 | 79.976 | 88.981 | - | 1.260 | 2.278 |
| 28. *Echinostoma trivolvis* complex Lin A (2) | 87.441 | 86.374 | 85.900 | 86.967 | 87.164 | 86.256 | 86.019 | 75.891 | 77.078 | 86.223 | 84.005 | 81.398 | 88.744 | 91.706 | - | 2.208 |
| 29. *Isthmiophora melis* (out) | 70.784 | 70.309 | 72.447 | 72.684 | 73.159 | 72.447 | 71.971 | 70.546 | 70.784 | 72.619 | 69.596 | 69.834 | 71.734 | 70.190 | 70.428 | - |

Table B7. Averaged percent similarities within the genera *Petasiger* and *Neopetasiger* (*nad1* gene). Standard error estimate(s) are shown above the diagonal and were obtained by a bootstrap procedure (500 replicates). Ambiguous positions were removed for each sequence pair. Values in parentheses indicate the number of representative sequences if > 1. Evolutionary analyses were conducted in MEGA (Tamura et al., 2021).

| | 1 | 2 | 3 | 4 | 5 | 6 | 7 | 8 | 9 | 10 | 11 |
|---|---|---|---|---|---|---|---|---|---|---|---|
| 1. BMC - *Petasiger* sp. 4 (2) | - | 2.208 | 1.965 | 2.255 | 2.125 | 2.506 | 1.925 | 2.351 | 0.377 | 2.393 | 2.317 |
| 2. *Fasciola hepatica* (out) | 73.146 | - | 2.273 | 2.257 | 2.342 | 2.273 | 1.882 | 2.143 | 2.208 | 2.303 | 2.166 |
| 3. *Neopetasiger islandicus* | 79.847 | 74.680 | - | 2.362 | 1.998 | 2.391 | 1.630 | 2.304 | 1.967 | 2.121 | 2.152 |
| 4. *Neopetasiger neocommense* | 72.564 | 75.064 | 71.282 | - | 2.315 | 2.394 | 2.079 | 2.286 | 2.270 | 2.439 | 2.230 |
| 5. *Neopetasiger sp. 5* | 74.169 | 69.744 | 81.074 | 69.565 | - | 2.436 | 1.888 | 2.345 | 2.121 | 2.407 | 2.360 |
| 6. *Petasiger* sp. 1 | 64.541 | 73.146 | 68.112 | 70.000 | 67.263 | - | 1.986 | 2.148 | 2.498 | 2.243 | 2.204 |
| 7. *Petasiger* sp. 2 (2) | 72.959 | 74.680 | 78.189 | 70.256 | 75.703 | 71.939 | - | 1.813 | 1.927 | 1.793 | 1.872 |
| 8. *Petasiger* sp. 3 (3) | 68.793 | 75.789 | 71.259 | 71.538 | 67.263 | 77.126 | 73.682 | - | 2.342 | 2.095 | 1.939 |
| 9. *Petasiger* sp. 4 (Alberta) | 99.490 | 73.146 | 79.592 | 72.564 | 73.913 | 64.286 | 72.704 | 68.537 | - | 2.397 | 2.321 |
| 10. *Petasiger* sp. 4 (Kenya) | 70.408 | 73.657 | 73.214 | 68.462 | 68.798 | 72.959 | 74.362 | 76.446 | 70.153 | - | 2.102 |
| 11. *Petasiger* sp. 5 (2) | 69.005 | 76.087 | 74.107 | 72.179 | 69.949 | 75.893 | 72.832 | 81.250 | 69.005 | 75.383 | - |

Table B8. Averaged percent similarities between the Leucochloridiidae. Standard error estimate(s) are shown above the diagonal and were obtained by a bootstrap procedure (500 replicates). Ambiguous positions were removed for each sequence pair. Values in parentheses indicate the number of representative sequences if > 1. Evolutionary analyses were conducted in MEGA (Tamura et al., 2021).

| | 1 | 2 | 3 | 4 | 5 | 6 | 7 | 8 | 9 | 10 | 11 |
|---|---|---|---|---|---|---|---|---|---|---|---|
| 1. BMC 1-1312 - *Leucochloridium* sp. | - | 2.099 | 2.048 | 1.893 | 1.953 | 2.077 | 1.983 | 1.903 | 1.903 | 1.893 | 1.901 |
| 2. *Brachylaima asakawai* (out) | 78.908 | - | 1.860 | 2.052 | 2.094 | 2.220 | 2.115 | 2.187 | 2.187 | 2.052 | 2.156 |
| 3. *B. ezohelicis* (out) | 79.156 | 84.367 | - | 2.057 | 2.100 | 2.098 | 2.114 | 2.146 | 2.146 | 2.057 | 2.110 |
| 4. *Leucochloridium* cf. *passeri* | 81.141 | 79.653 | 78.164 | - | 1.635 | 1.771 | 1.662 | 1.548 | 1.548 | 0.000 | 1.646 |
| 5. *Leucochloridium paradoxum* (6) | 80.521 | 77.006 | 75.931 | 86.642 | - | 1.737 | 0.535 | 1.755 | 1.755 | 1.635 | 1.799 |
| 6. *Leucochloridium perturbatum* (2) | 79.529 | 77.047 | 77.295 | 85.980 | 84.864 | - | 1.786 | 1.856 | 1.856 | 1.771 | 1.713 |
| 7. *Leucochloridium* sp. LC384429 | 80.645 | 77.171 | 75.682 | 86.849 | 98.222 | 84.615 | - | 1.802 | 1.802 | 1.662 | 1.829 |
| 8. *Leucochloridium* sp. LC648352 | 81.390 | 77.667 | 77.171 | 90.571 | 85.980 | 84.988 | 85.856 | - | 0.000 | 1.548 | 1.509 |
| 9. *Leucochloridium* sp. LC648354 | 81.390 | 77.667 | 77.171 | 90.571 | 85.980 | 84.988 | 85.856 | 100 | - | 1.548 | 1.509 |
| 10.*Leucochloridium* sp. (2) | 81.141 | 79.653 | 78.164 | 100 | 86.642 | 85.980 | 86.849 | 90.571 | 90.571 | - | 1.646 |
| 11. *Leucochloridium vogtianum* (4) | 82.010 | 77.667 | 77.792 | 88.089 | 83.912 | 85.732 | 84.119 | 89.454 | 89.454 | 88.089 | - |

Table B9. Averaged percent similarities of the Notocotylidae. Standard error estimate(s) are shown above the diagonal and were obtained by a bootstrap procedure (500 replicates). Ambiguous positions were removed for each sequence pair. Values in parentheses indicate the number of representative sequences if > 1. Evolutionary analyses were conducted in MEGA (Tamura et al., 2021).

| | 1 | 2 | 3 | 4 | 5 | 6 | 7 | 8 | 9 | 10 | 11 | 12 |
|---|---|---|---|---|---|---|---|---|---|---|---|---|
| 1. BMC *Notocotylus* sp. D (2) | - | 0.974 | 2.052 | 1.909 | 1.217 | 0.980 | 1.235 | 1.073 | 1.056 | 1.912 | 1.845 | 1.845 |
| 2. BMC *Notocotylus* sp. A (2) | 94.990 | - | 2.048 | 1.864 | 1.260 | 0.454 | 0.847 | 0.965 | 0.962 | 1.878 | 1.729 | 1.793 |
| 3. *Echinostoma hortense* (out) | 70.704 | 71.118 | - | 2.009 | 2.077 | 2.052 | 2.052 | 2.031 | 2.018 | 2.087 | 1.921 | 2.033 |
| 4. Notocotylidae sp. MSB | 76.440 | 78.292 | 68.880 | - | 1.928 | 1.886 | 1.835 | 1.913 | 1.894 | 1.842 | 1.795 | 1.864 |
| 5. *Notocotylus intestinalis* | 91.820 | 91.104 | 70.868 | 76.797 | - | 1.272 | 1.371 | 1.335 | 1.310 | 1.896 | 1.783 | 1.805 |
| 6. *Notocotylus* sp. A (2) | 95.297 | 98.875 | 70.868 | 77.823 | 91.020 | - | 0.969 | 0.986 | 0.977 | 1.921 | 1.747 | 1.801 |
| 7. *Notocotylus* sp. B (2) | 91.393 | 96.209 | 71.429 | 77.823 | 89.571 | 95.297 | - | 1.177 | 1.153 | 1.944 | 1.759 | 1.832 |
| 8. *Notocotylus* sp. BOLD | 93.865 | 95.194 | 70.807 | 78.189 | 91.411 | 95.092 | 92.213 | - | 0.296 | 1.882 | 1.795 | 1.814 |
| 9. *Notocotylus* sp. D | 93.865 | 95.194 | 71.074 | 78.439 | 91.633 | 95.102 | 92.434 | 99.591 | - | 1.868 | 1.774 | 1.799 |
| 10. *Ogmocotyle ailuri* | 77.812 | 77.914 | 71.281 | 77.413 | 78.571 | 77.347 | 77.096 | 78.119 | 78.367 | - | 1.584 | 1.583 |
| 11. *Ogmocotyle sikae* | 79.755 | 81.493 | 72.727 | 79.877 | 80.204 | 81.020 | 80.982 | 80.368 | 80.612 | 85.510 | - | 1.517 |
| 12. *Ogmocotyle* sp. | 78.279 | 79.816 | 71.222 | 77.823 | 79.755 | 79.346 | 79.592 | 79.098 | 79.346 | 85.276 | 86.503 | - |

Table B10. Averaged percent similarities of *Plagiorchis* between species. Standard error estimate(s) are shown above the diagonal and were obtained by a bootstrap procedure (500 replicates). Ambiguous positions were removed for each sequence pair. Evolutionary analyses were conducted in MEGA (Tamura et al., 2021).

| | 1 | 2 | 3 | 4 | 5 | 6 | 7 | 8 | 9 | 10 | 11 | 12 | 13 |
|---|---|---|---|---|---|---|---|---|---|---|---|---|---|
| 1. BMC - *Plagiorchis* sp. Lin 6 (2) | - | 2.040 | 1.875 | 1.926 | 2.030 | 1.880 | 1.980 | 2.278 | 1.702 | 1.814 | 1.862 | 2.116 | 1.815 |
| 2. BMC - *Plagiorchis* sp. Lin 1 (2) | 84.129 | - | 2.011 | 1.829 | 2.006 | 2.016 | 1.976 | 2.265 | 2.007 | 1.940 | 1.975 | 2.043 | 2.081 |
| 3. BMC - *Plagiorchis* sp. Lin 5 | 85.846 | 85.670 | - | 1.867 | 2.114 | 1.829 | 2.086 | 2.313 | 1.814 | 2.030 | 1.991 | 2.064 | 2.113 |
| 4. BMC - *Plagiorchis* sp. Lin 4 (2) | 86.923 | 87.519 | 86.154 | - | 1.890 | 1.878 | 2.012 | 2.245 | 1.850 | 2.051 | 1.906 | 1.956 | 1.986 |
| 5. BMC - *Plagiorchis* sp. Lin 2 (2) | 84.900 | 84.722 | 83.975 | 85.670 | - | 1.952 | 2.112 | 2.452 | 1.972 | 2.069 | 1.422 | 2.059 | 2.061 |
| 6. BMC - *Plagiorchis* sp. Lin 7 (2) | 86.864 | 84.831 | 87.018 | 85.010 | 85.229 | - | 1.876 | 2.432 | 1.179 | 2.051 | 1.829 | 2.066 | 1.843 |
| 7. BMC - *Plagiorchis* sp. Lin 9 | 84.877 | 85.781 | 83.333 | 84.414 | 83.617 | 84.521 | - | 2.384 | 1.903 | 1.937 | 2.124 | 2.210 | 2.113 |
| 8. *Haematoloechus* sp. (out) | 79.077 | 79.198 | 77.846 | 79.385 | 76.579 | 75.734 | 76.852 | - | 2.402 | 2.400 | 2.305 | 2.271 | 2.409 |
| 9. *Plagiorchis elegans* | 89.231 | 85.978 | 86.769 | 85.846 | 85.516 | 94.748 | 85.494 | 76.923 | - | 2.035 | 1.939 | 2.064 | 1.905 |
| 10. *Plagiorchis koreanus* | 86.769 | 86.133 | 84.923 | 86.154 | 83.975 | 83.155 | 85.494 | 77.846 | 83.077 | - | 2.025 | 2.108 | 1.875 |
| 11. *Plagiorchis maculosus* | 85.538 | 85.362 | 85.846 | 85.385 | 92.604 | 87.329 | 83.951 | 77.538 | 87.077 | 83.692 | - | 1.940 | 2.077 |
| 12. *Plagiorchis muelleri* | 83.077 | 84.745 | 84.000 | 86.615 | 83.975 | 82.537 | 82.407 | 80.000 | 82.769 | 84.923 | 84.923 | - | 2.104 |
| 13. *Plagiorchis* sp. 3 | 88.000 | 84.437 | 85.538 | 86.000 | 84.283 | 89.028 | 83.951 | 78.154 | 87.692 | 87.077 | 84.923 | 84.615 | - |
| 14. *Plagiorchis* sp. 5 | 85.846 | 89.368 | 84.923 | 91.077 | 83.975 | 86.089 | 84.259 | 78.462 | 86.462 | 84.308 | 86.154 | 85.846 | 88.000 |
| 15. *Plagiorchis* sp. 8 | 88.615 | 87.211 | 86.462 | 90.154 | 86.749 | 86.246 | 85.802 | 78.769 | 87.077 | 88.615 | 86.462 | 84.923 | 87.692 |
| 16. *Plagiorchis* sp. 10 | 84.923 | 85.978 | 86.154 | 87.231 | 85.208 | 84.700 | 82.099 | 79.692 | 84.308 | 82.462 | 86.154 | 88.615 | 83.692 |
| 17. *Plagiorchis* sp. 11 | 86.154 | 85.362 | 83.692 | 85.846 | 82.434 | 83.155 | 82.716 | 76.000 | 85.231 | 85.231 | 83.077 | 84.923 | 84.000 |
| 18. *Plagiorchis* sp. | 87.385 | 87.519 | 86.154 | 99.385 | 85.516 | 84.547 | 84.259 | 79.692 | 85.538 | 86.154 | 84.923 | 86.462 | 85.846 |
| 19. *Plagiorchis* sp. Lin 1 (2) | 84.154 | 99.692 | 85.692 | 87.538 | 84.746 | 84.855 | 85.340 | 79.231 | 86.000 | 85.692 | 85.385 | 84.769 | 84.462 |
| 20. *Plagiorchis* sp. Lin 2 | 84.615 | 83.205 | 84.000 | 85.385 | 95.069 | 83.927 | 83.333 | 77.538 | 85.231 | 84.308 | 92.615 | 84.615 | 82.769 |
| 21. *Plagiorchis* sp. Lin 3 | 84.615 | 86.286 | 82.154 | 86.462 | 82.434 | 83.310 | 82.407 | 76.615 | 85.231 | 83.692 | 84.615 | 83.385 | 84.308 |
| 22. *Plagiorchis* sp. Lin 4 (2) | 86.923 | 87.827 | 86.769 | 99.385 | 85.516 | 84.701 | 84.722 | 79.692 | 85.692 | 86.154 | 85.231 | 86.769 | 86.000 |
| 23. *Plagiorchis* sp. Lin 5 | 86.154 | 85.978 | 99.692 | 86.462 | 84.283 | 87.327 | 83.642 | 78.154 | 87.077 | 85.231 | 86.154 | 84.308 | 85.846 |
| 24. *Plagiorchis* sp. Lin 6 | 99.385 | 83.821 | 85.538 | 86.615 | 84.592 | 86.555 | 84.568 | 79.385 | 88.923 | 86.769 | 85.231 | 83.077 | 88.615 |
| 25. *Plagiorchis* sp. Lin 7 (2) | 86.615 | 84.591 | 86.923 | 85.231 | 85.054 | 99.230 | 84.722 | 75.538 | 94.923 | 83.231 | 87.846 | 82.615 | 89.077 |

*Continued on next page*

Table B10 – *Continued from previous page*

| | 14 | 15 | 16 | 17 | 18 | 19 | 20 | 21 | 22 | 23 | 24 | 25 | 26 | 27 | 28 |
|---|---|---|---|---|---|---|---|---|---|---|---|---|---|---|---|
| 1. BMC - *Plagiorchis* sp. Lin 6 (2) | 1.868 | 1.779 | 2.005 | 1.910 | 1.859 | 2.053 | 2.036 | 2.006 | 1.918 | 1.828 | 0.435 | 1.895 | 1.860 | 1.976 | 1.875 |
| 2. BMC - *Plagiorchis* sp. Lin 1 (2) | 1.665 | 1.762 | 2.013 | 1.976 | 1.845 | 0.235 | 2.124 | 2.012 | 1.817 | 1.993 | 2.062 | 2.038 | 2.077 | 1.947 | 1.924 |
| 3. BMC - *Plagiorchis* sp. Lin 5 | 1.957 | 2.027 | 1.913 | 2.054 | 1.891 | 2.007 | 2.071 | 2.243 | 1.848 | 0.304 | 1.911 | 1.868 | 2.055 | 2.076 | 1.895 |
| 4. BMC - *Plagiorchis* sp. Lin 4 (2) | 1.527 | 1.574 | 1.883 | 1.891 | 0.367 | 1.833 | 1.941 | 1.929 | 0.307 | 1.846 | 1.951 | 1.864 | 2.038 | 2.029 | 1.785 |
| 5. BMC - *Plagiorchis* sp. Lin 2 (2) | 2.020 | 1.822 | 1.961 | 2.075 | 1.893 | 2.015 | 1.227 | 2.169 | 1.895 | 2.098 | 2.065 | 1.969 | 2.144 | 2.121 | 1.972 |
| 6. BMC - *Plagiorchis* sp. Lin 7 (2) | 1.909 | 1.852 | 1.954 | 2.076 | 1.900 | 2.018 | 2.041 | 2.038 | 1.883 | 1.800 | 1.917 | 0.322 | 1.920 | 1.889 | 1.744 |
| 7. BMC - *Plagiorchis* sp. Lin 9 | 2.079 | 1.847 | 2.173 | 2.082 | 2.023 | 2.018 | 2.169 | 2.115 | 2.009 | 2.074 | 2.016 | 1.879 | 1.631 | 0.316 | 1.877 |
| 8. *Haematoloechus* sp. (out) | 2.330 | 2.370 | 2.286 | 2.439 | 2.258 | 2.262 | 2.262 | 2.398 | 2.248 | 2.308 | 2.258 | 2.439 | 2.295 | 2.377 | 2.266 |
| 9. *Plagiorchis elegans* | 1.855 | 1.836 | 2.048 | 1.974 | 1.885 | 2.009 | 2.023 | 2.005 | 1.866 | 1.787 | 1.735 | 1.195 | 1.931 | 1.894 | 1.781 |
| 10. *Plagiorchis koreanus* | 2.047 | 1.680 | 2.154 | 1.968 | 2.088 | 1.982 | 2.139 | 2.138 | 2.055 | 1.999 | 1.814 | 2.051 | 1.988 | 1.890 | 2.006 |
| 11. *Plagiorchis maculosus* | 1.923 | 1.778 | 1.871 | 1.920 | 1.920 | 1.974 | 1.459 | 2.035 | 1.904 | 1.979 | 1.901 | 1.832 | 2.009 | 2.143 | 1.798 |
| 12. *Plagiorchis muelleri* | 1.901 | 2.082 | 1.760 | 1.993 | 1.966 | 2.048 | 1.949 | 2.072 | 1.937 | 2.041 | 2.116 | 2.065 | 2.132 | 2.190 | 1.967 |
| 13. *Plagiorchis* sp. 3 | 1.900 | 1.879 | 2.046 | 2.163 | 2.002 | 2.099 | 2.199 | 2.021 | 1.979 | 2.084 | 1.738 | 1.876 | 2.030 | 2.085 | 1.791 |
| 14. *Plagiorchis* sp. 5 | - | 1.798 | 1.947 | 1.945 | 1.564 | 1.672 | 1.975 | 1.972 | 1.515 | 1.939 | 1.901 | 1.888 | 2.072 | 2.105 | 1.799 |
| 15. *Plagiorchis* sp. 8 | 87.692 | - | 2.079 | 1.567 | 1.618 | 1.780 | 1.919 | 1.958 | 1.580 | 2.010 | 1.815 | 1.861 | 1.900 | 1.853 | 1.865 |
| 16. *Plagiorchis* sp. 10 | 85.846 | 86.154 | - | 1.952 | 1.894 | 2.021 | 1.986 | 2.221 | 1.865 | 1.890 | 1.972 | 1.969 | 2.022 | 2.175 | 1.891 |
| 17. *Plagiorchis* sp. 11 | 85.231 | 90.462 | 86.154 | - | 1.927 | 1.989 | 2.108 | 2.060 | 1.889 | 2.027 | 1.924 | 2.095 | 1.931 | 2.077 | 2.002 |
| 18. *Plagiorchis* sp. | 91.077 | 89.846 | 87.077 | 85.538 | - | 1.848 | 1.944 | 1.968 | 0.371 | 1.871 | 1.886 | 1.883 | 2.057 | 2.041 | 1.813 |
| 19. *Plagiorchis* sp. Lin 1 (2) | 89.385 | 87.231 | 86.000 | 85.077 | 87.538 | - | 2.121 | 2.017 | 1.819 | 1.990 | 2.077 | 2.038 | 2.096 | 1.991 | 1.929 |
| 20. *Plagiorchis* sp. Lin 2 | 84.000 | 86.154 | 84.615 | 81.538 | 85.231 | 83.231 | - | 2.186 | 1.925 | 2.059 | 2.036 | 2.055 | 2.212 | 2.189 | 1.983 |
| 21. *Plagiorchis* sp. Lin 3 | 85.538 | 85.846 | 81.538 | 84.308 | 86.154 | 86.308 | 82.769 | - | 1.903 | 2.217 | 2.009 | 2.040 | 2.212 | 2.080 | 2.041 |
| 22. *Plagiorchis* sp. Lin 4 (2) | 91.385 | 89.846 | 87.385 | 85.846 | 99.385 | 87.846 | 85.538 | 86.769 | - | 1.826 | 1.944 | 1.869 | 2.030 | 2.023 | 1.757 |
| 23. *Plagiorchis* sp. Lin 5 | 85.231 | 86.769 | 86.462 | 84.000 | 86.462 | 86.000 | 84.308 | 82.462 | 87.077 | - | 1.867 | 1.839 | 2.030 | 2.062 | 1.886 |
| 24. *Plagiorchis* sp. Lin 6 | 85.538 | 88.308 | 85.231 | 85.846 | 87.077 | 83.846 | 84.615 | 84.308 | 86.615 | 85.846 | - | 1.930 | 1.916 | 2.013 | 1.875 |
| 25. *Plagiorchis* sp. Lin 7 (2) | 85.692 | 86.308 | 84.462 | 82.923 | 84.769 | 84.615 | 84.154 | 83.538 | 84.923 | 87.231 | 86.308 | - | 1.936 | 1.899 | 1.772 |

*Continued on next page*

Table B10 – *Continued from previous page*

| | 1 | 2 | 3 | 4 | 5 | 6 | 7 | 8 | 9 | 10 | 11 | 12 | 13 |
|---|---|---|---|---|---|---|---|---|---|---|---|---|---|
| 26. *Plagiorchis* sp. Lin 8 | 87.692 | 84.592 | 83.692 | 85.077 | 81.818 | 85.317 | 91.049 | 78.154 | 85.538 | 86.462 | 83.692 | 84.308 | 86.462 |
| 27. *Plagiorchis* sp. Lin 9 (2) | 84.462 | 85.979 | 83.385 | 84.231 | 83.205 | 83.773 | 99.383 | 76.923 | 85.231 | 86.000 | 83.231 | 82.462 | 83.538 |
| 28. *Plagiorchis vespertilionis* | 87.077 | 86.286 | 85.538 | 88.462 | 85.824 | 87.018 | 86.111 | 79.077 | 87.077 | 84.615 | 87.385 | 87.077 | 87.077 |

| | 14 | 15 | 16 | 17 | 18 | 19 | 20 | 21 | 22 | 23 | 24 | 25 | 26 | 27 | 28 |
|---|---|---|---|---|---|---|---|---|---|---|---|---|---|---|---|
| 26. *Plagiorchis* sp. Lin 8 | 85.538 | 85.846 | 84.308 | 85.231 | 84.923 | 84.462 | 80.923 | 81.846 | 85.077 | 84.000 | 87.077 | 84.769 | - | 1.635 | 1.824 |
| 27. *Plagiorchis* sp. Lin 9 (2) | 83.692 | 85.538 | 81.692 | 82.769 | 84.154 | 85.538 | 82.615 | 82.769 | 84.615 | 83.692 | 84.154 | 83.846 | 90.769 | - | 1.882 |
| 28. *Plagiorchis vespertilionis* | 88.308 | 86.462 | 87.692 | 84.615 | 88.308 | 86.308 | 85.538 | 84.615 | 88.615 | 85.846 | 87.077 | 86.615 | 88.000 | 85.692 | - |

Table B11. Percent similarities of the available *Manodistomum* species from the Plagiorchiidae along with *Plagiorchis* sp. Lineages from Alberta. Values were not averaged due to the limited availability of sequences. Standard error estimate(s) are shown above the diagonal. Ambiguous positions were removed for each sequence pair. Evolutionary analyses were conducted in MEGA (Tamura et al., 2021).

| | 1 | 2 | 3 | 4 | 5 | 6 | 7 | 8 | 9 | 10 | 11 | 12 |
|---|---|---|---|---|---|---|---|---|---|---|---|---|
| 1. BMC 1973 - *Manodistomum* sp. | - | 2.646 | 2.175 | 1.927 | 2.573 | 2.490 | 2.546 | 2.467 | 2.475 | 2.504 | 2.519 | 2.460 |
| 2. *Haematoloechus* sp. (KM538096) (out) | 73.214 | - | 2.661 | 2.661 | 2.497 | 2.511 | 2.482 | 2.515 | 2.452 | 2.651 | 2.482 | 2.565 |
| 3. *Manodistomum* sp. (HOSL287-19) | 84.286 | 72.598 | - | 1.945 | 2.539 | 2.452 | 2.421 | 2.459 | 2.482 | 2.482 | 2.405 | 2.421 |
| 4. *Manodistomum* sp. (TREMA2456-10) | 88.214 | 72.598 | 87.9 | - | 2.591 | 2.497 | 2.565 | 2.488 | 2.452 | 2.525 | 2.511 | 2.482 |
| 5. *Plagiorchis* sp. Lineage 1 (MH369420) | 75.812 | 77.698 | 76.619 | 75.18 | - | 2.229 | 1.965 | 2.105 | 2.229 | 2.189 | 2.189 | 2.105 |
| 6. *Plagiorchis* sp. Lineage 2 (MH369467) | 77.978 | 77.338 | 78.777 | 77.698 | 83.453 | - | 2.169 | 2.209 | 2.169 | 2.209 | 2.303 | 2.229 |
| 7. *Plagiorchis* sp. Lineage 4 (MH369418) | 76.534 | 78.058 | 79.496 | 75.899 | 87.77 | 84.532 | - | 2.083 | 2.148 | 2.229 | 2.127 | 2.148 |
| 8. *Plagiorchis* sp. Lineage 5 (MH369419) | 78.214 | 76.868 | 78.292 | 77.58 | 85.612 | 83.813 | 85.971 | - | 2.127 | 2.037 | 2.209 | 2.189 |
| 9. *Plagiorchis* sp. Lineage 6 (MH369470) | 78.339 | 78.777 | 78.058 | 78.777 | 83.453 | 84.532 | 84.892 | 85.252 | - | 2.060 | 1.940 | 2.189 |
| 10. *Plagiorchis* sp. Lineage 7 (MH369458) | 77.617 | 73.381 | 78.058 | 76.978 | 84.173 | 83.813 | 83.453 | 86.691 | 86.331 | - | 2.105 | 2.209 |
| 11. *Plagiorchis* sp. Lineage 8 (MH369449) | 77.256 | 78.058 | 79.856 | 77.338 | 84.173 | 82.014 | 85.252 | 83.813 | 88.129 | 85.612 | - | 1.805 |
| 12. *Plagiorchis* sp. Lineage 9 (MH369424) | 78.700 | 75.899 | 79.496 | 78.058 | 85.612 | 83.453 | 84.892 | 84.173 | 84.173 | 83.813 | 89.928 | - |

Table B12. Percent similarities of Psilostomidae. Standard error estimate(s) are shown above the diagonal and were obtained by a bootstrap procedure (500 replicates). Ambiguous positions were removed for each sequence pair. Values in parentheses indicate the number of representative sequences if > 1. Evolutionary analyses were conducted in MEGA (Tamura et al., 2021).

| | 1 | 2 | 3 | 4 | 5 | 6 | 7 |
|---|---|---|---|---|---|---|---|
| 1. BMC Psilostomidae gen. sp. A (2) | - | 1.975 | 1.637 | 0.105 | 2.454 | 0.230 | 2.091 |
| 2. *Echinocasmus japonicus* (out) | 78.571 | - | 2.012 | 1.972 | 2.520 | 1.975 | 1.975 |
| 3. *Pseudosilostoma varium* | 85.459 | 79.372 | - | 1.626 | 2.369 | 1.654 | 2.153 |
| 4. Psilostomidae gen. sp. A (2) | 99.889 | 78.683 | 85.570 | - | 2.448 | 0.260 | 2.093 |
| 5. *R. ondatrae* (Johnson et al. 2021) (2) | 64.212 | 58.161 | 62.824 | 64.341 | - | 2.454 | 2.592 |
| 6. *R. ondatrae* (Keller et al., 2021) | 99.778 | 78.571 | 85.235 | 99.667 | 64.212 | - | 2.074 |
| 7. *Sphaeridiotrema globulus* | 76.615 | 79.688 | 74.330 | 76.726 | 55.426 | 76.837 | - |

Table B13. Percent similarities of the avian schistosomes. Standard error estimate(s) are shown above the diagonal and were obtained by a bootstrap procedure (500 replicates). Ambiguous positions were removed for each sequence pair. Values in parentheses indicate the number of representative sequences if > 1. Evolutionary analyses were conducted in MEGA (Tamura et al., 2021).

| | 1 | 2 | 3 | 4 | 5 | 6 | 7 | 8 | 9 | 10 | 11 |
|---|---|---|---|---|---|---|---|---|---|---|---|
| 1. *Allobilharzia visceralis* | - | 1.764 | 1.971 | 1.875 | 1.771 | 1.633 | 1.786 | 1.885 | 1.913 | 1.736 | 1.977 |
| 2. *Anserobilharzia brantae* | 84.729 | - | 1.835 | 1.958 | 1.773 | 1.667 | 1.741 | 1.960 | 2.030 | 1.808 | 1.850 |
| 3. Avian schistosomatid sp. A | 81.034 | 83.744 | - | 1.687 | 1.846 | 1.912 | 1.909 | 1.692 | 1.933 | 1.890 | 0.344 |
| 4. Avian schistosomatid sp. B | 82.266 | 81.034 | 85.222 | - | 1.838 | 1.977 | 1.915 | 0.121 | 2.038 | 1.777 | 1.698 |
| 5. Avian schistosomatid sp. C | 84.975 | 83.251 | 82.266 | 82.020 | - | 1.907 | 1.802 | 1.844 | 2.004 | 1.795 | 1.875 |
| 6. BMC - *Trichobilharzia* sp. A (2) | 85.450 | 85.451 | 82.244 | 80.272 | 81.875 | - | 1.569 | 1.981 | 1.768 | 1.672 | 1.929 |
| 7. BMC - *Trichobilharzia physellae* (2) | 84.094 | 85.944 | 81.505 | 81.998 | 84.340 | 89.137 | - | 1.918 | 1.894 | 1.749 | 1.922 |
| 8. BMC - Avian schistosomatid sp. B (2) | 82.143 | 80.911 | 85.099 | 99.877 | 81.897 | 80.148 | 81.875 | - | 2.045 | 1.778 | 1.704 |
| 9. BMC - *Gigantobilharzia huronensis* (2) | 80.172 | 79.187 | 80.172 | 79.433 | 78.695 | 82.121 | 80.395 | 79.310 | - | 1.966 | 1.956 |
| 10. BMC - *Trichobilharzia szidati* (2) | 84.483 | 85.468 | 83.005 | 83.498 | 84.236 | 87.176 | 86.067 | 83.374 | 79.310 | - | 1.907 |
| 11. BMC - Avian schistosomatid sp. A | 80.542 | 83.251 | 99.507 | 84.975 | 81.773 | 81.751 | 81.011 | 84.852 | 79.680 | 82.512 | - |
| 12. *Drenditobilharzia pulverulenta* | 80.846 | 82.836 | 79.851 | 78.856 | 80.100 | 79.702 | 81.196 | 78.731 | 75.373 | 82.463 | 79.353 |
| 13. *Gigantobilharzia huronensis* | 80.296 | 79.064 | 80.296 | 80.296 | 79.064 | 81.998 | 80.272 | 80.172 | 98.645 | 79.680 | 79.803 |
| 14. *Nasusbilharzia melancorypha* | 76.601 | 80.296 | 78.079 | 81.773 | 79.803 | 79.655 | 79.778 | 81.650 | 78.941 | 79.803 | 78.079 |
| 15. Schistosomatidae sp. (out) (2) | 83.251 | 82.635 | 81.158 | 80.049 | 83.374 | 83.847 | 81.998 | 79.926 | 79.557 | 84.606 | 80.665 |
| 16. *Trichobilharzia franki* | 83.005 | 83.005 | 80.542 | 79.557 | 81.034 | 89.272 | 89.026 | 79.433 | 78.202 | 87.192 | 80.049 |
| 17. *Trichobilharzia mergi* | 85.961 | 84.729 | 81.773 | 81.527 | 85.468 | 85.574 | 86.683 | 81.404 | 80.419 | 88.547 | 81.281 |
| 18. *Trichobilharzia novaseelandiae* | 86.207 | 86.453 | 82.020 | 80.542 | 84.729 | 88.410 | 87.300 | 80.419 | 79.680 | 88.054 | 82.020 |
| 19. *Trichobilharzia physellae* | 83.990 | 85.714 | 81.527 | 82.020 | 84.236 | 89.026 | 99.260 | 81.897 | 80.172 | 86.084 | 81.034 |
| 20. *Trichobilharzia querquedulae* | 84.691 | 86.173 | 83.457 | 82.222 | 83.457 | 91.595 | 91.842 | 82.099 | 80.864 | 88.519 | 82.963 |
| 21. *Trichobilharzia regenti* | 86.946 | 87.685 | 81.527 | 80.296 | 84.975 | 88.903 | 88.533 | 80.172 | 81.897 | 87.192 | 81.034 |
| 22. *Trichobilharzia* sp. A | 86.207 | 85.961 | 83.005 | 81.034 | 82.512 | 99.014 | 89.766 | 80.911 | 82.882 | 87.562 | 82.512 |
| 23. *Trichobilharzia* sp. B | 87.438 | 85.468 | 81.773 | 80.788 | 83.498 | 92.972 | 91.492 | 80.665 | 82.882 | 87.808 | 81.281 |
| 24. *Trichobilharzia* sp. C | 84.483 | 86.946 | 81.281 | 82.020 | 81.773 | 90.136 | 90.753 | 81.897 | 79.926 | 87.069 | 80.788 |

*Continued on next page*

Table B13 – *Continued from previous page*

| | 12 | 13 | 14 | 15 | 16 | 17 | 18 | 19 | 20 | 21 | 22 | 23 | 24 |
|---|---|---|---|---|---|---|---|---|---|---|---|---|---|
| 1. *Allobilharzia visceralis* | 2.070 | 1.932 | 2.076 | 1.798 | 1.814 | 1.743 | 1.640 | 1.822 | 1.725 | 1.666 | 1.634 | 1.549 | 1.814 |
| 2. *Anserobilharzia brantae* | 1.928 | 2.055 | 1.994 | 1.864 | 1.852 | 1.731 | 1.678 | 1.797 | 1.752 | 1.660 | 1.661 | 1.771 | 1.647 |
| 3. Avian schistosomatid sp. A | 2.046 | 1.929 | 2.011 | 1.943 | 1.934 | 1.838 | 1.827 | 1.922 | 1.818 | 1.888 | 1.892 | 1.915 | 1.932 |
| 4. Avian schistosomatid sp. B | 2.026 | 2.000 | 1.957 | 1.953 | 2.031 | 1.883 | 1.989 | 1.918 | 1.961 | 1.972 | 1.980 | 2.016 | 1.895 |
| 5. Avian schistosomatid sp. C | 2.084 | 2.016 | 1.974 | 1.863 | 1.904 | 1.746 | 1.806 | 1.829 | 1.929 | 1.789 | 1.929 | 1.864 | 1.941 |
| 6. BMC - *Trichobilharzia* sp. A (2) | 2.091 | 1.800 | 1.939 | 1.772 | 1.516 | 1.701 | 1.538 | 1.590 | 1.402 | 1.549 | 0.402 | 1.218 | 1.486 |
| 7. BMC - *Trichobilharzia physellae* (2) | 2.023 | 1.917 | 1.949 | 1.880 | 1.490 | 1.678 | 1.640 | 0.345 | 1.358 | 1.589 | 1.549 | 1.363 | 1.479 |
| 8. BMC - Avian schistosomatid sp. B (2) | 2.023 | 2.005 | 1.960 | 1.961 | 2.031 | 1.893 | 1.998 | 1.923 | 1.963 | 1.980 | 1.985 | 2.023 | 1.899 |
| 9. BMC - *Gigantobilharzia huronensis* (2) | 2.261 | 0.579 | 1.955 | 1.935 | 1.956 | 1.988 | 1.940 | 1.926 | 1.863 | 1.994 | 1.786 | 1.792 | 2.010 |
| 10. BMC - *Trichobilharzia szidati* (2) | 1.837 | 1.918 | 1.966 | 1.687 | 1.693 | 1.463 | 1.607 | 1.740 | 1.659 | 1.638 | 1.690 | 1.716 | 1.723 |
| 11. BMC - Avian schistosomatid sp. A | 2.072 | 1.949 | 2.002 | 1.959 | 1.934 | 1.846 | 1.806 | 1.936 | 1.840 | 1.892 | 1.906 | 1.921 | 1.946 |
| 12. *Drenditobilharzia pulverulenta* | - | 2.192 | 2.122 | 1.947 | 2.073 | 1.915 | 2.070 | 2.045 | 1.996 | 2.029 | 2.089 | 2.040 | 2.017 |
| 13. *Gigantobilharzia huronensis* | 76.368 | - | 1.954 | 1.980 | 2.003 | 1.969 | 1.982 | 1.941 | 1.902 | 2.016 | 1.816 | 1.837 | 2.006 |
| 14. *Nasusbilharzia melancorypha* | 75.871 | 79.064 | - | 2.001 | 2.054 | 1.962 | 2.094 | 1.948 | 1.975 | 2.093 | 1.940 | 1.983 | 2.014 |
| 15. Schistosomatidae sp. (out) (2) | 79.975 | 79.433 | 79.557 | - | 1.837 | 1.785 | 1.864 | 1.902 | 1.752 | 1.797 | 1.754 | 1.816 | 1.833 |
| 16. *Trichobilharzia franki* | 79.851 | 77.586 | 78.079 | 82.389 | - | 1.651 | 1.713 | 1.516 | 1.393 | 1.727 | 1.530 | 1.486 | 1.627 |
| 17. *Trichobilharzia mergi* | 82.090 | 80.542 | 80.049 | 84.236 | 86.700 | - | 1.572 | 1.715 | 1.602 | 1.457 | 1.703 | 1.607 | 1.714 |
| 18. *Trichobilharzia novaseelandiae* | 78.856 | 79.310 | 79.310 | 83.374 | 87.192 | 89.163 | - | 1.680 | 1.587 | 1.398 | 1.548 | 1.464 | 1.709 |
| 19. *Trichobilharzia physellae* | 81.095 | 80.049 | 79.557 | 82.143 | 89.163 | 86.700 | 87.192 | - | 1.384 | 1.623 | 1.566 | 1.398 | 1.507 |
| 20. *Trichobilharzia querquedulae* | 82.294 | 80.741 | 81.235 | 84.074 | 91.605 | 88.889 | 89.136 | 91.852 | - | 1.554 | 1.370 | 1.226 | 1.389 |
| 21. *Trichobilharzia regenti* | 81.343 | 81.773 | 79.064 | 84.606 | 85.961 | 89.901 | 91.626 | 88.424 | 88.889 | - | 1.578 | 1.471 | 1.639 |
| 22. *Trichobilharzia* sp. A | 80.100 | 82.759 | 80.296 | 84.606 | 89.409 | 86.207 | 88.916 | 89.655 | 92.099 | 89.163 | - | 1.180 | 1.458 |
| 23. *Trichobilharzia* sp. B | 80.846 | 82.512 | 80.542 | 83.621 | 90.148 | 88.670 | 89.901 | 91.379 | 93.333 | 90.394 | 93.842 | - | 1.410 |
| 24. *Trichobilharzia* sp. C | 80.597 | 80.049 | 80.788 | 84.113 | 88.916 | 86.700 | 87.192 | 90.640 | 91.852 | 88.916 | 90.887 | 92.118 | - |
| 25. *Trichobilharzia* sp. D | 82.090 | 81.281 | 80.296 | 84.360 | 85.714 | 88.670 | 87.438 | 86.700 | 88.642 | 87.931 | 87.685 | 88.424 | 87.438 |

*Continued on next page*

Table B13 – *Continued from previous page*

| | 25 | 26 | 27 | 28 |
|---|---|---|---|---|
| 1. *Allobilharzia visceralis* | 1.766 | 1.779 | 1.696 | 1.720 |
| 2. *Anserobilharzia brantae* | 1.740 | 1.792 | 1.681 | 1.715 |
| 3. Avian schistosomatid sp. A | 1.947 | 1.846 | 1.756 | 1.875 |
| 4. Avian schistosomatid sp. B | 1.877 | 1.950 | 1.871 | 1.770 |
| 5. Avian schistosomatid sp. C | 1.766 | 1.712 | 1.692 | 1.766 |
| 6. BMC - *Trichobilharzia* sp. A (2) | 1.603 | 1.702 | 1.648 | 1.653 |
| 7. BMC - *Trichobilharzia physellae* (2) | 1.619 | 1.659 | 1.806 | 1.696 |
| 8. BMC - Avian schistosomatid sp. B (2) | 1.885 | 1.950 | 1.874 | 1.774 |
| 9. BMC - *Gigantobilharzia huronensis* (2) | 1.891 | 1.984 | 1.985 | 1.930 |
| 10. BMC - *Trichobilharzia szidati* (2) | 1.622 | 1.537 | 1.648 | 1.139 |
| 11. BMC - Avian schistosomatid sp. A | 1.956 | 1.863 | 1.774 | 1.892 |
| 12. *Drenditobilharzia pulverulenta* | 1.956 | 1.828 | 1.942 | 1.898 |
| 13. *Gigantobilharzia huronensis* | 1.930 | 2.015 | 1.967 | 1.896 |
| 14. *Nasusbilharzia melancorypha* | 2.003 | 2.045 | 2.042 | 1.912 |
| 15. Schistosomatidae sp. (out) (2) | 1.748 | 1.775 | 1.856 | 1.700 |
| 16. *Trichobilharzia franki* | 1.686 | 1.703 | 1.811 | 1.738 |
| 17. *Trichobilharzia mergi* | 1.523 | 1.655 | 1.768 | 1.454 |
| 18. *Trichobilharzia novaseelandiae* | 1.551 | 1.622 | 1.727 | 1.680 |
| 19. *Trichobilharzia physellae* | 1.647 | 1.642 | 1.785 | 1.699 |
| 20. *Trichobilharzia querquedulae* | 1.536 | 1.646 | 1.577 | 1.588 |
| 21. *Trichobilharzia regenti* | 1.566 | 1.466 | 1.662 | 1.581 |
| 22. *Trichobilharzia* sp. A | 1.619 | 1.713 | 1.647 | 1.648 |
| 23. *Trichobilharzia* sp. B | 1.596 | 1.675 | 1.649 | 1.675 |
| 24. *Trichobilharzia* sp. C | 1.681 | 1.691 | 1.697 | 1.717 |
| 25. *Trichobilharzia* sp. D | - | 1.456 | 1.663 | 1.514 |

*Continued on next page*

Table B13 – *Continued from previous page*

| | 1 | 2 | 3 | 4 | 5 | 6 | 7 | 8 | 9 | 10 | 11 |
|---|---|---|---|---|---|---|---|---|---|---|---|
| 26. *Trichobilharzia* sp. E | 83.744 | 83.744 | 82.759 | 80.542 | 84.236 | 84.834 | 85.204 | 80.419 | 79.433 | 88.300 | 82.266 |
| 27. *Trichobilharzia stagnicolae* | 85.961 | 85.714 | 83.251 | 82.512 | 85.222 | 85.944 | 84.711 | 82.389 | 79.680 | 86.330 | 82.759 |
| 28. *Trichobilharzia szidati* | 85.468 | 85.961 | 83.005 | 83.498 | 85.222 | 87.546 | 87.670 | 83.374 | 81.404 | 93.966 | 82.512 |

| | 12 | 13 | 14 | 15 | 16 | 17 | 18 | 19 | 20 | 21 | 22 | 23 | 24 |
|---|---|---|---|---|---|---|---|---|---|---|---|---|---|
| 26. *Trichobilharzia* sp. E | 81.592 | 79.064 | 77.833 | 83.251 | 84.729 | 86.453 | 85.961 | 85.714 | 86.667 | 88.424 | 85.468 | 85.961 | 85.714 |
| 27. *Trichobilharzia stagnicolae* | 81.343 | 80.049 | 79.557 | 83.005 | 83.744 | 86.207 | 86.946 | 85.222 | 87.654 | 87.685 | 86.700 | 86.207 | 86.700 |
| 28. *Trichobilharzia szidati* | 81.095 | 81.773 | 80.296 | 84.483 | 85.961 | 88.670 | 86.700 | 87.685 | 89.136 | 88.177 | 88.424 | 88.670 | 87.685 |

| | 25 | 26 | 27 | 28 |
|---|---|---|---|---|
| 26. *Trichobilharzia* sp. E | 88.916 | - | 1.492 | 1.514 |
| 27. *Trichobilharzia stagnicolae* | 87.931 | 88.670 | - | 1.656 |
| 28. *Trichobilharzia szidati* | 89.655 | 89.163 | 87.438 | - |

Table B14. Percent similarities of mammalian schistosomes. Standard error estimate(s) are shown above the diagonal and were obtained by a bootstrap procedure (500 replicates). Ambiguous positions were removed for each sequence pair. Values in parentheses indicate the number of representative sequences if > 1. Evolutionary analyses were conducted in MEGA (Tamura et al., 2021).

| | 1 | 2 | 3 | 4 | 5 | 6 |
|---|---|---|---|---|---|---|
| 1. BMC - *Schistosomatium douthitti* | - | 1.864 | 1.992 | 2.073 | 2.038 | 0.313 |
| 2. *Heterobilharzia americana* | 79.000 | - | 1.958 | 1.922 | 2.043 | 1.857 |
| 3. *Schistosoma bovis* (out) | 72.925 | 74.651 | - | 1.621 | 1.641 | 1.999 |
| 4. *Schistosoma indicum* | 71.739 | 76.248 | 83.826 | - | 1.761 | 2.056 |
| 5. *Schistosoma nasale* | 73.123 | 73.253 | 83.235 | 82.249 | - | 2.017 |
| 6. *Schistosomatium douthitti* (2) | 98.912 | 78.821 | 72.556 | 71.471 | 73.346 | - |

Table B15. Averaged percent similarities of the Strigeidae I. Standard error estimate(s) are shown above the diagonal and were obtained by a bootstrap procedure (500 replicates). Ambiguous positions were removed for each sequence pair. Evolutionary analyses were conducted in MEGA (Tamura et al., 2021).

| | 1 | 2 | 3 | 4 | 5 | 6 | 7 | 8 | 9 | 10 | 11 | 12 |
|---|---|---|---|---|---|---|---|---|---|---|---|---|
| 1. BMC - *Cotylurus* sp. A (2) | - | 1.639 | 1.594 | 1.694 | 1.624 | 1.621 | 1.751 | 1.771 | 1.614 | 1.581 | 0.217 | 1.613 |
| 2. BMC - *Cotylurus* sp. E (2) | 86.418 | - | 1.526 | 1.344 | 1.271 | 1.341 | 1.850 | 1.821 | 1.321 | 1.271 | 1.629 | 1.530 |
| 3. BMC - *Cotylurus* sp. B (2) | 86.637 | 87.144 | - | 1.513 | 1.511 | 1.615 | 1.830 | 1.720 | 1.602 | 1.553 | 1.580 | 0.270 |
| 4. BMC - *Cotylurus* sp. C (2) | 85.980 | 91.685 | 87.965 | - | 1.190 | 1.402 | 1.840 | 1.840 | 1.176 | 1.136 | 1.685 | 1.518 |
| 5. BMC - *Cotylurus* sp. F (2) | 86.418 | 92.888 | 87.309 | 92.998 | - | 1.359 | 1.819 | 1.818 | 0.908 | 1.006 | 1.614 | 1.523 |
| 6. BMC - *Cotylurus strigeoides* (2) | 85.950 | 91.447 | 84.978 | 91.009 | 90.680 | - | 1.785 | 1.701 | 1.327 | 1.363 | 1.607 | 1.626 |
| 7. *Cardiocephaloides medioconiger* | 82.694 | 82.932 | 83.260 | 83.151 | 82.495 | 82.457 | - | 1.527 | 1.768 | 1.711 | 1.749 | 1.820 |
| 8. *Cardiocephaloides physalis* | 83.790 | 82.385 | 84.136 | 82.713 | 82.495 | 83.553 | 87.965 | - | 1.761 | 1.843 | 1.790 | 1.739 |
| 9. *Cotylurus cornutus* | 87.076 | 92.341 | 85.996 | 93.217 | 95.842 | 91.119 | 83.151 | 83.370 | - | 0.916 | 1.613 | 1.612 |
| 10. *Cotylurus flabelliformis* | 87.295 | 92.670 | 86.871 | 93.873 | 95.624 | 91.119 | 84.464 | 82.713 | 96.061 | - | 1.580 | 1.559 |
| 11. *Cotylurus* sp. A | 99.781 | 86.652 | 86.871 | 86.214 | 86.652 | 86.185 | 82.713 | 83.589 | 87.090 | 87.309 | - | 1.599 |
| 12. *Cotylurus* sp. B | 86.418 | 86.871 | 99.562 | 87.746 | 87.090 | 84.759 | 83.151 | 83.807 | 85.777 | 86.652 | 86.652 | - |
| 13. *Cotylurus* sp. C | 85.980 | 91.685 | 87.965 | 100 | 92.998 | 91.009 | 83.151 | 82.713 | 93.217 | 93.873 | 86.214 | 87.746 |
| 14. *Cotylurus* sp. D | 87.295 | 91.247 | 85.996 | 92.341 | 90.810 | 93.641 | 84.026 | 83.151 | 91.028 | 91.685 | 87.527 | 85.558 |
| 15. *Cotylurus* sp. E | 86.637 | 99.234 | 87.746 | 92.341 | 92.998 | 91.776 | 83.589 | 82.713 | 92.341 | 92.998 | 86.871 | 87.527 |
| 16. *Cotylurus* sp. F | 86.418 | 92.888 | 87.309 | 92.998 | 100 | 90.680 | 82.495 | 82.495 | 95.842 | 95.624 | 86.652 | 87.090 |
| 17. *Cotylurus strigeoides* (2) | 85.761 | 91.575 | 84.902 | 91.138 | 90.591 | 99.561 | 82.276 | 83.151 | 90.810 | 91.028 | 85.996 | 84.683 |
| 18. *Ichthyocotylurus pileatus* | 84.193 | 86.404 | 83.333 | 84.430 | 84.649 | 84.945 | 83.991 | 82.895 | 85.307 | 85.746 | 84.211 | 83.114 |
| 19. *Ichthyocotylurus* sp. 2 | 85.291 | 87.610 | 85.746 | 85.526 | 83.772 | 85.055 | 83.991 | 84.430 | 84.868 | 85.088 | 85.088 | 85.307 |
| 20. *Ichthyocotylurus* sp. 3 | 85.104 | 85.996 | 84.683 | 86.652 | 86.214 | 86.184 | 85.120 | 84.683 | 86.652 | 86.433 | 85.120 | 84.464 |
| 21. *Tylodelphys scheuringi* (out) | 81.998 | 80.592 | 80.263 | 80.044 | 79.605 | 79.780 | 81.798 | 81.140 | 79.825 | 80.702 | 81.798 | 80.702 |

*Continued on next page*

Table B15 – *Continued from previous page*

| | 13 | 14 | 15 | 16 | 17 | 18 | 19 | 20 | 21 |
|---|---|---|---|---|---|---|---|---|---|
| 1. BMC - *Cotylurus* sp. A (2) | 1.694 | 1.530 | 1.628 | 1.624 | 1.632 | 1.765 | 1.623 | 1.699 | 1.729 |
| 2. BMC - *Cotylurus* sp. E (2) | 1.344 | 1.379 | 0.321 | 1.271 | 1.328 | 1.621 | 1.533 | 1.656 | 1.842 |
| 3. BMC - *Cotylurus* sp. B (2) | 1.513 | 1.565 | 1.521 | 1.511 | 1.626 | 1.796 | 1.600 | 1.680 | 1.910 |
| 4. BMC - *Cotylurus* sp. C (2) | 0.000 | 1.253 | 1.314 | 1.190 | 1.407 | 1.769 | 1.686 | 1.655 | 1.898 |
| 5. BMC - *Cotylurus* sp. F (2) | 1.190 | 1.430 | 1.312 | 0 | 1.365 | 1.745 | 1.763 | 1.648 | 1.953 |
| 6. BMC - *Cotylurus strigeoides* (2) | 1.402 | 1.135 | 1.341 | 1.359 | 0.227 | 1.689 | 1.734 | 1.606 | 1.920 |
| 7. *Cardiocephaloides medioconiger* | 1.840 | 1.733 | 1.839 | 1.819 | 1.801 | 1.710 | 1.750 | 1.709 | 1.795 |
| 8. *Cardiocephaloides physalis* | 1.840 | 1.772 | 1.840 | 1.818 | 1.729 | 1.858 | 1.777 | 1.735 | 1.835 |
| 9. *Cotylurus cornutus* | 1.176 | 1.421 | 1.351 | 0.908 | 1.354 | 1.692 | 1.714 | 1.629 | 1.916 |
| 10. *Cotylurus flabelliformis* | 1.136 | 1.348 | 1.266 | 1.006 | 1.379 | 1.645 | 1.660 | 1.639 | 1.882 |
| 11. *Cotylurus* sp. A | 1.685 | 1.515 | 1.619 | 1.614 | 1.617 | 1.762 | 1.644 | 1.698 | 1.752 |
| 12. *Cotylurus* sp. B | 1.518 | 1.597 | 1.524 | 1.523 | 1.636 | 1.802 | 1.620 | 1.679 | 1.906 |
| 13. *Cotylurus* sp. C | - | 1.253 | 1.314 | 1.190 | 1.407 | 1.769 | 1.686 | 1.655 | 1.898 |
| 14. *Cotylurus* sp. D | 92.341 | - | 1.362 | 1.430 | 1.133 | 1.711 | 1.668 | 1.673 | 1.907 |
| 15. *Cotylurus* sp. E | 92.341 | 91.685 | - | 1.312 | 1.331 | 1.612 | 1.525 | 1.624 | 1.856 |
| 16. *Cotylurus* sp. F | 92.998 | 90.810 | 92.998 | - | 1.365 | 1.745 | 1.763 | 1.648 | 1.953 |
| 17. *Cotylurus strigeoides* (2) | 91.138 | 93.654 | 91.904 | 90.591 | - | 1.681 | 1.724 | 1.596 | 1.910 |
| 18. *Ichthyocotylurus pileatus* | 84.430 | 84.868 | 86.842 | 84.649 | 85.197 | - | 1.370 | 1.421 | 1.808 |
| 19. *Ichthyocotylurus* sp. 2 | 85.526 | 84.868 | 87.939 | 83.772 | 85.197 | 90.570 | - | 1.450 | 1.726 |
| 20. *Ichthyocotylurus* sp. 3 | 86.652 | 85.777 | 86.652 | 86.214 | 86.324 | 90.351 | 90.570 | - | 1.686 |
| 21. *Tylodelphys scheuringi* (out) | 80.044 | 79.605 | 81.140 | 79.605 | 79.825 | 82.675 | 82.456 | 84.430 | - |

Table B16. Averaged percent similarities of the Strigeidae II. Standard error estimate(s) are shown above the diagonal and were obtained by a bootstrap procedure (500 replicates). Values in parentheses indicate the number of representative sequences if > 1. Ambiguous positions were removed for each sequence pair. Evolutionary analyses were conducted in MEGA (Tamura et al., 2021).

| | 1 | 2 | 3 | 4 | 5 | 6 | 7 | 8 | 9 | 10 | 11 | 12 |
|---|---|---|---|---|---|---|---|---|---|---|---|---|
| 1. *Apatemon* sp. '*jamiesoni*' | - | 1.637 | 1.547 | 1.552 | 1.789 | 1.993 | 2.001 | 1.969 | 1.929 | 1.921 | 2.000 | 1.950 |
| 2. *Apatemon* sp. A | 86.408 | - | 1.218 | 1.497 | 1.858 | 1.883 | 1.900 | 1.970 | 1.902 | 1.786 | 1.840 | 1.873 |
| 3. *Apatemon* sp. B | 87.864 | 93.220 | - | 1.398 | 1.783 | 1.813 | 1.835 | 1.906 | 1.810 | 1.705 | 1.810 | 1.828 |
| 4. *Apatemon* sp. C | 89.078 | 88.862 | 89.831 | - | 1.868 | 1.693 | 1.714 | 1.847 | 1.850 | 1.717 | 1.879 | 1.843 |
| 5. *Apharyngostrigea pipientis* (out) | 84.709 | 81.840 | 84.262 | 82.809 | - | 1.769 | 1.798 | 1.876 | 1.939 | 1.802 | 1.896 | 1.783 |
| 6. *Australapatemon burti* | 83.738 | 84.019 | 84.262 | 85.956 | 83.777 | - | 0.243 | 1.576 | 1.478 | 1.495 | 1.510 | 1.629 |
| 7. *Australapatemon burti* complex sp. Lin 1 | 83.495 | 83.777 | 84.019 | 85.714 | 83.535 | 99.758 | - | 1.600 | 1.506 | 1.527 | 1.546 | 1.583 |
| 8. *Australapatemon mclaughlini* | 83.495 | 82.324 | 83.051 | 84.746 | 83.051 | 89.588 | 89.346 | - | 1.269 | 1.486 | 1.586 | 1.522 |
| 9. *Australapatemon* sp. | 84.223 | 83.535 | 84.262 | 84.746 | 82.809 | 90.557 | 90.315 | 92.494 | - | 1.538 | 1.331 | 1.438 |
| 10. *Australapatemon* sp. LIN 3 | 82.524 | 83.293 | 85.230 | 85.714 | 82.809 | 90.557 | 90.315 | 89.104 | 88.620 | - | 1.519 | 1.545 |
| 11. *Australapatemon* sp. LIN 4 | 82.039 | 83.777 | 83.051 | 83.535 | 82.809 | 89.588 | 89.346 | 89.104 | 91.768 | 88.136 | - | 1.583 |
| 12. *Australapatemon* sp. LIN 5 | 81.311 | 82.567 | 83.051 | 84.504 | 83.051 | 88.136 | 88.378 | 89.346 | 90.557 | 87.651 | 88.136 | - |
| 13. *Australapatemon* sp. LIN 6 | 84.223 | 83.535 | 84.262 | 84.746 | 83.535 | 91.525 | 91.283 | 92.252 | 99.031 | 88.862 | 92.010 | 90.073 |
| 14. *Australapatemon* sp. LIN 8 | 84.951 | 85.230 | 85.472 | 84.262 | 85.714 | 89.346 | 89.104 | 86.441 | 88.620 | 87.167 | 88.136 | 84.746 |
| 15. *Australapatemon* sp. LIN 9A | 84.709 | 84.988 | 85.956 | 85.714 | 84.019 | 89.831 | 89.588 | 88.862 | 89.104 | 87.651 | 87.893 | 87.167 |
| 16. *Australapatemon* sp. LIN 9B | 84.223 | 84.988 | 85.714 | 87.893 | 82.567 | 89.831 | 89.588 | 89.588 | 89.588 | 89.831 | 87.893 | 88.620 |
| 17. *Australapatemon* sp. LIN 10 | 84.466 | 84.746 | 84.988 | 85.956 | 84.019 | 90.799 | 90.557 | 94.915 | 94.673 | 90.073 | 91.041 | 91.768 |
| 18. *Australapatemon* sp. | 84.223 | 83.535 | 84.262 | 84.746 | 82.809 | 90.557 | 90.315 | 92.494 | 100 | 88.620 | 91.768 | 90.557 |
| 19. BMC - *Australapatemon burti* (2) | 83.738 | 83.051 | 84.262 | 85.714 | 83.777 | 98.063 | 97.821 | 88.862 | 90.315 | 89.831 | 89.346 | 89.104 |
| 20. BMC - *Australapatemon burti* complex sp. Lin 1 (2) | 83.738 | 82.567 | 81.840 | 84.504 | 83.777 | 95.400 | 95.157 | 88.378 | 88.620 | 88.862 | 87.651 | 86.683 |
| 21. BMC - *Australapatemon* sp. Lin 9A (2) | 85.194 | 85.351 | 86.077 | 85.835 | 84.019 | 89.588 | 89.346 | 88.378 | 88.620 | 87.167 | 87.409 | 86.925 |
| 22. BMC - *Australapatemon* sp. (2) | 84.102 | 83.656 | 84.383 | 84.867 | 83.414 | 90.436 | 90.194 | 92.373 | 99.395 | 88.741 | 92.131 | 90.436 |
| 23. BMC - *Australapatemon mclaughlini* (2) | 83.495 | 82.446 | 83.172 | 84.746 | 83.172 | 89.588 | 89.346 | 99.758 | 92.494 | 89.104 | 89.346 | 89.588 |
| 24. BMC - *Australapatemon* sp. Lin 10 (2) | 84.223 | 84.504 | 84.746 | 85.472 | 83.777 | 90.557 | 90.315 | 95.400 | 94.915 | 90.315 | 90.799 | 92.252 |
| 25. BMC - *Australapatemon* sp. Lin 8 (2) | 84.709 | 84.988 | 85.230 | 84.019 | 85.714 | 89.104 | 88.862 | 86.199 | 88.620 | 86.925 | 88.136 | 84.504 |
| 26. BMC - *Australapatemon* sp. Lin 9C (2) | 84.102 | 83.898 | 84.746 | 84.625 | 82.567 | 88.741 | 88.499 | 87.530 | 88.499 | 88.741 | 88.741 | 87.772 |
| 27. BMC - *Australapatemon* sp. Lin 6 (2) | 84.102 | 83.656 | 84.625 | 84.867 | 82.930 | 90.799 | 90.557 | 92.494 | 99.516 | 88.862 | 91.768 | 90.799 |

*Continued on next page*

Table B16 – *Continued from previous page*

| | 13 | 14 | 15 | 16 | 17 | 18 | 19 | 20 | 21 | 22 | 23 |
|---|---|---|---|---|---|---|---|---|---|---|---|
| 1. *Apatemon* sp. '*jamiesoni*' | 1.951 | 1.886 | 1.915 | 1.858 | 1.888 | 1.929 | 1.978 | 1.999 | 1.860 | 1.912 | 1.946 |
| 2. *Apatemon* sp. A | 1.904 | 1.820 | 1.858 | 1.775 | 1.840 | 1.902 | 1.906 | 1.939 | 1.764 | 1.887 | 1.951 |
| 3. *Apatemon* sp. B | 1.830 | 1.753 | 1.751 | 1.725 | 1.751 | 1.810 | 1.821 | 1.961 | 1.685 | 1.776 | 1.885 |
| 4. *Apatemon* sp. C | 1.850 | 1.881 | 1.769 | 1.646 | 1.780 | 1.850 | 1.690 | 1.879 | 1.708 | 1.823 | 1.824 |
| 5. *Apharyngostrigea pipientis* (out) | 1.891 | 1.753 | 1.842 | 1.960 | 1.837 | 1.939 | 1.764 | 1.805 | 1.846 | 1.922 | 1.873 |
| 6. *Australapatemon burti* | 1.408 | 1.423 | 1.437 | 1.480 | 1.463 | 1.478 | 0.707 | 1.066 | 1.453 | 1.469 | 1.555 |
| 7. *Australapatemon burti* complex sp. Lin 1 | 1.436 | 1.450 | 1.464 | 1.514 | 1.493 | 1.506 | 0.758 | 1.086 | 1.476 | 1.501 | 1.580 |
| 8. *Australapatemon mclaughlini* | 1.264 | 1.629 | 1.488 | 1.517 | 1.065 | 1.269 | 1.597 | 1.666 | 1.536 | 1.259 | 0.172 |
| 9. *Australapatemon* sp. | 0.488 | 1.564 | 1.518 | 1.546 | 1.055 | 0.000 | 1.472 | 1.607 | 1.569 | 0.356 | 1.245 |
| 10. *Australapatemon* sp. LIN 3 | 1.517 | 1.617 | 1.508 | 1.484 | 1.413 | 1.538 | 1.486 | 1.565 | 1.540 | 1.501 | 1.470 |
| 11. *Australapatemon* sp. LIN 4 | 1.329 | 1.554 | 1.591 | 1.634 | 1.482 | 1.331 | 1.475 | 1.636 | 1.624 | 1.299 | 1.560 |
| 12. *Australapatemon* sp. LIN 5 | 1.464 | 1.762 | 1.616 | 1.600 | 1.373 | 1.438 | 1.571 | 1.700 | 1.652 | 1.428 | 1.502 |
| 13. *Australapatemon* sp. LIN 6 | - | 1.538 | 1.450 | 1.530 | 1.072 | 0.488 | 1.415 | 1.587 | 1.501 | 0.492 | 1.241 |
| 14. *Australapatemon* sp. LIN 8 | 88.862 | - | 1.431 | 1.502 | 1.556 | 1.564 | 1.446 | 1.445 | 1.455 | 1.560 | 1.608 |
| 15. *Australapatemon* sp. LIN 9A | 90.073 | 90.799 | - | 1.137 | 1.454 | 1.518 | 1.467 | 1.548 | 0.354 | 1.502 | 1.478 |
| 16. *Australapatemon* sp. LIN 9B | 89.588 | 89.104 | 93.947 | - | 1.465 | 1.546 | 1.504 | 1.617 | 1.129 | 1.532 | 1.503 |
| 17. *Australapatemon* sp. LIN 10 | 94.673 | 88.378 | 89.831 | 90.799 | - | 1.055 | 1.491 | 1.650 | 1.508 | 1.029 | 1.033 |
| 18. *Australapatemon* sp. | 99.031 | 88.620 | 89.104 | 89.588 | 94.673 | - | 1.472 | 1.607 | 1.569 | 0.356 | 1.245 |
| 19. BMC - *Australapatemon burti* (2) | 91.283 | 89.104 | 89.104 | 88.620 | 90.073 | 90.315 | - | 1.139 | 1.479 | 1.466 | 1.574 |
| 20. BMC - *Australapatemon burti* complex sp. Lin 1 (2) | 89.104 | 88.862 | 88.620 | 87.893 | 88.136 | 88.620 | 94.431 | - | 1.558 | 1.638 | 1.638 |
| 21. BMC - *Australapatemon* sp. Lin 9A (2) | 89.588 | 90.557 | 99.153 | 93.705 | 89.346 | 88.620 | 88.862 | 88.499 | - | 1.549 | 1.527 |
| 22. BMC - *Australapatemon* sp. (2) | 98.910 | 88.499 | 88.983 | 89.467 | 95.036 | 99.395 | 90.194 | 88.015 | 88.499 | - | 1.237 |
| 23. BMC - *Australapatemon mclaughlini* (2) | 92.252 | 86.562 | 88.862 | 89.588 | 94.915 | 92.494 | 88.862 | 88.378 | 88.378 | 92.373 | - |
| 24. BMC - *Australapatemon* sp. Lin 10 (2) | 94.915 | 88.136 | 89.831 | 91.283 | 99.516 | 94.915 | 89.831 | 87.893 | 89.346 | 95.278 | 95.400 |
| 25. BMC - *Australapatemon* sp. Lin 8 (2) | 88.862 | 99.637 | 90.678 | 88.983 | 88.136 | 88.620 | 88.862 | 88.620 | 90.436 | 88.499 | 86.320 |
| 26. BMC - *Australapatemon* sp. Lin 9C (2) | 88.983 | 88.257 | 91.404 | 92.615 | 89.467 | 88.499 | 88.741 | 87.046 | 90.920 | 88.378 | 87.772 |
| 27. BMC - *Australapatemon* sp. Lin 6 (2) | 99.031 | 88.136 | 89.104 | 89.588 | 94.673 | 99.516 | 90.799 | 88.378 | 88.620 | 99.395 | 92.494 |

*Continued on next page*

Table B16 – *Continued from previous page*

| | 24 | 25 | 26 | 27 |
|---|---|---|---|---|
| 1. *Apatemon* sp. '*jamiesoni*' | 1.888 | 1.902 | 1.839 | 1.926 |
| 2. *Apatemon* sp. A | 1.849 | 1.824 | 1.826 | 1.889 |
| 3. *Apatemon* sp. B | 1.754 | 1.759 | 1.793 | 1.794 |
| 4. *Apatemon* sp. C | 1.810 | 1.891 | 1.883 | 1.843 |
| 5. *Apharyngostrigea pipientis* (out) | 1.842 | 1.753 | 1.915 | 1.920 |
| 6. *Australapatemon burti* | 1.484 | 1.437 | 1.512 | 1.444 |
| 7. *Australapatemon burti* complex sp. Lin 1 | 1.513 | 1.464 | 1.537 | 1.477 |
| 8. *Australapatemon mclaughlini* | 1.008 | 1.637 | 1.555 | 1.254 |
| 9. *Australapatemon* sp. | 1.019 | 1.552 | 1.561 | 0.300 |
| 10. *Australapatemon* sp. LIN 3 | 1.391 | 1.624 | 1.535 | 1.500 |
| 11. *Australapatemon* sp. LIN 4 | 1.493 | 1.551 | 1.531 | 1.299 |
| 12. *Australapatemon* sp. LIN 5 | 1.335 | 1.764 | 1.613 | 1.422 |
| 13. *Australapatemon* sp. LIN 6 | 1.035 | 1.526 | 1.519 | 0.436 |
| 14. *Australapatemon* sp. LIN 8 | 1.557 | 0.220 | 1.562 | 1.566 |
| 15. *Australapatemon* sp. LIN 9A | 1.448 | 1.411 | 1.295 | 1.495 |
| 16. *Australapatemon* sp. LIN 9B | 1.422 | 1.491 | 1.284 | 1.542 |
| 17. *Australapatemon* sp. LIN 10 | 0.341 | 1.563 | 1.456 | 1.050 |
| 18. *Australapatemon* sp. | 1.019 | 1.552 | 1.561 | 0.300 |
| 19. BMC - *Australapatemon burti* (2) | 1.506 | 1.458 | 1.484 | 1.430 |
| 20. BMC - *Australapatemon burti* complex sp. Lin 1 (2) | 1.670 | 1.460 | 1.683 | 1.609 |
| 21. BMC - *Australapatemon* sp. Lin 9A (2) | 1.499 | 1.436 | 1.340 | 1.541 |
| 22. BMC - *Australapatemon* sp. (2) | 0.992 | 1.548 | 1.539 | 0.310 |
| 23. BMC - *Australapatemon mclaughlini* (2) | 0.982 | 1.616 | 1.535 | 1.232 |
| 24. BMC - *Australapatemon* sp. Lin 10 (2) | - | 1.564 | 1.444 | 1.007 |
| 25. BMC - *Australapatemon* sp. Lin 8 (2) | 87.893 | - | 1.557 | 1.553 |
| 26. BMC - *Australapatemon* sp. Lin 9C (2) | 89.709 | 88.136 | - | 1.533 |
| 27. BMC - *Australapatemon* sp. Lin 6 (2) | 94.915 | 88.136 | 88.499 | - |
